# Accurate detection of circulating tumor DNA using nanopore consensus sequencing

**DOI:** 10.1101/2020.07.14.202010

**Authors:** Alessio Marcozzi, Myrthe Jager, Martin Elferink, Roy Straver, Joost H. van Ginkel, Boris Peltenburg, Li-Ting Chen, Ivo Renkens, Joyce van Kuik, Chris Terhaard, Remco de Bree, Lot A. Devriese, Stefan M. Willems, Wigard P. Kloosterman, Jeroen de Ridder

**Affiliations:** Center for Molecular Medicine and Oncode Institute, University Medical Center Utrecht, Utrecht University, Heidelberglaan 100, 3584 CX Utrecht, The Netherlands; Cyclomics, Universiteitsweg 100, 3584 CG Utrecht, The Netherlands; Department of Genetics, University Medical Center Utrecht, Utrecht University, Heidelberglaan 100, 3584 CX Utrecht, The Netherlands; Department of pathology, University Medical Center Utrecht, Utrecht University, Heidelberglaan 100, 3584 CX Utrecht, The Netherlands; Department of Oral and Maxillofacial Surgery, University Medical Center Utrecht, Utrecht University, Heidelberglaan 100, 3584 CX Utrecht, The Netherlands; Department of Radiotherapy, UMC Utrecht Cancer Center, University Medical Center Utrecht, Utrecht, The Netherlands; Department of Head and Neck Surgical Oncology, UMC Utrecht Cancer Center, University Medical Center Utrecht, Utrecht, The Netherlands; Department of Medical Oncology, UMC Utrecht Cancer Center, University Medical Center Utrecht, Utrecht, The Netherlands; Department of Pathology and Medical Biology, University Medical Center Groningen, Rijksuniversiteit Groningen, Hanzeplein 1, 9713 GZ Groningen, The Netherlands; Center for Molecular Medicine, University Medical Center Utrecht, Utrecht University, Heidelberglaan 100, 3584 CX Utrecht, The Netherlands

## Abstract

Levels of circulating tumor DNA (ctDNA) in liquid biopsies may serve as a sensitive biomarker for real-time, minimally-invasive tumor diagnostics and monitoring. However, detecting ctDNA is challenging, as much fewer than 5% of the cell-free DNA in the blood typically originates from the tumor. To detect lowly abundant ctDNA molecules based on somatic variants, extremely sensitive sequencing methods are required. Here, we describe a new technique, CyclomicsSeq, which is based on Oxford Nanopore sequencing of concatenated copies of a single DNA molecule. Consensus calling of the DNA copies increased the base-calling accuracy ∼60x, enabling accurate detection of *TP53* mutations at frequencies down to 0.02%. We demonstrate that a *TP53*-specific CyclomicsSeq assay can be successfully used to monitor tumor burden during treatment for head-and-neck cancer patients. CyclomicsSeq can be applied to any genomic locus and offers an accurate diagnostic liquid biopsy approach that can be implemented in point-of-care clinical workflows.

## INTRODUCTION

Solid tumors constantly shed small DNA molecules into the bloodstream, which are cleared within a few hours^1,2^. Determining the circulating tumor DNA (ctDNA) content in the blood of cancer patients offers a unique opportunity for real-time detection and monitoring of solid tumors^3,4^, as levels of these ctDNA molecules are associated with tumor presence, tumor type, tumor size, tumor stage, prognosis, response to therapy, and recurrent disease^2,5–9^. Furthermore, obtaining blood from a patient is minimally-invasive and therefore, in contrast to biopsies of solid tumors, more suited to generate serial measurements of the tumor within the same patient. Moreover, tumor locations (primary tumors or metastases) are not always easily accessible for taking biopsies and complications can occur. In this context, it has been shown that ctDNA detection in blood and other fluids (“liquid biopsies”) is complementary to solid biopsies for detection of targets for precision medicine^10^.

The presence of somatic mutations in cell-free DNA (cfDNA) molecules is commonly used to approximate ctDNA content^8,11^. However, detection of ctDNA is challenging, since noncancerous cells also shed cfDNA into the blood. The fraction of tumor-derived molecules in the blood is typically much lower than 5% and fractions as low as 0.1% have been observed^5,12,13^. Therefore, a diagnostic ctDNA assay must be fast and cheap as well as highly sensitive. ctDNA can be detected with good sensitivity by digital droplet PCR (ddPCR), but this technique requires quite some time since it can typically only interrogate a single locus per assay and variants must be known *a priori*^2,14,15^. Alternatively, next-generation sequencing (NGS) approaches are used, but these suffer from a lower sensitivity and require highly optimized lab workflows to become cost-effective^9,16^.

Oxford Nanopore Technology (ONT) recently emerged as a powerful sequencing platform that offers advantages in terms of speed (real-time sequencing), cost-efficiency (low capital investment), and flexibility (distributed sequencing instead of centralized sequencing)^17^. ONT sequencing could, therefore, be very relevant for rapid and point-of-care clinical liquid biopsy testing. There are, however, two important limitations for ONT sequencing that hamper its use in a clinical setting. Firstly, current protocols are optimized for long DNA molecules. The shortest fragment sequenced on this platform to date is ∼425 bp, which is much longer than the average 145bp ctDNA^18,19^. Secondly, the basal error rate is ∼5-10%, which is too high to reliably detect ctDNA^20,21^. Several studies have shown that reading the same molecule multiple times can reduce the sequencing error rate^22–25^. However, some of these methods can only detect ctDNA fractions of >10%^25^, while others rely on self-circularization which is not possible for short ctDNA molecules^26^.

Here, we present a new technique, called CyclomicsSeq, that utilizes circularization and concatemerization of short DNA molecules and an optimized DNA backbone sequence in combination with ONT sequencing. As proof of concept, we developed a *TP53*-specific CyclomicsSeq protocol and a dedicated software pipeline to determine the mutation burden of a series of cfDNA samples obtained from liquid biopsies from patients with Human Papilloma Virus (HPV) negative head-and-neck squamous cell carcinoma (HNSCC). *TP53* is the most commonly mutated tumor suppressor gene in human cancer and therefore serves as a widely applicable target for cancer monitoring based on liquid biopsies^27,28^. There are relatively few hotspot mutations^29^, making this gene especially suitable for NGS-based approaches. The application to HPV-negative HNSCC is motivated by the fact that five-year survival rates are relatively low and substantial treatment benefits may be obtained by early diagnosis of recurrent disease and/or treatment response^30–32^. Moreover, differentiation between residual or recurrent tumor and radiation effects is often difficult during response evaluation or in case of suspicion of recurrency, even using modern imaging techniques. Approximately 90% of the HPV-negative HNSCC patients have a somatic mutation in *TP53^33^*. These *TP53* mutations occur early in the tumorigenesis of HNSCC and as such are present in (virtually) all tumor cells including subclones that metastasize^31,34^. For this reason, the detection of mutated *TP53* ctDNA molecules in liquid biopsies is suggested to be an ideal biomarker for HNSCC^14,35^.

We demonstrate that CyclomicsSeq leads to highly accurate consensus sequences, suitable for mutation detection at single-molecule resolution. Longitudinal liquid biopsy testing using CyclomicsSeq correctly identifies the presence and absence of ctDNA content, which could be informative for the management of HNSCC patients. CyclomicsSeq can be applied to a single or multiple genomic regions of choice, in principle, thereby representing a new liquid biopsy test that is relevant for diagnostic monitoring of any solid tumor for which ctDNA is a suitable biomarker.

## RESULTS

### CyclomicsSeq generates long concatemers

CyclomicsSeq is a protocol designed to produce and sequence long (>1Kb) DNA concatemers with a linear repetition of a sequence of interest called “insert”, and a DNA adaptor, referred to as “backbone”. The molecular protocol of CyclomicsSeq is divided into four main steps: 1) circularization of insert and backbone, 2) rolling circle amplification (RCA), 3) long-read sequencing, and 4) data processing (Fig. 1a-b; Supplementary Fig. 1). In step 4, the long reads are split based on the backbone and insert sequences and individual copies are extracted. Based on these individual copies, a consensus sequence is constructed for the backbone and insert separately. The backbones are optimized for e.g. flexibility while retaining a short length of around ∼250 bp (Methods; Supplementary Fig. 2, Supplementary Table S1). They serve as a molecular adaptor to mediate the circularization of the insert and are used to split and filter the reads during the data processing step. Backbones also includes barcodes and restriction sites utilized for quality control of the concatemers before sequencing (Methods; Supplementary Fig. 1). The inserts can be, in principle, any double-stranded DNA fragment. We have tested the method with inserts ranging from 90 to 700 bp. In this study, short (<200bp) PCR amplicons from the *TP53* gene amplified from (cf)DNA were used.

**Figure 1.**
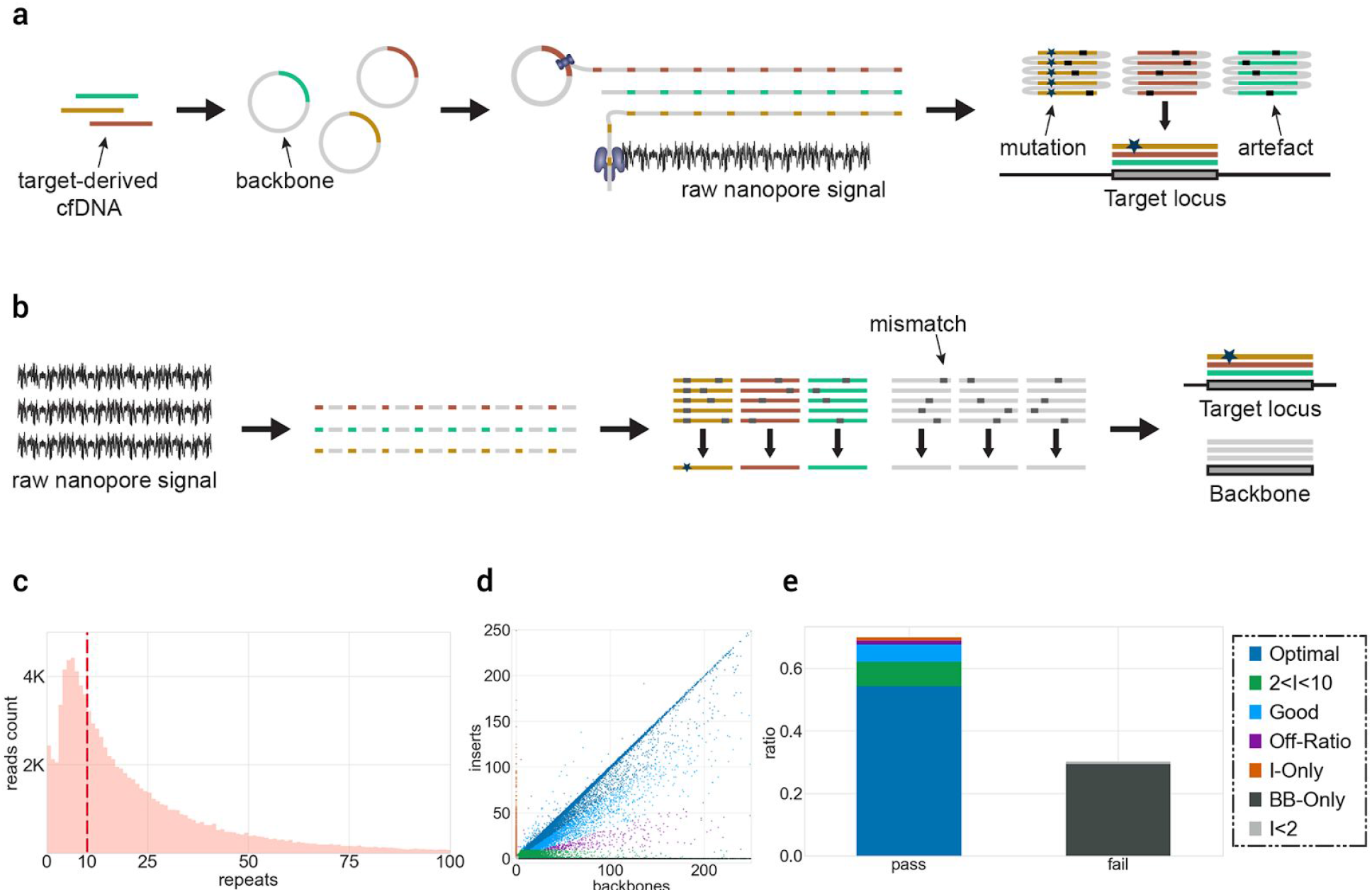
CyclomicsSeq protocol. **a** Experimental setup of CyclomicsSeq. PCR-amplified (‘target-derived) cfDNA is circularized with an optimized DNA backbone. Rolling circle amplification generates a long DNA molecule with alternating insert and backbone sequences, which is sequenced using ONT sequencing. Consensus calling of the DNA sequence allows discrimination between mutations and sequencing artifacts. **b** Schematic overview of the bioinformatic pipeline. **c** Distribution of insert copies versus the number of reads for a representative CyclomicsSeq run (**#CY_SM_PC_HN_0002_001_000**). **d** Ratio of insert versus backbone for CyclomicsSeq reads for a representative CyclomicsSeq run (**#CY_SM_PC_HN_0002_001_000**). Each read is represented by a data point (dot). Colours, noted in the legend, represent the different categories a read can belong to. Optimal CyclomicsSeq reads result from a one to one ratio of insert and backbone copies and contain at least 10 repeats (Blue). The other categories include: reads with fewer repeats (Green), reads without a backbone (Orange), reads without the target insert (Gray). Reads with BB:I ratios between 0.35 and 3 are defined as “Good” (Cyan), while the others are classified as “Off-ratio” (Purple). **e** Ratio of sequencing data grouped by read type for a representative CyclomicsSeq run (**#CY_SM_PC_HN_0002_001_000**). In this case, more than 60% of the data was used to generate consensus reads. The remnant data was discarded because it contained backbone-only sequences.

As a proof-of-principle, we performed a CyclomicsSeq test with a *TP53* insert and backbone BB24 (Methods; Sample CY_SM_PC_HN_0002_001_000; Supplementary Table S2-S3), and sequenced the resulting concatemeric DNA molecules on a Nanopore MinION instrument. This MinION run (Fig. 1c-e) yielded 7.2Gb of data, with reads containing concatemers of up to 250 repeats and an average number of concatemers of 24 repeats (Fig. 1c). The majority of the data (70%) consisted of concatemers with alternating backbone and insert sequences (Fig. 1d). The main byproducts of the RCA reaction were backbone-only concatemers (30%) that are filtered out during data processing (Fig. 1e).

In addition to the single amplicon used in the above pilot test, we tested whether CyclomicsSeq can be paired with amplicon panels covering multiple genomic loci and entire coding regions of genes. As an example, a multiplex PCR method was used to amplify all the *TP53* exons from cfDNA (Sample CY_SM_PC_HC_0004_003; Supplementary Data, Supplementary Table S3). Consensus reads spanned across all *TP53* exons, with a relatively even distribution of the coverage across all exons (Fig. 2a). Using a single MinION workflow, we obtained a coverage >1,000X for the vast majority (74%) of the exonic bases of *TP53* (Fig, 2b).

**Figure 2.**
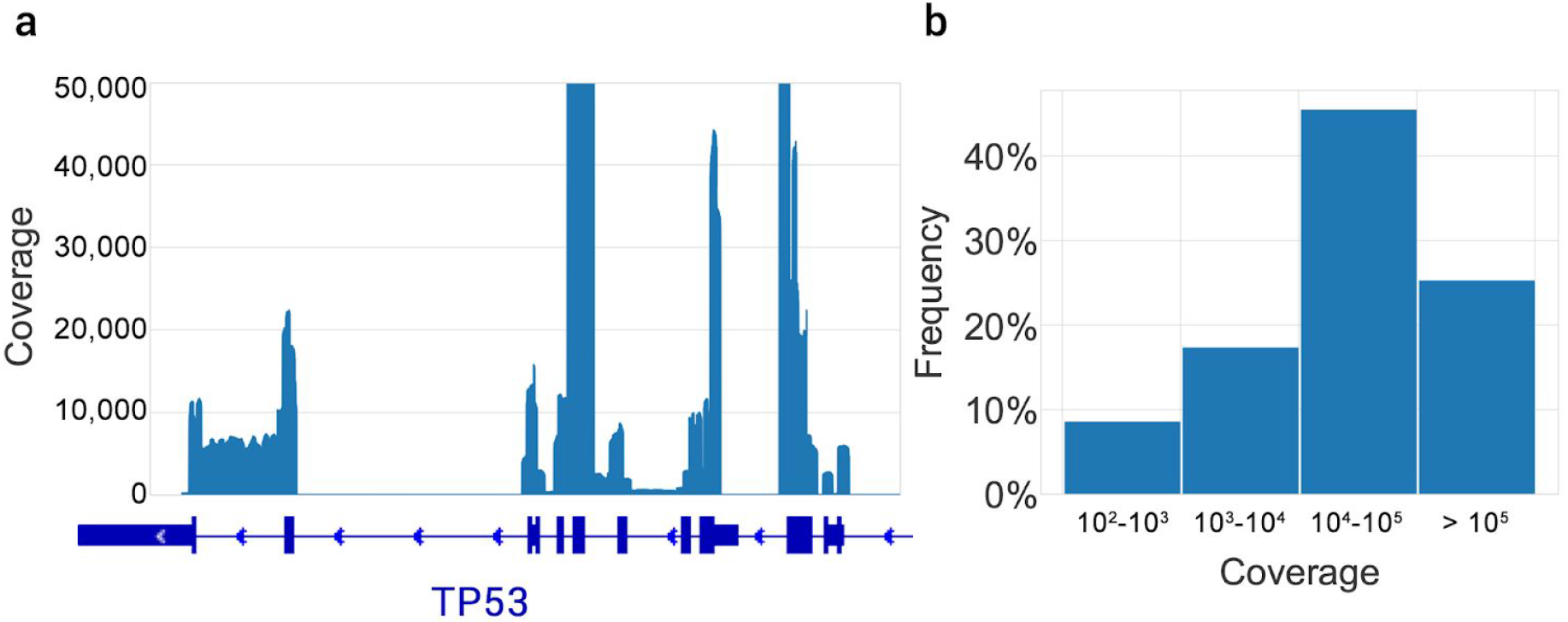
Coverage of consensus reads across the *TP53* gene. **a** Coverage profile, aligned with a schematic representation of the *TP53* gene. The bold blue boxes indicate the exons. The Y-axis was limited to 50,000X coverage. **b** Frequency of coverage grouped by intervals.

To evaluate person-, time- and sequencing-dependent variability in CyclomicsSeq results, CyclomicsSeq was performed three times by two different operators (only one of which had experience with the protocol) on two different days using the same insert and backbone, and subsequently each CyclomicsSeq product was sequenced on two separate MinION flow cells (Supplementary Fig. 3, Supplementary Table S2-S3). The insert used for these experiments was a mixture of four versions of a 151bp synthetic insert with 0 - 4 mutations across the insert. In total, between 12,242 and 125,446 reads were obtained. The ratio of PASS and FAIL reads and the read length distribution were highly similar between runs, although there is some inter-individual difference (Supplementary Fig. 3). Nevertheless, the observed ratios of the four inserts were highly similar as well, indicating that CyclomicSeq provides reproducible results (Supplementary Fig. 3).

### Consensus calling improves the accuracy

To evaluate the effect of CyclomicsSeq consensus calling on Nanopore sequencing accuracy, we performed 19 CyclomicsSeq experiments with three different backbone sequences, one backbone for each experiment (Supplementary Data, Supplementary Table S2-S3). In total, between 0.86 and 11.9 million (mean 3.09 million) sequencing reads were obtained (Supplementary Table S3). For each experiment, we determined the false positive rate of single-nucleotide errors (snFP rate) in the consensus backbone sequences as a function of the number of copies of the backbone in a read (Fig. 3a). For reads with a single copy of the backbone, the mean snFP rate was 0.0184 (minimum and maximum values were 0.0166 - 0.0210) (Fig. 3a). Consensus calling reduced the snFP rate to 0.0038 (0.0028 - 0.0057) for 5 and 0.0016 (0.001 - 0.0024) for 10 repeats (Fig. 3a). The snFP rate did not decrease substantially after ∼10 repeats. Similar to false positive single-nucleotide errors, the number of short deletions decreased with an increased number of repeats. This reduction plateaus after ∼10 repeats (Supplementary Fig. 4). This indicates that applying a threshold of at least 10 repeats for consensus calling will result in accurate mutation calls without unnecessary loss of data. Using this threshold, the mean false positive rate for single-nucleotide errors was 5.10^−4^ (2.10^−4^ - 6.10^−4^) in the backbone sequences.

**Figure 3.**
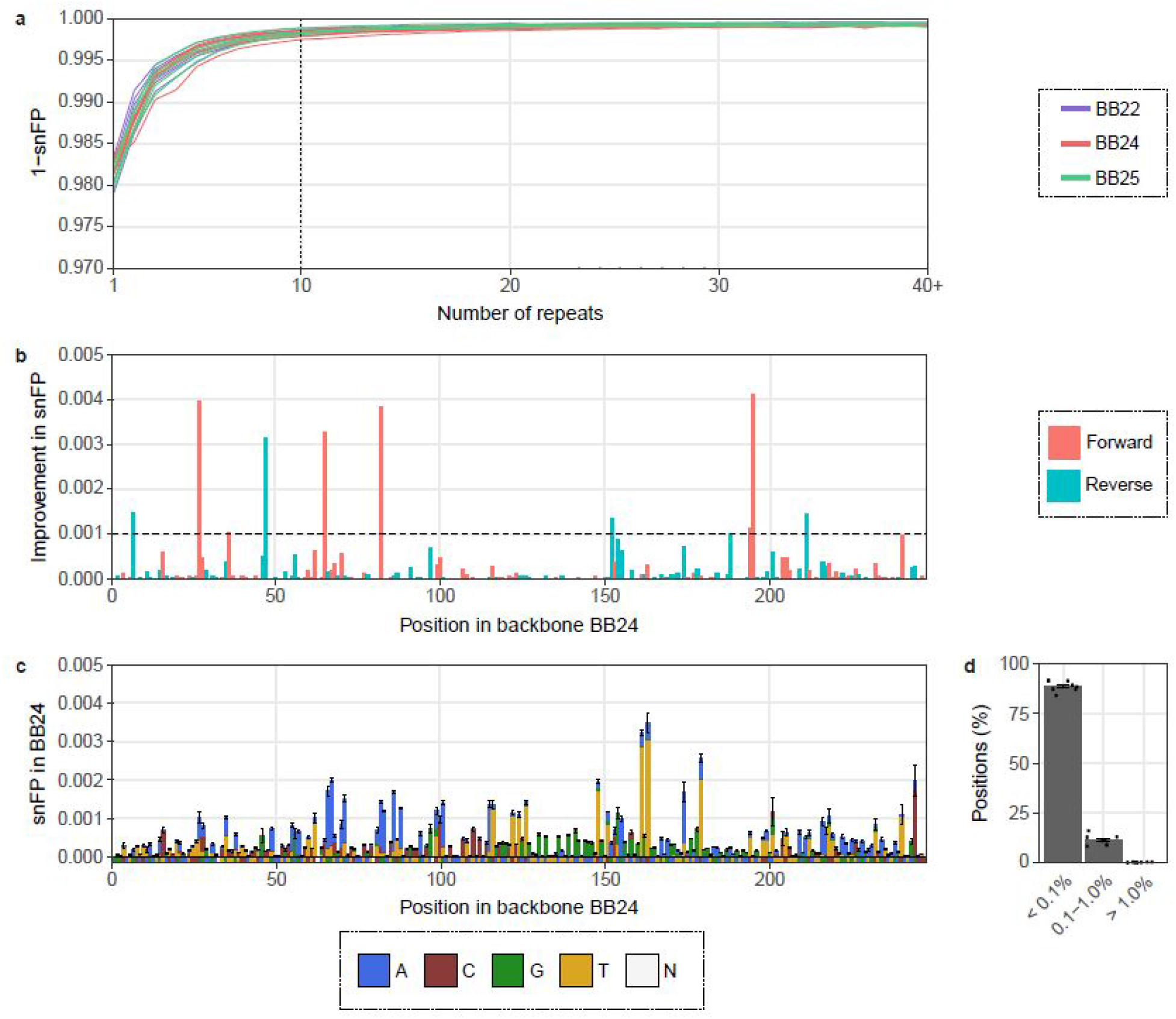
Consensus calling increases the accuracy of base calling in the backbone. **a** 1 - single-nucleotide false positive (snFP) rate in backbones (BB22, BB24, and BB25) per number of repeats. The dashed line indicates 10 repeats. Colours represent backbone type. **b** Improvement in snFP rate in BB24 if only the forward or the reverse reads are taken into account. The dashed line indicates an improvement of 0.1%. Colours represent read orientation. **c** Mean snFP rate across BB24 in reads with at least 10 repeats. Reference sequence is depicted below the x-axis. Colours represent base type. N = any. **d** Percentage of positions in BB24 with indicated snFP percentage. Data points represent individual sequencing runs. 6 BB22, 8 BB24, and 5 BB25 runs were used for the calculations. Error bars indicate the standard deviation (sd).

Although 91.9% of the positions in the backbone sequences had an snFP rate below 0.001, some positions had an snFP rate exceeding 0.004 (Supplementary Fig. 5). This suggests that there were non-random sequencing errors in the sequencing data that cannot be resolved by standard consensus calling. Non-random sequencing errors can depend on the sequence context and, therefore, considering only reads with a forward or a reverse orientation for some positions might reduce these non-random errors. Indeed, the snFP rate could be further improved by at least 0.1% at 11 of the 243 positions in BB24 by considering only forward or reverse reads for those positions (Fig. 3b). This especially reduced the number of false positives at positions with a high snFP rate. The improvement was consistent between sequencing runs, confirming the non-randomness of errors at these positions (Supplementary Fig. 5). After correction for forward or reverse orientation, 92.7% of the positions had a mean snFP rate < 0.001 in consensus called reads with at least 10 repeats of the insert, and 0% of the positions had an snFP rate >0.01 (Fig. 3c-d, Supplementary Fig. 6). Furthermore, only 2.1% of the positions had a combined snFP and deletion rate >0.01 (Supplementary Fig. 4). Using both the threshold for consensus calling (at least 10 repeats) and the forward/reverse orientation correction, the snFP rate was 3.10^−4^ (minimum and maximum values were 2.10^−4^ - 5.10^−4^) in the backbone sequences. CyclomicsSeq thus lowers the sequence error rate of ONT sequencing by ∼60x, which is a rate compatible with mutation frequencies in circulating DNA of cancer patients^5,12,13^.

Recently, ONT released the Flongle flow cell with R9-like pores and reusable parts^36^. Although Flongle flow cells have 4 times fewer pores, the flow cell is more cost-efficient and, therefore, may be more suitable for diagnostic approaches. Eight samples were sequenced using a Flongle flow cell to determine base calling accuracy after consensus calling (Supplementary Data, Supplementary Table S2-S3). Similar to the normal R9 flow cell, the snFP rate decreases through consensus calling in the Flongle flow cell and this plateaus at ∼10 repeats (Supplementary Fig. 7). Furthermore, the snFP profile across the backbone showed similar features between Flongle and R9 flow cells (BB25; Supplementary Fig. 6, Supplementary Fig. 7). We observed that the snFP rate of backbone sequences was ∼1.4x higher. This confirmed that the Flongle may be a cost-effective alternative for the R9 flow cell for some diagnostic approaches.

ONT also released a beta version of a new flow cell with a higher accuracy in March 2019^37^. Two other samples were tested on this R10 flow cell (Supplementary Data, Supplementary Table S2-S3). Similarly as observed before, the snFP rate and deletion rate decreased through consensus calling and reached a plateau at ∼10 repeats (Supplementary Fig. 8). However, the mean error rate was ∼1.3x higher (determined from the backbone; Supplementary Fig. 8) and in comparison to the R9 flow cell, the error profile was very different. Therefore, R10 flow cells may provide a valuable alternative to R9 flow cells if the oncogenic mutations occur at any of the few positions with a relatively high snFP rate in R9.

### Detection of COSMIC mutations in*TP53*

To evaluate the use of CyclomicsSeq for detection of cancer mutations in liquid biopsies from cancer patients, we focused on sequencing the *TP53* gene in cfDNA. In a first experiment, we estimated the false positive rate for detection of known *TP53* mutations as catalogued in the COSMIC database^38^ in four sequencing runs based on a *TP53* amplicon covering one, multiple or all *TP53* exons, amplified from control cfDNA samples from individuals without cancer (Fig. 4). For all four runs, the median snFP rate was less than 6.10^−4^ across the *TP53* exon(s). For ∼90% of the COSMIC mutations the snFP rate was lower than 1.10^−3^ and between 20% and 30% of all COSMIC bases have a snFP rate lower than 1.10^−4^.

**Figure 4.**
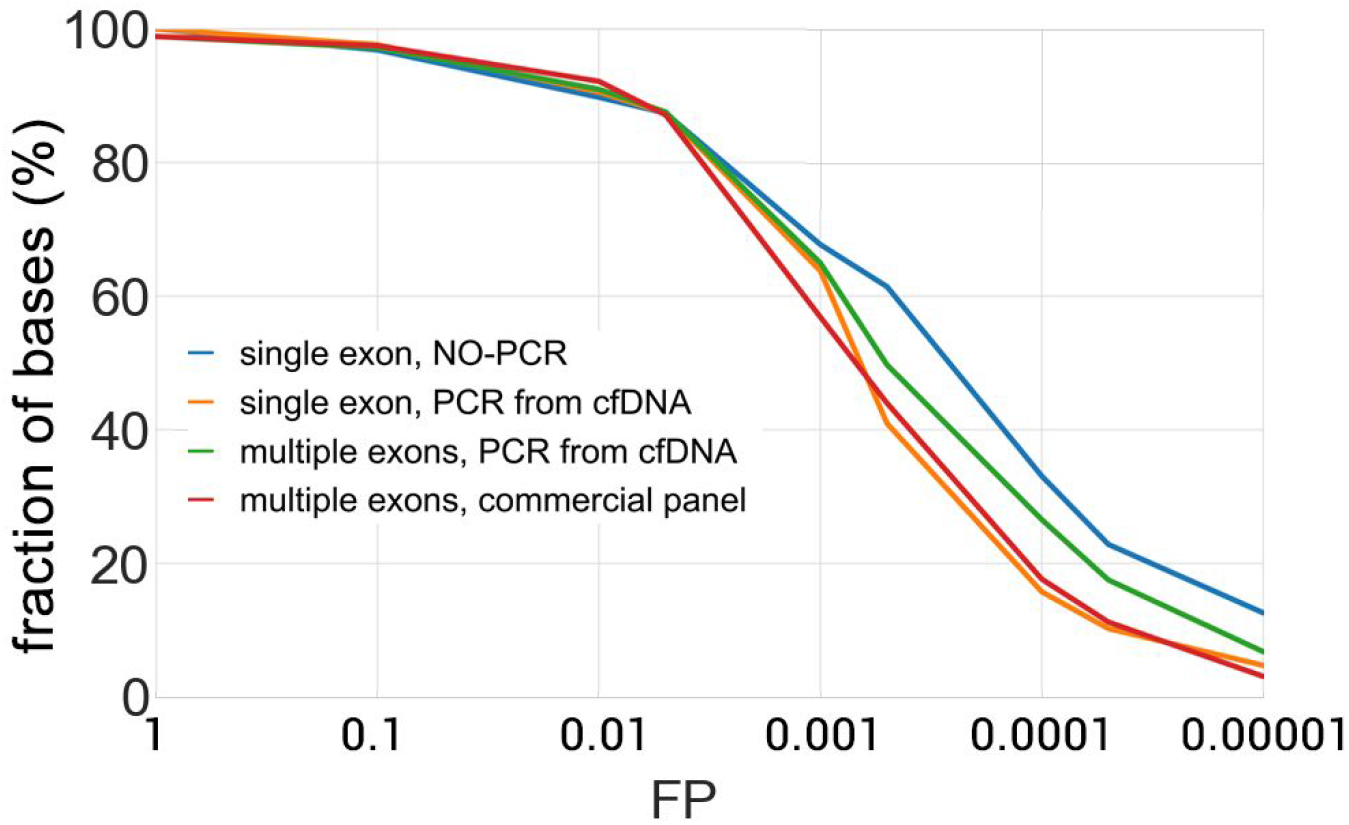
False positive rate for COSMIC mutations in *TP53*. False positive (FP) rate was obtained for four representative runs: two runs based on a single *TP53* exon, one run covering multiple *TP53* exons, and one run covering all *TP53* exons. For each COSMIC position in the target insert, the FP was calculated. The y-axis represents the percentage of bases having a snFP value lower than the FP indicated at the corresponding X-value. Blue is CY_PJET_12WT_0001_000, yellow is CY_SS_PC_HC_0001_001_000, green is CY_SM_PC_HC_0002_001_000, and red is CY_SM_PC_HC_0004_001_000.

Next, we aimed to test CyclomicsSeq in a situation which mimics low ctDNA amounts in the blood. To this end, we generated a 141 bp (17:7577010-7577150 in GRCh37, covering a *TP53* exon) synthetic ‘WT’ molecule without mutations and a ‘MUT’ insert of the same genomic locus with three cancer hotspot mutations in *TP53.* Both samples were mixed to obtain a low-abundant mutant sample of 99.9% WT and 0.1% MUT molecules (‘WT/MUT’; Supplementary Fig. 9). To create RCA template, the WT, MUT and mix of WT and MUT molecules were cloned into pJET, instead of the CyclomicsSeq backbone, as this allows amplification of the insert by replication in *E. coli*, thus preventing accumulation of errors due to PCR (Supplementary Fig. 9). In total, between 2.5 and 3.9 million sequencing reads were obtained for WT, MUT and the mixed WT/MUT sample (Supplementary Data, Supplementary Table S3). Because pJET is ∼10x longer than the backbone, a threshold of at least 5 repeats was applied during consensus calling. Even so, only 7.8 - 10.9% of these reads contained enough copies of the insert and were useful for data analysis. We found that for molecules with only one insert (i.e. without consensus calling) the snFP rate was ∼1.08x lower compared to inserts amplified with PCR (Supplementary Fig. 9). This indicates that the PCR used to amplify insert prior to CyclomicsSeq introduces errors. In the consensus called reads (i.e. in molecules with at least 5 repeats of the insert), the snFP rate was ∼1.26x lower compared to inserts that underwent a PCR step (Fig. 5a). A PCR-free approach can thus improve the results obtained by CyclomicsSeq even further, at the cost of sequencing depth, simplicity of the protocol and sample processing time.

**Figure 5.**
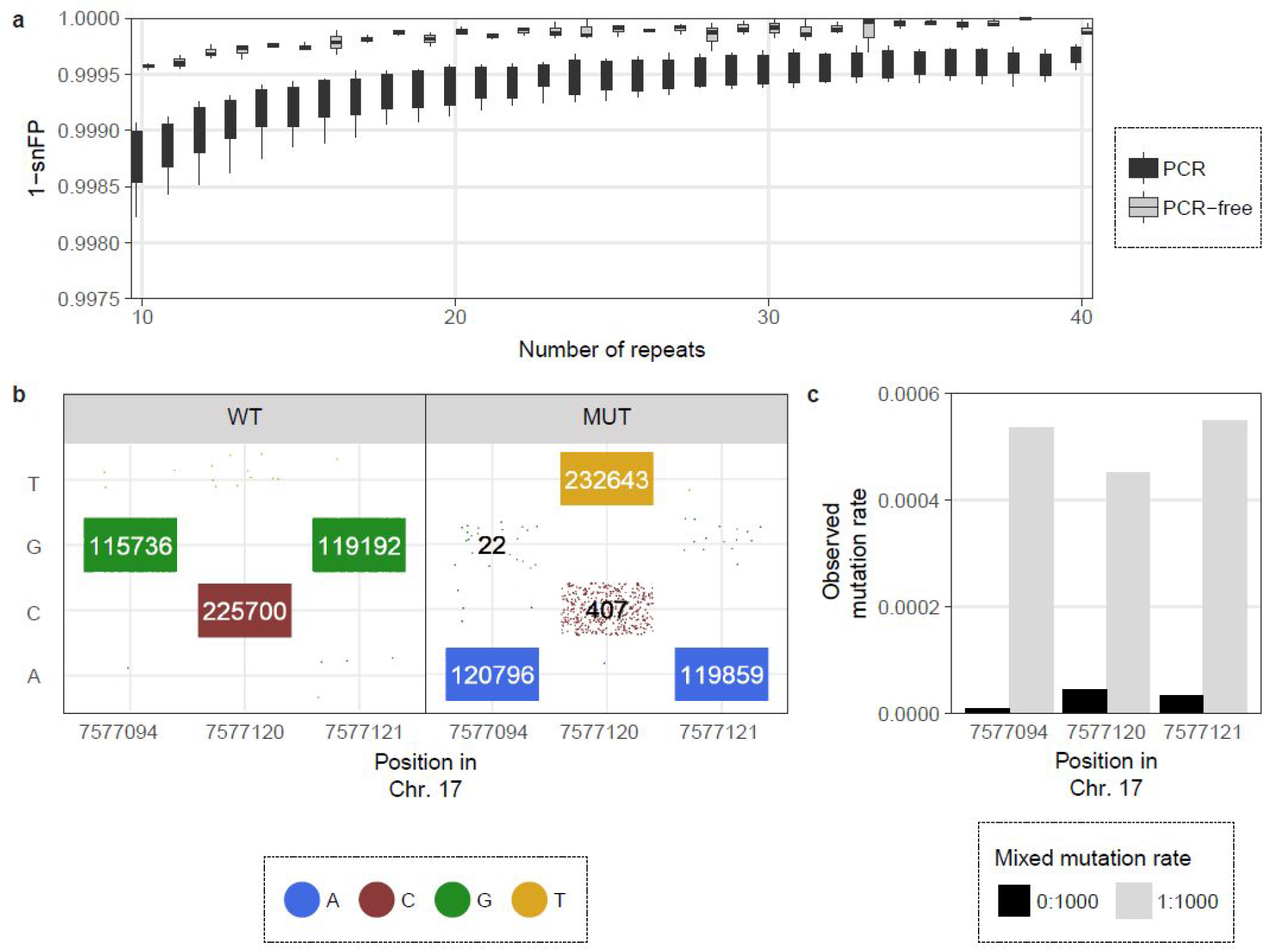
Detecting mutations in a synthetic *TP53* exon using CyclomicsSeq. **a** Box plots (center line = median; box limits = 25th and 75th percentiles; whiskers = 1.5x interquartile range; data points = outliers) depicting 1 - single-nucleotide false positive (snFP) rate in the insert (17:7577010-7577150 in GRCh37) per number of repeats for 8 PCR and 3 PCR-free inserts. **b** Calls in WT and MUT at the three mutant positions in *TP53*. Data points represent single reads. Colours indicate base call in the consensus-called read. Numbers >20 reads are shown, of which true negatives and true positives are indicated in white. **c** Observed mutation rate in WT and mixed WT/MUT at the three mutant positions in *TP53*. Expected mutation rates are 0.000 in WT and 0.001 in mixed WT/MUT.

In the MUT sample, 0.018%, 0.17%, and 0.013% of the reads contained a false positive WT call at the three assessed positions, respectively (Fig. 5b). Furthermore, 99.9%, 99.5% and 99.9% of the reads contained true positive mutation calls in the MUT sample, at the three assessed positions (Fig. 5b). The three synthetic mutations were observed in less than 0.004% of the reads in the WT sample (Fig. 5b). In the mixed WT/MUT, the observed ctDNA fraction was notably higher than in the WT sample for all three positions (Fig. 5c). These experiments confirm that CyclomicsSeq can be used to accurately detect low amounts of mutated ctDNA in the blood.

### CyclomicsSeq enables detection of ctDNA

To confirm whether CyclomicsSeq can be used to detect mutated ctDNA in the blood of patients, we focused on HPV-negative HNSCC patients, because 90% of these tumors contain *TP53* mutations^33^. We isolated cfDNA from the blood of three advanced stage HPV-negative HNSCC patients (denoted as patient A, B and C) before, during and after treatment (2 - 6 time points per patient) and performed CyclomicsSeq on each sample (Fig. 6a-c). Each patient’s HNSCC tumor contained a known *TP53* mutation, as determined by sequencing of tumor tissue (Supplementary Table S2). All three patients received daily radiotherapy treatment for five to seven weeks. In addition, patient A (multiple doses of cisplatin) and B (1 dose of cisplatin & carboplatin) also received concomitant chemotherapy treatment (‘chemoradiation’). The presence/absence of *TP53* mutations in ctDNA derived from the liquid biopsies was confirmed using ddPCR with primers designed to target the variant observed from performing NGS on the corresponding solid tumor biopsy of each patient (Fig. 6d-f). Furthermore, Magnetic Resonance Imaging (MRI) scans of patient A and B were also available to assess gross tumor volume (GTV; Fig. 6g-h).

**Figure 6.**
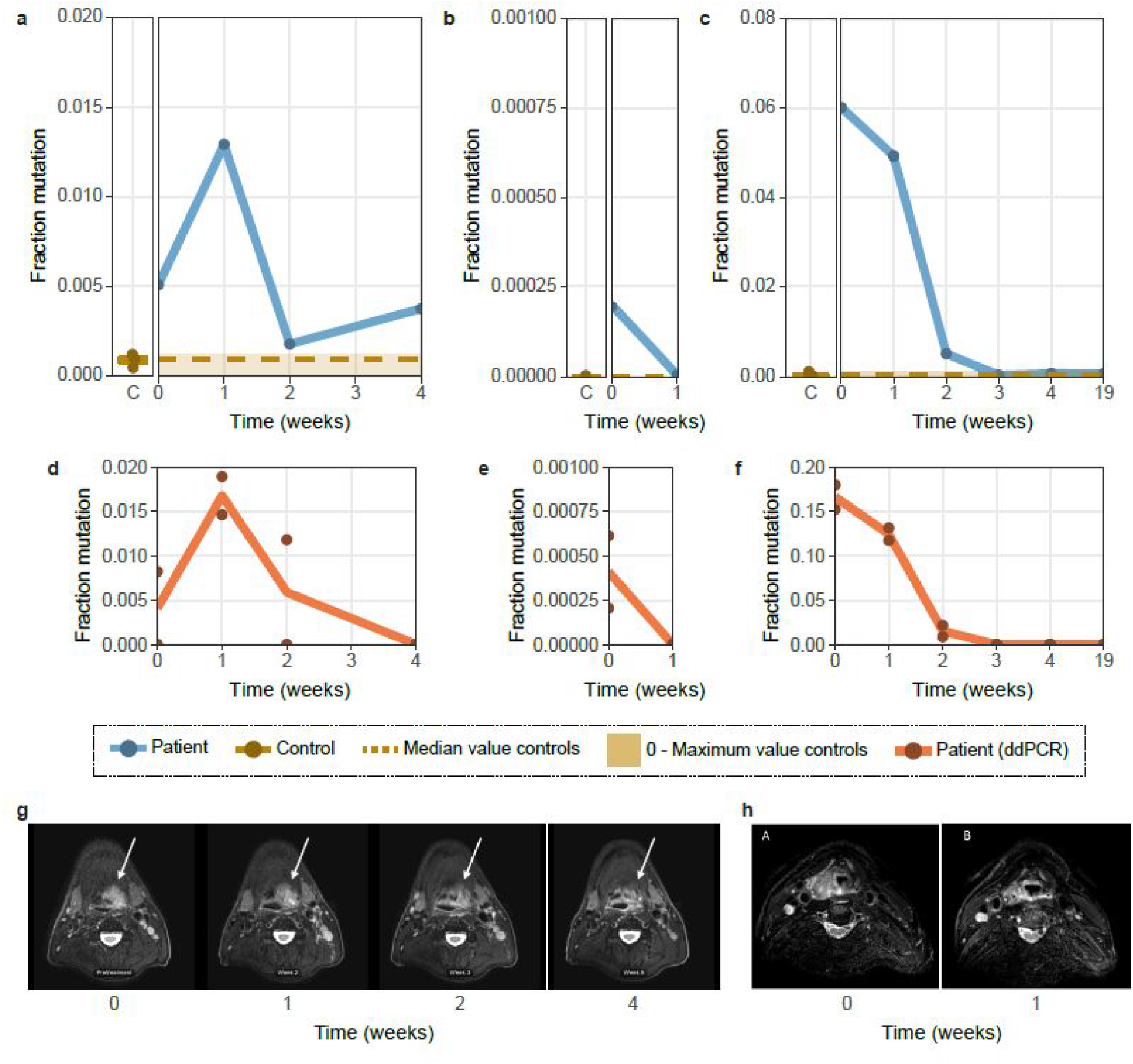
Mutated ctDNA in the blood of patients and controls. **a** 17:7577121 G>A in Patient A (right panel of figure **a**) and three controls (left panel of figure **a**). **b** 17:7577095-7577123 deletion in Patient B (right panel of figure **b**) and two controls (left panel of figure **b**). **c** 17:7578403 C>T in Patient C (right panel of figure **c**) and five controls (left panel of figure **c**). **d** 17:7577121 G>A in Patient A in ddPCR. **e** 17:7576870 C>A in Patient B in ddPCR **f** 17:7578403 C>T in Patient C in ddPCR. **g** MRIs of patient A. White arrow indicates the primary tumor. **h** MRIs of patient B. Each data point is a single measurement and lines show the mean measurement per time in weeks. C indicates ‘controls’. Time indicates the time in weeks after treatment initiation. Median and 0-Maximum values in controls are depicted in yellow in the patient panels of CyclomicsSeq, to support a clear comparison between patients and controls.

Patient A, 57 years of age, presented with a stage II oropharyngeal squamous cell carcinoma with a GTV of 15.5 cm^3^. Chemoradiation reduced the GTV to 1.8 cm^3^. The patient developed locoregional recurrent disease within 10 months after treatment. Patient A had a 17:7577121 G>A (in GRCh37; *TP53*c.817 C>T) missense mutation in *TP53* with a variant allele frequency (VAF) of 0.60 in the tumor. CyclomicsSeq and ddPCR were performed before treatment (time = 0), and 1, 2, and 4 weeks into treatment. Both CyclomicsSeq and ddPCR detected 0.5% ctDNA before treatment (Fig. 6a,d). After an initial increase, the amount of mutated ctDNA dropped but never reached 0% in the CyclomicsSeq measurements. Observations in ddPCR were similar to CyclomicsSeq, but the ddPCR measurement of time point 4 was negative. Unlike ddPCR, the CyclomicsSeq measurement of time point 4 is in line with the observations on MRI that residual tumor was still present at the end of treatment (Fig. 6a,d,g).

Patient B, 56 years of age, presented with a stage IV hypopharyngeal squamous cell carcinoma with a GTV of 12.8 cm^3^. During chemoradiation, GTV initially increased to 17.6 cm^3^, and subsequently reduced to 1.7 cm^3^ five weeks into treatment. On clinical examination and MRI, differentiation between residual disease and post treatment effects was difficult. Patient B died 4 months after treatment of tumor- and/or treatment-associated complications. Patient B had a 17:7576870 C>A (in GRCh37; *TP53*c.976 G>T) nonsense variant with a VAF of 0.22 in the tumor and a 17:7577095-7577123 deletion (in GRCh37; TP53c.815del29) in *TP53* with a VAF 0.34 in the tumor. CyclomicsSeq (aimed at the exon containing the deletion) and ddPCR (aimed at the nonsense mutation) were performed before treatment (time = 0), and 1 week into treatment. Although the assays measured different mutations, both assays detect an initial mutated ctDNA amount of ∼0.02%, which drops below the detection limit 1 week after treatment initiation (Fig. 6b,e,h).

Patient C presented with a stage II oropharyngeal squamous cell carcinoma at the age of 82. Radiotherapy resulted in a recurrence-free survival during one year of follow-up. No GTV-data were available due to lack of patient consent for performing additional MRI measurements. Patient C had a 17:7578403 C>T (in GRCh37; *TP53*c.527 G>A) missense mutation in *TP53* with a VAF of 0.55 in the tumor. CyclomicsSeq and ddPCR were performed before treatment (time = 0), at multiple time points (1, 2, 3 and 4 weeks) during treatment and 19 weeks after treatment initiation. Both CyclomicsSeq and ddPCR showed the presence of ctDNA at time points 0, 1 and 2. Although the observed amounts of mutated ctDNA differ between CyclomicsSeq and ddPCR, both assays detect 0% ctDNA three weeks after treatment initiation, in line with the observed recurrence-free survival (Fig. 6c,f).

## DISCUSSION

Here, we present CyclomicsSeq, a method for detecting ctDNA in liquid biopsies of cancer patients. We show that CyclomicsSeq substantially lowers the error-rate of ONT sequencing by ∼60-fold and, thereby, can facilitate the detection of mutations with a frequency of at least 0.02% in cfDNA in blood of cancer patients. The generation of a consensus sequence allows discrimination between artifacts and true mutations. Similar approaches like INC-Seq, CircSeq and R2C2 have not been optimized to enable efficient sequencing of short molecules such as ctDNA^22–24^. The CyclomicsSeq protocol takes only ∼3 days, including sequencing and data analysis and is universally applicable to any target genomic locus or gene. This is a major advantage over the use of ddPCR to detect mutations in ctDNA, which also requires new primer design (and validation) based on prior knowledge of the mutation present in the tumor for each individual patient. CyclomicsSeq can interrogate complete amplicon panels and leverage real-time ONT sequencing. Therefore, CyclomicsSeq is very suitable for point-of-care clinical workflows.

Although the snFP rate is compatible with the detection of ctDNA in the majority of cancer patients, some genomic positions still suffer from a relatively higher snFP rate. In addition, 0.25% of the bases have a high deletion rate (>10%, Supplementary Fig. 4) that will decrease sensitivity to detect single nucleotide variants at those positions. We aim to lower the error rate further by e.g. removing the PCR step from the protocol. Furthermore, implementation of forward/reverse correction for deletions will likely reduce these mutation rates as well. Finally, the implementation of the 4-nucleotide interspersed barcode sequence in the PCR can aid in deduplication of PCR-amplified molecules and removal of chimeric reads^39^.

We demonstrate CyclomicsSeq using the ONT sequencing platform in the current study, but other long-read sequencing platforms, such as PacBio, can straightforwardly be used to sequence CyclomicsSeq molecules. The technique can also be combined with any PCR kit or protocol, including commercially available PCR amplification panels. In this study, we focused on the applicability in liquid biopsies, but CyclomicsSeq can facilitate cancer diagnostics using tissue-biopsies as well. Additionally, CyclomicsSeq can be expanded to other blood-based measurements, such as foetal cfDNA in non-invasive prenatal diagnostics (NIPD) and potentially even viral cfDNA for the detection and monitoring of viral infections (e.g. COVID-19). In conclusion, CyclomicsSeq is a widely applicable technique that gives reproducible results and can be used in combination with other sequencing technologies to sensitively detect lowly abundant DNA molecules in (liquid) biopsies.

## METHODS

### Human cfDNA

We included three HNSCC patients with known *TP53* mutations (Patient A, B and C). Blood of these HNSCC patients was obtained in the UMC Utrecht within the PREDICT study (NL57164.041.16). Blood of healthy donors was obtained from the Mini Donor Dienst and from the Cyclomics study of the UMC Utrecht. Blood was collected in 10 ml K_2_EDTA blood collection tubes (BD Vacutainer). Use of the human specimens for research purposes was approved by the Medical Ethics Committee of the UMC Utrecht (16-331, 07/125 and 20/055). Informed consent was provided by all participants.

Plasma was isolated within a few hours after blood collection. First, the blood was centrifuged at 800g for 10 minutes. The upper layer was subsequently centrifuged at >14,000g for 1 minute (except for the blood of Patient C), after which supernatant plasma was retrieved and stored at −80°C for further processing After thawing, cfDNA was isolated from 0.5 - 10 ml of plasma using the Quick-cfDNA Serum & Plasma Kit centrifugation protocol according to the manufacturer’s protocol (Zymo Research). cfDNA was eluted twice in the provided elution buffer or in MilliQ and subsequently stored at −80°C. DNA quantity measurement of isolated DNA samples took place using a Qubit fluorometer with double stranded DNA (dsDNA) High Sensitivity Assay Kit (Thermo Fisher Scientific).

### Backbone design principles

The backbones used in CyclomicsSeq are flexible to facilitate circularization of the short DNA molecules, while being as short as possible to limit loss of sequencing capacity to repeated sequencing of the backbone in concatemeric RCA molecules. The backbone sequences were generated with the aid of a genetic algorithm (Supplementary Fig. 2) and selected over a variety of parameters including flexibility, GC content, sequence entropy. The sequences were also checked for the absence of repeated kmers and predicted hairpins. The extremities of the backbone sequence encode for the half of a palindromic restriction site (SrfI). The full restriction site is formed when two backbones are ligated together or a single backbone is self-circularized. This feature facilitates the linearization and subsequent removal of backbone-only circles that may form during the circularization reaction (see ‘Circularization reaction’ paragraph). The backbones also contain four variable bases acting as a barcode. The barcode can be used to detect and resolve chimeric reads and to pool multiple samples in a single run. The barcode bases are not consecutive (like other barcodes used for Nanopore sequencing), but interspersed along the sequence to minimize base-call errors. When a DNA molecule translocates through the protein nanopore during the sequencing process, multiple adjacent bases affect the signal. Some combinations of bases are intrinsically hard to discriminate because they generate a similar signal pattern, thus, placing four consecutive barcode bases would lead to errors in discriminating each of the 256 possible barcodes. Interspersing the barcode bases with multiple (non-barcode) bases guarantees that their signal is well isolated and easily discerned.

### Preparation of the backbone

The backbones were generated by annealing and fill-in of two semi-complementary synthetic phosphorylated oligos purchased from Integrated DNA Technologies (https://www.idtdna.com). A polymerase with error-correction activity was used for the fill-in reaction in order to obtain blunt-end products, with phosphorylated ends. The fill-in reaction consisted of a 25 μl Phusion High-Fidelity Master Mix 2X (New England Biolabs), 23 μl of water and 1ul of each oligo at a concentration of 10 μM. The reaction was subjected to 5 cycles of DNA melting (1 minute at 98°C), annealing (30 seconds at 65°C) and elongation (15 seconds at 72°C). All the backbones were gel-purified.

### Preparation of Insert

Per sample, 2 to 10 ng of cfDNA was used for PCR-based enrichment of *TP53* sequences. Briefly, 20 μl reaction mixture composed by 10 μl of Phusion High-Fidelity Master Mix 2X (NEB), 2 μl of Betaine 5M (Sigma-Aldrich), 0.5 μl of pure DMSO (Sigma-Aldrich), 0.5 μM of the forward and the reverse primers each, cfDNA, and MilliQ (to a volume of 20 μl)) was prepared. If necessary, the volume of the cfDNA was reduced by SpeedVac at medium temperature prior to preparation of the PCR mix. The PCR reaction consisted of 1 minute incubation at 98°C, 30 cycles (Supplementary Table) of 30 seconds at 98°C and 15 seconds at 59°C, and finally 2 minutes incubation at 72°C. PCR products were gel-purified using the Wizard SV Gel and PCR Clean-Up System (Promega) according to the manufacturer’s protocol. PCR products were kept at −20°C. Sample CY_SM_PC_HC_0004_003 and CY_SM_PC_HC_0004_004 were amplified using the CleanPlex *TP53* Panel of Paragon Genomics according to manufacturer’s protocol.

### Circularization reaction

The reaction mix for circularization of the backbone and insert (3:1 backbone:insert ratio, 20 to 60 ng of insert), 5 μl 10X CutSmart Buffer (New England Biolabs), 10 μl 10mM ATP (New England Biolabs), 2 μl T4 ligase (New England Biolabs), 2 μl SrfI (New England Biolabs), and MilliQ (until a volume of 50 μl)) was prepared on ice. Circularization was performed by 8 cycles of 10 minutes at 16°C and 10 minutes at 37°C, followed by 20 minutes at 70°C. To digest any residual backbone-backbone byproducts, 1 μl of SrfI (New England Biolabs) was added and the mixture was incubated for 10 minutes at 37°C, followed by 20 minutes at 70°C. The linear DNA was then removed using Plasmid-Safe DNase (Lucigen). Briefly, the circularization mixture was combined with 6 μl 10X Plasmid-Safe Buffer (Lucigen), 2 μl Plasmid-Safe Enzyme (Lucigen), and 6 μl 10mM ATP (New England Biolabs). Linear DNA was then digested by 30 minute incubation at 37°C, followed by 30 minutes inactivation at 70°C. Circular DNA was purified using the QIAquick nucleotide removal kit according to the manufacturer’s protocol (Qiagen).

### Rolling circle amplification (RCA)

Circular DNA obtained by the circularization reaction was combined with 12 μl 5X Annealing buffer (50 mM Tris @ pH 7.5-8.0, 250 mM NaCl, 5 mM EDTA) and 1 μl Exo-resistant random primers (Thermofisher), heated for 5 minutes at 98°C and then cooled down at room temperature. Subsequently, the RCA mix (previous reaction mixture, 10 μl 10X Phi29 Buffer (Thermofisher), 2 μl BSA (New England Biolabs), 10 μl dNTPs (Thermofisher), 4 μl pyrophosphatase (Thermofisher), 2 μl Phi29 Polymerase (Thermofisher), and MQ (to a volume of 100 μl)) was prepared. RCA was performed overnight at 30°C. The RCA-reaction was inactivated by 10 minute incubation at 70°C.

To test whether CyclomicsSeq worked, 4 μl of RCA mixture was incubated with a restriction enzyme that specifically cuts backbone-backbone interactions, but not backbone-insert interactions. Briefly, 4 μl of RCA mixture was combined with 4 μl Restriction enzyme buffer (New England Biolabs), 13 μl MilliQ, and 1 μl BglII (New England Biolabs). The reaction mixture was incubated for 1 hour at 37°C and then ran on a 1.5% Agarose gel.

### Production of plasmids for PCR-free experiments

Synthetic sense and antisense oligos were purchased from Integrated DNA Technologies in order to produce the following two dsDNA strands, encoding for a single exon of the *TP53* gene.

~~~
**>Amplicon_12 - WT**(length: 141)
CTTGCTTACCTCGCTTAGTGCTCCCTGGGGGCAGCTCGTGGTGAGGCTCCCCTTTCTTGCGGAGATTCTCTTCC
TCTGTGCGCC**<u>G</u>**GTCTCTCCCAGGACAGGCACAAACA**<u>CG</u>**CACCTCAAAGCTGTTCCGTCCCAGTAGAT
~~~

~~~
**>Amplicon_12 - MUT**(length: 141)
CTTGCTTACCTCGCTTAGTGCTCCCTGGGGGCAGCTCGTGGTGAGGCTCCCCTTTCTTGCGGAGATTCTCTTCC
TCTGTGCGCC**<u>A</u>**GTCTCTCCCAGGACAGGCACAAACA**<u>TA</u>**CACCTCAAAGCTGTTCCGTCCCAGTAGAT
~~~

The complementary strands, 0.5 μM each, were mixed in 1X CutSmart buffer (NEB) and annealed by keeping the reaction at 98°C for 5 minutes and then let it cool down at room temperature. The annealed product was gel purified and cloned into a pJet vector using the CloneJET PCR Cloning Kit (Thermofisher). The vector was used to transform chemically-competent *E. coli* TOP10 cells. The cells were selected on LB plates supplemented with Ampicillin. Single colonies were picked used to inoculate fresh LB plates. This second expansion was done to ensure the monoclonality of the subsequent cultures. Single colonies were picked and cultured in liquid medium (LB with Ampicillin) for 16 hours. The plasmid DNA was extracted from each culture using the QIAprep Spin Miniprep Kit (Qiagen) and sequenced using the Sanger method to guarantee the correctness of the amplified sequences.

The plasmid DNA was quantified using a NanoDrop spectrophotometer (Thermofisher), all the preps were adjusted to the same concentration of 25 ng/μl. Three solutions were prepared, one containing only the WT sequence, one constraining only the MUT sequence and one containing 0.1% of MUT in WT. These samples were used as input for a rolling circle amplification reaction.

### Repeatability assay

Four different inserts with a sequence matching a region of *TP53* (WT) and mutant sequences derived from the WT sequence (M0, M1, M2; four mutations each) were produced by annealing and elongation of synthetic oligos as described in the previous paragraph.

~~~
**>WT | 17:7579205-7579355**
GAAGCCAAAGGGTGAAGAGGAATCCCAAAGTTCCAAACAAAAGAAATGCAGGGGGATACGGCCAGGCATTGAAG
TCTCATGGAAGCCAGCCCCTCAGGGCAACTGACCGTGCAAGTCACAGACTTGGCTGTCCCAGAATGCAAGAAGC
CCA
~~~

~~~
**>M0 | [‘G_7579255:A’, ‘G_7579257:A’, ‘C_7579263:A’, ‘G_7579302:C’]**
GAAGCCAAAGGGTGAAGAGGAATCCCAAAGTTCCAAACAAAAGAAATGCA**<u>A</u>**G
**<u>A</u>**GGATA**<u>A</u>**GGCCAGGCATTGAAG
TCTCATGGAAGCCAGCCCCTCAG**<u>C</u>**GCAACTGACCGTGCAAGTCACAGACTTGGCTGTCCCAGAATGCAAGAAGC
CCA
~~~

~~~
**>M1 | [‘T_7579261:C’, ‘A_7579272:C’, ‘G_7579293:A’, ‘C_7579304:T’]**
GAAGCCAAAGGGTGAAGAGGAATCCCAAAGTTCCAAACAAAAGAAATGCAGGGGGA**<u>C</u>**ACGGCCAGGC**<u>C</u>**TTGAAG
TCTCATGGAAGCCA**<u>A</u>**CCCCTCAGGG**<u>T</u>**AACTGACCGTGCAAGTCACAGACTTGGCTGTCCCAGAATGCAAGAAGC
CCA
~~~

~~~
**>M2 | [‘G_7579252:A’, ‘C_7579266:A’, ‘A_7579268:G’, ‘C_7579295:T’]**
GAAGCCAAAGGGTGAAGAGGAATCCCAAAGTTCCAAACAAAAGAAAT**<u>A</u>**CAGGGGGATACGG**<u>A</u>**C**<u>G</u>**GGCATTGAAG
TCTCATGGAAGCCAGC**<u>T</u>**CCTCAGGGCAACTGACCGTGCAAGTCACAGACTTGGCTGTCCCAGAATGCAAGAAGC
CCA
~~~

The inserts were gel purified and diluted in water to a concentration of 4 ng/μl and mixed in a ratio 8:4:2:1 (WT:M0:M1:M2). 10 μl of such a mix was used for a circularization reaction with 14 ul of BB25 (14 ng/μl), followed by RCA.

### ONT sequencing

RCA products are purified using AMPure beads. Subsequently, branched DNA (which can be a consequence of the RCA) was resolved by a 1 hour incubation at 37°C with 4 μl T7 endonuclease (New England Biolabs) and re-purified using AMPure beads. ONT libraries were prepared according to manufacturer’s protocol version SQK-LSK109 using 1500ng as input DNA, extending the DNA repair step to 50 minutes and the adapter ligation to 30 min.

### Data processing

Sequencing data was processed using the cyclomics consensus pipeline available in our GitHub repository (https://github.com/UMCUGenetics/Cyclomics_consensus_pipeline). Individual concatemer sequence reads were mapped to a targeted reference genome which only included the backbone and insert loci (e.g. *TP53*) sequences. Mapping was performed using LAST (v921) to separate individual copies (http://last.cbrc.jp/). Primary mapping by lastal^40^ (parameters -Q 1 -p {last_param}, last_param is available in the Github repository) was followed by lastsplit^41^ (default settings) and maf-convert (default settings) to obtain a SAM file. SAM files were sorted and converted to BAM files using Sambamba^42^. These BAM files contain mapping information of each individual copy of the backbone or insert that was present in the original concatemer sequence reads. The targeted-mapped BAM files were used as a basis for consensus calling in three strategies: default consensus calling, repeat count analysis, and forward and reverse splitting.

### Default consensus calling

For the default consensus calling BAM files were converted to the m5 format using bam2m5 (https://github.com/sein-tao/bam2m5, commit 0ef1a930b6a0426c55e8de950bf1ac22eef61bdf) which severed as input for the pbdagcon tool (https://github.com/PacificBiosciences/pbdagcon, commit 3c382f2673fbf3c5305f5323188e790dc396ac9d) to construct consensus reads. Settings for pbdagcon were -m 35 (Minimum length for correction), -t 0 (Trim alignments on either size), and -c 10 (Minimum coverage for correction). For pJET experiments, -c 5 was used. Resulting consensus reads were added to a run specific FASTA file and subsequently mapped to the entire human reference genome (hg19) including the backbone sequences as separate contigs. Reference genome mapping was performed using bwa-mem v0.7.17 (https://github.com/lh3/bwa) using options -c 100 and -M. Sambamba was used to sort and convert the SAM to BAM^42^.

### Repeat count analysis

For the repeat count analysis BAM files were binned based on the repeat count (1 – 39 and 40+) for the locus of interest (either insert or backbone). Repeat count is defined as maximum coverage at the locus of interest to circumvent possible splitting of an individual copy during last mapping. Consensus calling was performed without a coverage threshold (-c 1) to ensure consensus calling in all repeat bins. Resulting consensus reads were added to a bin specific fasta file. Full reference mapping and allele counting was performed for each repeat bin as mentioned in the ‘default consensus calling’ section.

### Forward and reverse splitting

For the forward and reverse splitting BAM files were binned based on forward or reverse orientation for the locus of interest (either insert or backbone). Forward or reverse is defined as the majority of reads in the locus of interest that map either on the forward (bitwise flag 0) or reverse (bitwise flag 16) orientation. Consensus calling, full reference mapping and allele counting was performed for each forward or reverse bin as mentioned in the ‘default consensus calling’ section.

### Run stats

The file structure.txt, generated by the pipeline, is parsed to determine the read length distribution and the ratio between backbone and insert for each read of a run. These features are then used to group the reads into the categories found in Fig. 1c-e and in the SupplementaryData. The code used is available in the jupyter notebook *Stats_from_structure.ipynb*, available in the GitHub repository (https://github.com/UMCUGenetics/CyclomicsManuscript).

### *TP53* coverage

The coverage of consensus reads on *TP53* is computed using samtools depth, without coverage limit (option -d=0), on the bam files generated by the pipeline, suffixed with *“full_consensus.sorted.bam”*. The resulting table was used to generate the plot of Figure 2 using the jupyter notebook *TP53_panel_coverage.ipynb*, available in the shared GitHub repository.

### COSMIC analysis

To determine the false positive rate specifically for COSMIC mutations (Figure 4), the number of the consensus bases called at each COSMIC position was counted. The false-positive rate, for each position, was calculated as the percentage of COSMIC mutation over the total coverage. For the position for which there exists a bias in the sequencing accuracy between the forward and the reverse strand, the consensus was computed separately, and the base counts and coverage from the most accurate strand were used. The code used is available in the jupyter notebook *COSMIC_analysis.ipynb*, available in the shared GitHub repository.

### snFP rate, error rate and forward/reverse correction

For each base position in each sample, allele frequencies were determined using Sambamba v0.6.5 depth (base -L {region of interest} --min-coverage=0)^42^. Subsequently, the snFPrate and the combined snFP&deletion rate were determined by dividing the errors by the total coverage. For the files with 1 to 40+ repeats (obtained at section ‘repeat count analysis’) separately, these rates were computed per table as a sum of all errors in the table divided by the sum of the coverage in a table. For the consensus called files with a cutoff of at least 10 repeats (all reads obtained in the section ‘default consensus calling’ and forward & reverse reads obtained in the section ‘forward and reverse splitting’), these rates were calculated per base position for the bases with at least 100X coverage. Mutated bases and barcode positions were blacklisted in these analyses.

For each base position in the 10+ consensus called files, we next determined whether taking only forward or reverse reads would reduce the mean snFP rate by more than 1/1000. If so, only forward or reverse measurements were considered for these positions specifically. Mean snFP and deletion rates and standard deviations were calculated per base position across the samples. Furthermore, the mean number of bases per sample with error rates <0.1%, 0.1-1% and >1% were computed.

### Forward/reverse correction reproducibility test

For this analysis all samples with BB24 that were sequenced with the R9 flow cell were included. Four samples were chosen at random as ‘training’ samples and the remaining four samples were the ‘testing’ samples. First, the bases that need forward/reverse correction were defined using the ‘training’ samples only, similar as described previously. Subsequently, the ‘testing’ samples were forward/reverse corrected. We then plotted both the uncorrected and the forward/reverse corrected snFP rate.

### Droplet digital PCR

ddPCR was performed as described previously^14^. Briefly, a ddPCR reaction was prepared (13 μl mastermix and 9 μl cfDNA) and subsequently ran on a QX200 ddPCR system according to protocol (Bio-Rad Laboratories). Data analysis was performed using QuantaSoft v1.7.4.0917 (Bio-Rad Laboratories). Each experiment was carried out in duplicate, and mean number of positive droplets were used as a proxy for ctDNA concentrations.

## Supporting information

TOC

## Data availability

The sequencing datasets generated during the current study are available at EGA (https://www.ebi.ac.uk/ega/), under accession number EGAS00001003759. The processed datasets analyzed during the current study are available at Zenodo (https://doi.org/10.5281/zenodo.3925250).

## Code availability

CyclomicsSeq scripts for processing raw data to consensus called data are available through Github (https://github.com/UMCUGenetics/Cyclomics_consensus_pipeline). Data analysis scripts used to process and plot the data for the current study are available through Github (https://github.com/UMCUGenetics/CyclomicsManuscript).

## ACKNOWLEDGEMENTS

The authors would like to thank Manon Huibers for input on the study, Floris Reinders for input on the manuscript and the Utrecht Sequencing Facility for sequencing. The Utrecht Sequencing Facility is subsidized by the University Medical Center Utrecht, Hubrecht Institute, and Utrecht University. This study was financially supported by the Oncode Institute (project number P2018-004) and is part of the Oncode Institute, which is partly financed by the Dutch Cancer Society.

## AUTHOR INFORMATION

### Co-first authors

Alessio Marcozzi and Myrthe Jager contributed equally to this work.

### Author contributions

W.P.K. and J.d.R. were involved in the conceptual design of the CyclomicsSeq protocol. A.M. developed the wet-lab protocol. A.M., M.J., M.E. and R.S. developed the bioinformatics data processing protocol. A.M., M.J. and B.P. collected the samples. A.M., M.J., I.R. and W.P.K. performed CyclomicsSeq wet-lab experiments. I.R. performed ONT sequencing. A.M., M.J., M.E., R.S. and L.C. performed bioinformatic analyses. J.H.G. and J.K. performed ddPCR experiments. B.P. collected MRIs. C.T., R.B., L.A.D. and S.M.W., were involved in the conceptual design of the HNSCC patient study. A.M., M.J., W.P.K. and J.d.R. were involved in the conceptual design of this study. A.M., M.J., W.P.K. and J.d.R. wrote the manuscript. All authors provided textual comments and have approved the manuscript. W.P.K. and J.d.R. supervised this study.

