## Supplementary figures and images for "Accurate detection of circulating tumor DNA using nanopore consensus sequencing"

### CY_SS_PC_HN_0003_001_000_BB-I_.pdf

# CY_SS_PC_HN_0003_001_000

#### BB:I Ratio

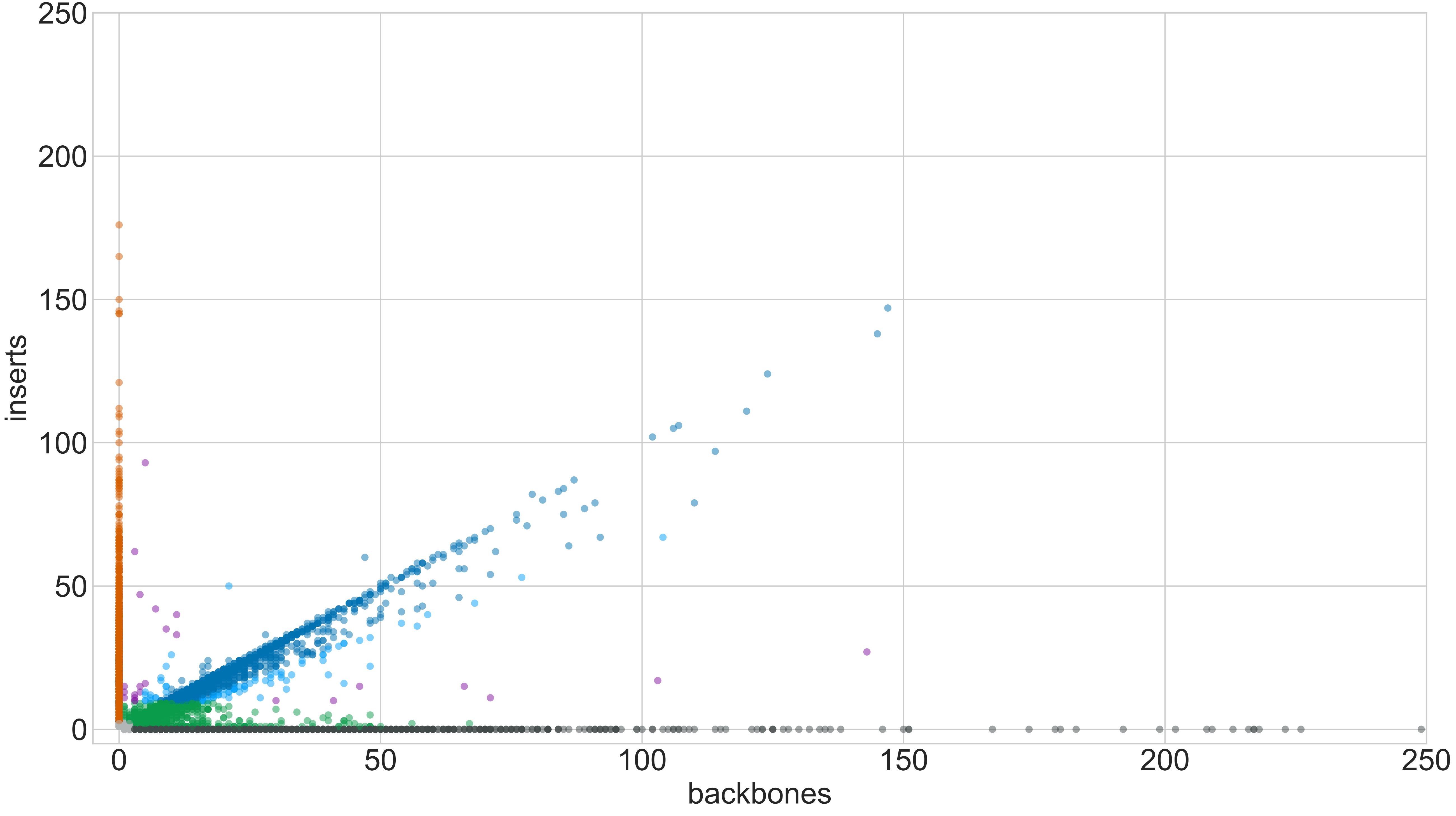

### CY_SS_PC_HN_0003_001_000_GROUPS_STACKED_NORM-LEN_.pdf

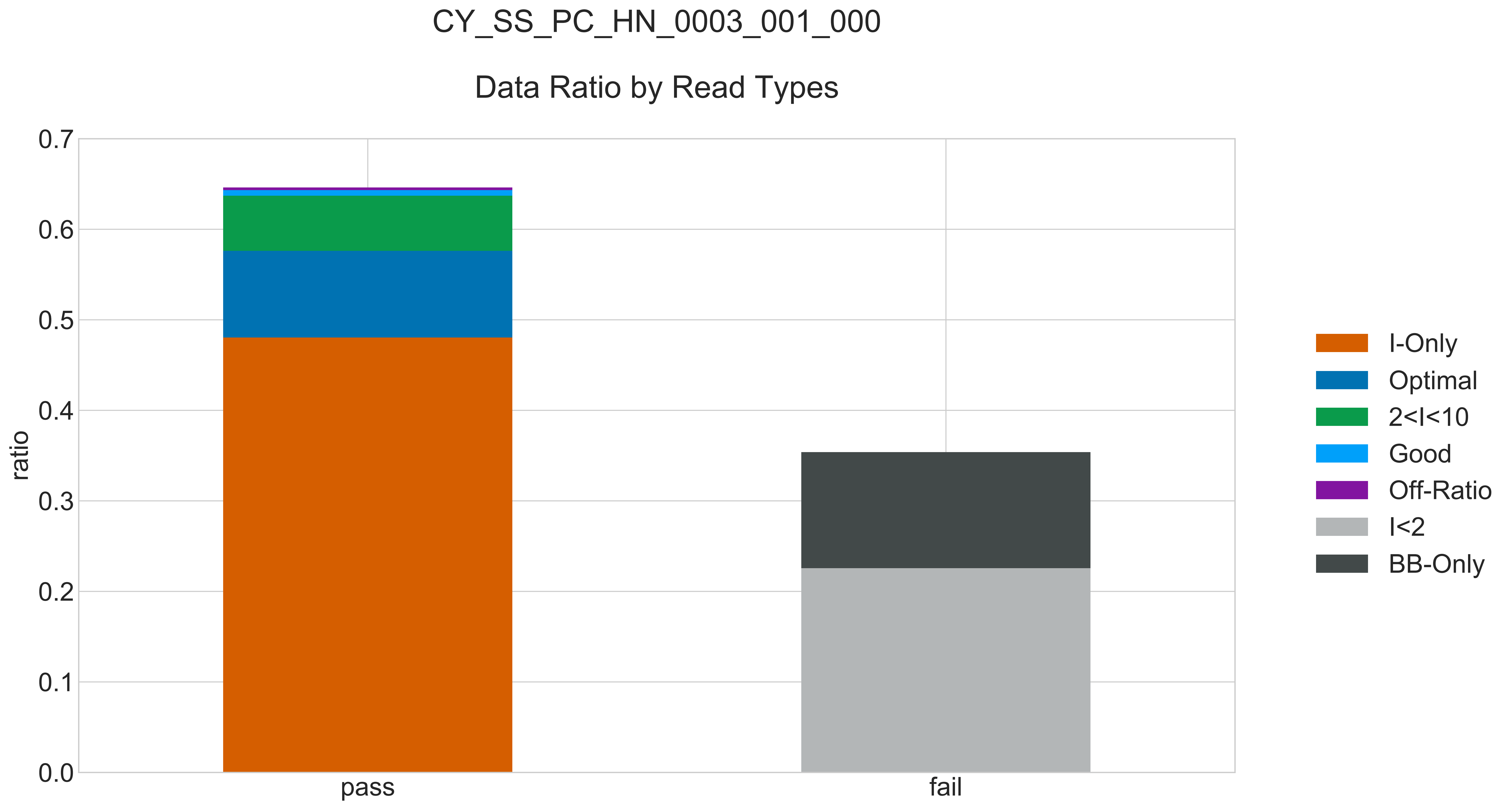

### CY_SS_PC_HN_0003_001_000_READ_LEN_.pdf

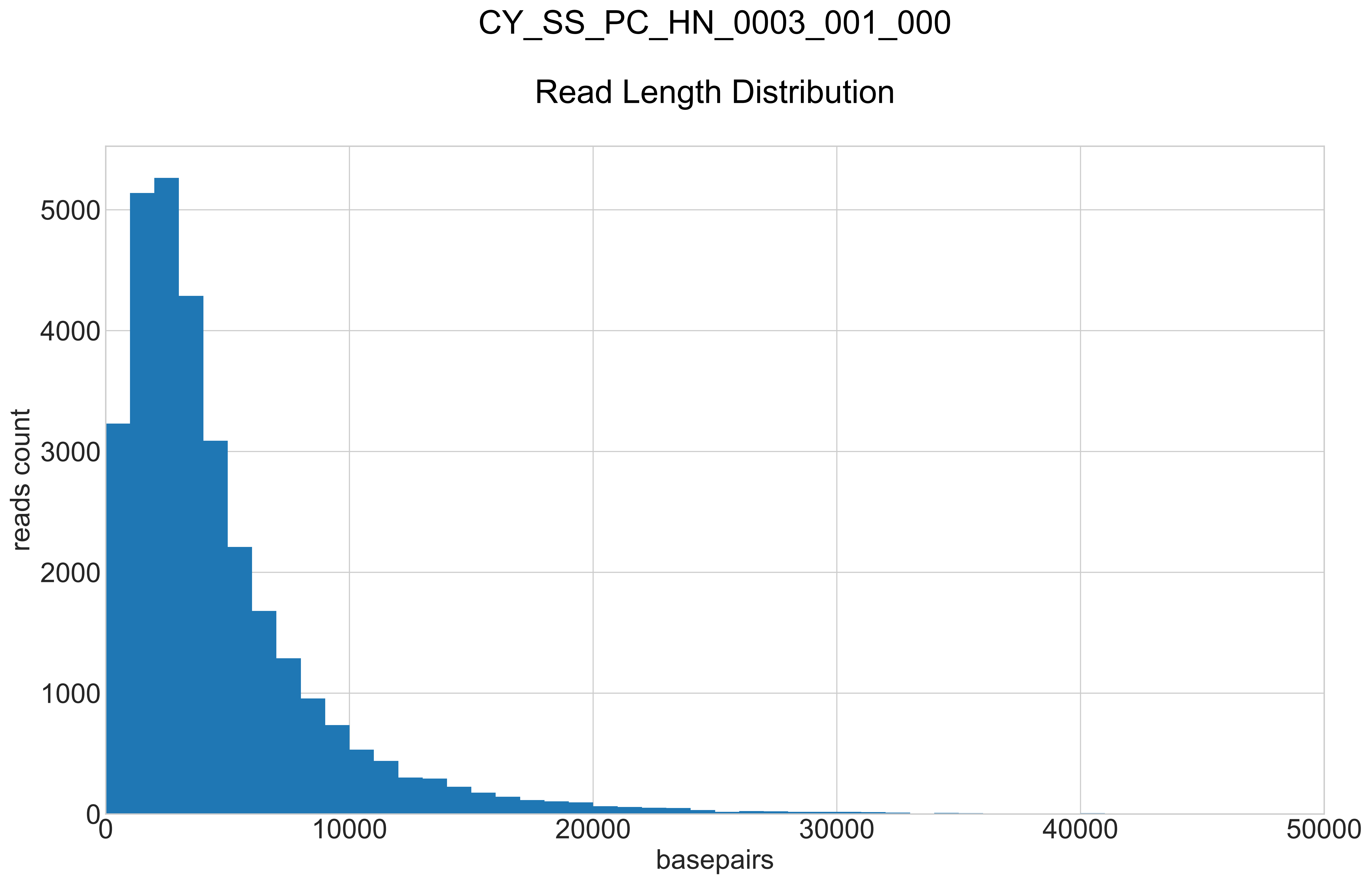

### CY_SS_PC_HN_0003_001_000_REPEATS_COUNT_.pdf

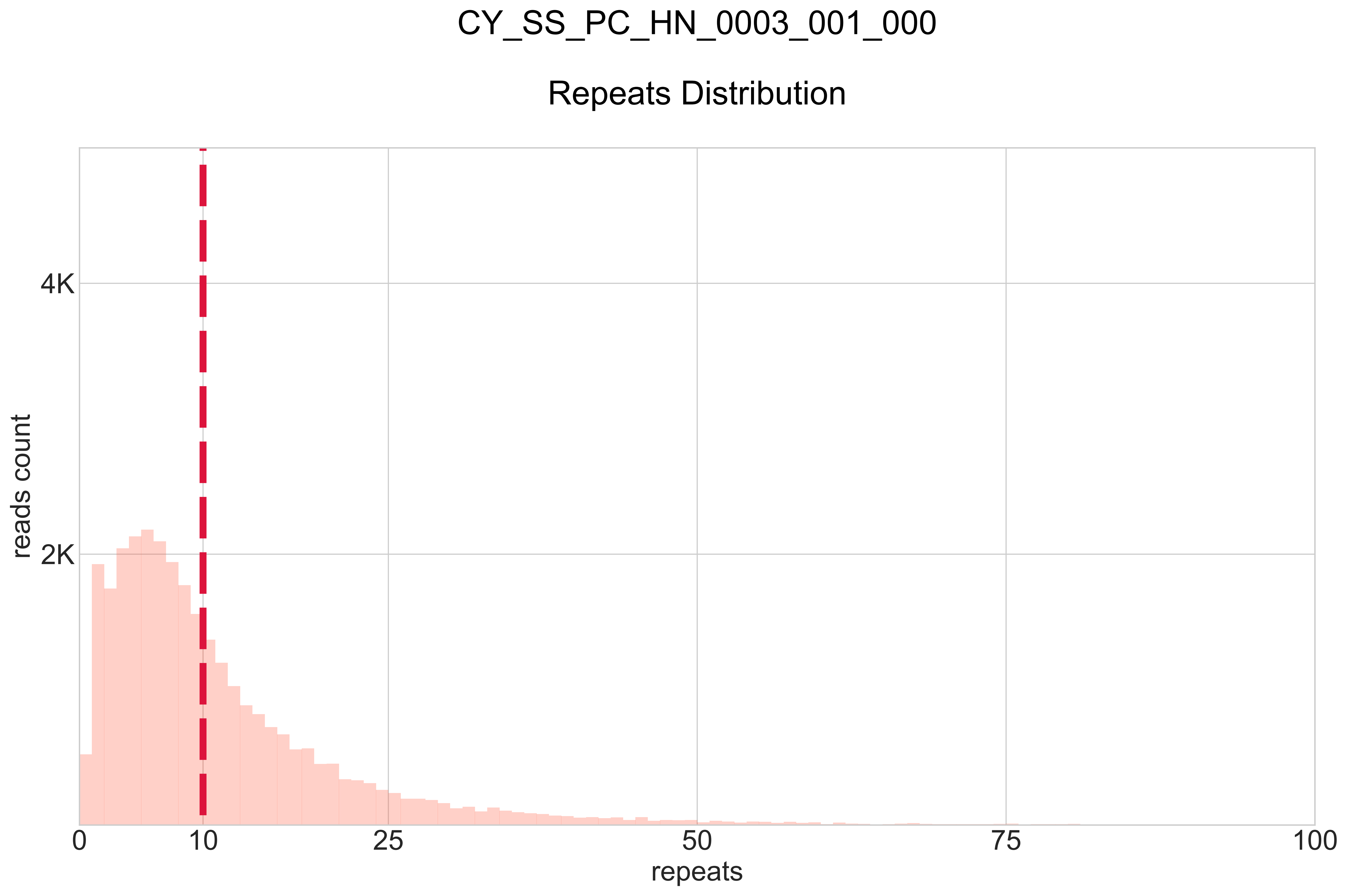

### CY_SS_PC_HN_0003_002_000_BB-I_.pdf

# CY_SS_PC_HN_0003_002_000

#### BB:I Ratio

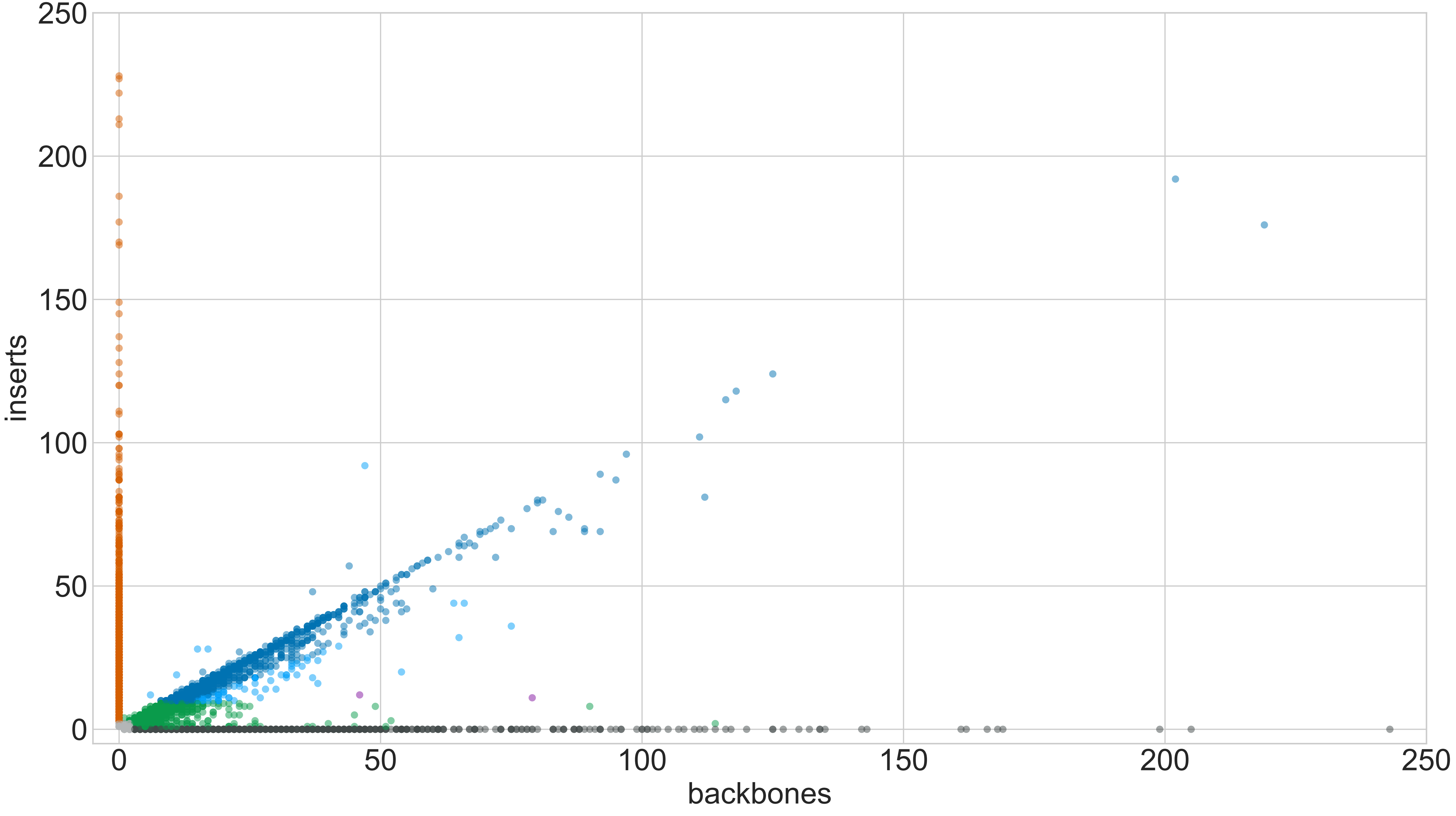

### CY_SS_PC_HN_0003_002_000_GROUPS_STACKED_NORM-LEN_.pdf

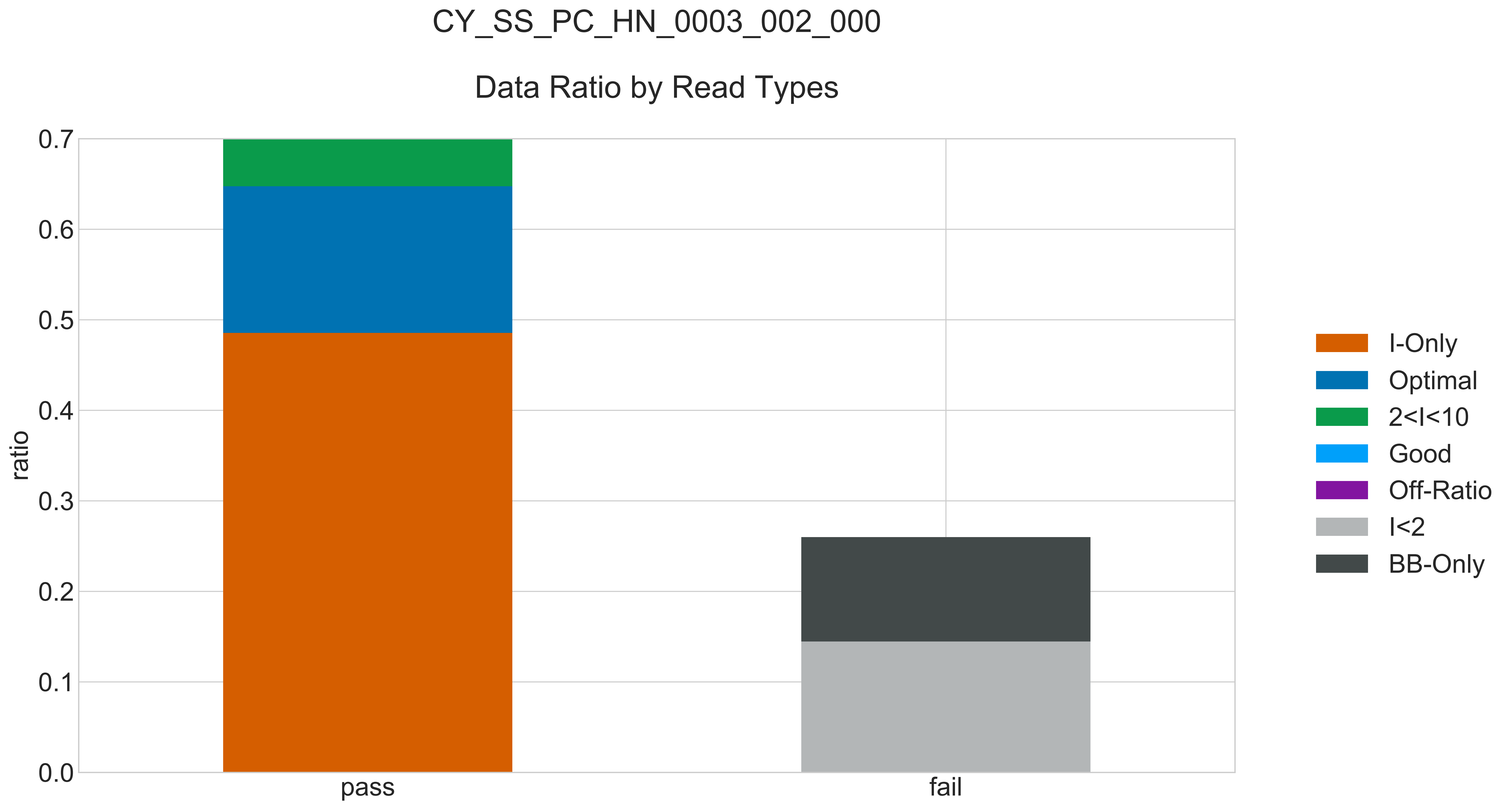

### CY_SS_PC_HN_0003_002_000_READ_LEN_.pdf

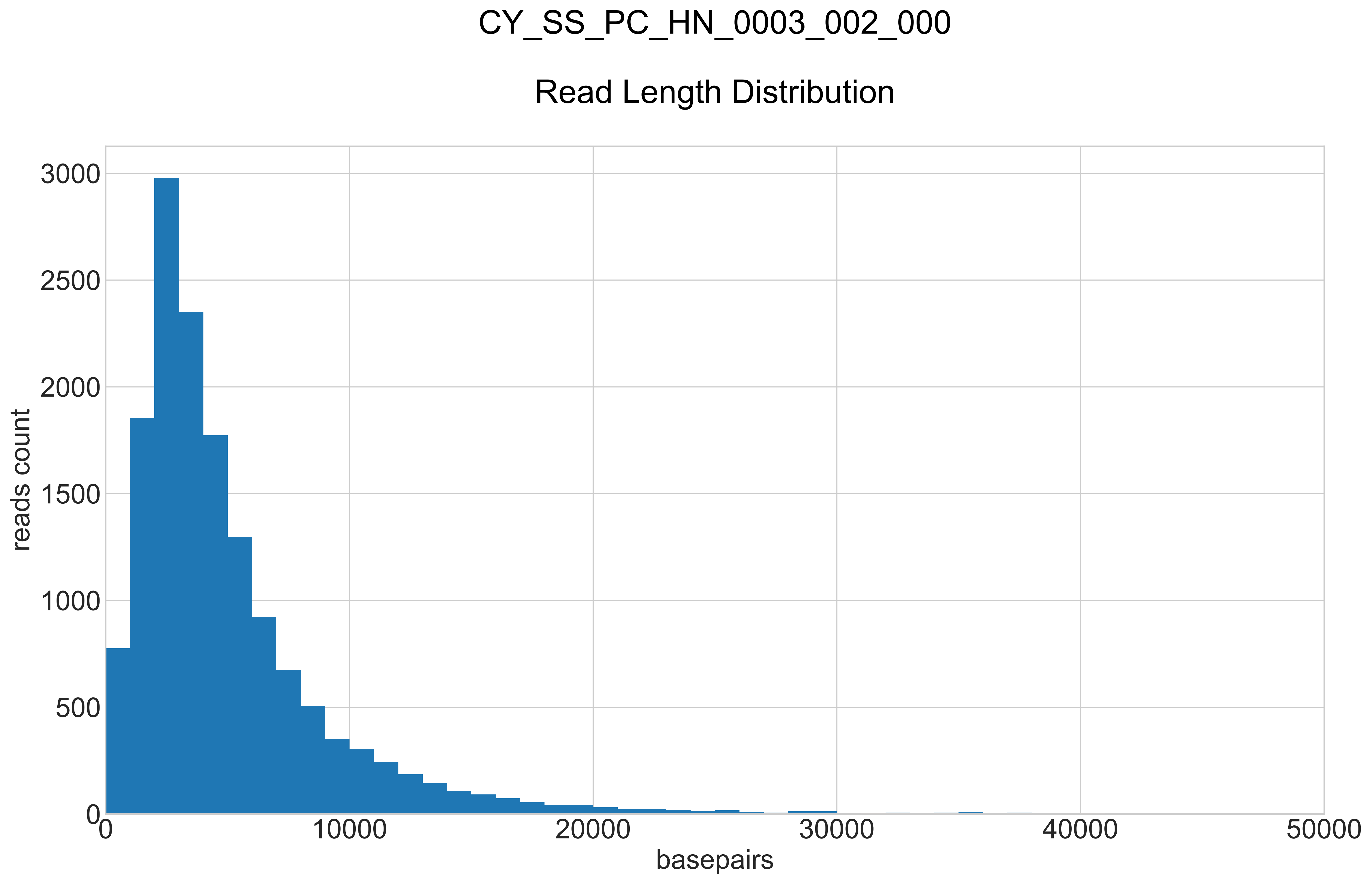

### CY_SS_PC_HN_0003_002_000_REPEATS_COUNT_.pdf

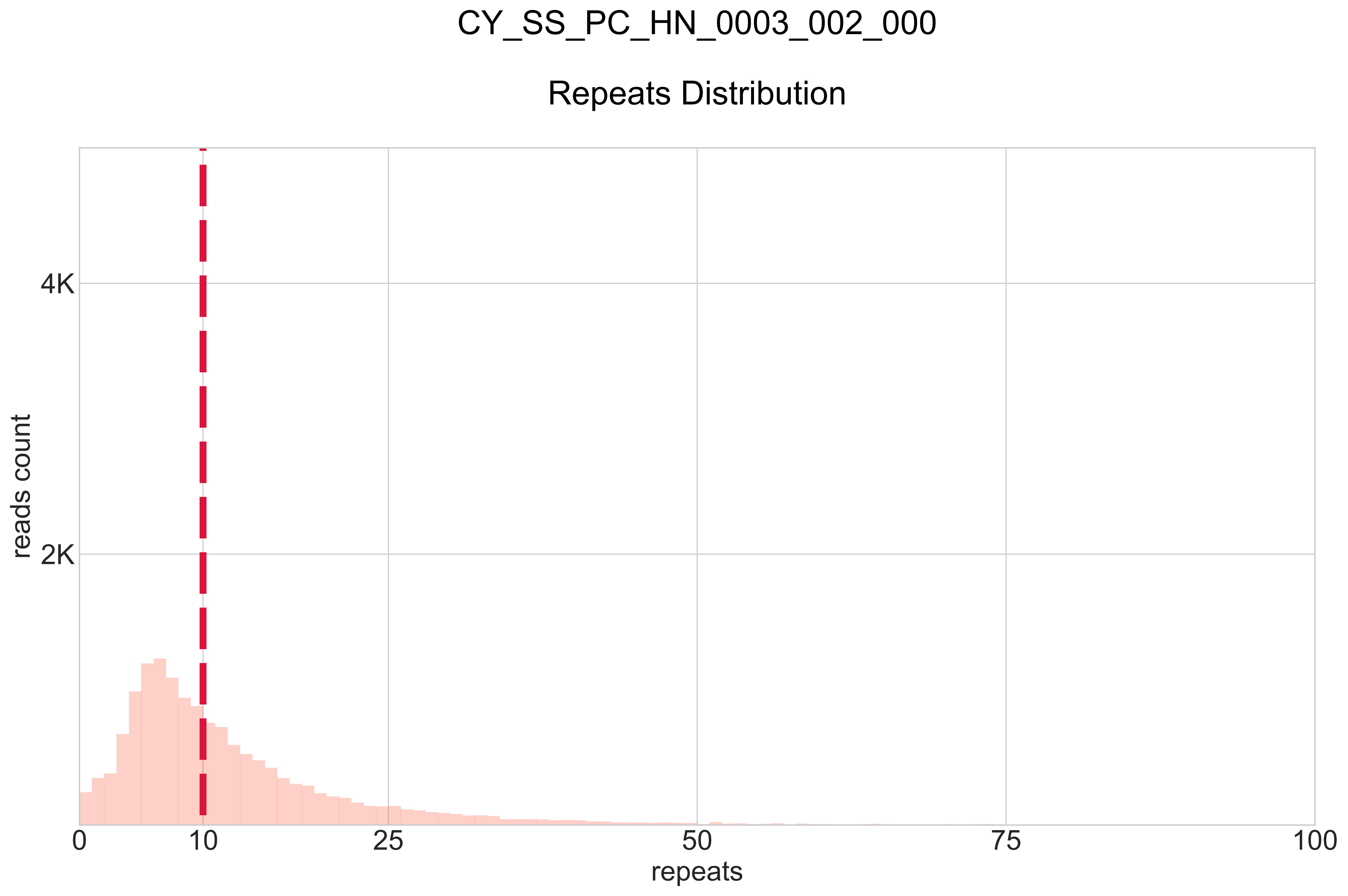

### CY_SS_PC_HN_0003_003_000_BB-I_.pdf

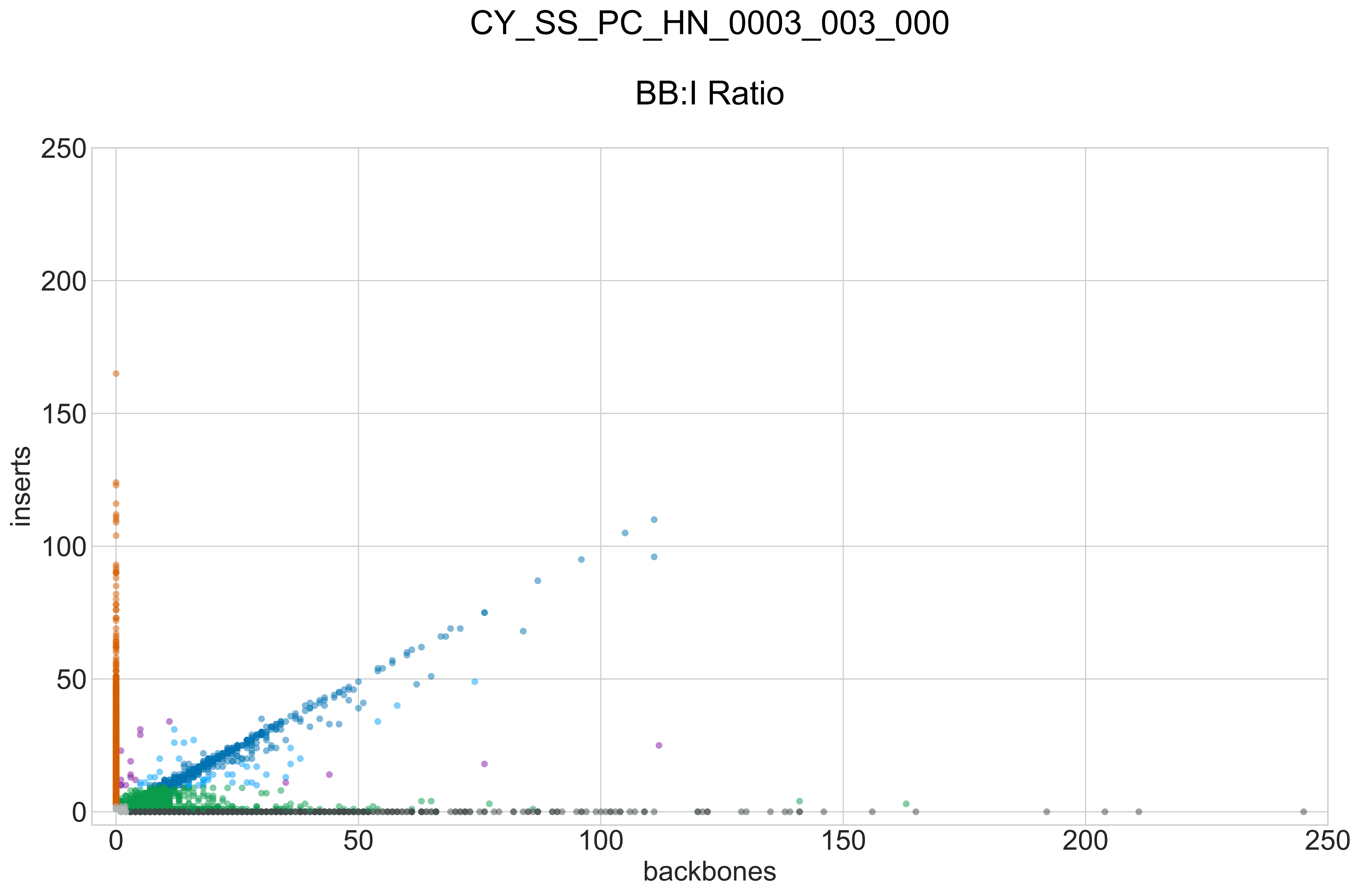

### CY_SS_PC_HN_0003_003_000_GROUPS_STACKED_NORM-LEN_.pdf

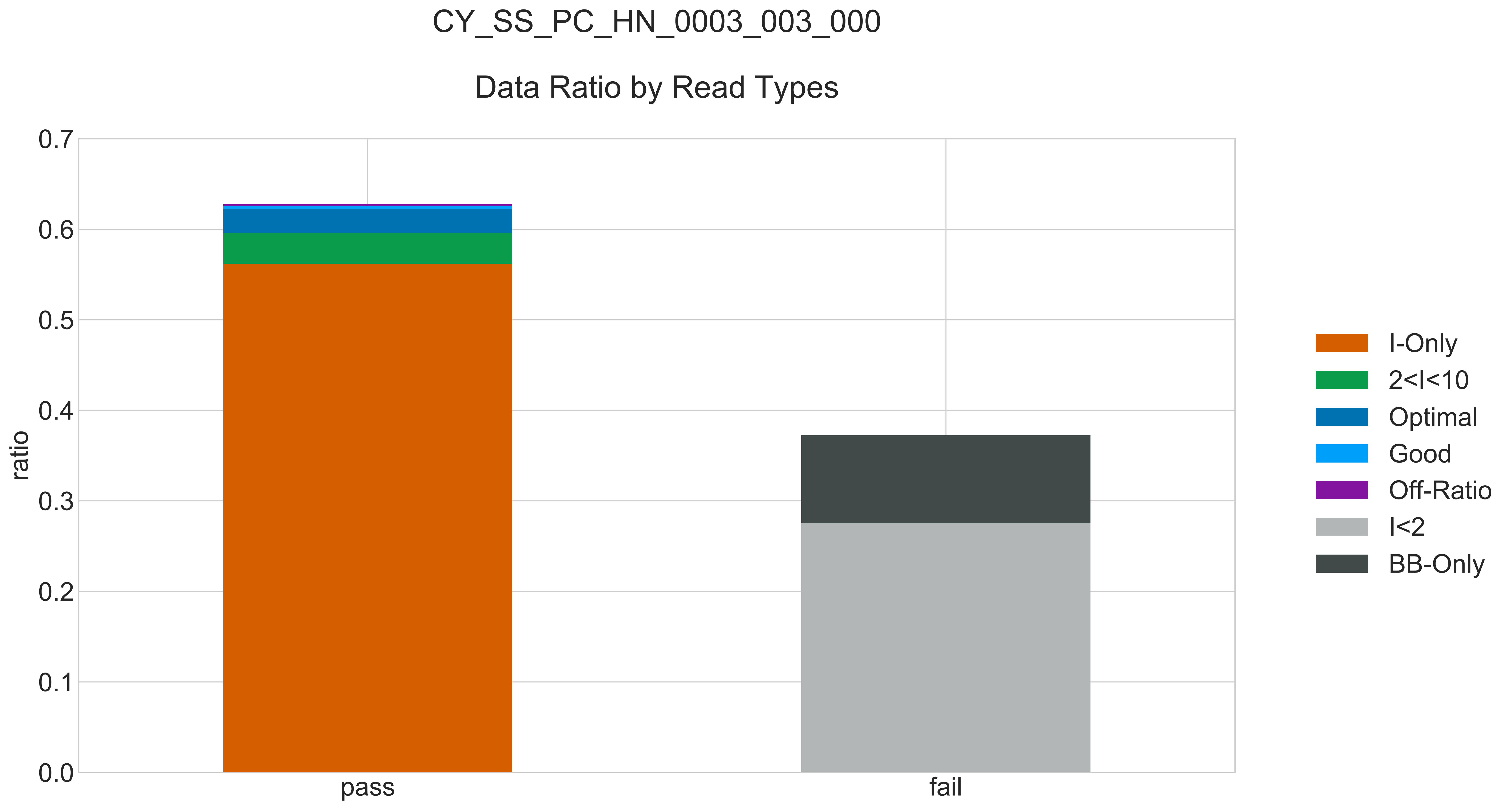

### CY_SS_PC_HN_0003_003_000_READ_LEN_.pdf

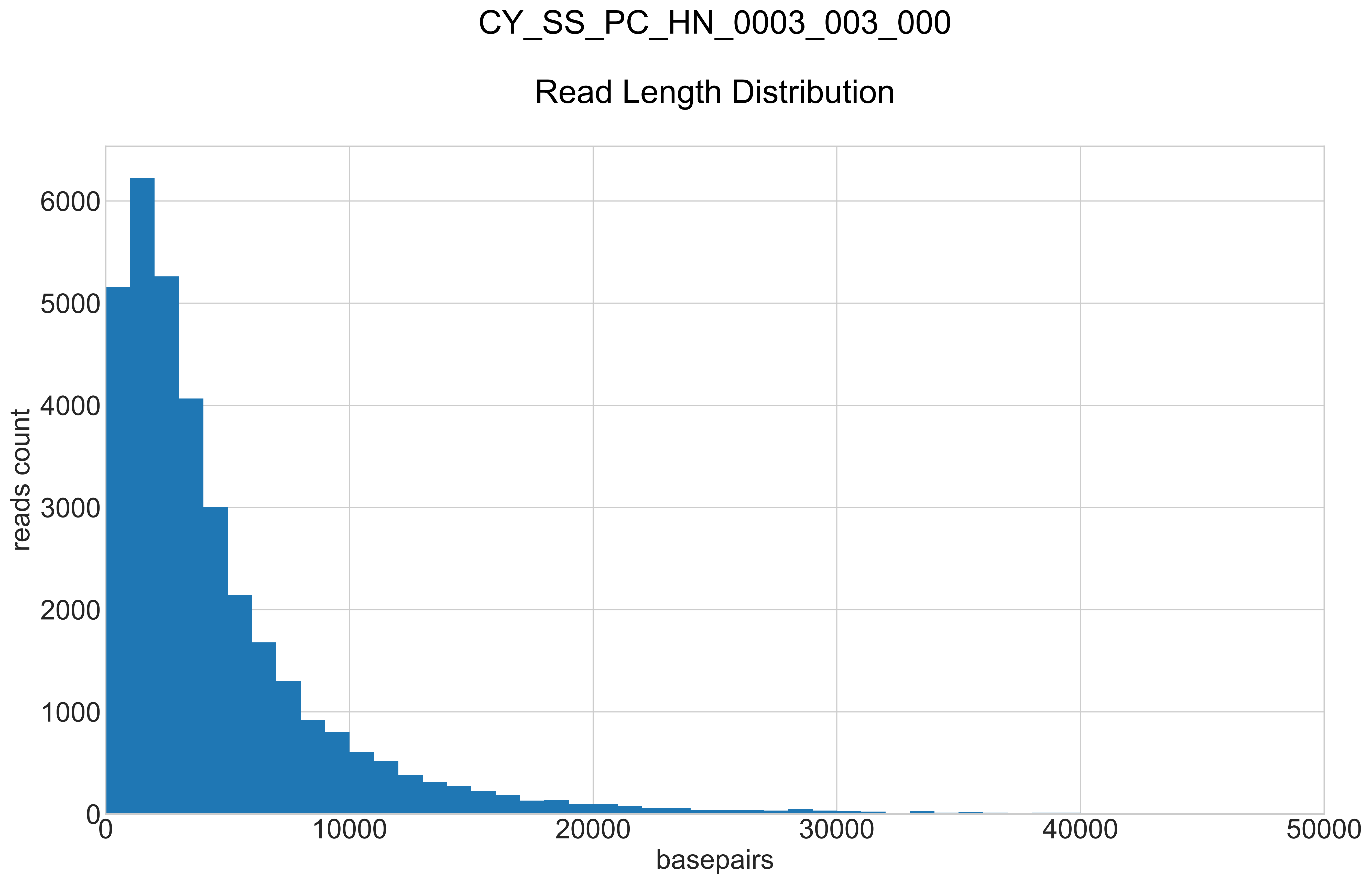

### CY_SS_PC_HN_0003_003_000_REPEATS_COUNT_.pdf

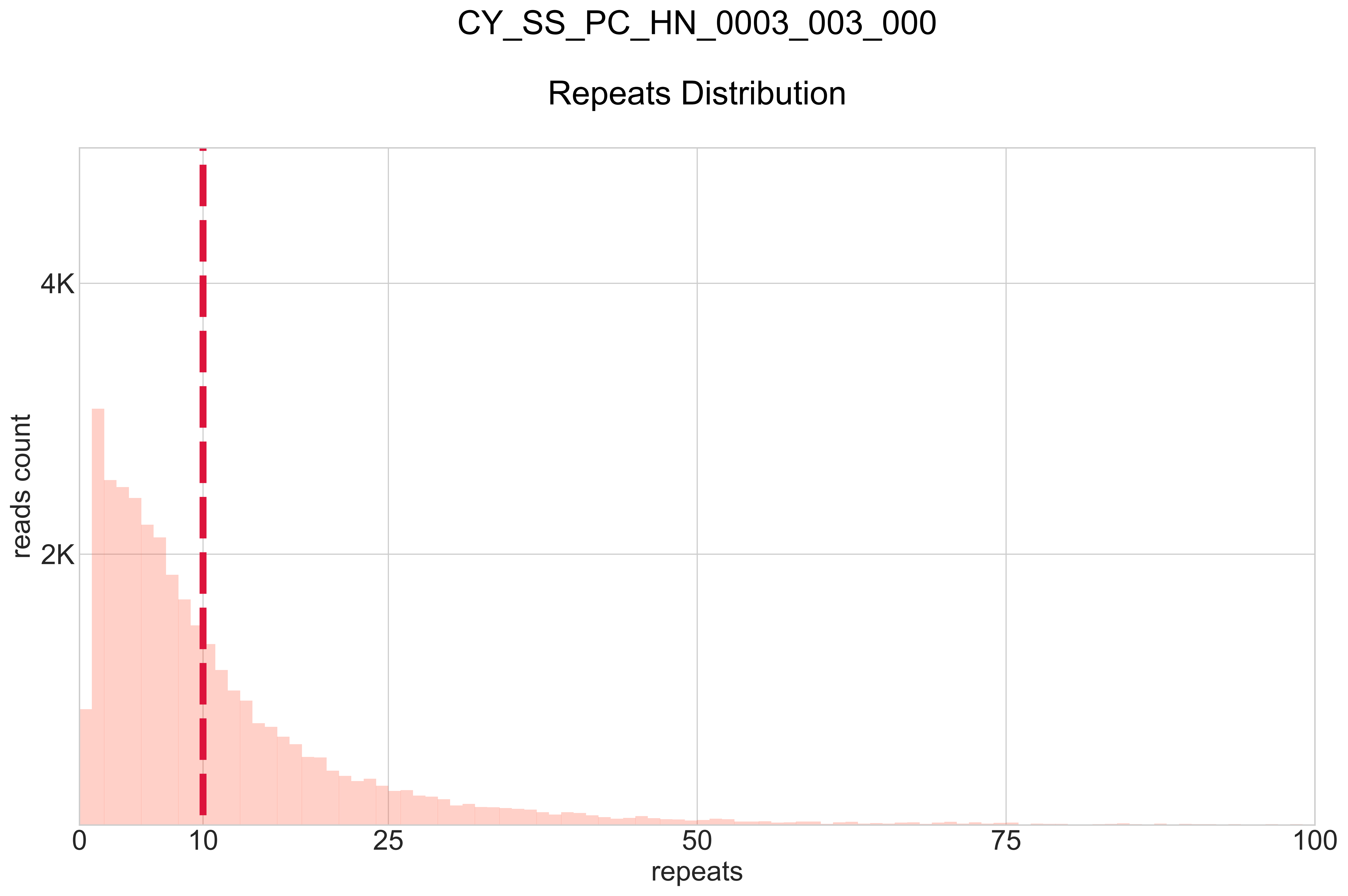

### CY_SS_PC_HN_0003_004_000_BB-I_.pdf

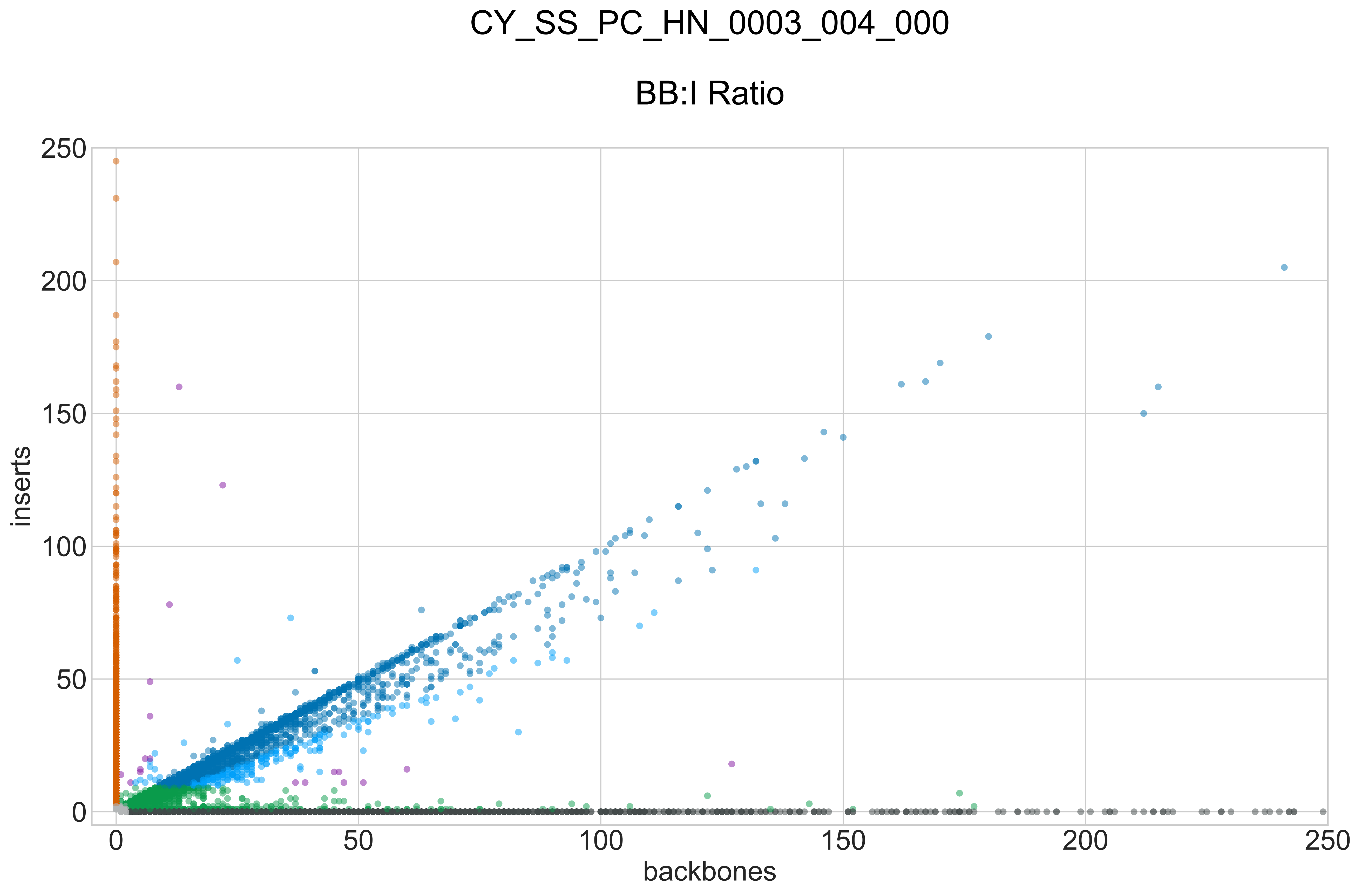

### CY_SS_PC_HN_0003_004_000_GROUPS_STACKED_NORM-LEN_.pdf

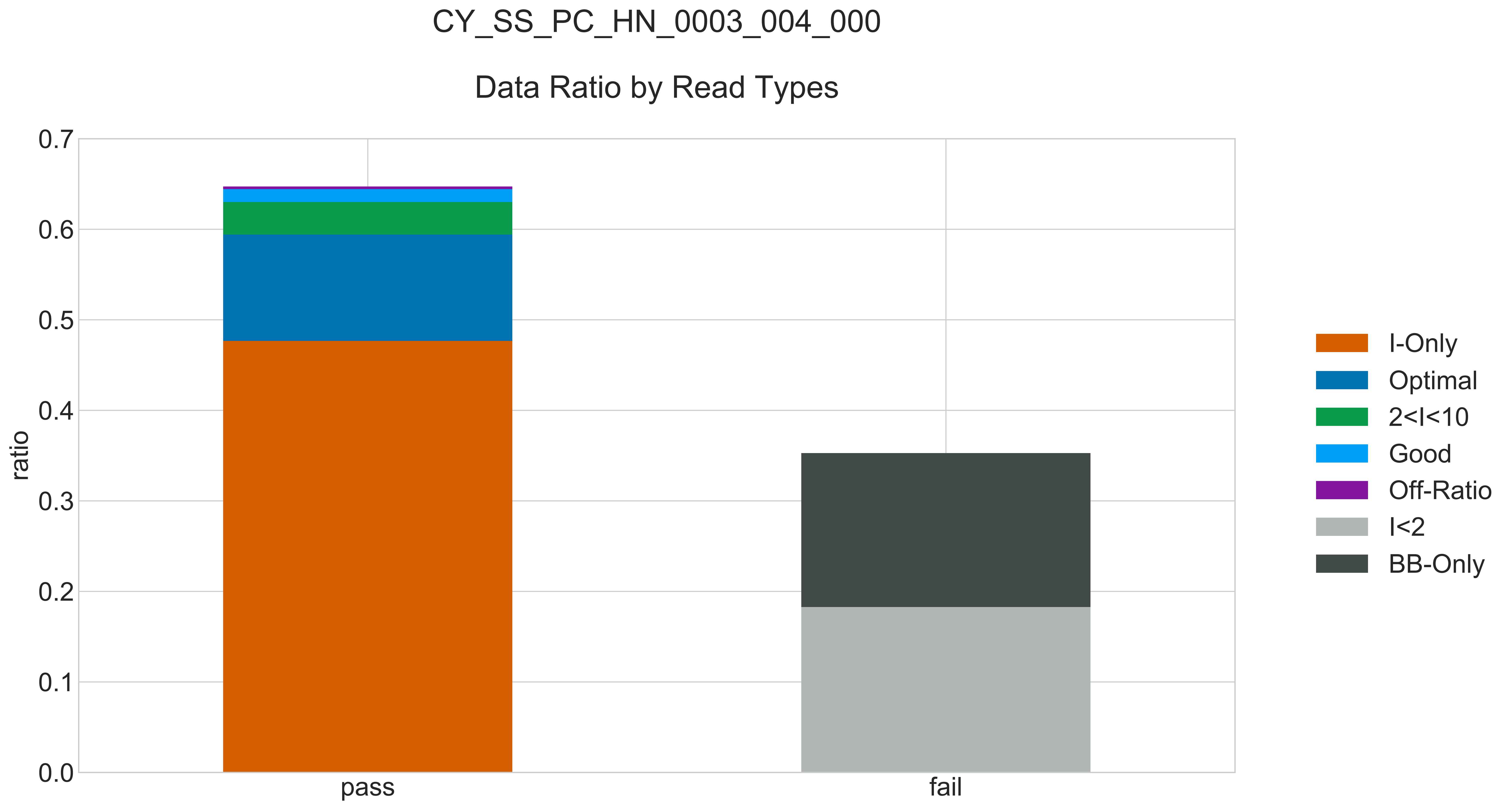

### CY_SS_PC_HN_0003_004_000_READ_LEN_.pdf

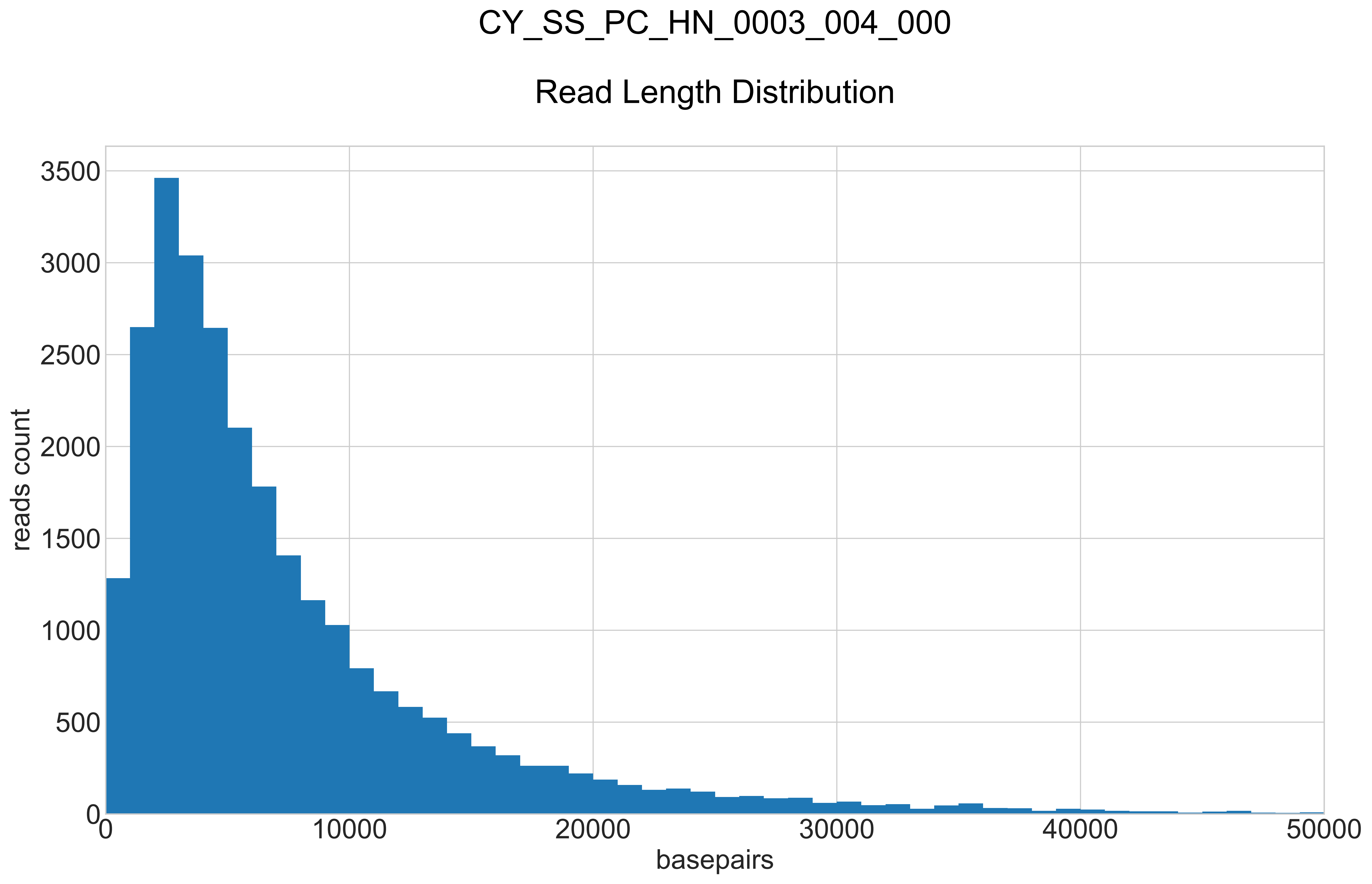

### CY_SS_PC_HN_0003_004_000_REPEATS_COUNT_.pdf

# CY_SS_PC_HN_0003_004_000

#### Repeats Distribution

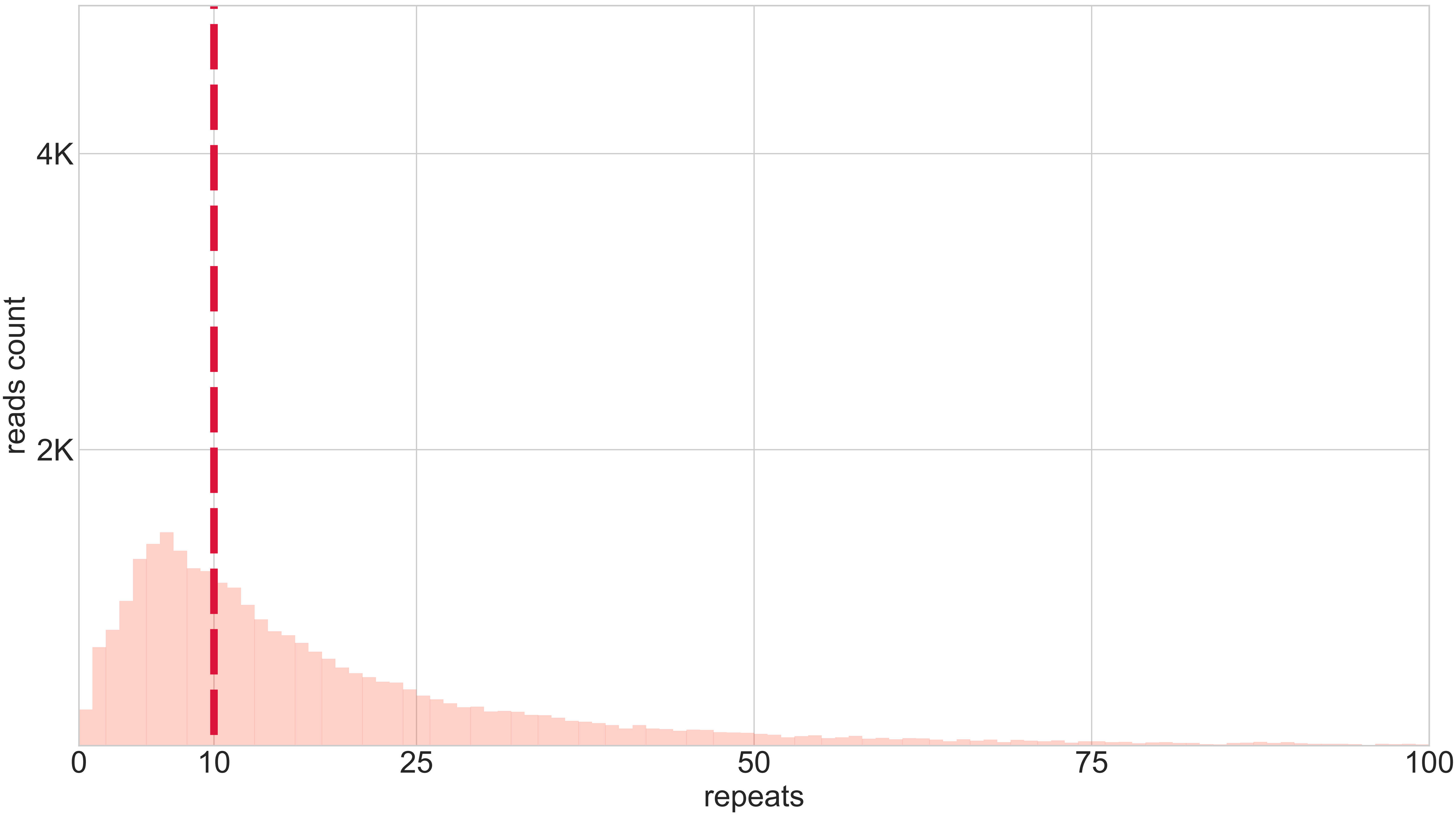

### CY_SS_PC_HN_0003_005_000_BB-I_.pdf

# CY_SS_PC_HN_0003_005_000

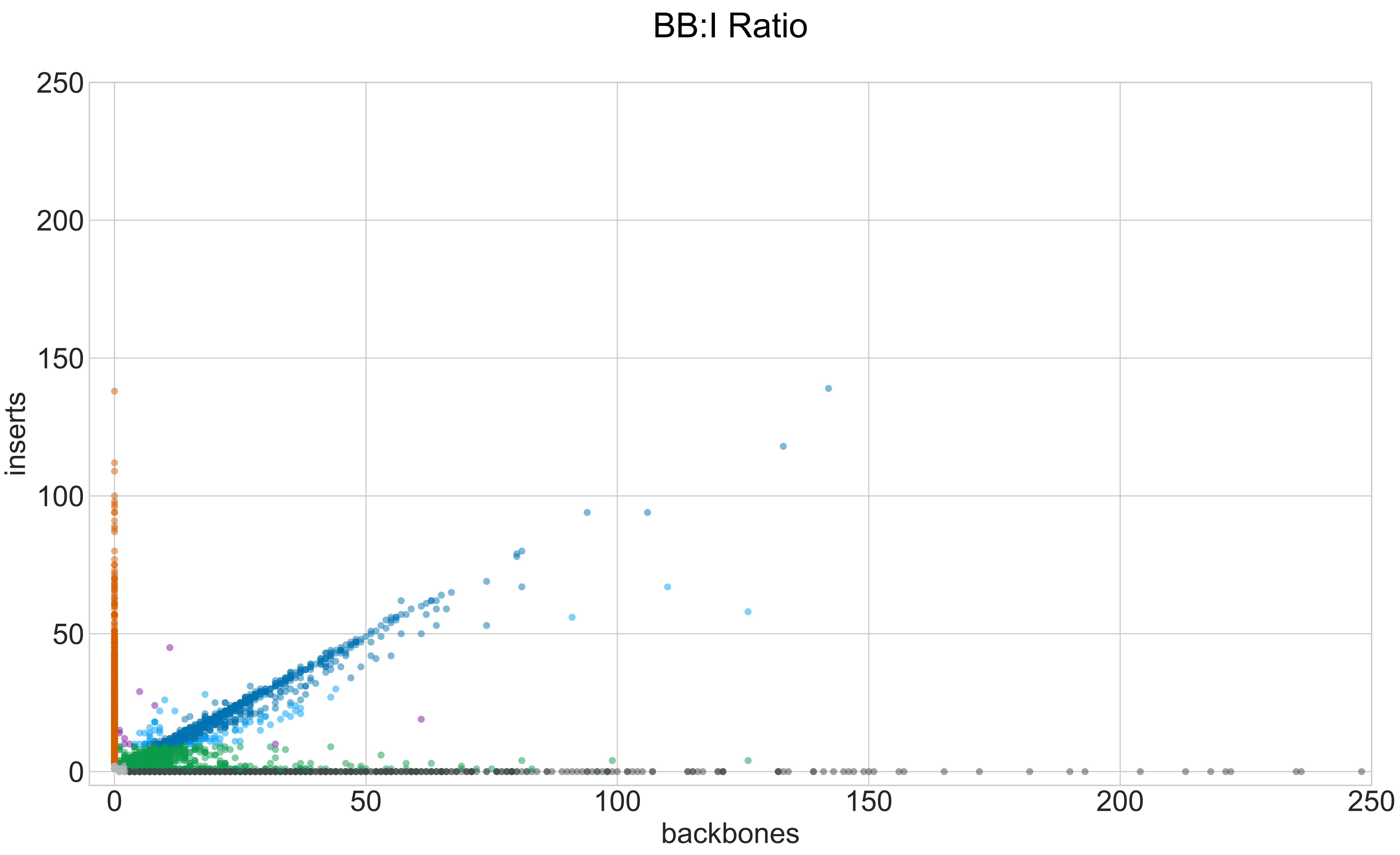

### CY_SS_PC_HN_0003_005_000_GROUPS_STACKED_NORM-LEN_.pdf

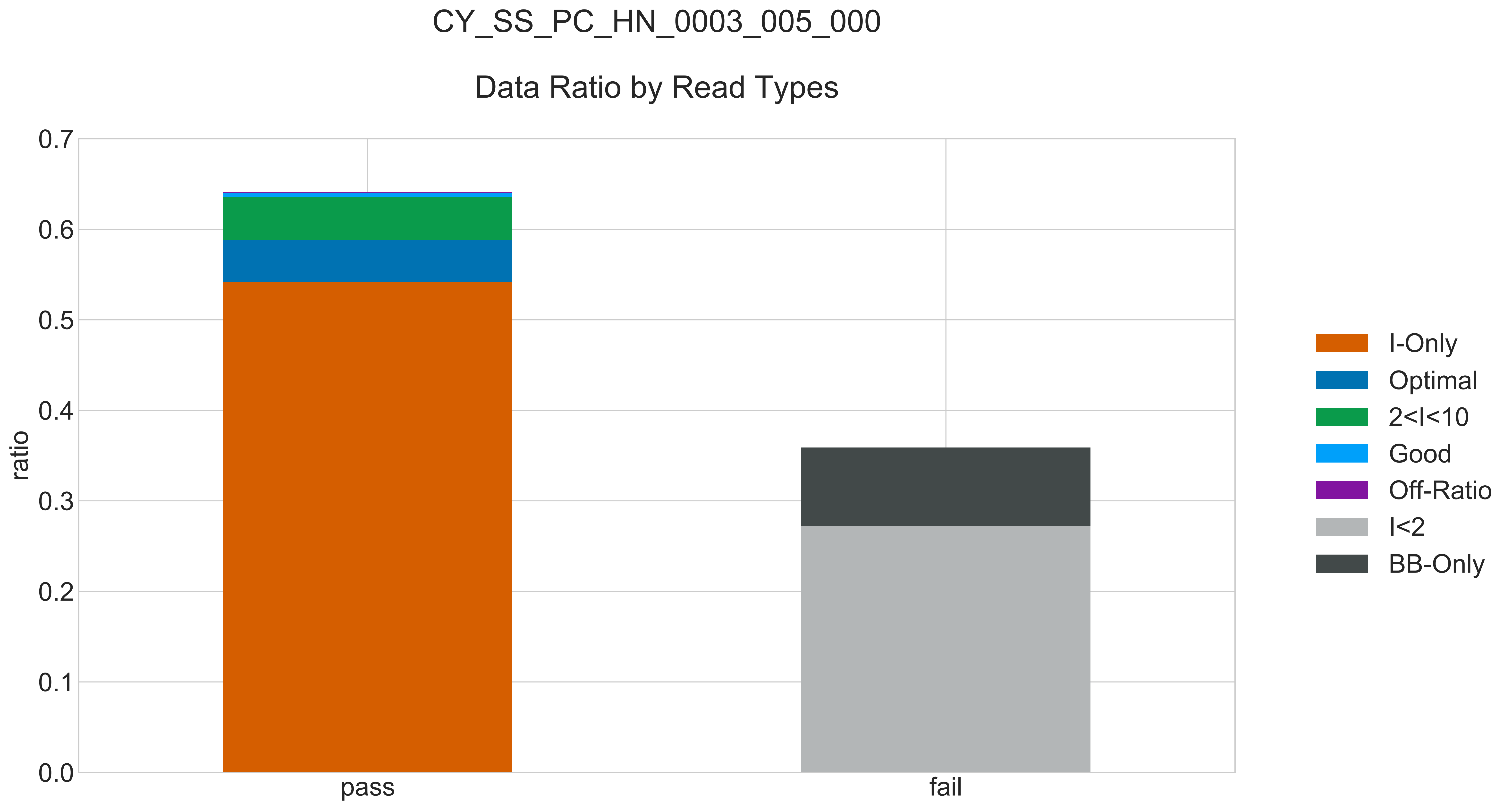

### CY_SS_PC_HN_0003_005_000_READ_LEN_.pdf

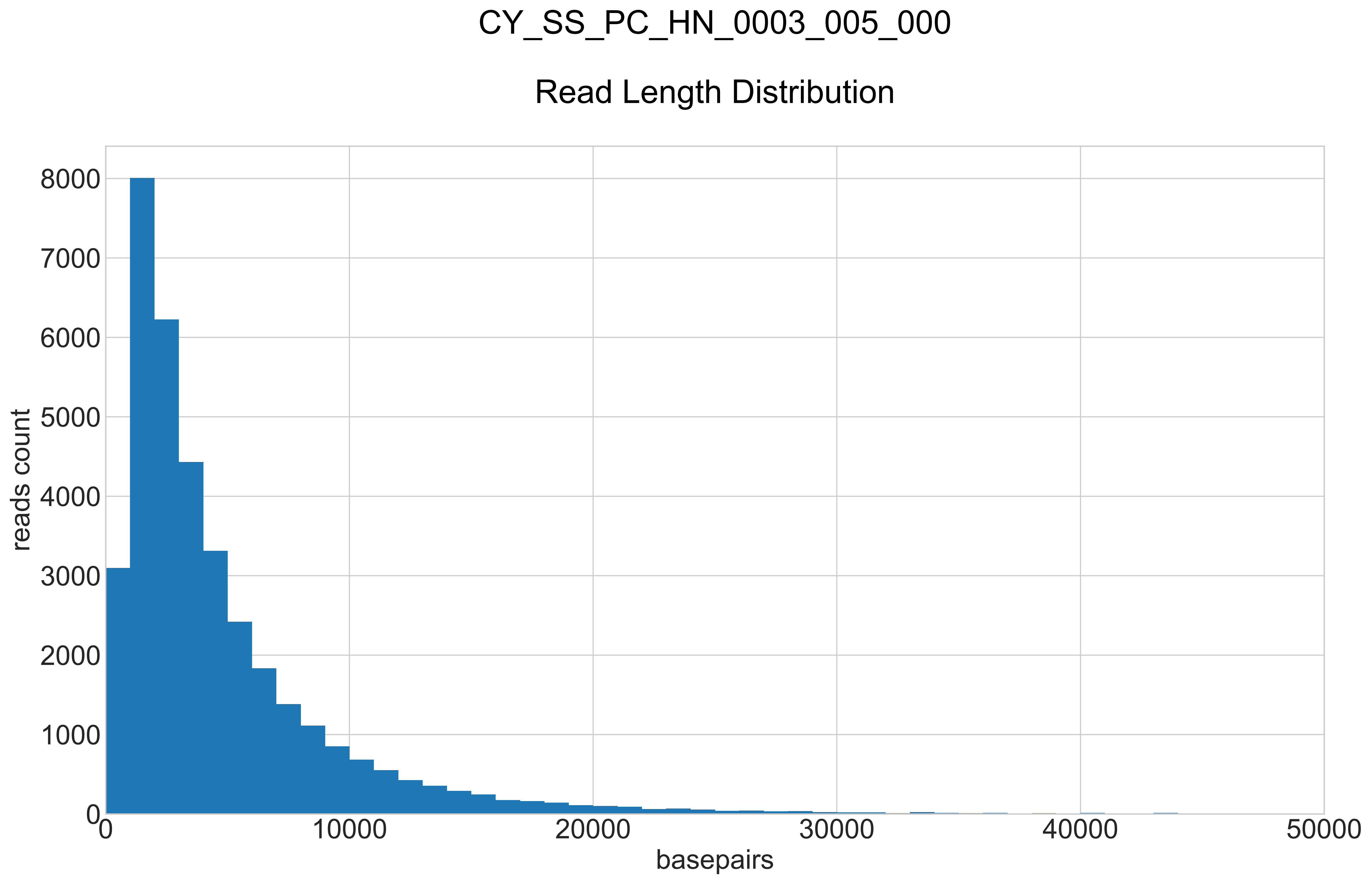

### CY_SS_PC_HN_0003_005_000_REPEATS_COUNT_.pdf

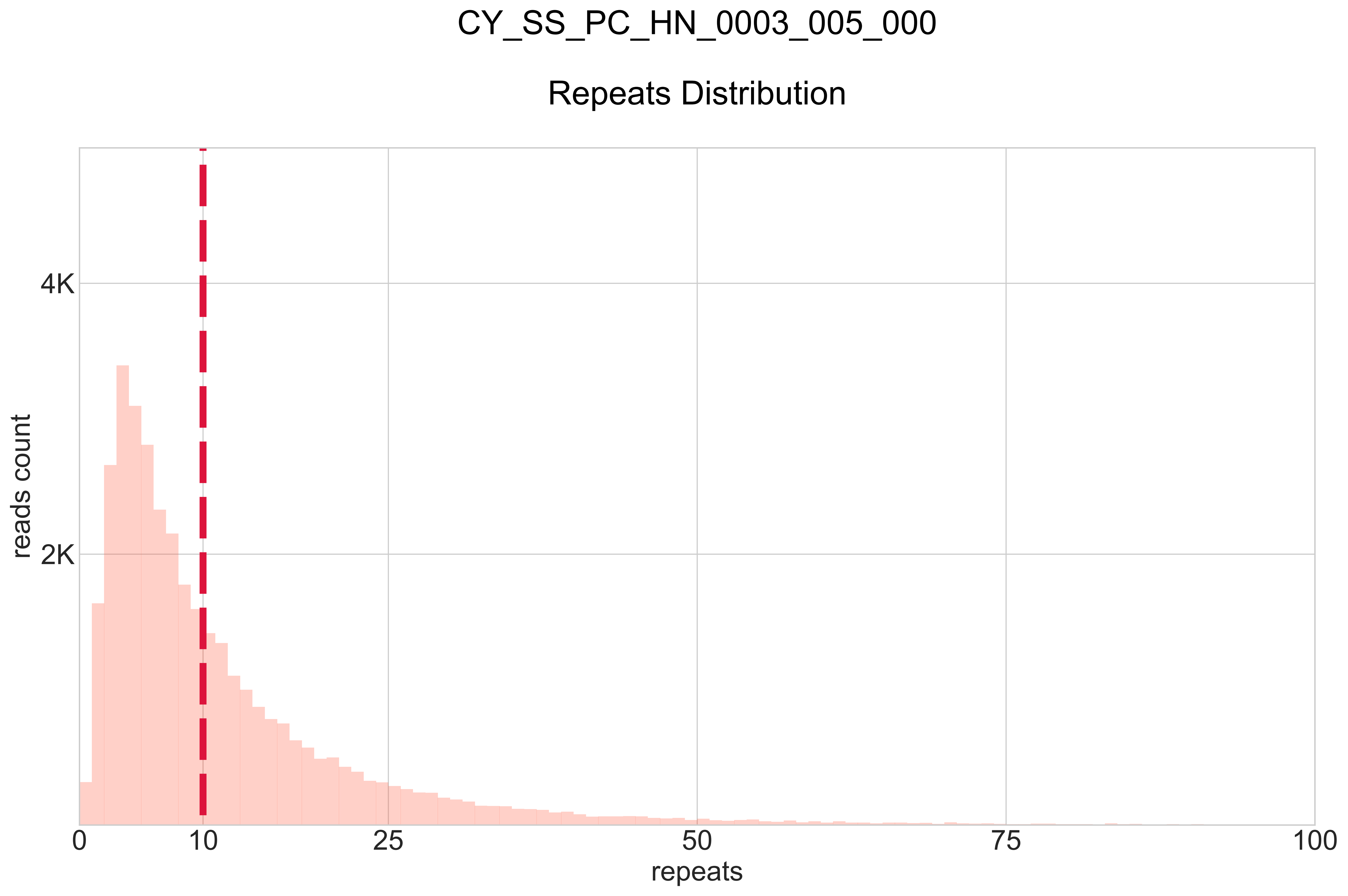

### CY_SS_PC_HN_0003_006_000_BB-I_.pdf

# CY_SS_PC_HN_0003_006_000

#### BB:I Ratio

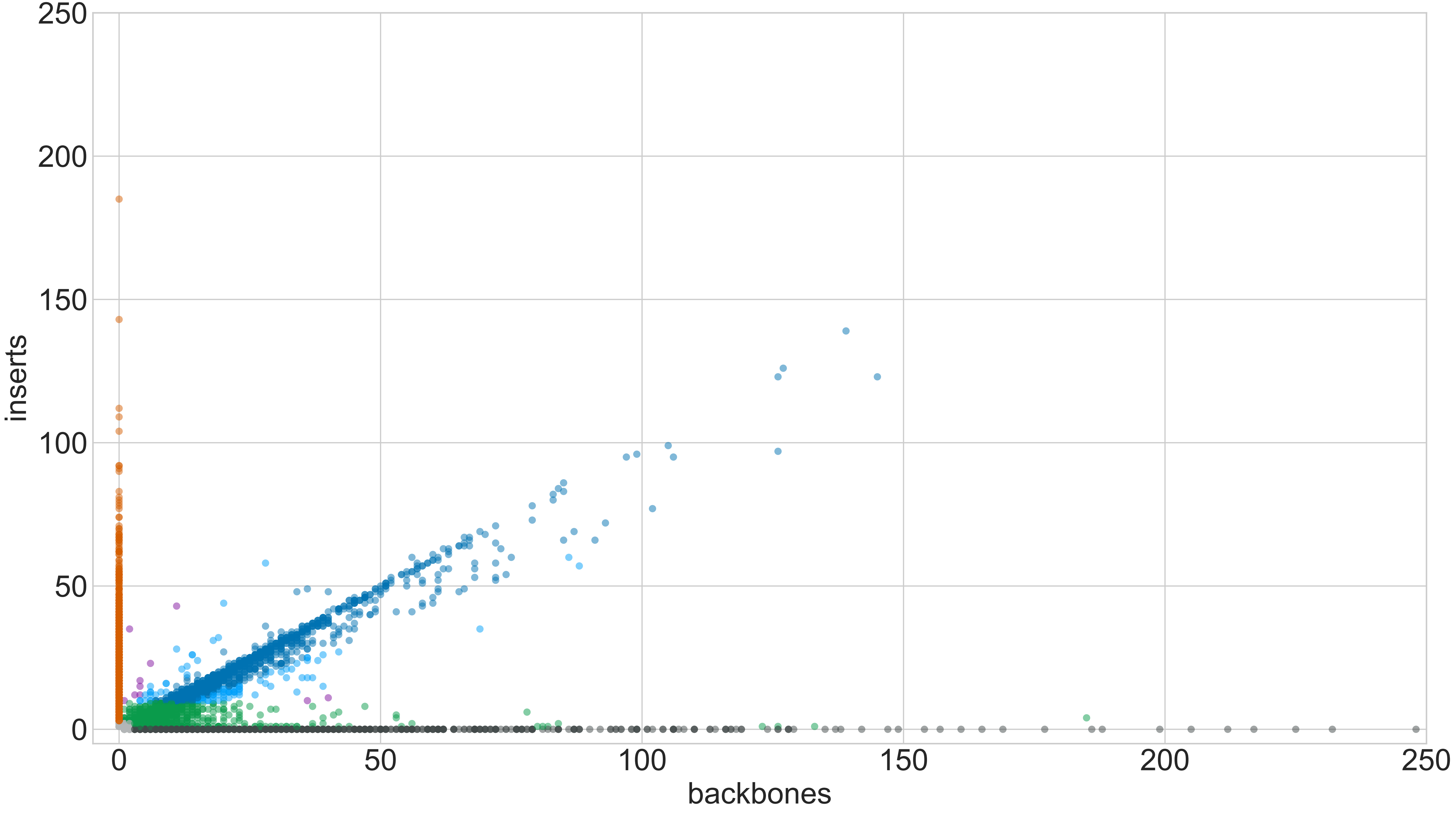

### CY_SS_PC_HN_0003_006_000_GROUPS_STACKED_NORM-LEN_.pdf

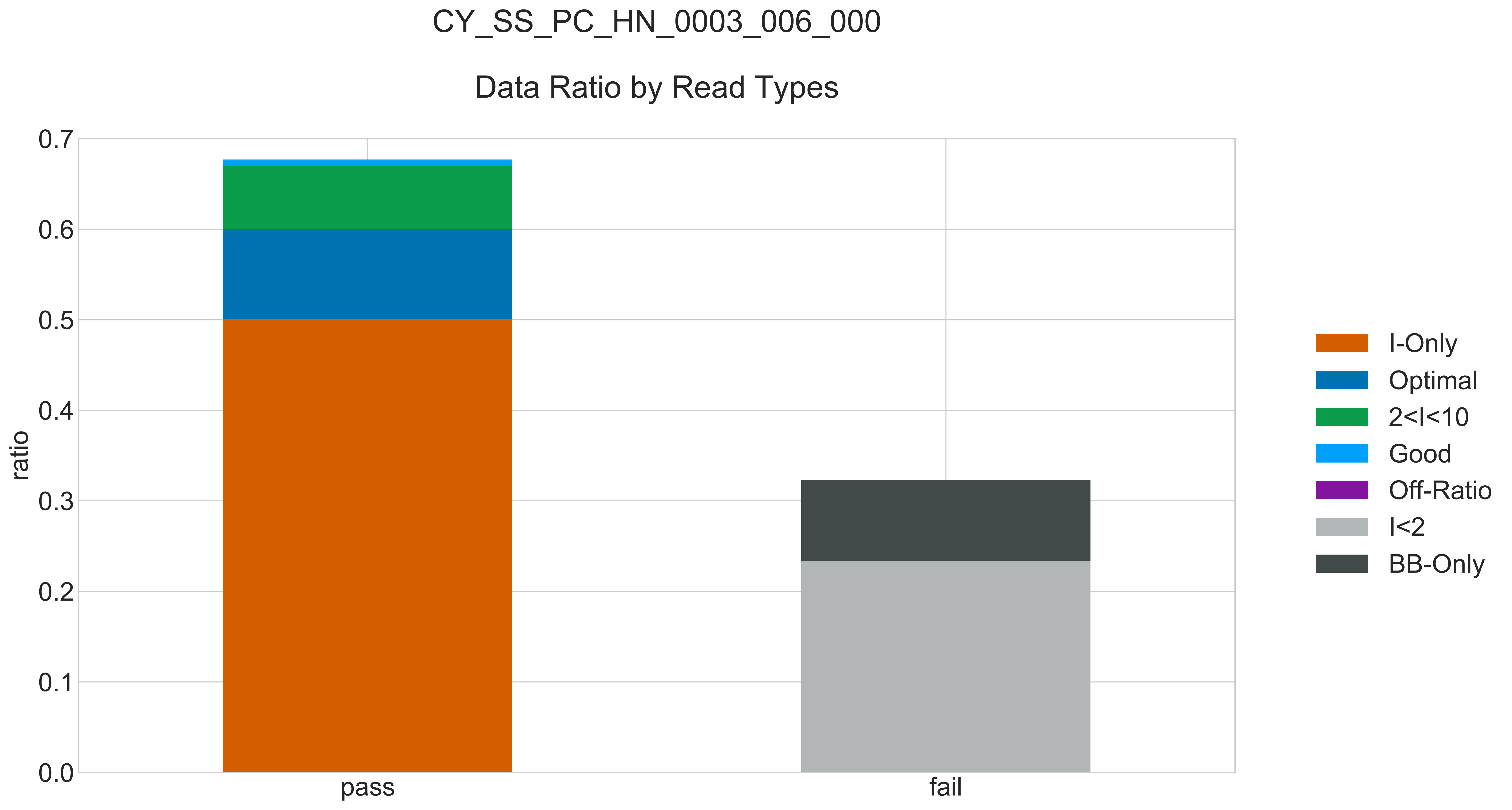

### CY_SS_PC_HN_0003_006_000_READ_LEN_.pdf

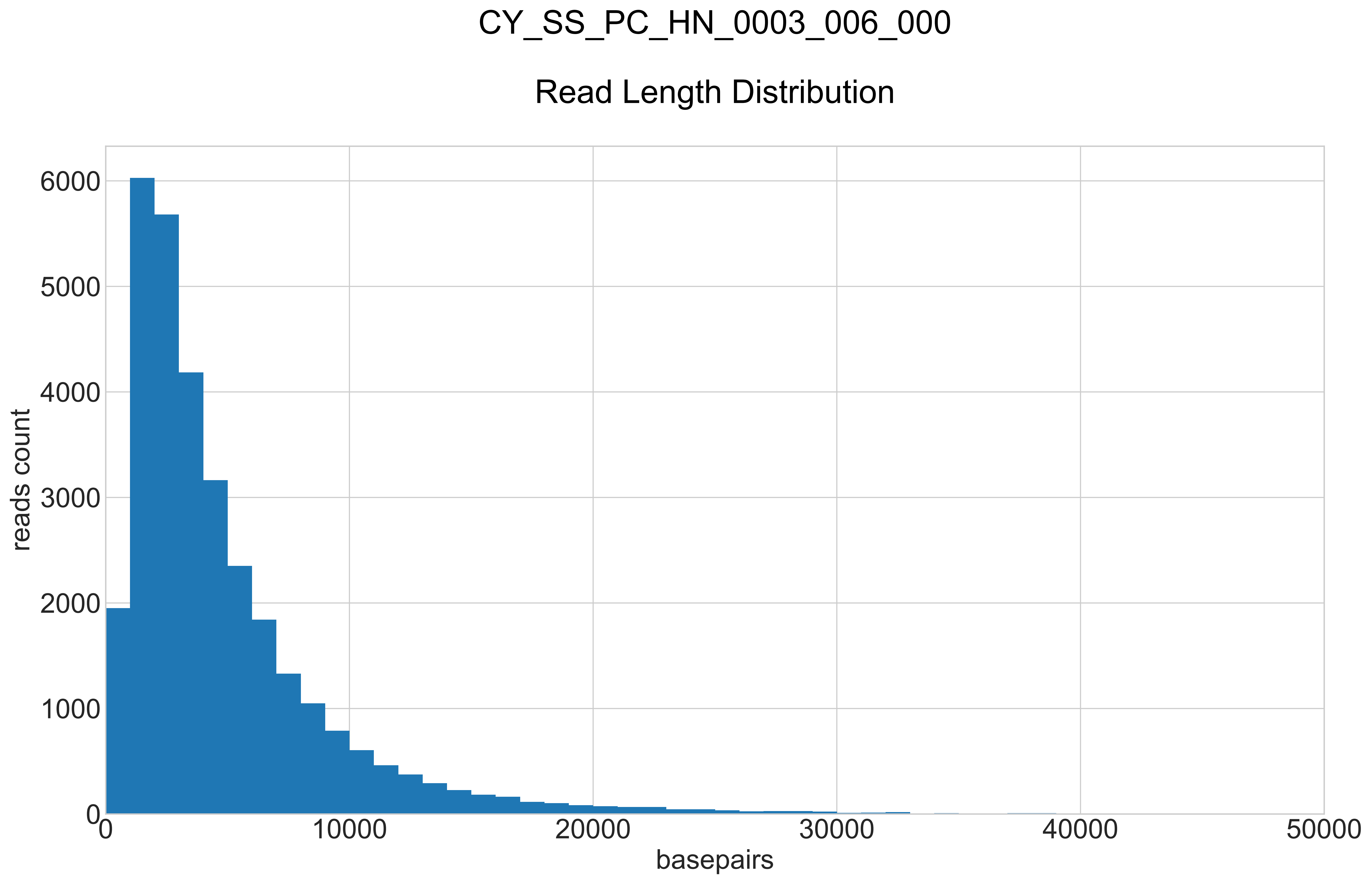

### CY_SS_PC_HN_0003_006_000_REPEATS_COUNT_.pdf

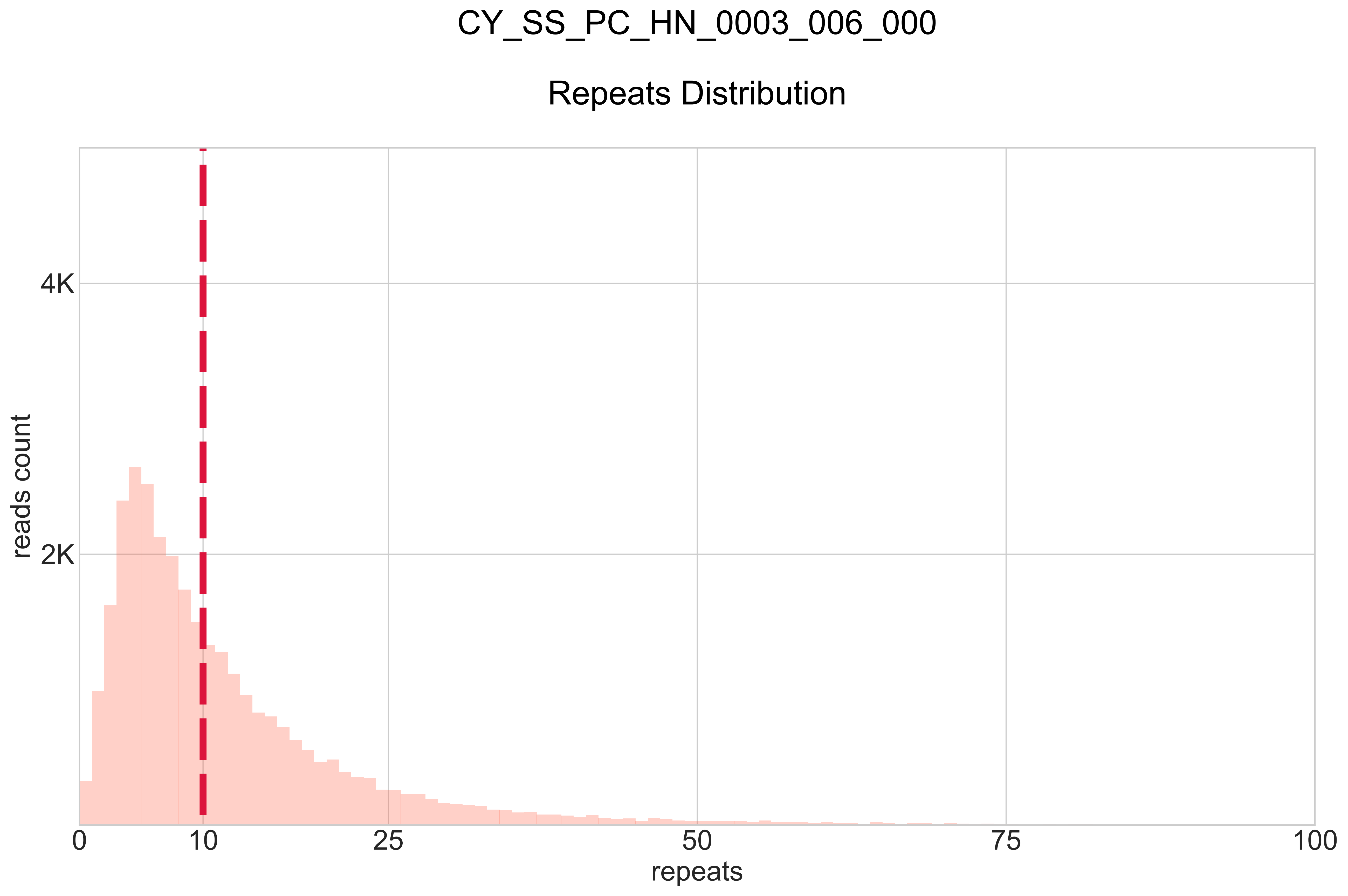

### CY_SS_SC_HN_0001_006_000_BB-I_.pdf

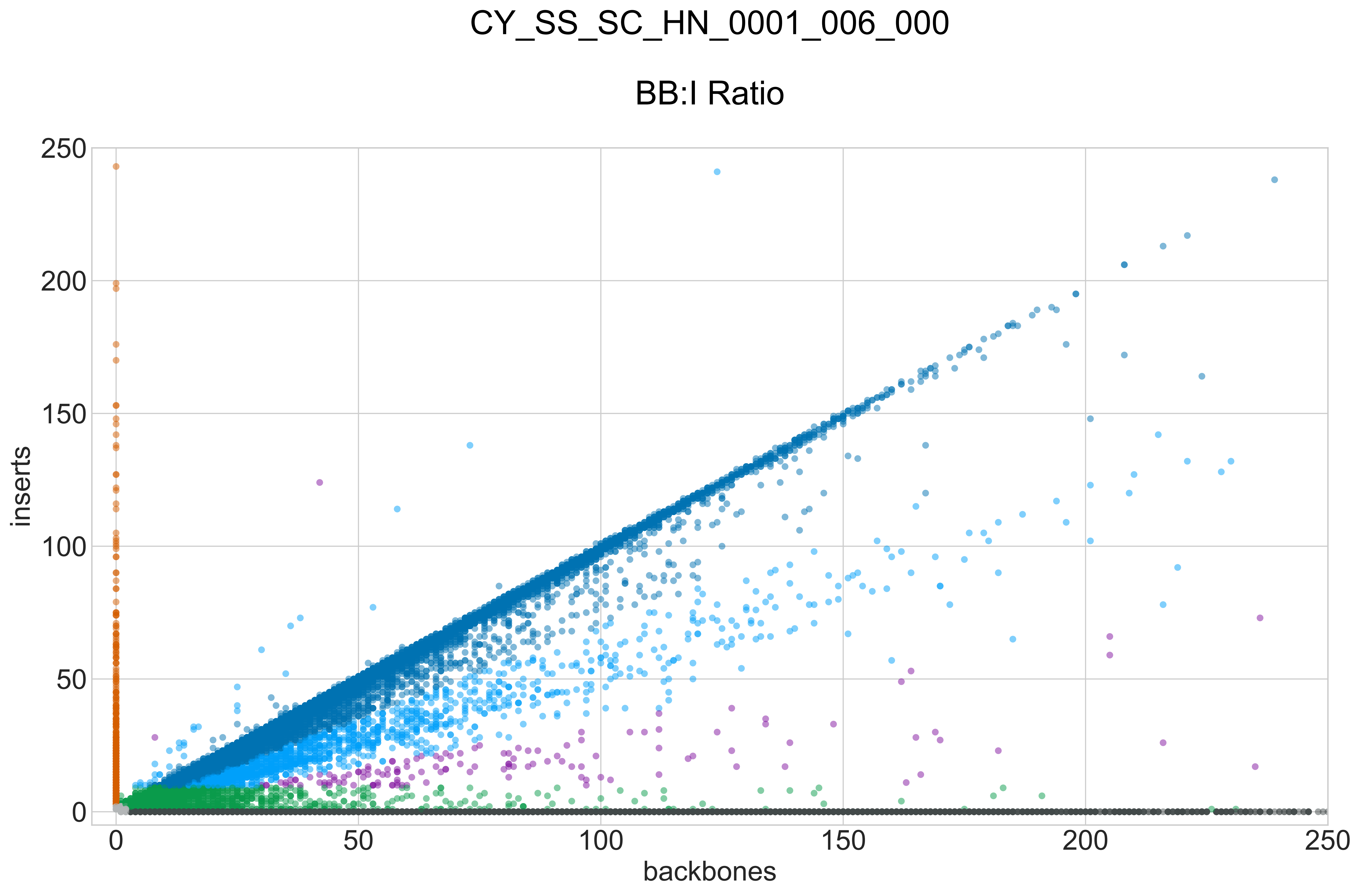

### CY_SS_SC_HN_0001_006_000_GROUPS_STACKED_NORM-LEN_.pdf

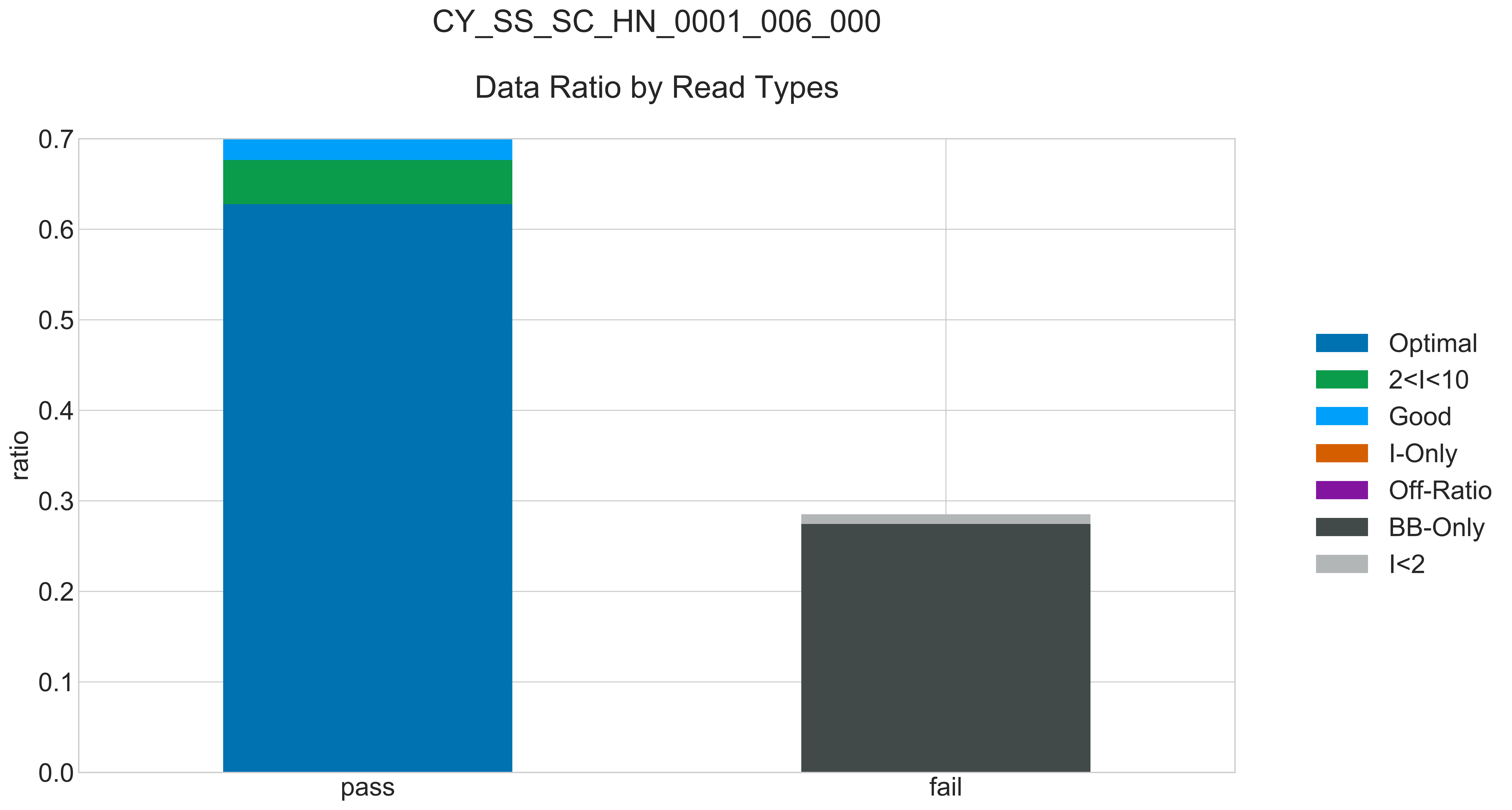

### CY_SS_SC_HN_0001_006_000_READ_LEN_.pdf

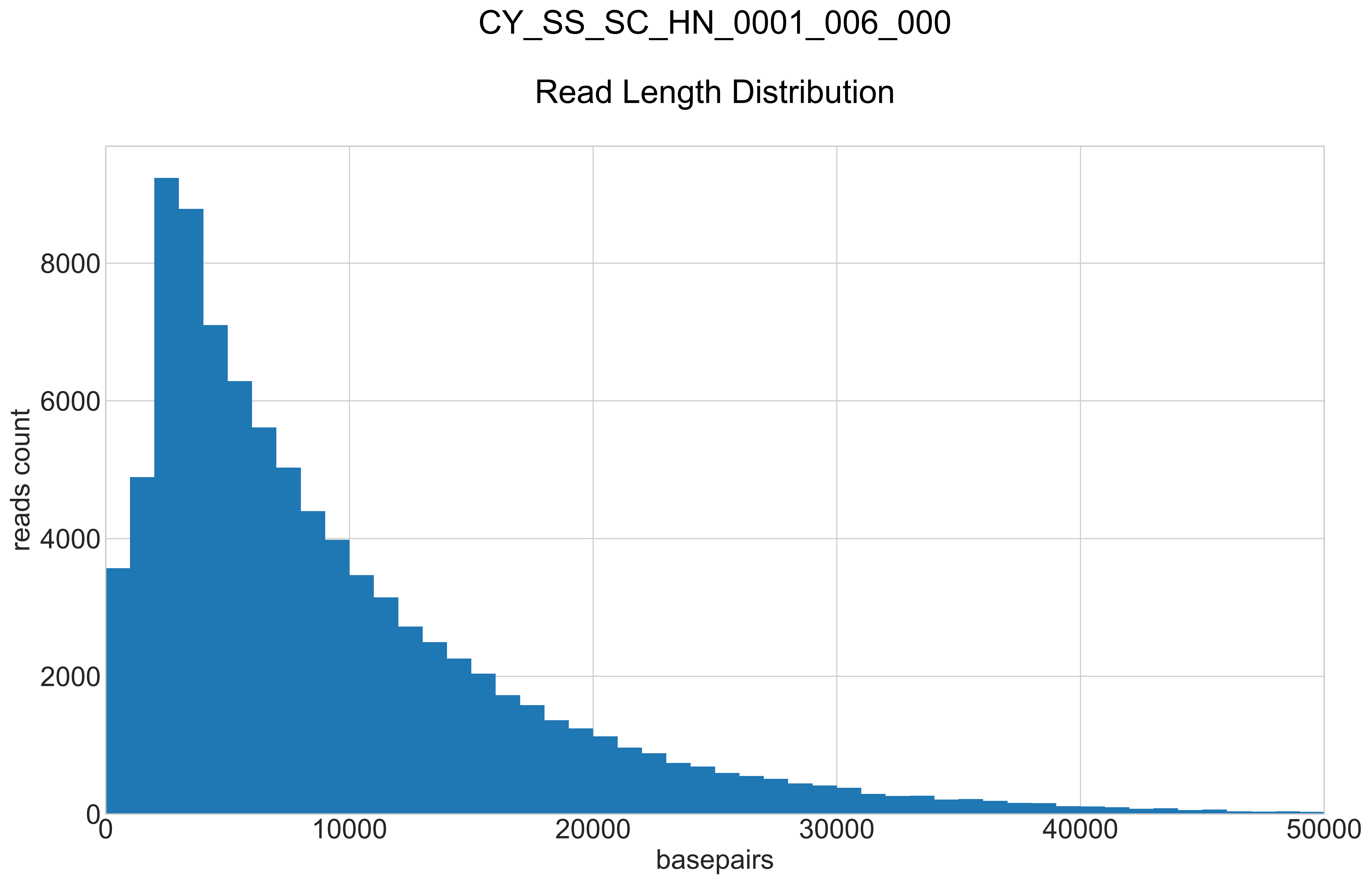

### CY_SS_SC_HN_0001_006_000_REPEATS_COUNT_.pdf

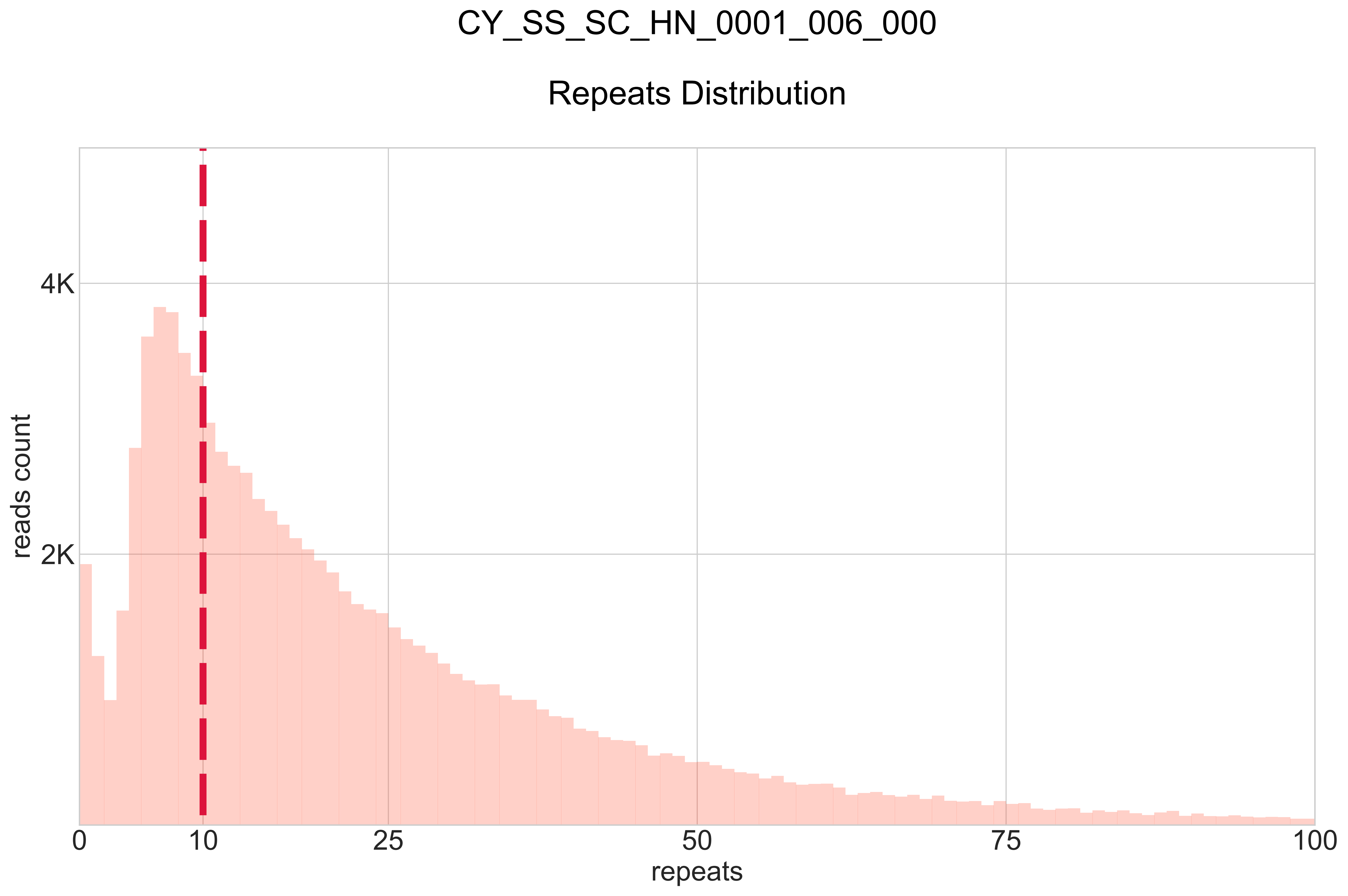

### Supplementary Figure 2

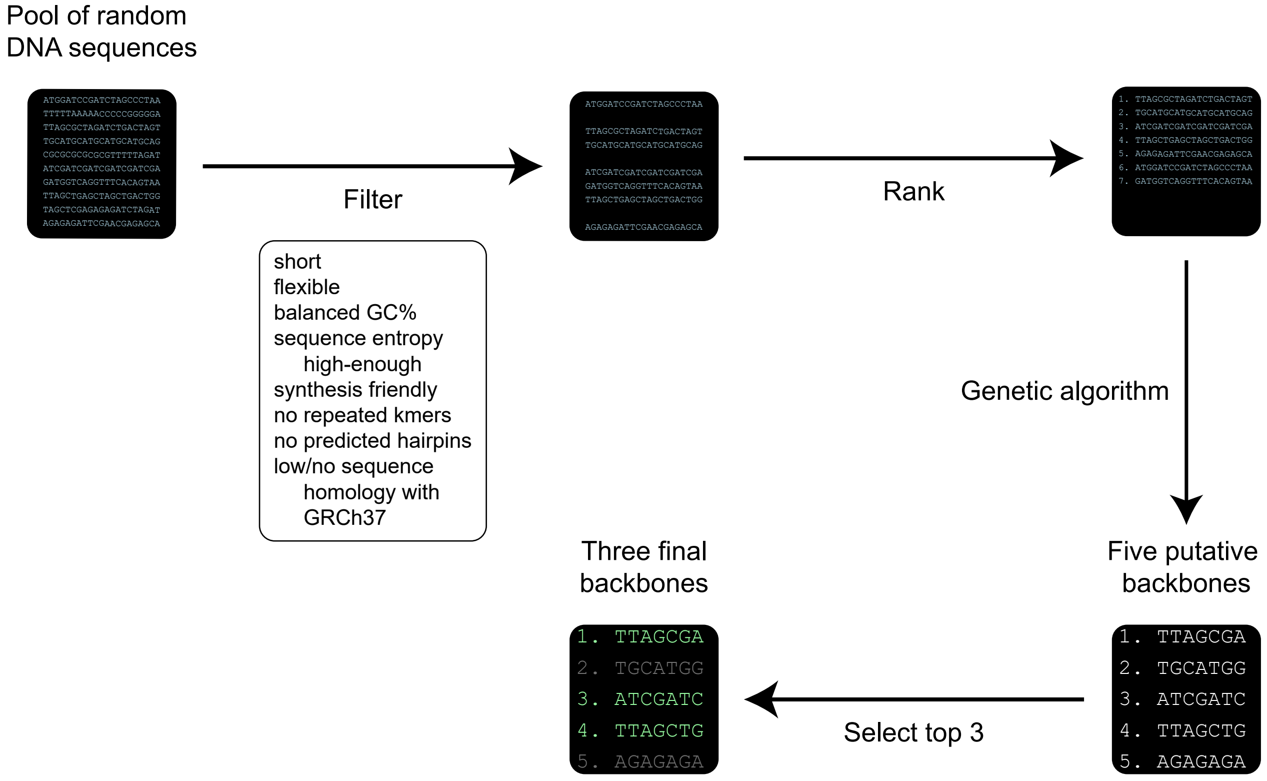


**Supplementary Fig. 2** Approach to design the three backbones used: BB22, BB24 and BB25.
