## Supplementary figures and images for "Accurate detection of circulating tumor DNA using nanopore consensus sequencing"

### CY_BB25_19WT_0001_000_BB-I_.pdf

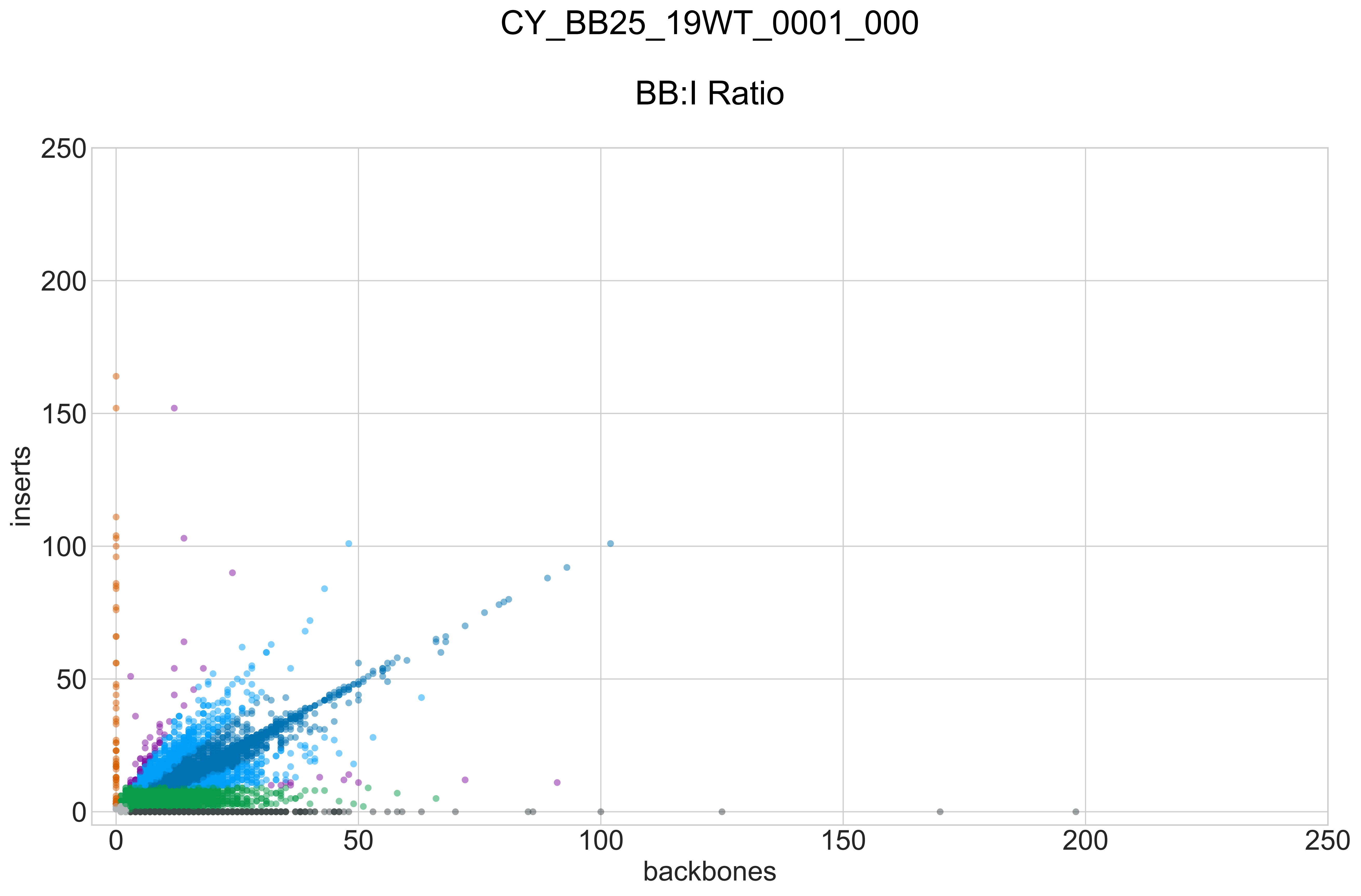

### CY_BB25_19WT_0001_000_GROUPS_STACKED_NORM-LEN_.pdf

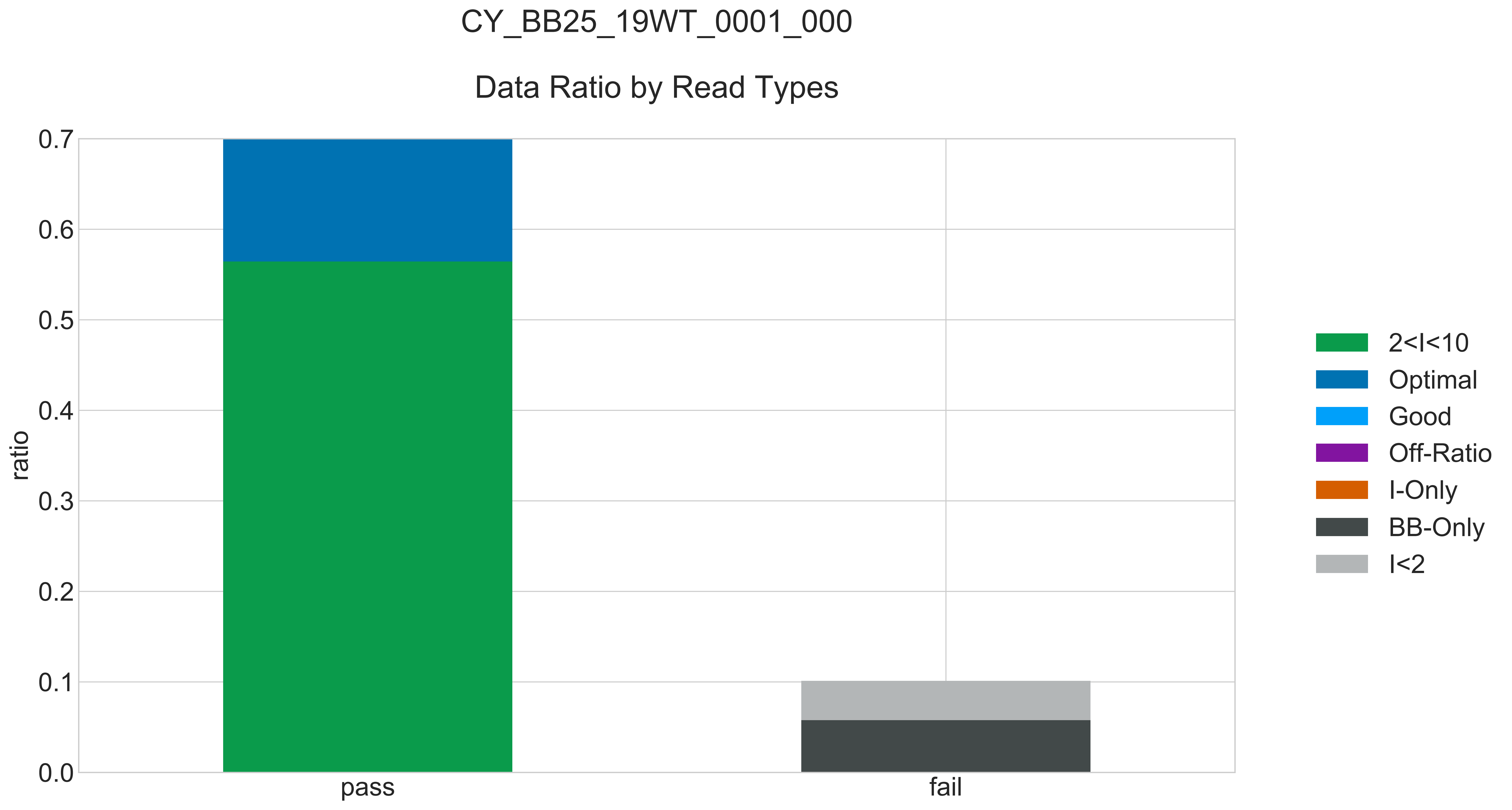

### CY_BB25_19WT_0001_000_READ_LEN_.pdf

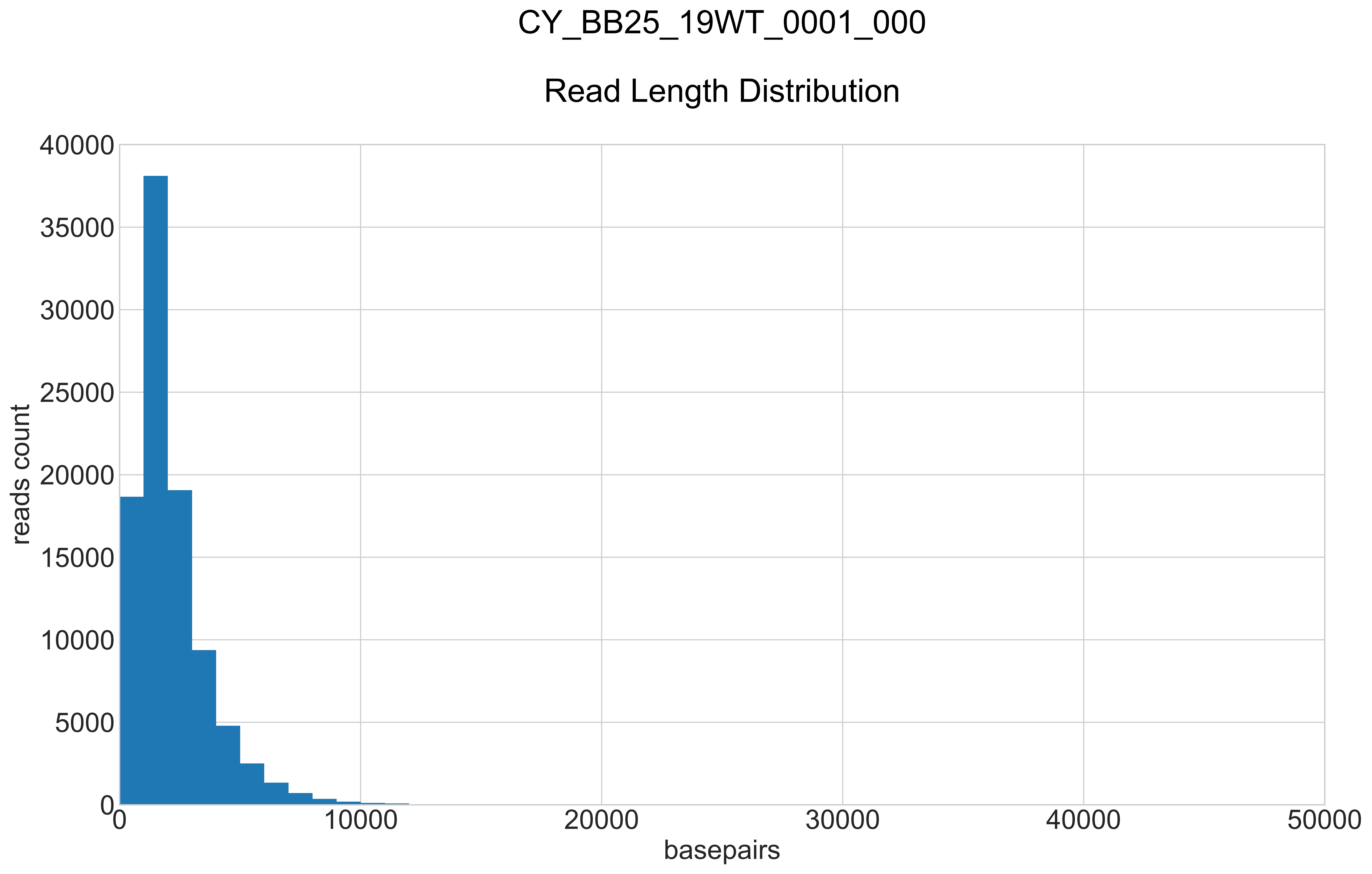

### CY_BB25_19WT_0001_000_REPEATS_COUNT_.pdf

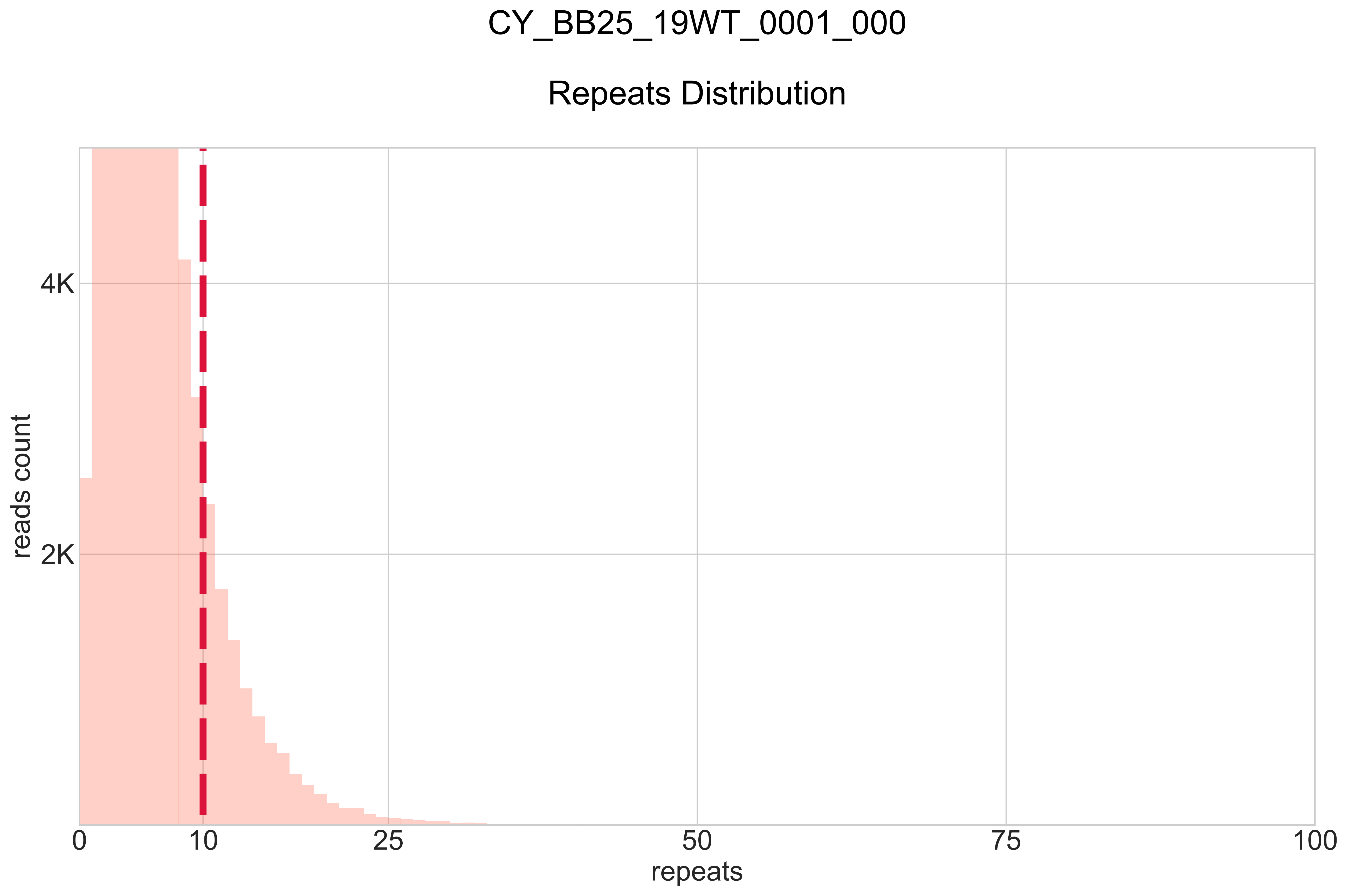

### CY_LOT1_QC_0001_000_BB-I_.pdf

#### CY_LOT1_QC_0001_000

##### BB:I Ratio

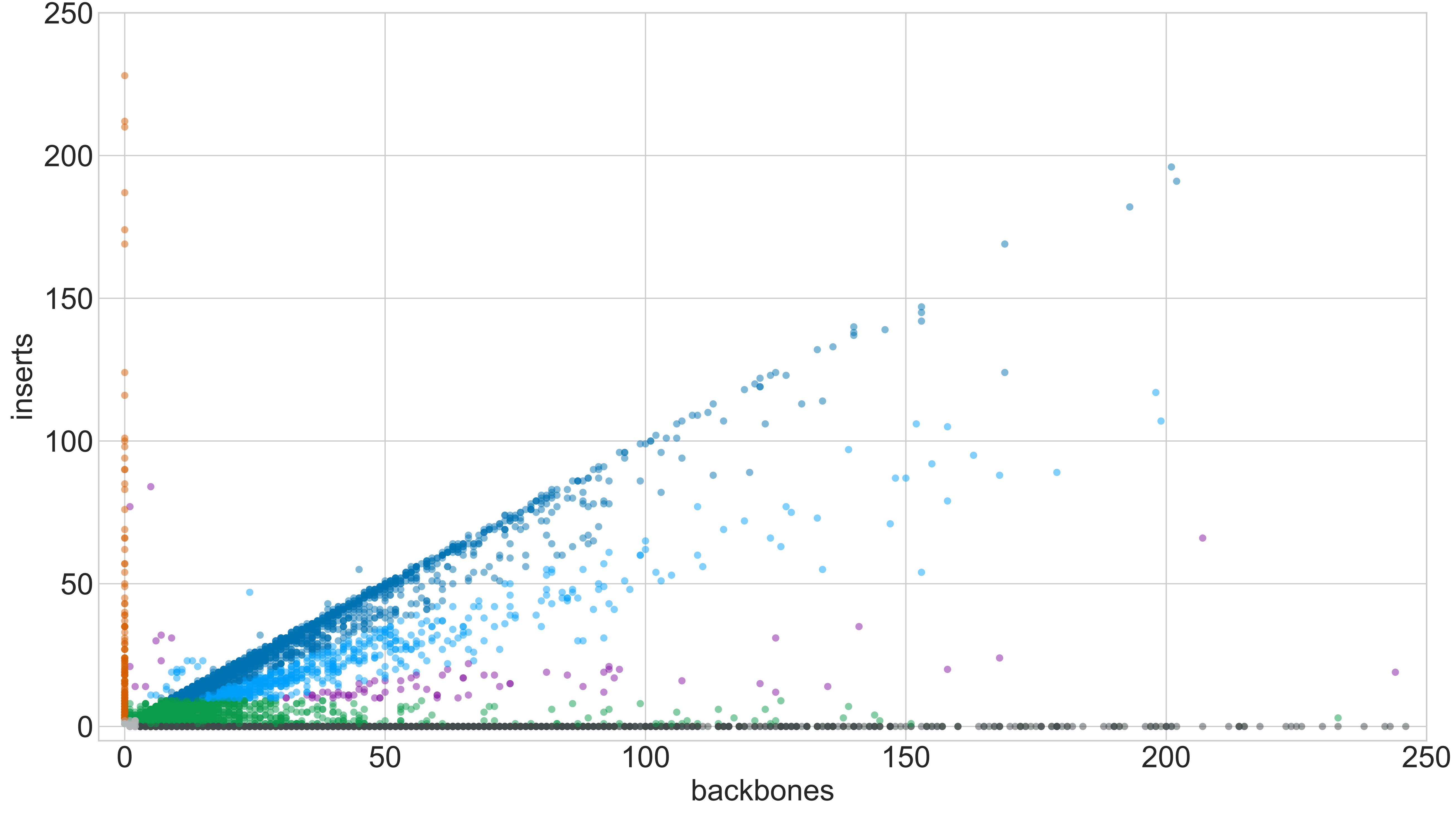

### CY_LOT1_QC_0001_000_GROUPS_STACKED_NORM-LEN_.pdf

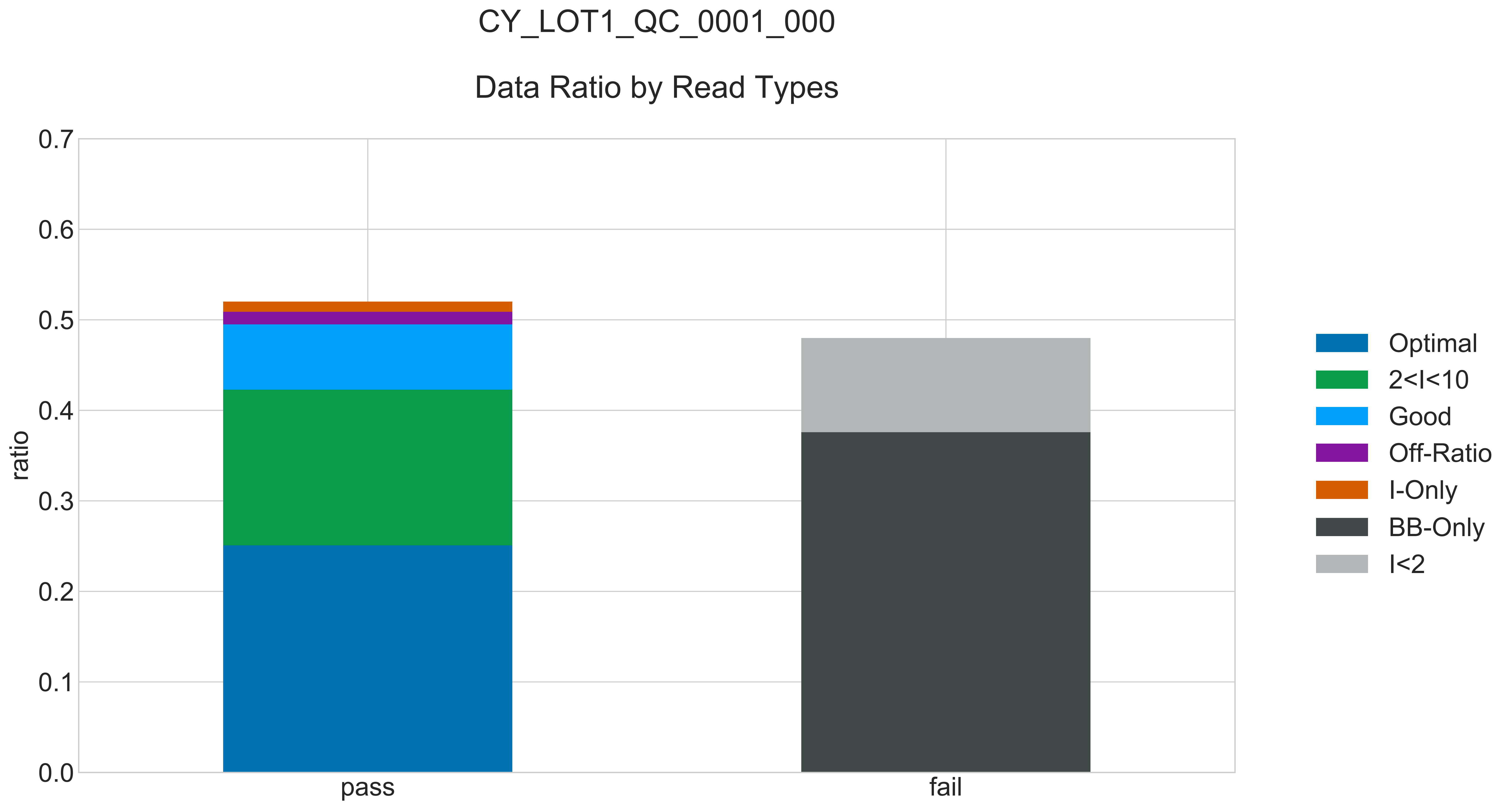

### CY_LOT1_QC_0001_000_READ_LEN_.pdf

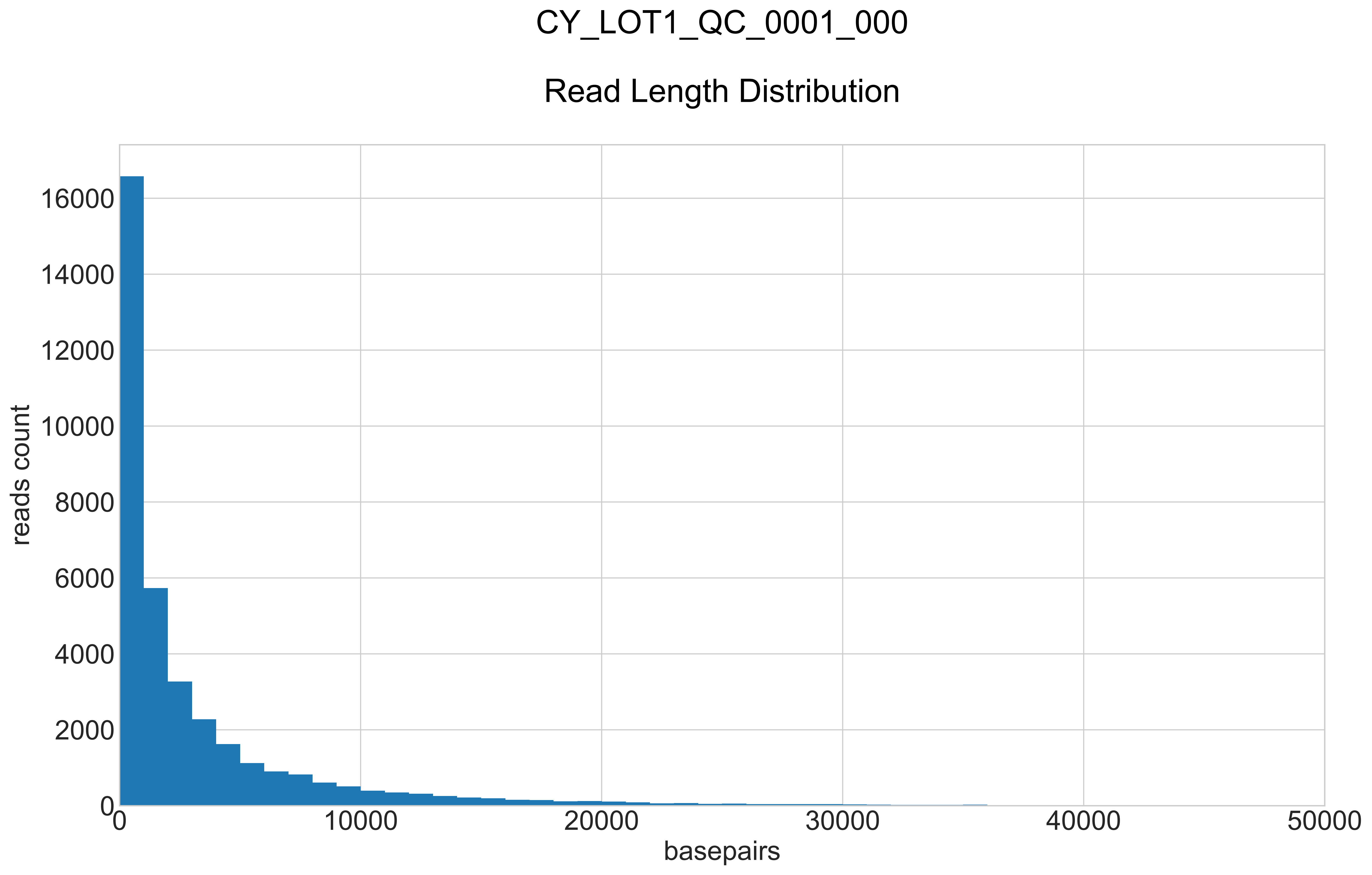

### CY_LOT1_QC_0001_000_REPEATS_COUNT_.pdf

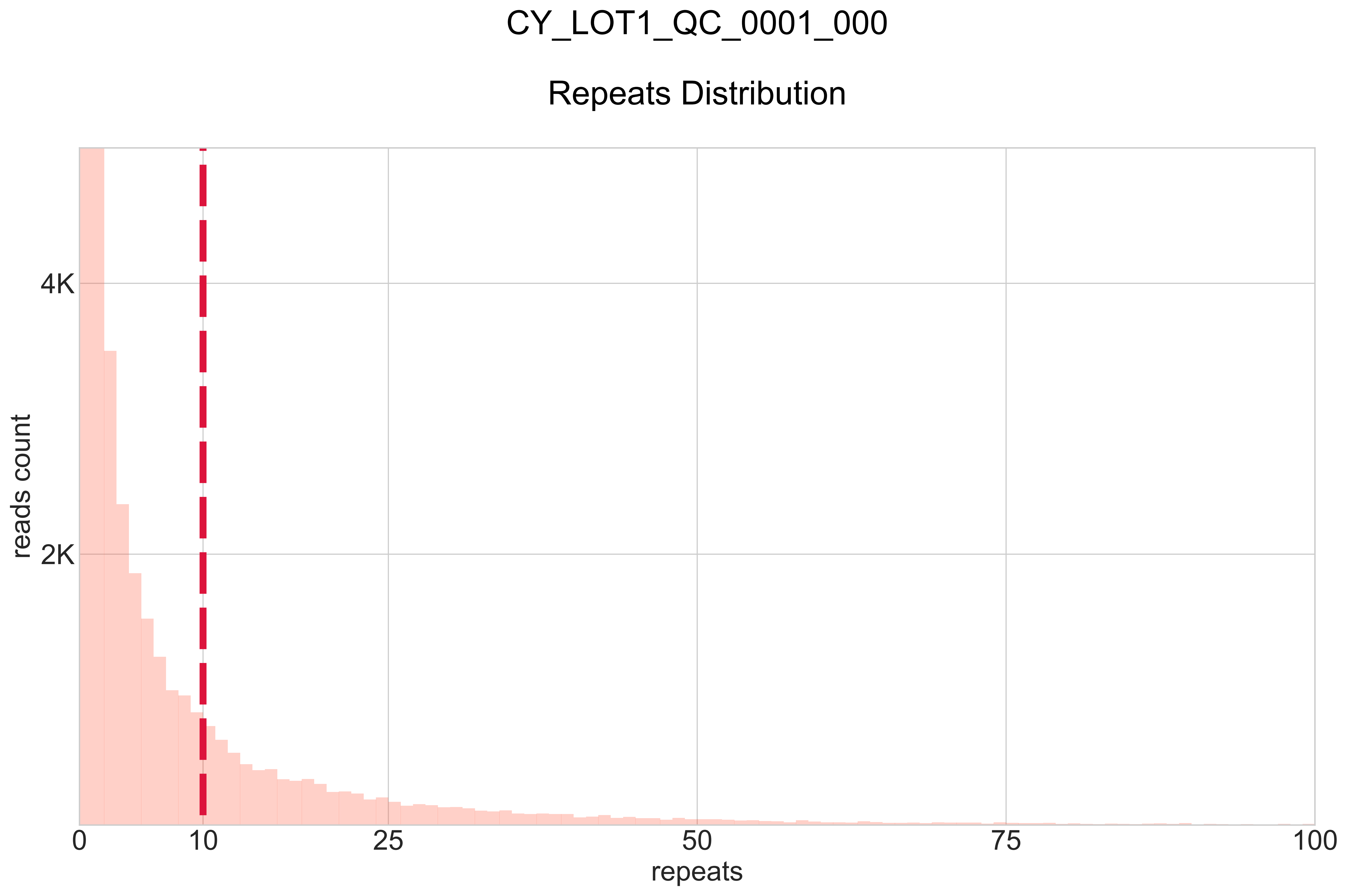

### CY_LOT1_QC_0001_001_BB-I_.pdf

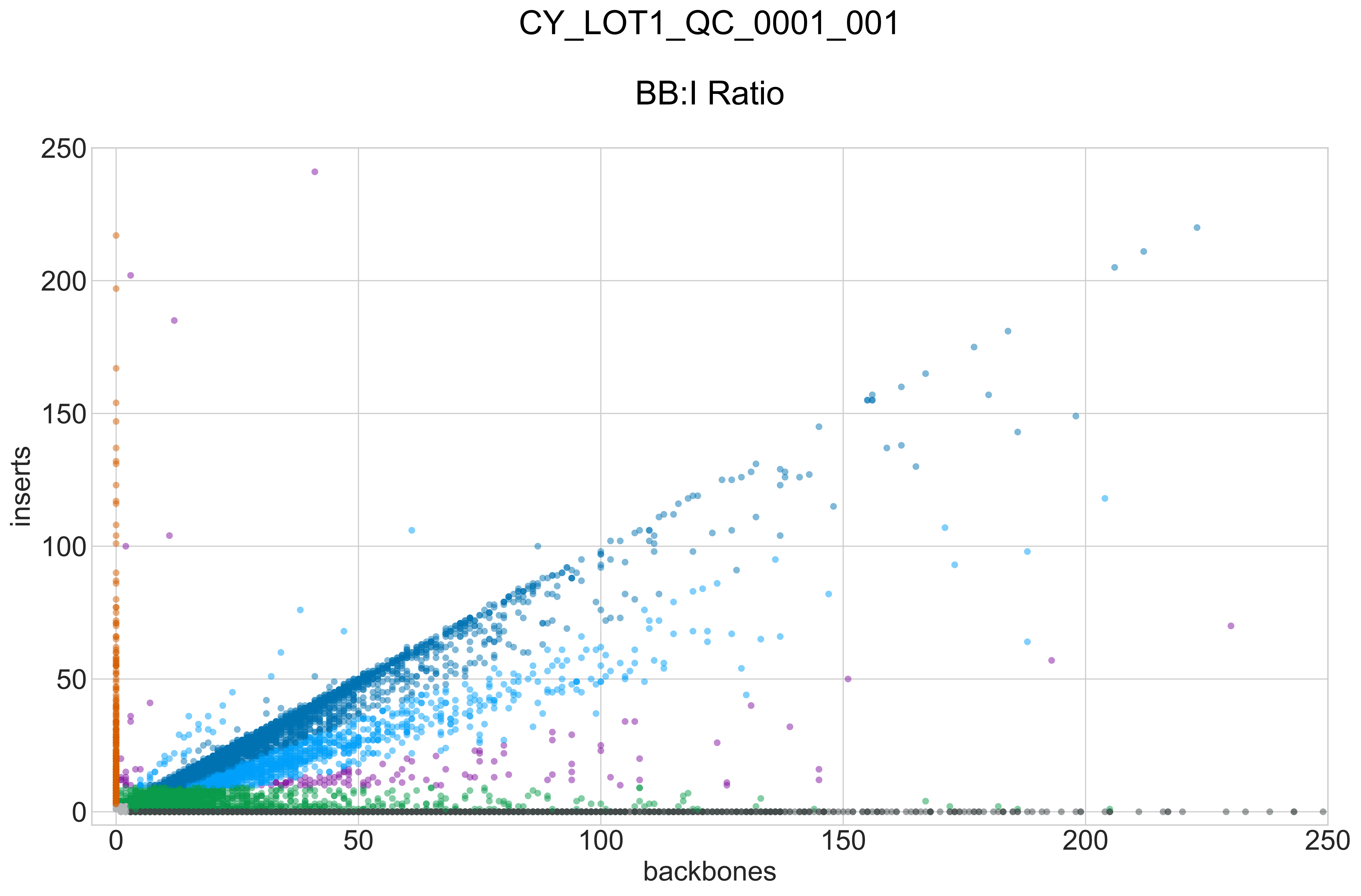

### CY_LOT1_QC_0001_001_GROUPS_STACKED_NORM-LEN_.pdf

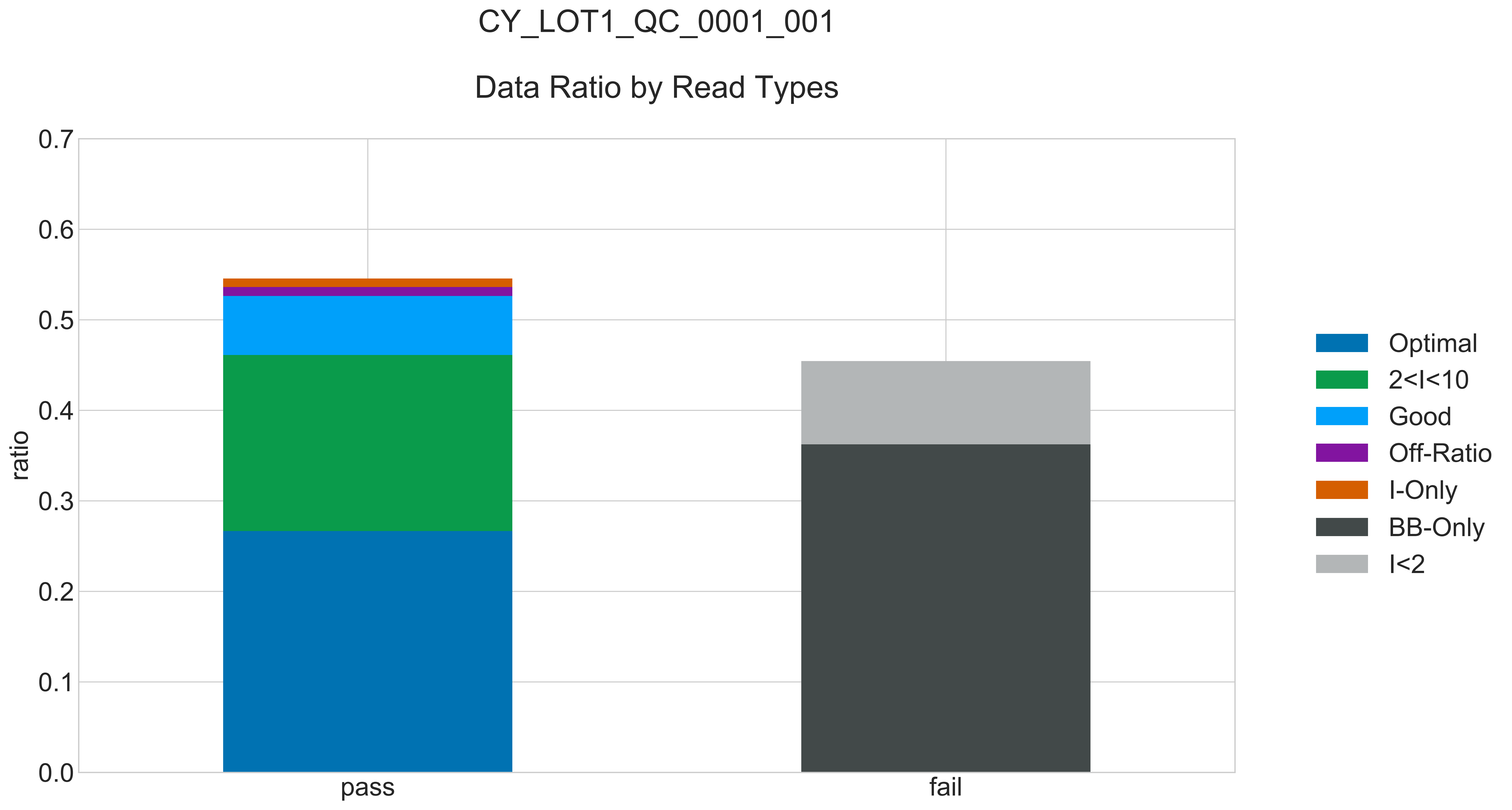

### CY_LOT1_QC_0001_001_READ_LEN_.pdf

#### CY_LOT1_QC_0001_001

##### Read Length Distribution

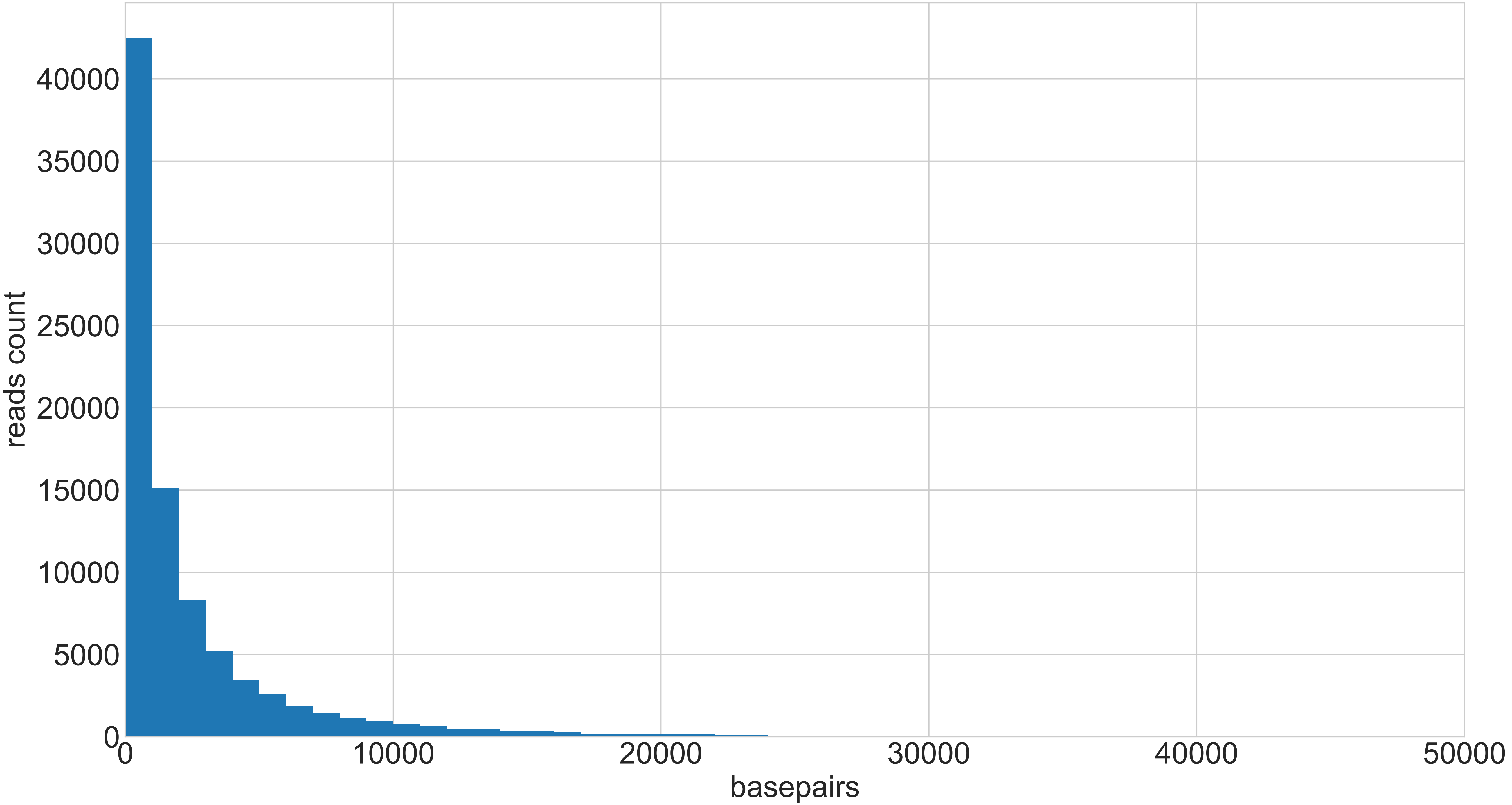

### CY_LOT1_QC_0001_001_REPEATS_COUNT_.pdf

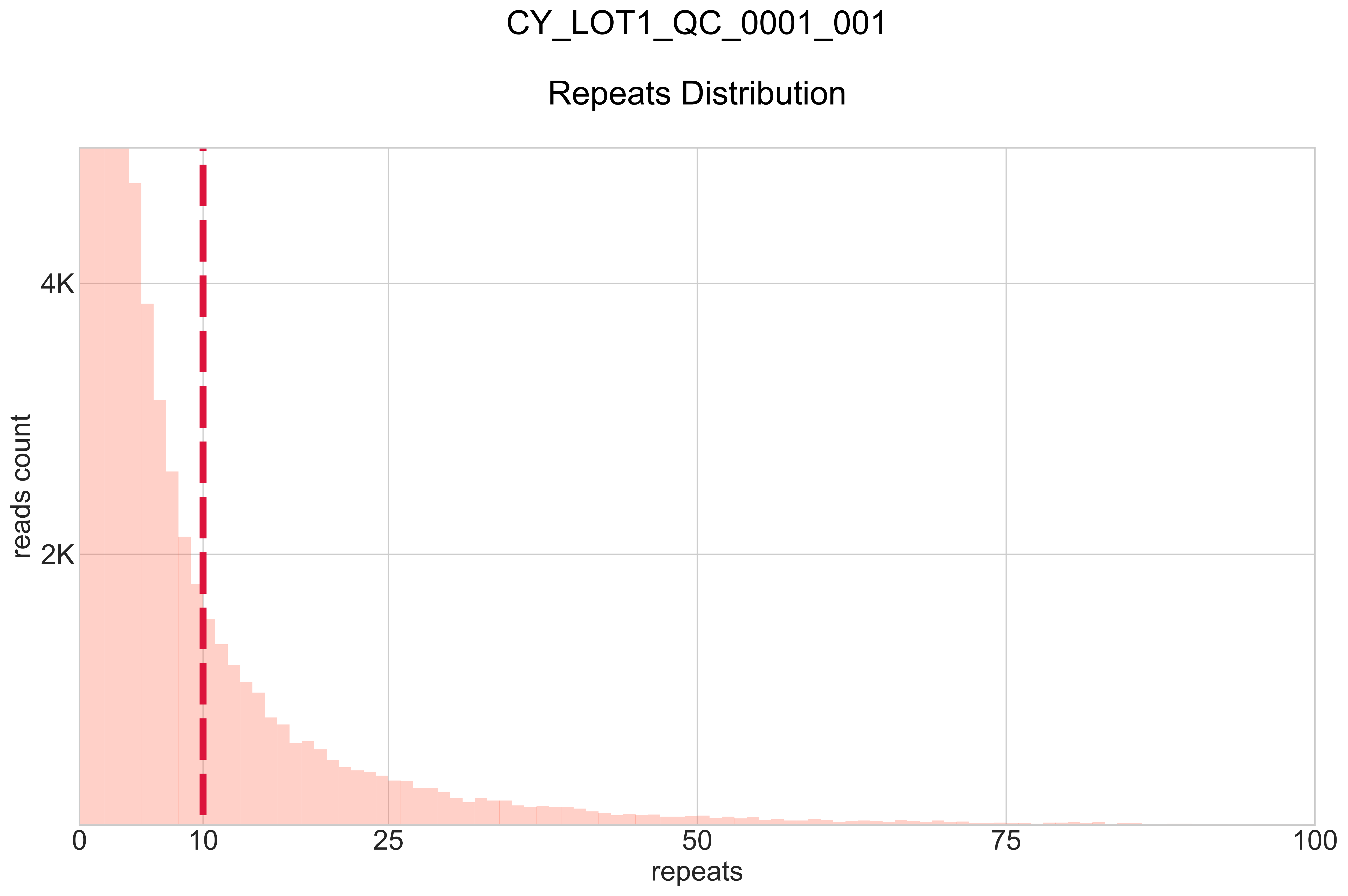

### CY_LOT1_QC_0002_000_BB-I_.pdf

#### CY_LOT1_QC_0002_000

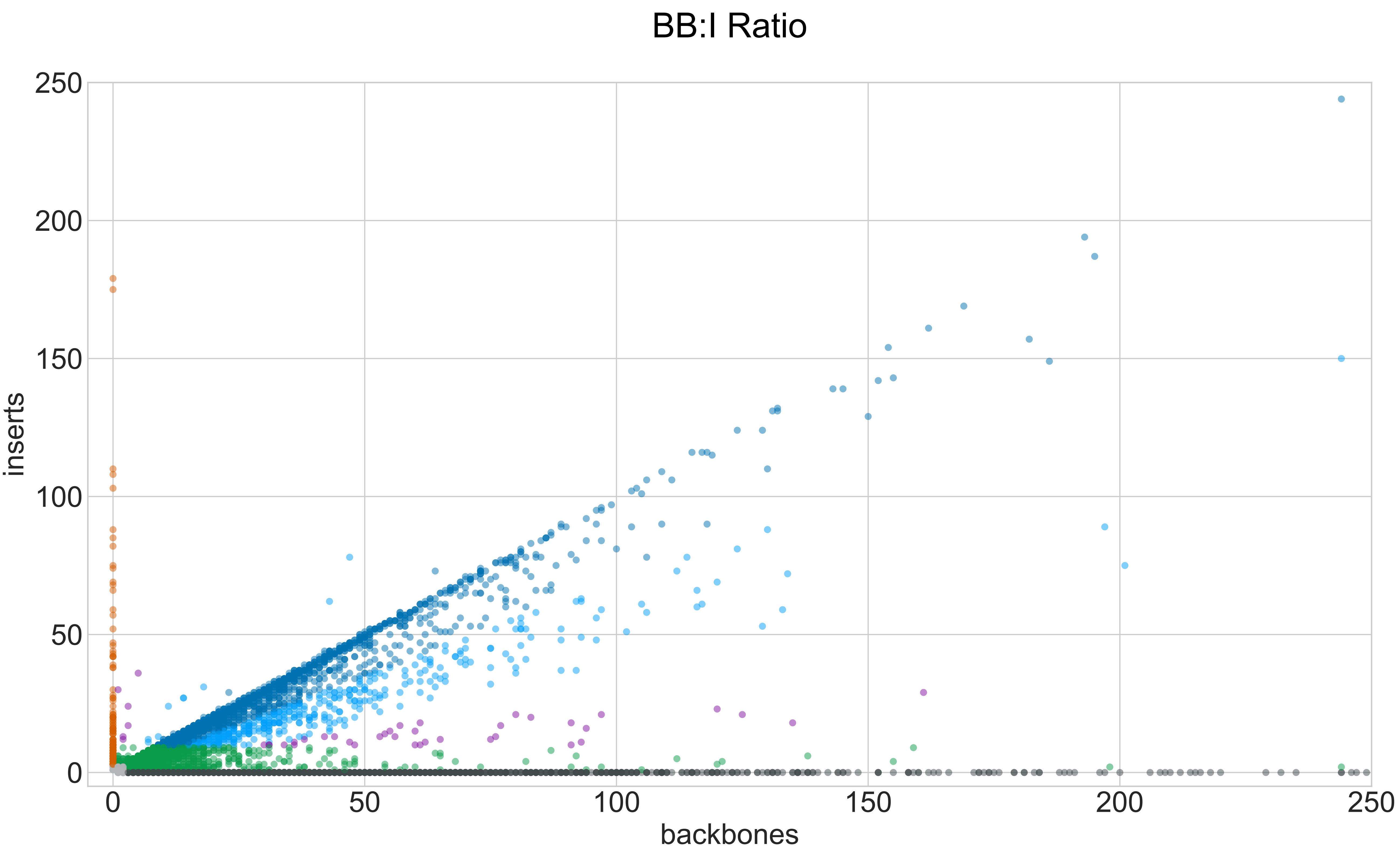

### CY_LOT1_QC_0002_000_GROUPS_STACKED_NORM-LEN_.pdf

#### CY_LOT1_QC_0002_000

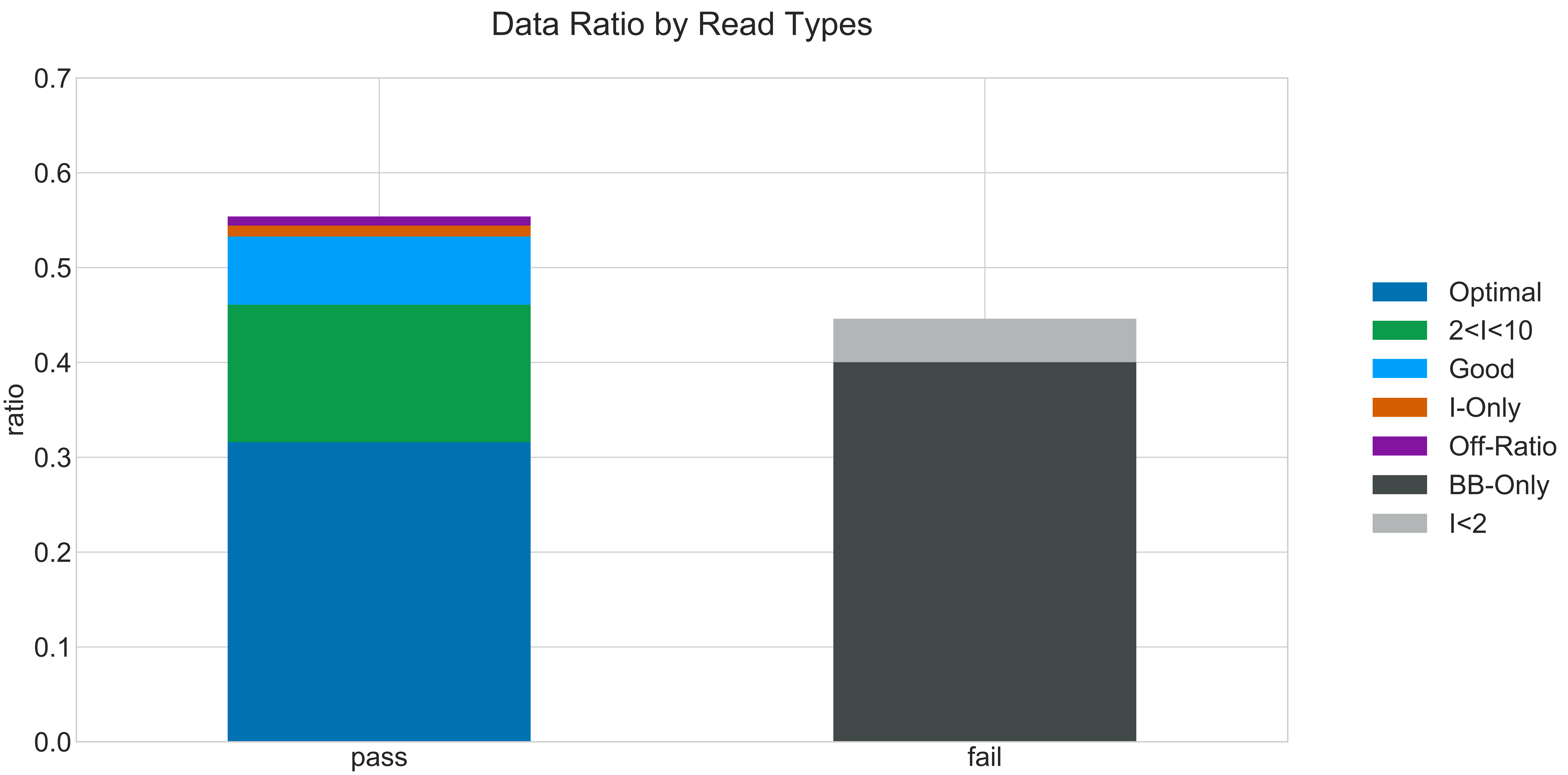

### CY_LOT1_QC_0002_000_READ_LEN_.pdf

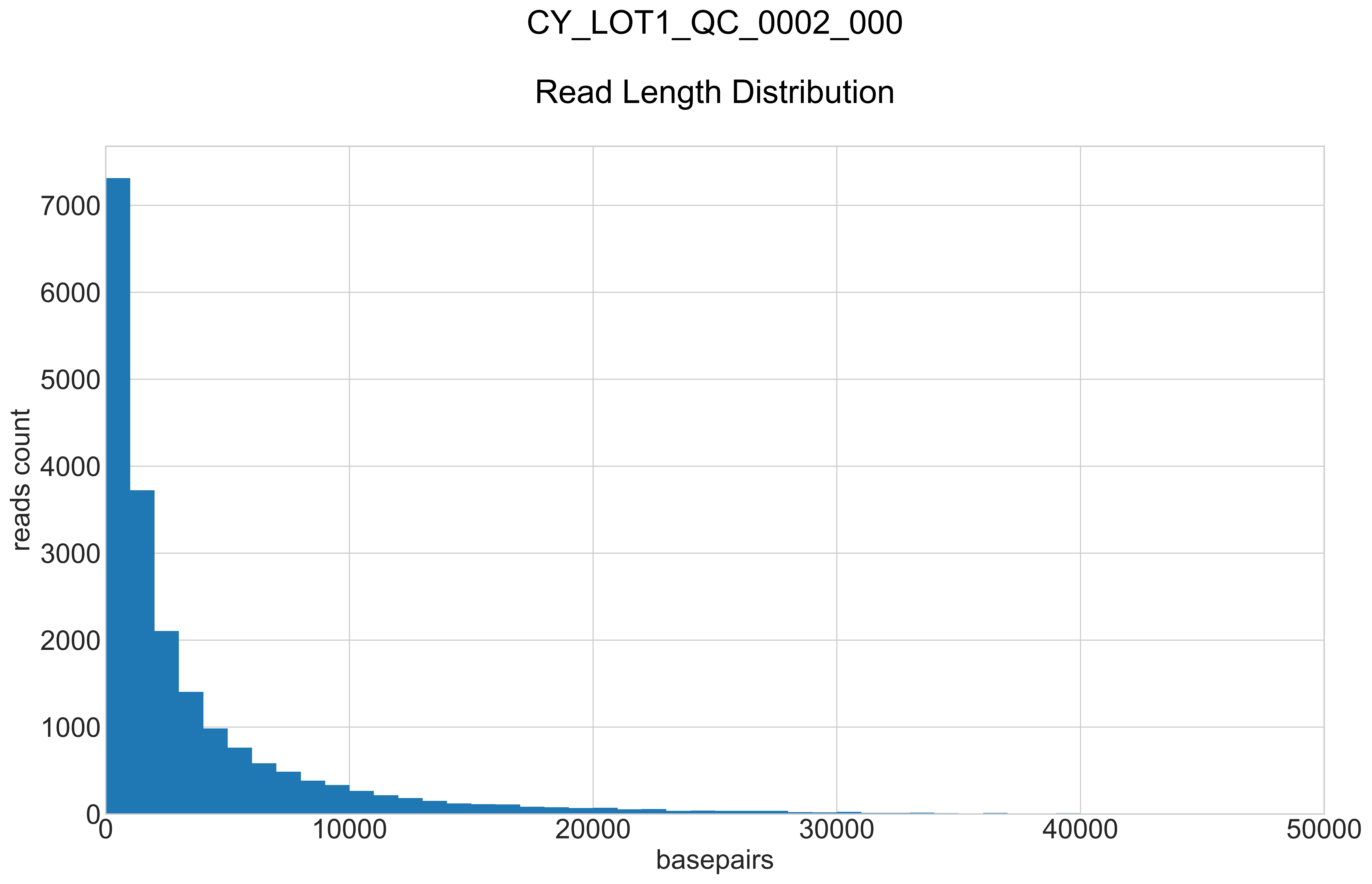

### CY_LOT1_QC_0002_000_REPEATS_COUNT_.pdf

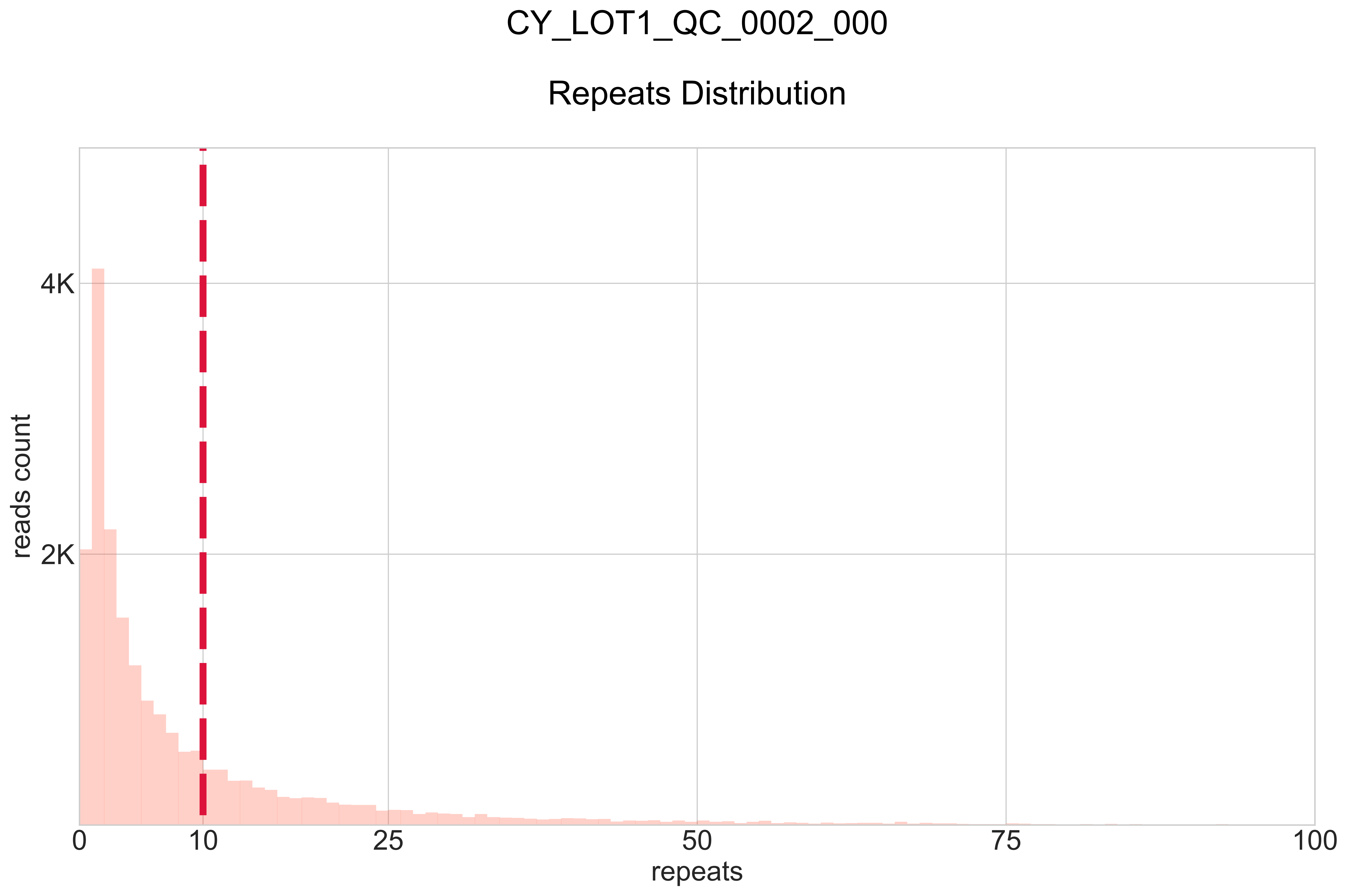

### CY_LOT1_QC_0002_001_BB-I_.pdf

#### CY_LOT1_QC_0002_001

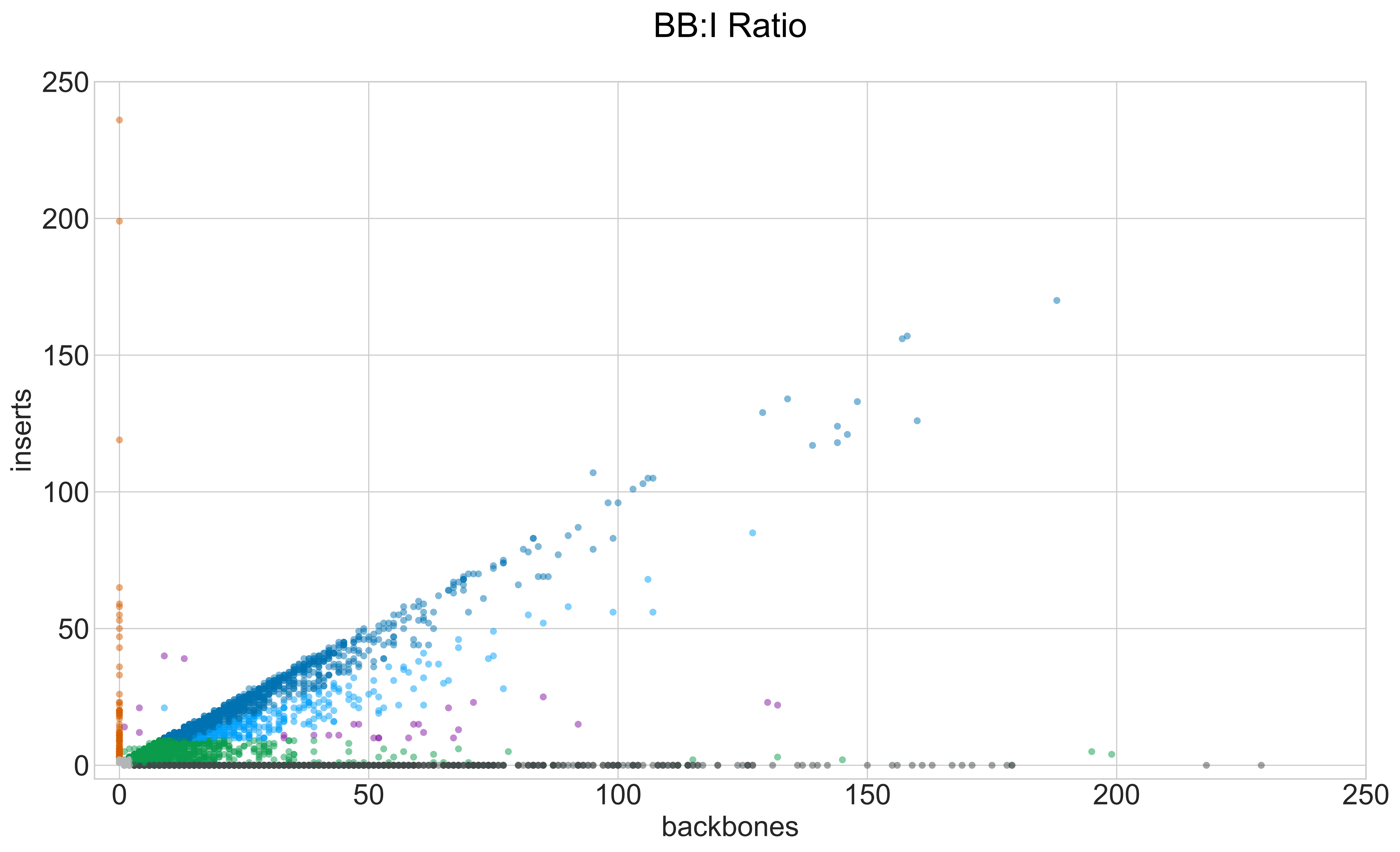

### CY_LOT1_QC_0002_001_GROUPS_STACKED_NORM-LEN_.pdf

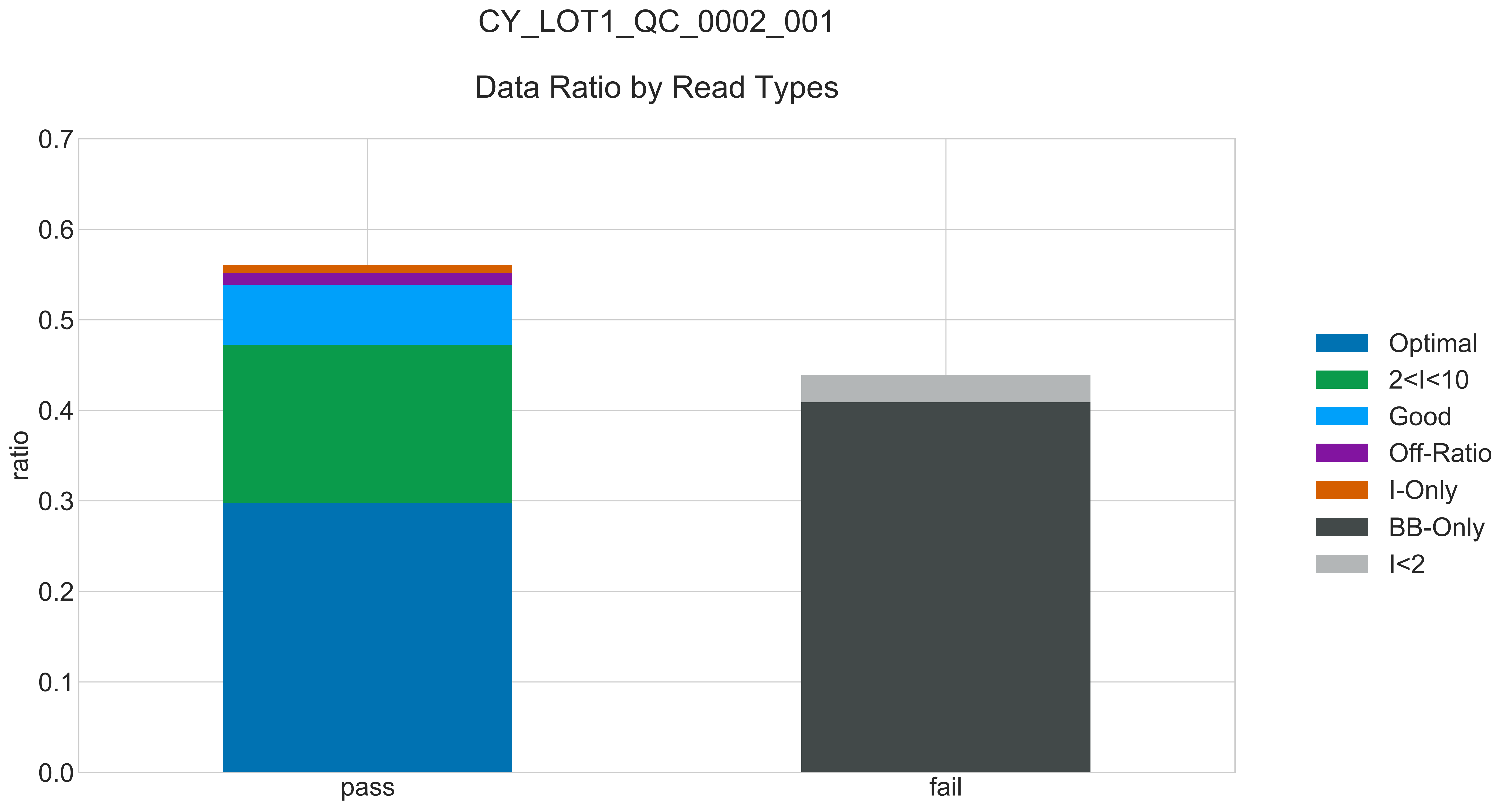

### CY_LOT1_QC_0002_001_READ_LEN_.pdf

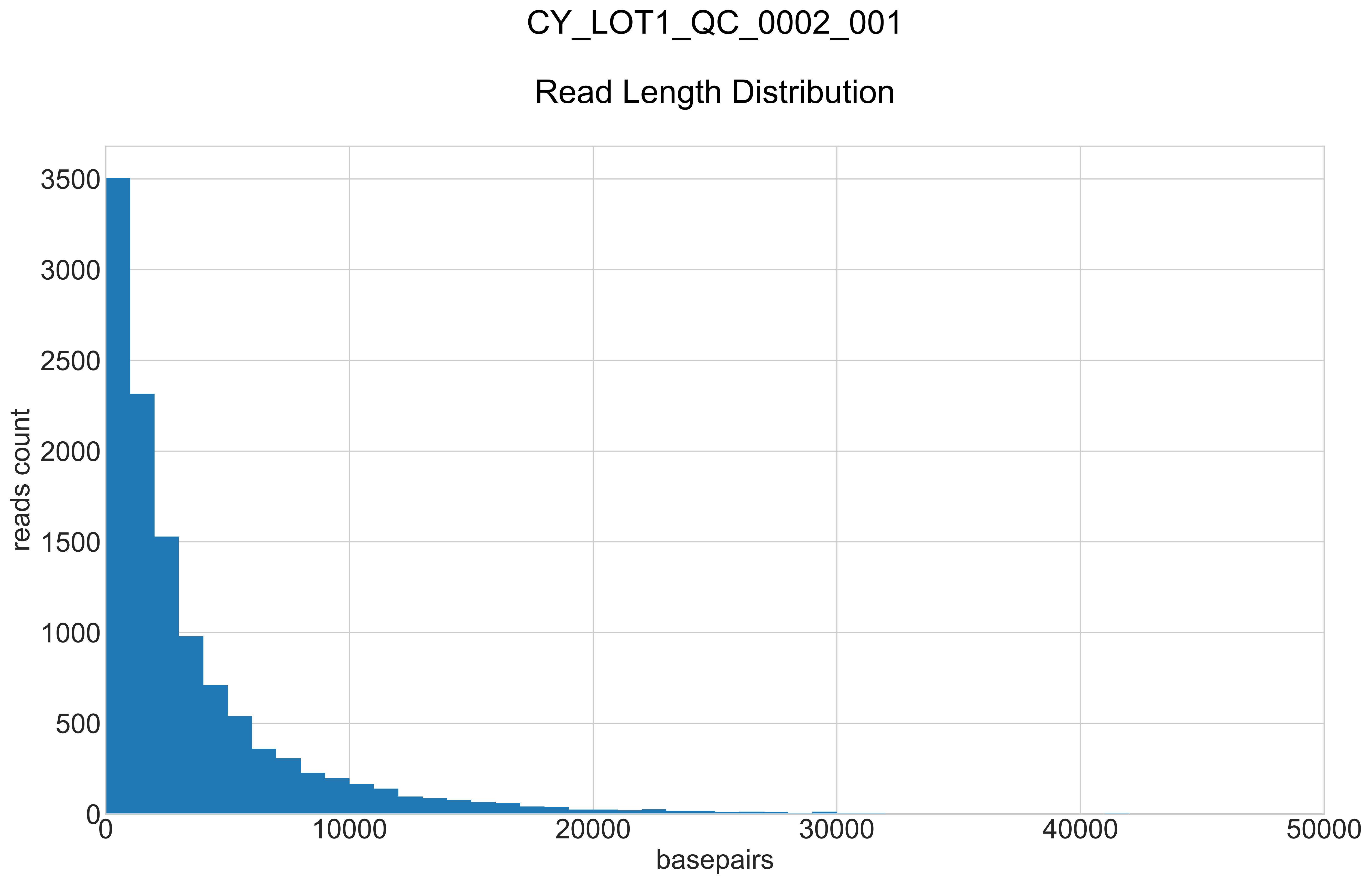

### CY_LOT1_QC_0002_001_REPEATS_COUNT_.pdf

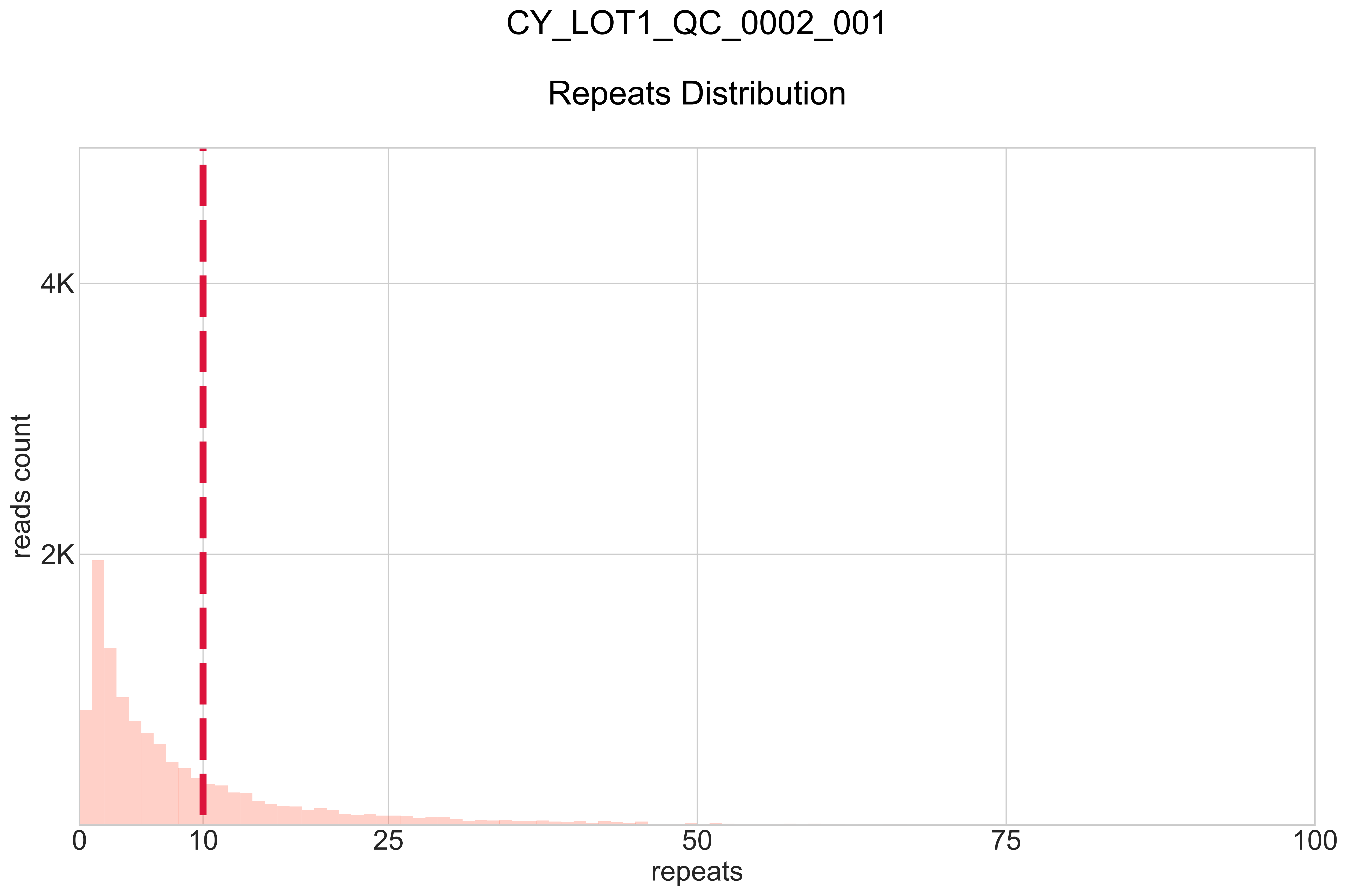

### CY_LOT1_QC_0003_000_BB-I_.pdf

#### CY_LOT1_QC_0003_000

##### BB:I Ratio

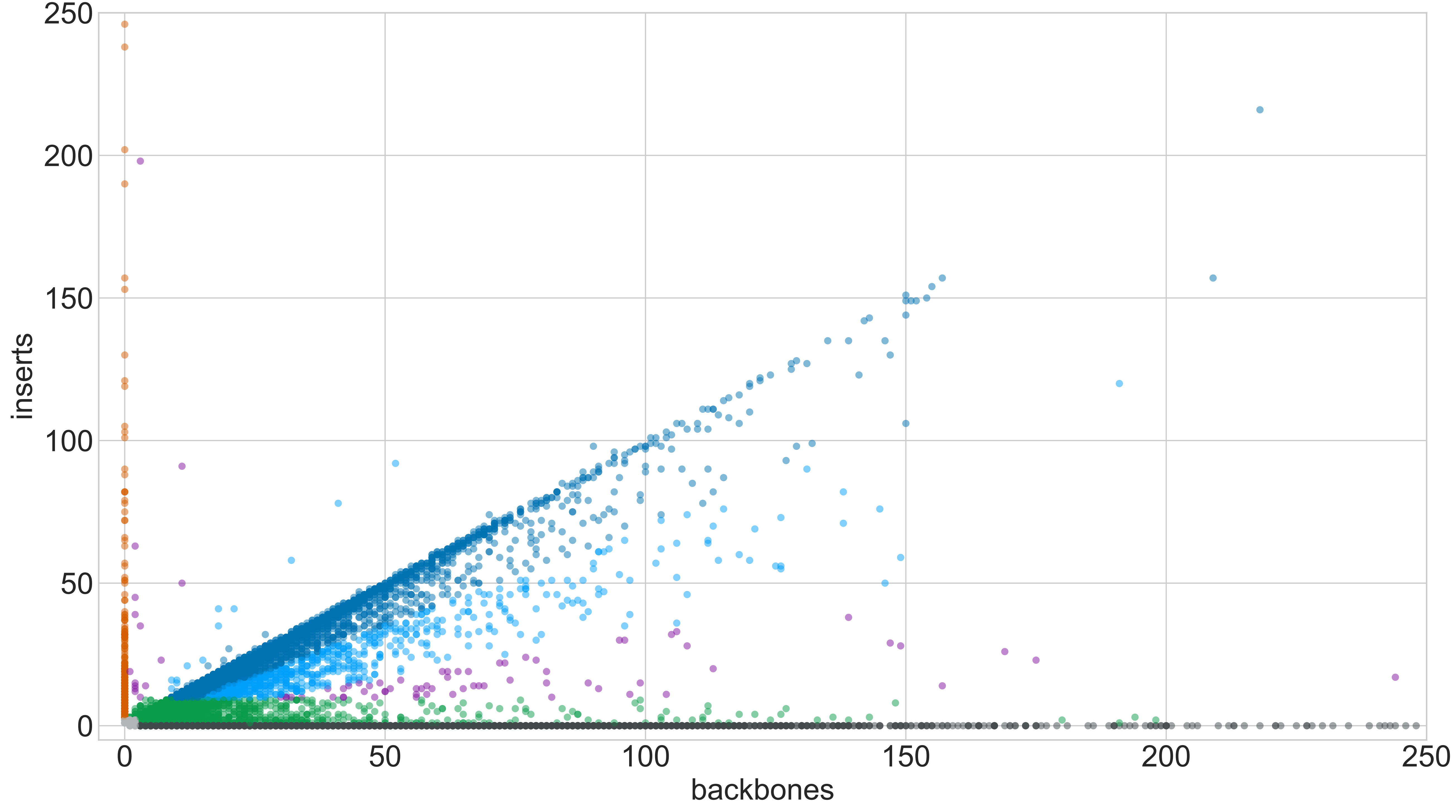

### CY_LOT1_QC_0003_000_GROUPS_STACKED_NORM-LEN_.pdf

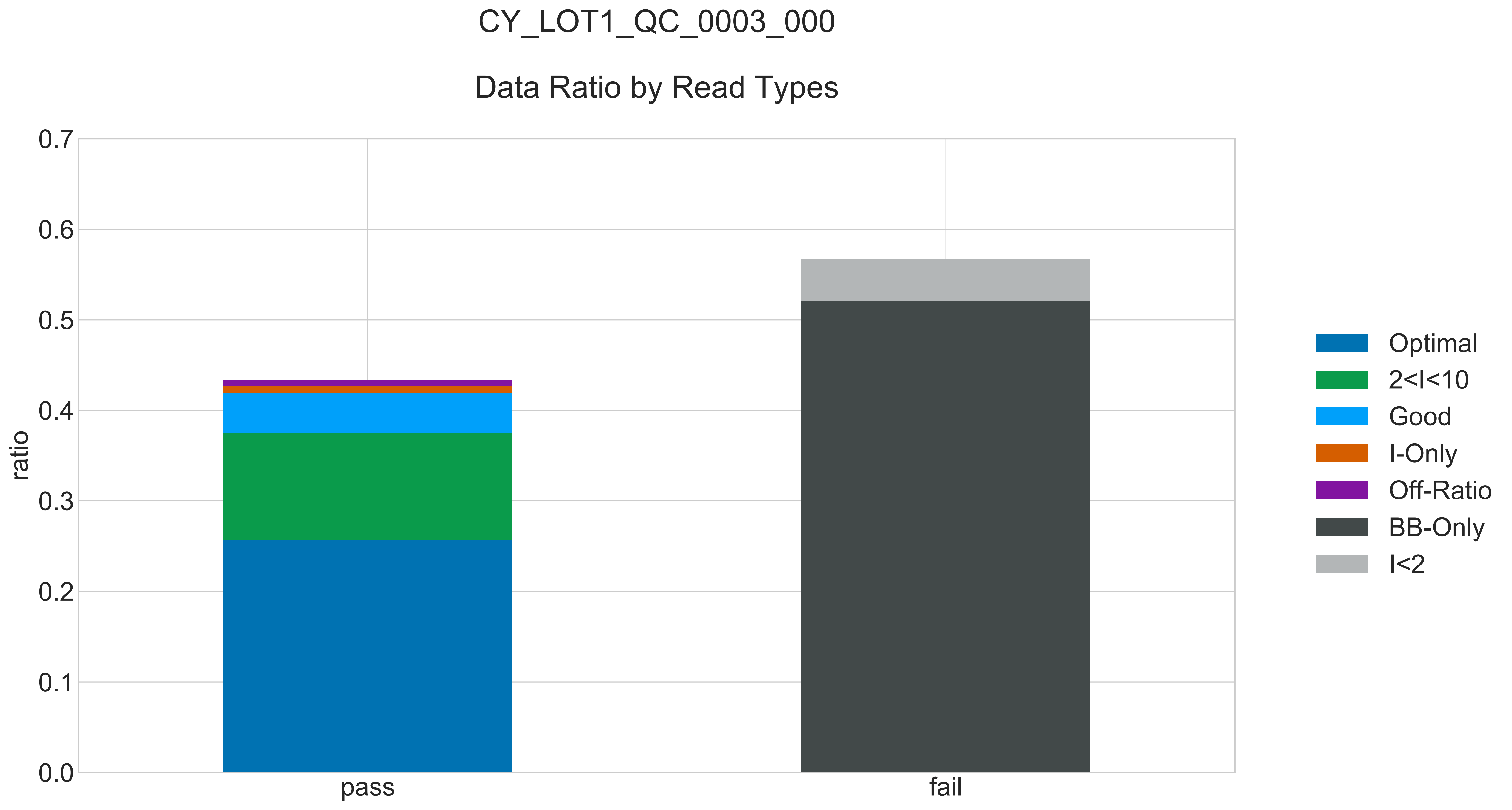

### CY_LOT1_QC_0003_000_READ_LEN_.pdf

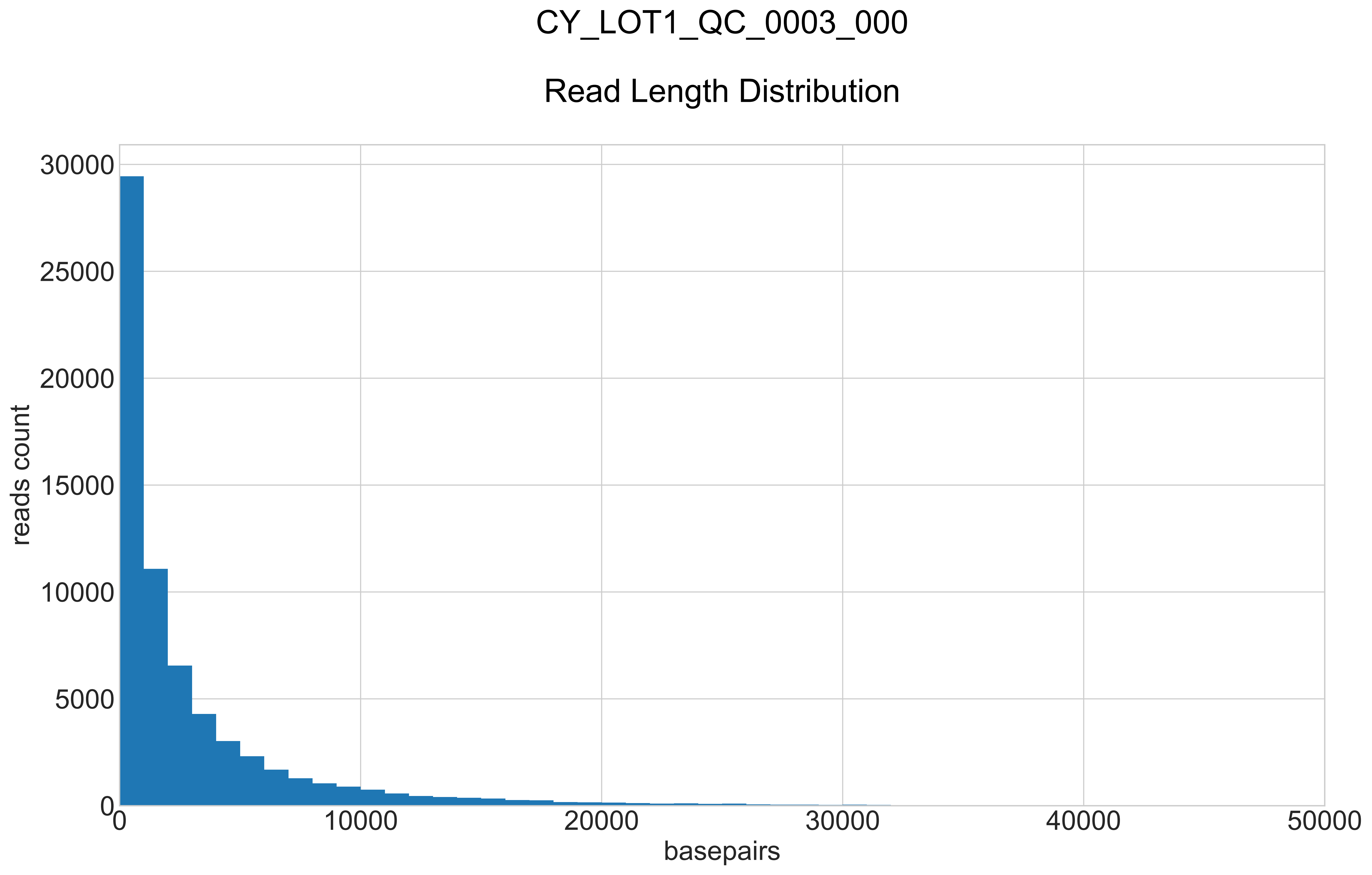

### CY_LOT1_QC_0003_000_REPEATS_COUNT_.pdf

#### CY_LOT1_QC_0003_000

##### Repeats Distribution

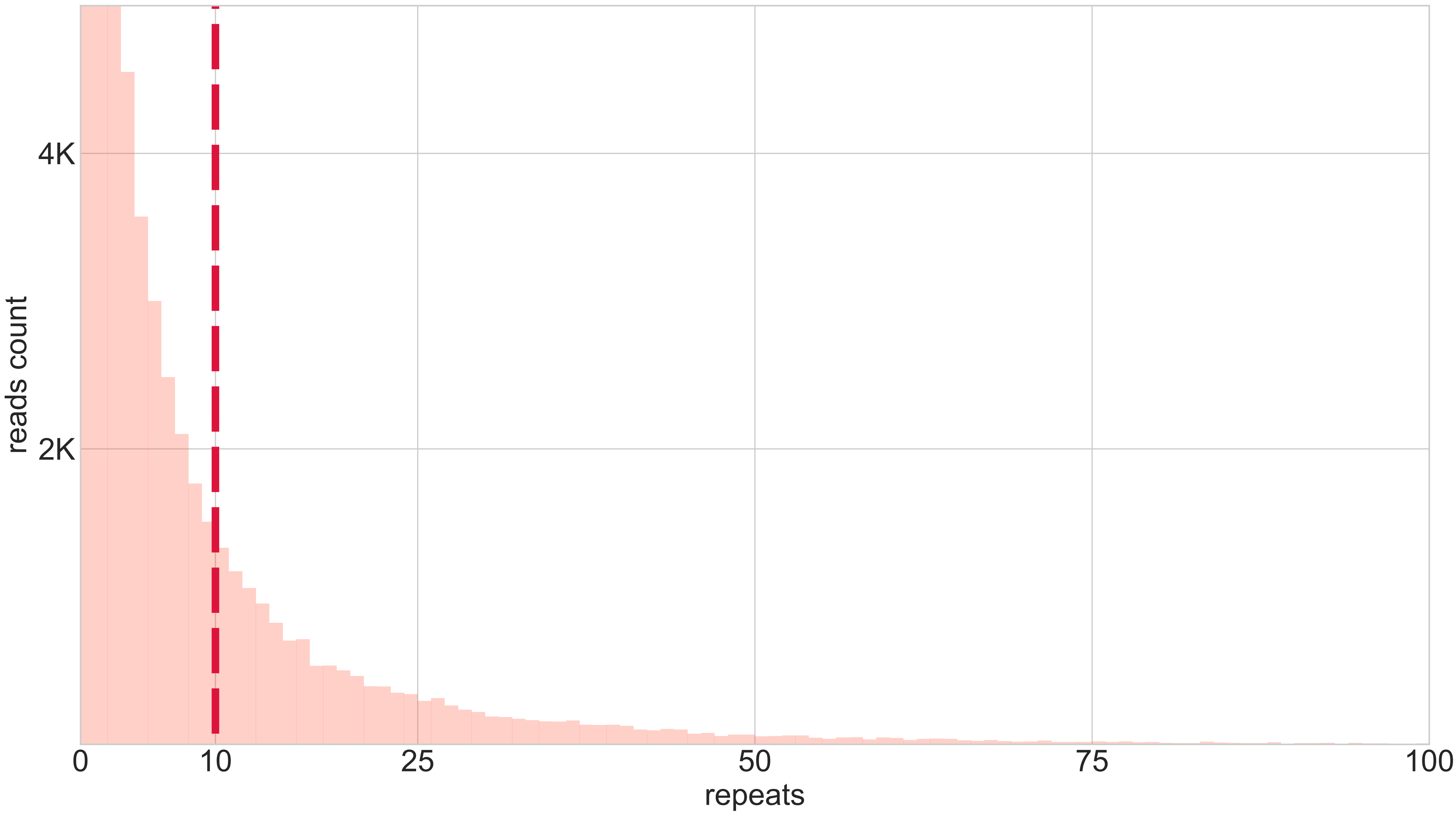

### CY_LOT1_QC_0003_001_BB-I_.pdf

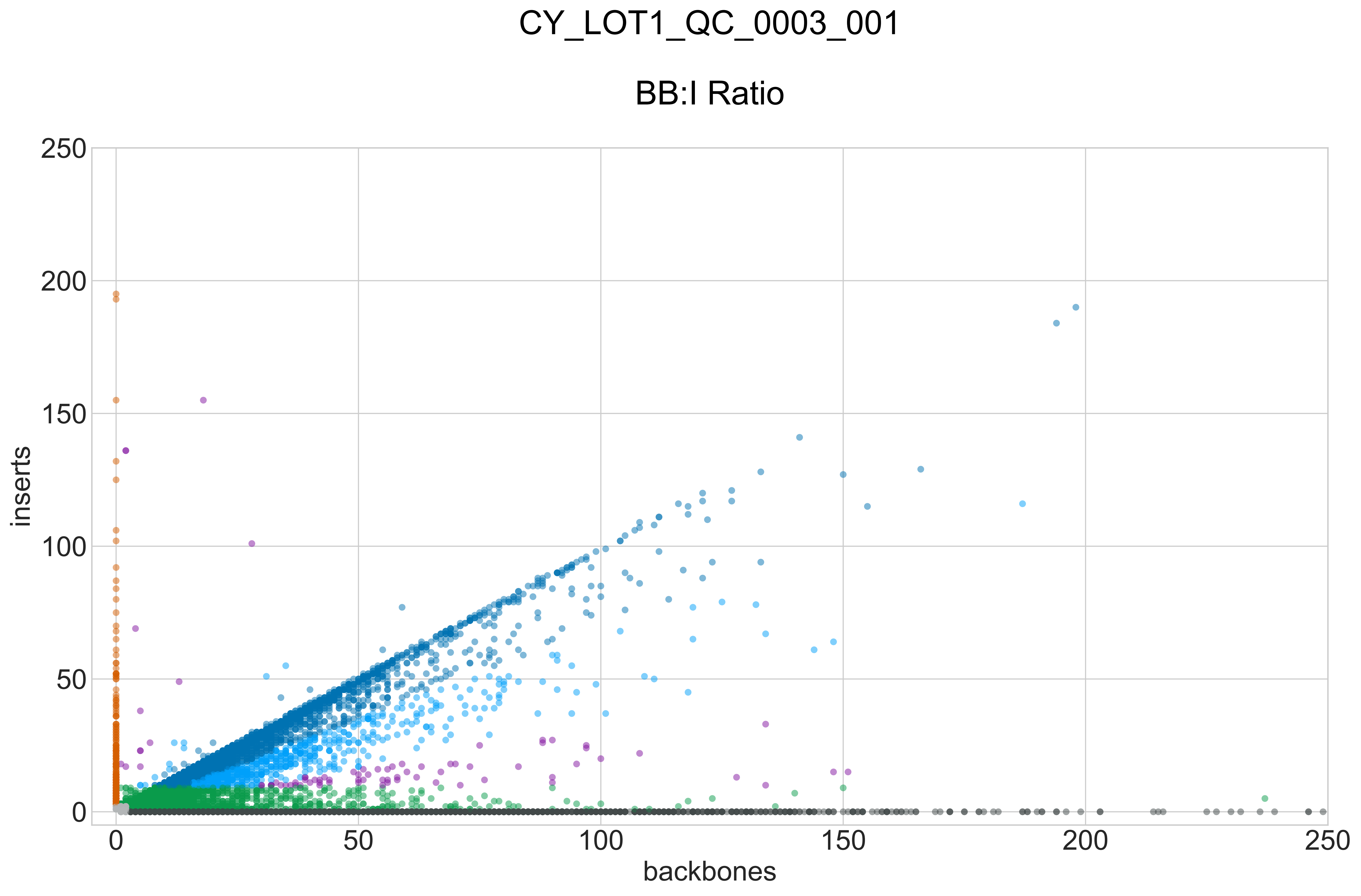

### CY_LOT1_QC_0003_001_GROUPS_STACKED_NORM-LEN_.pdf

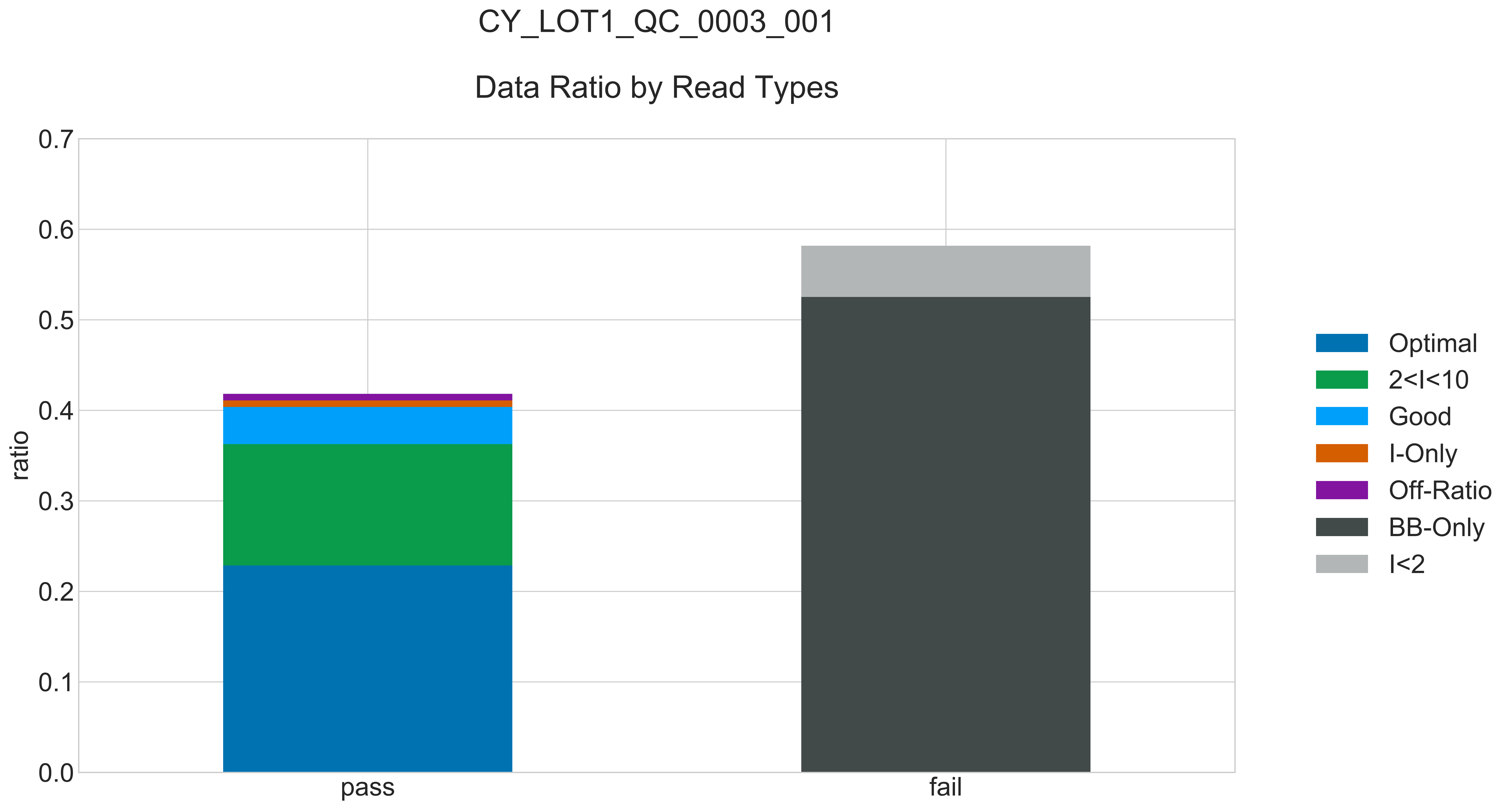

### CY_LOT1_QC_0003_001_READ_LEN_.pdf

#### CY_LOT1_QC_0003_001

##### Read Length Distribution

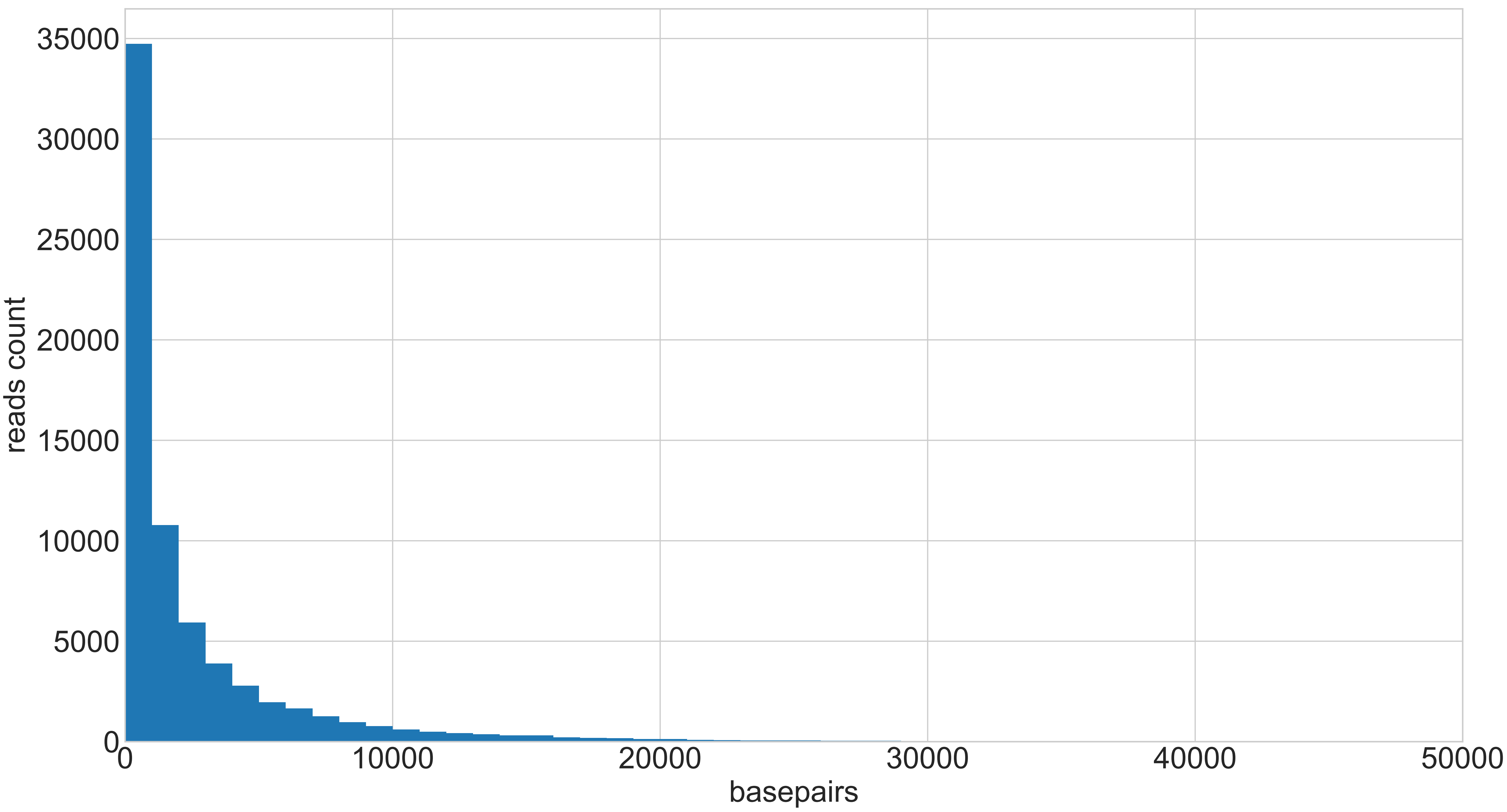

### CY_LOT1_QC_0003_001_REPEATS_COUNT_.pdf

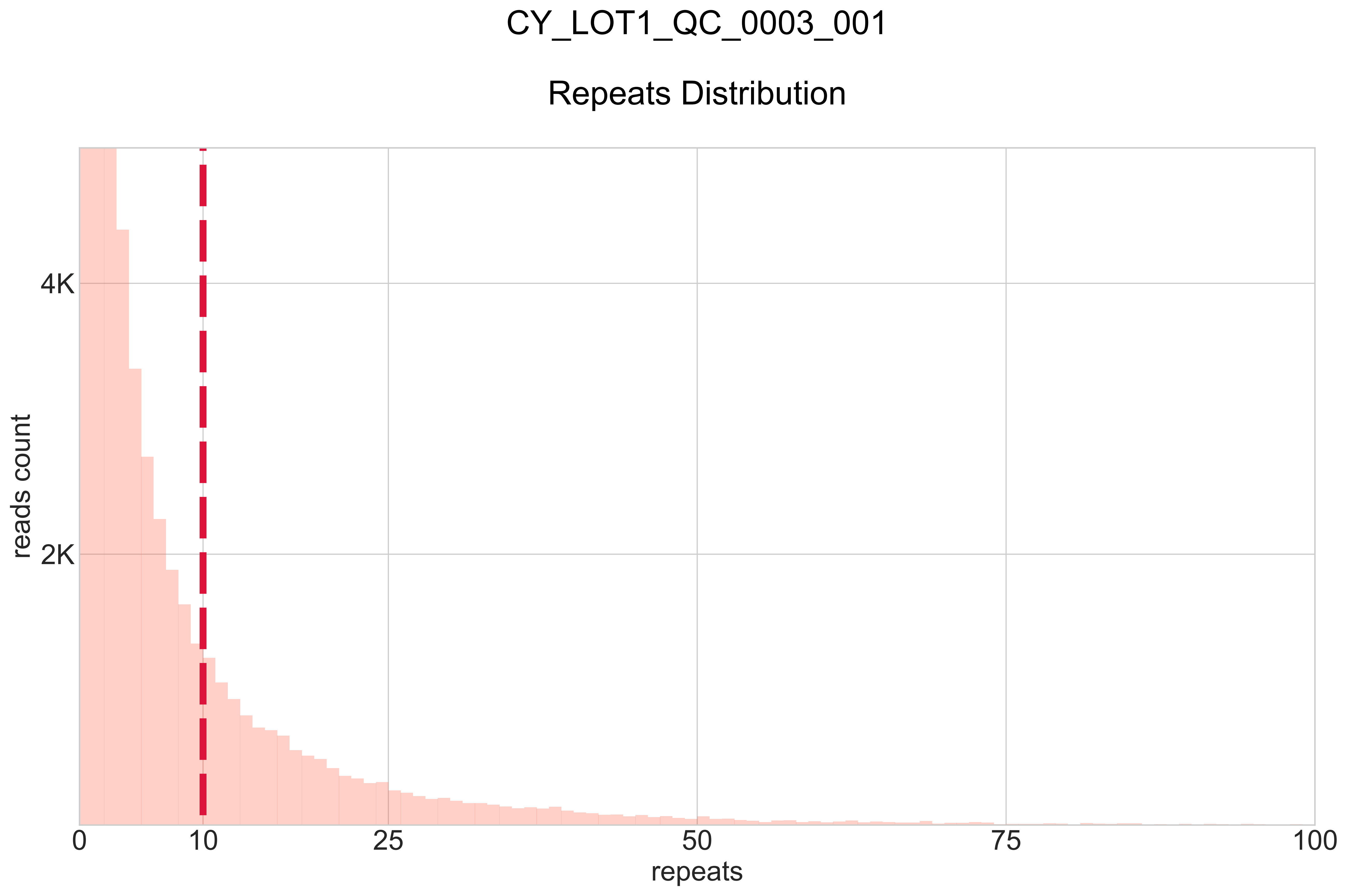

### CY_PJET_12MU_0001_000_BB-I_.pdf

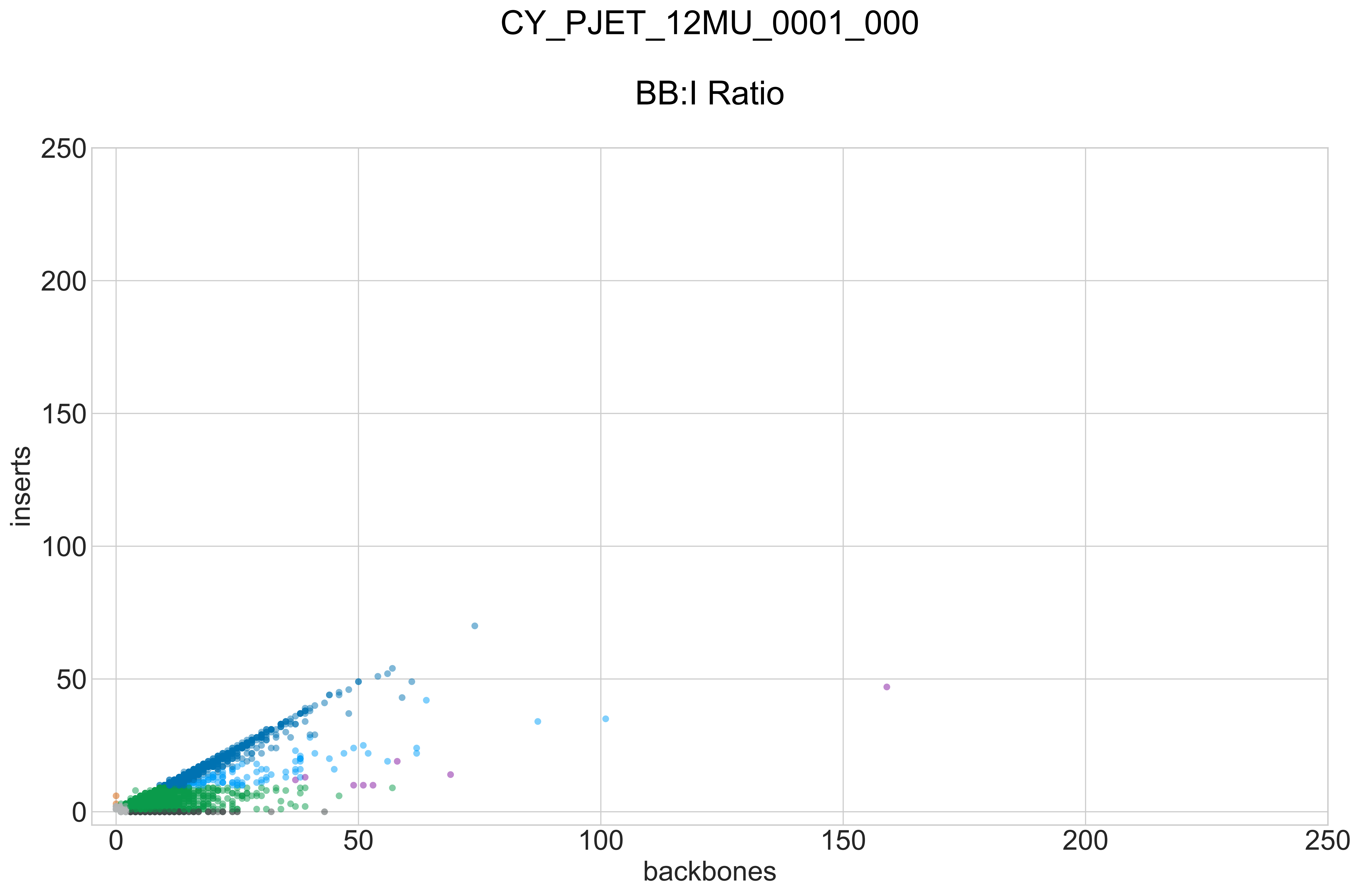

### CY_PJET_12MU_0001_000_GROUPS_STACKED_NORM-LEN_.pdf

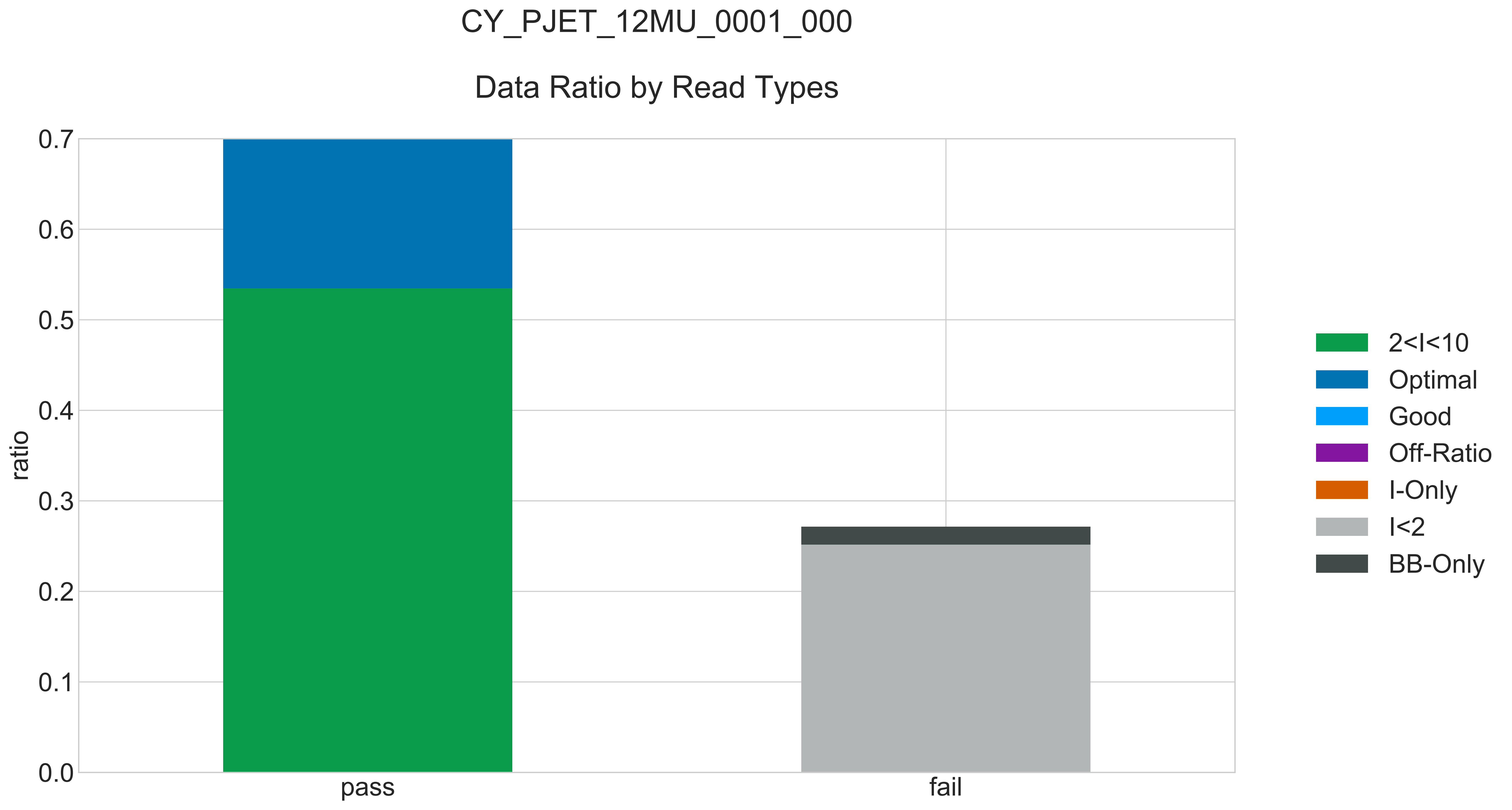

### CY_PJET_12MU_0001_000_READ_LEN_.pdf

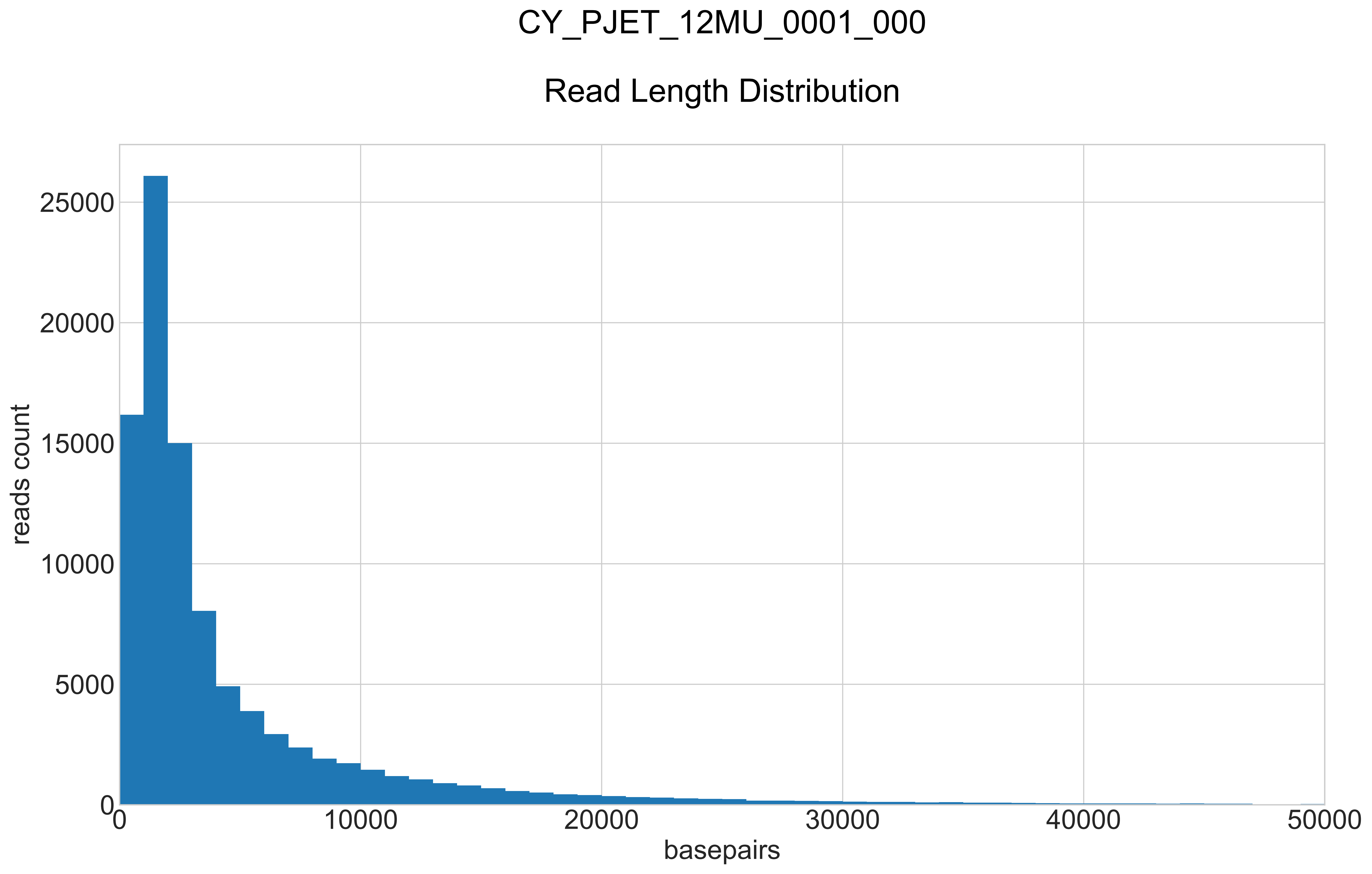

### CY_PJET_12MU_0001_000_REPEATS_COUNT_.pdf

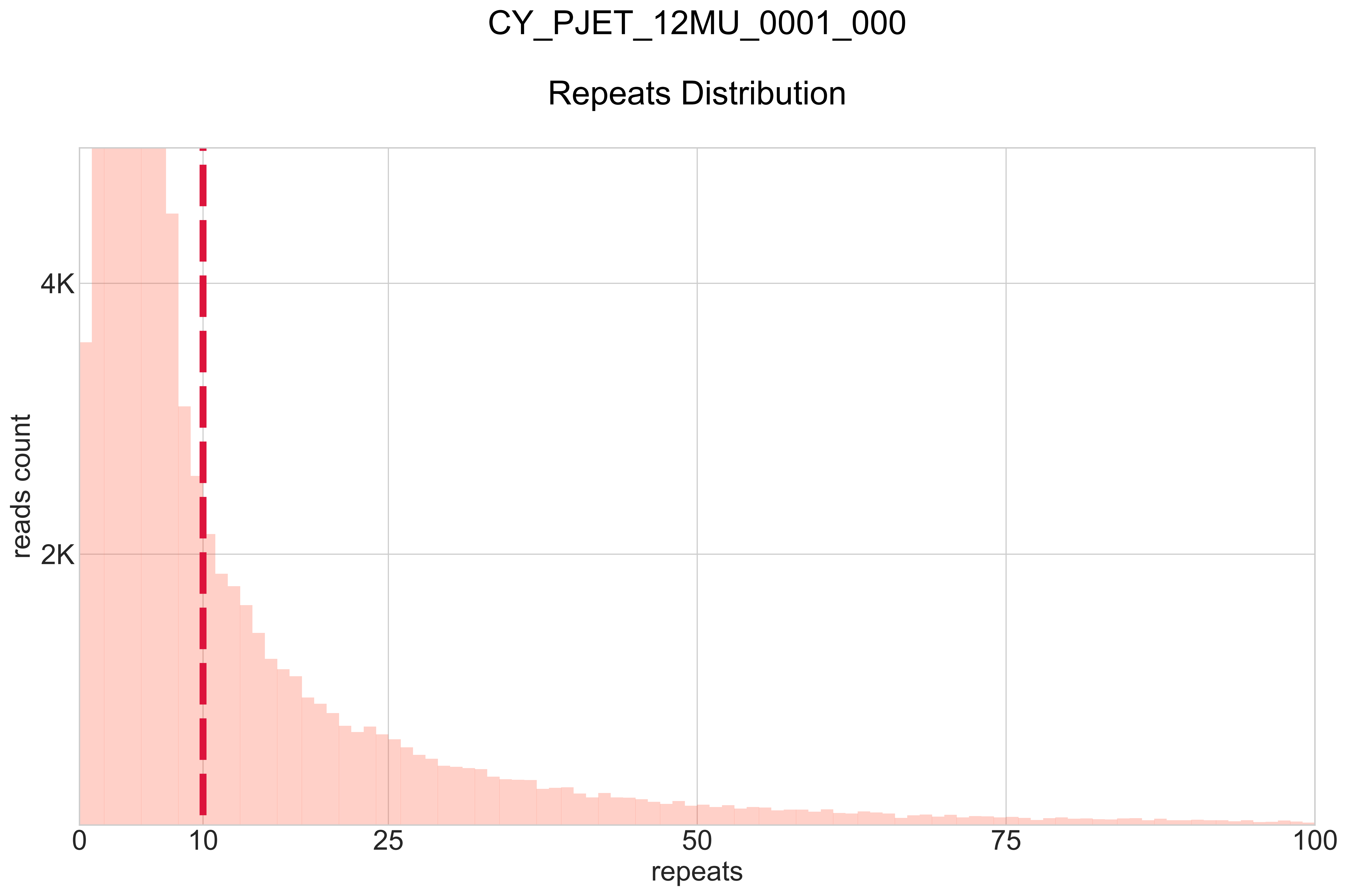

### CY_PJET_12WT_0001_000_BB-I_.pdf

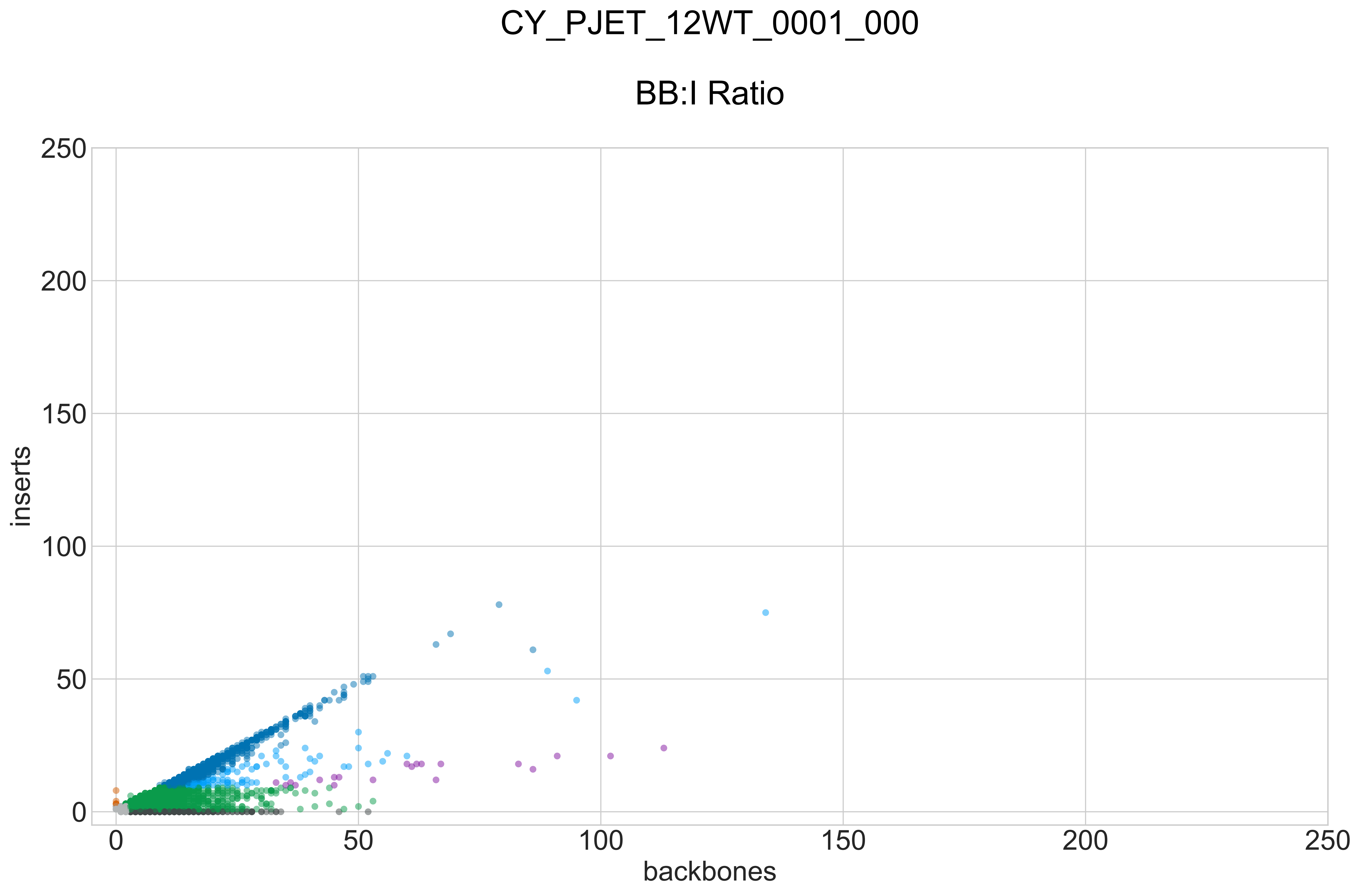

### CY_PJET_12WT_0001_000_GROUPS_STACKED_NORM-LEN_.pdf

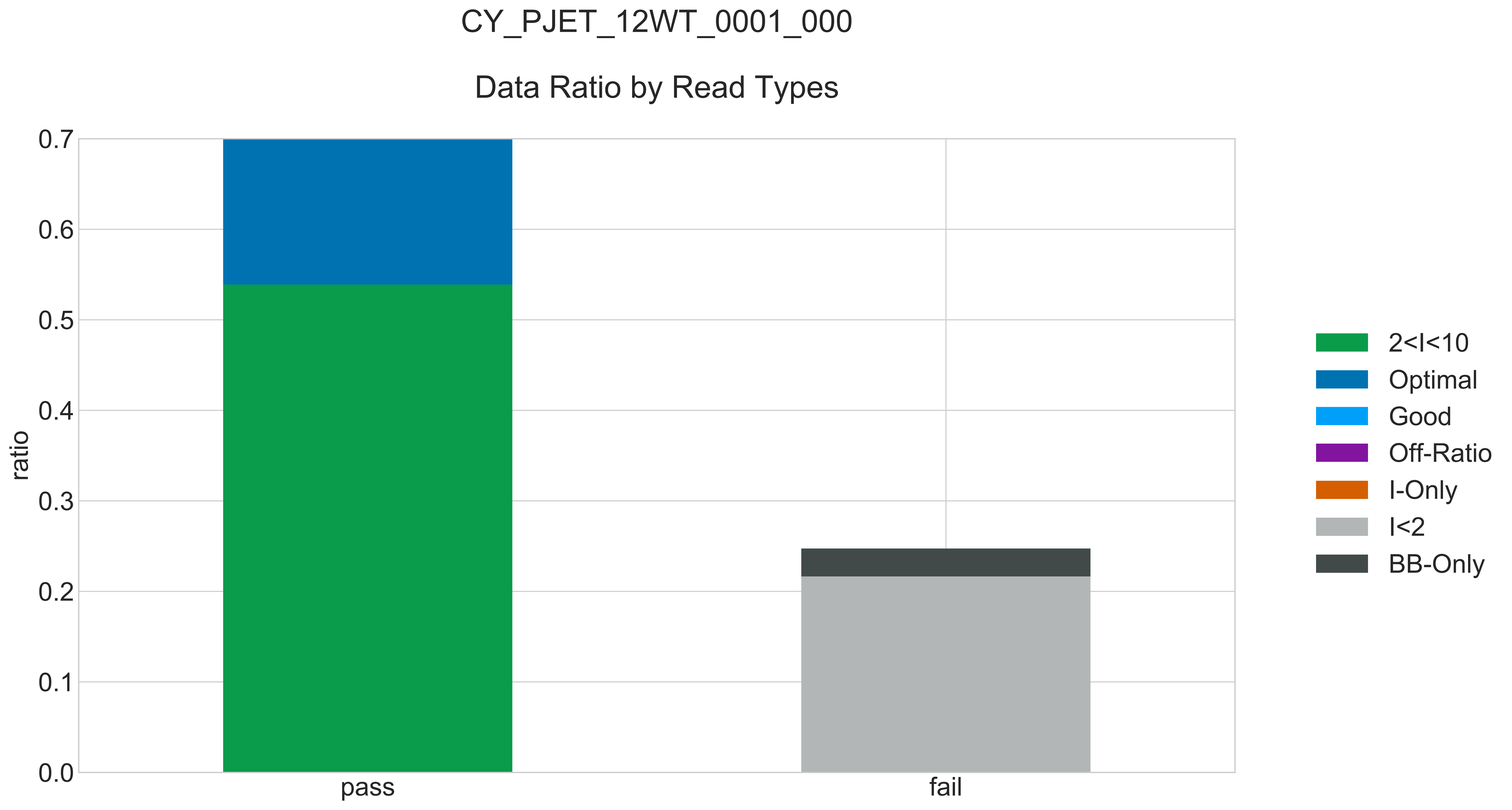

### CY_PJET_12WT_0001_000_READ_LEN_.pdf

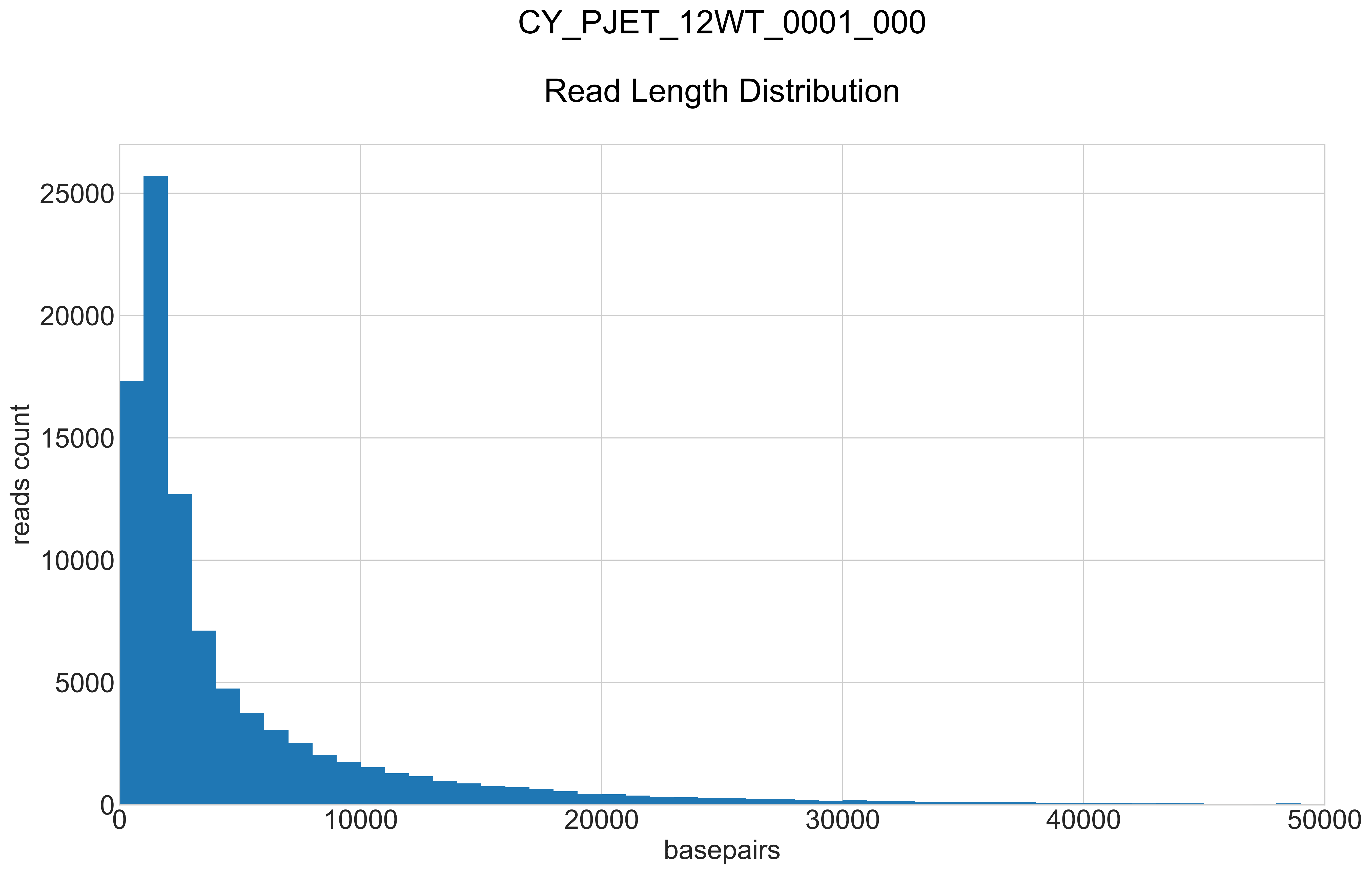

### CY_PJET_12WT_0001_000_REPEATS_COUNT_.pdf

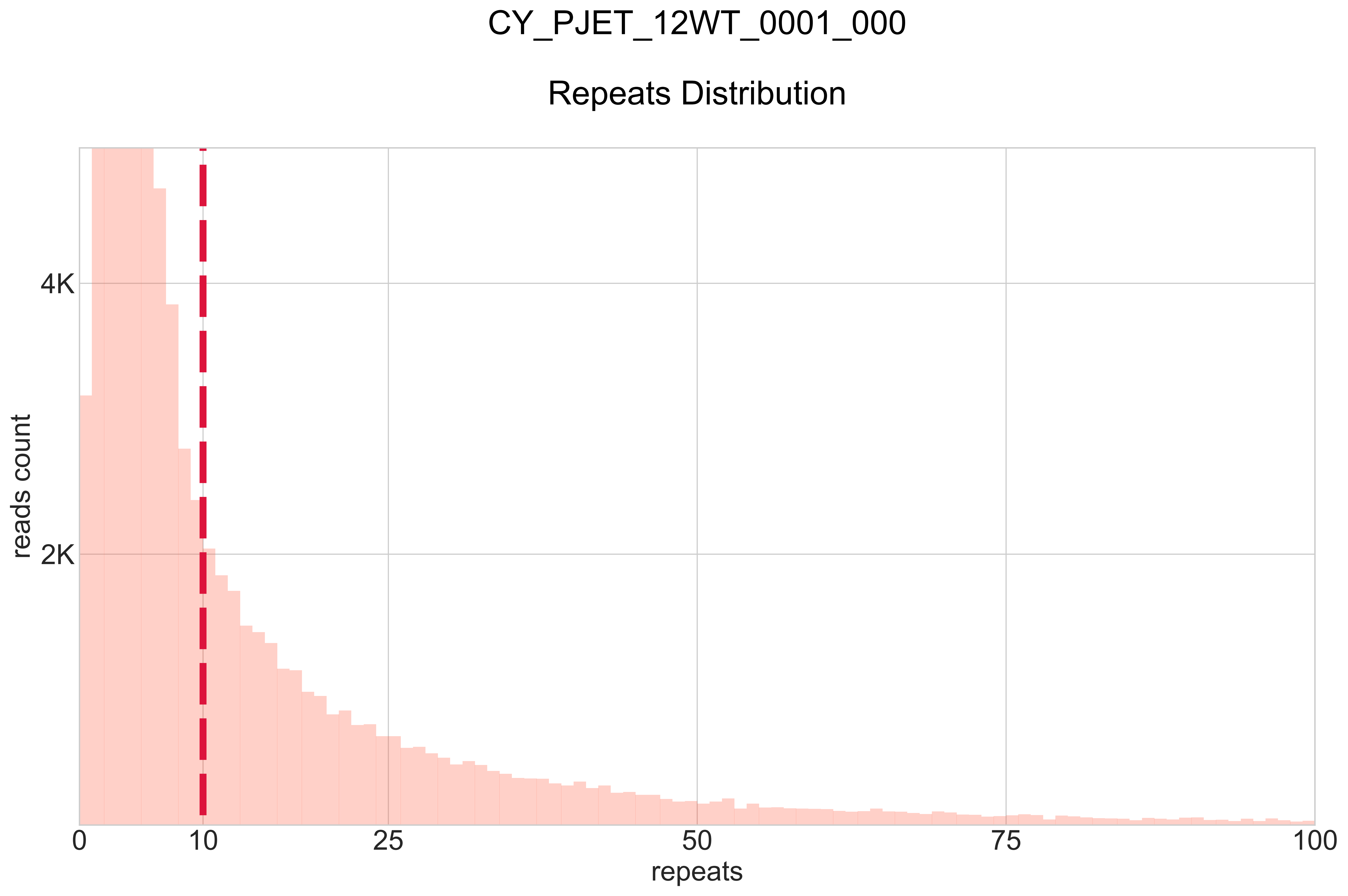

### CY_PJET_RATI_0001_000_BB-I_.pdf

#### CY_PJET_RATI_0001_000

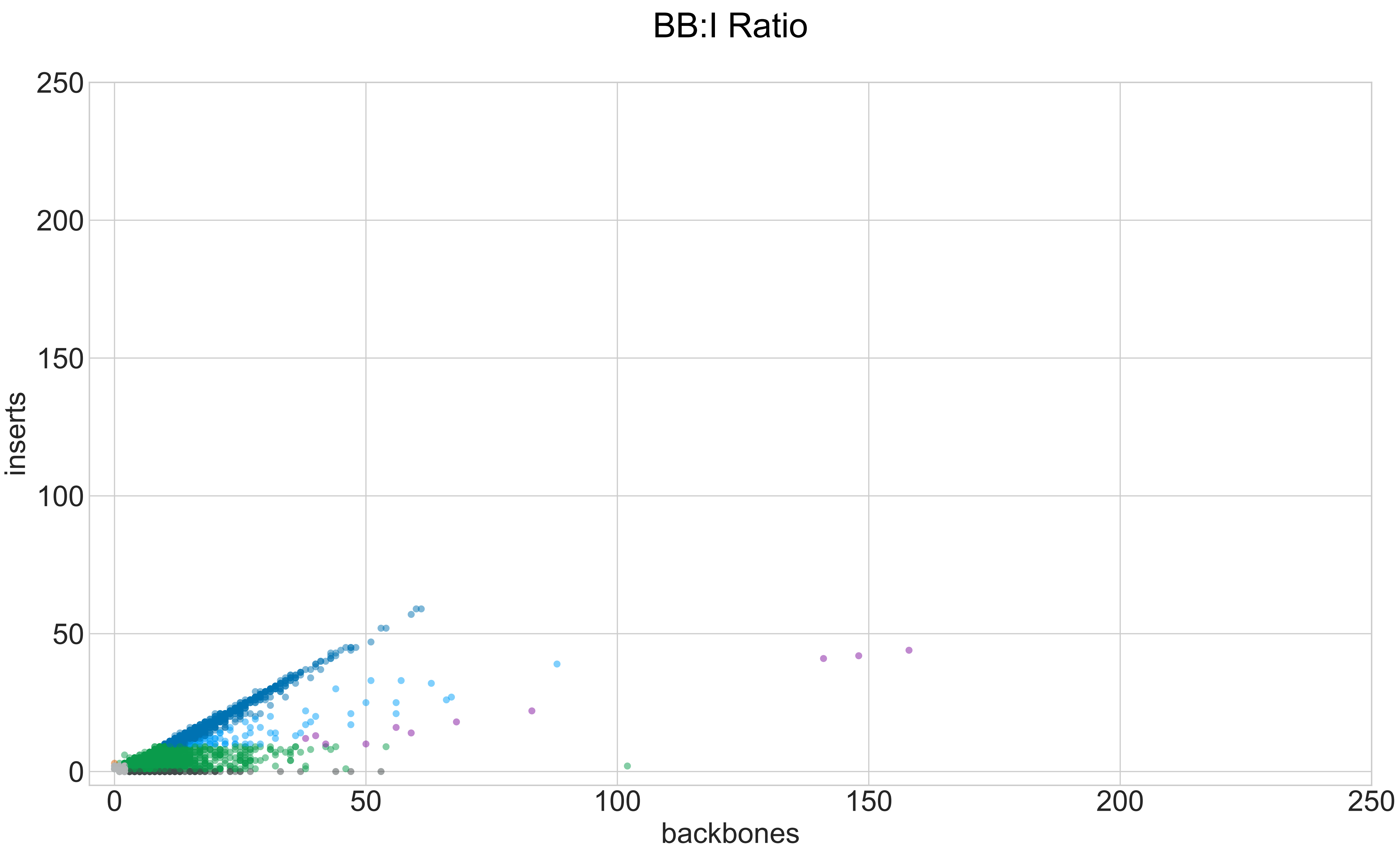

### CY_PJET_RATI_0001_000_GROUPS_STACKED_NORM-LEN_.pdf

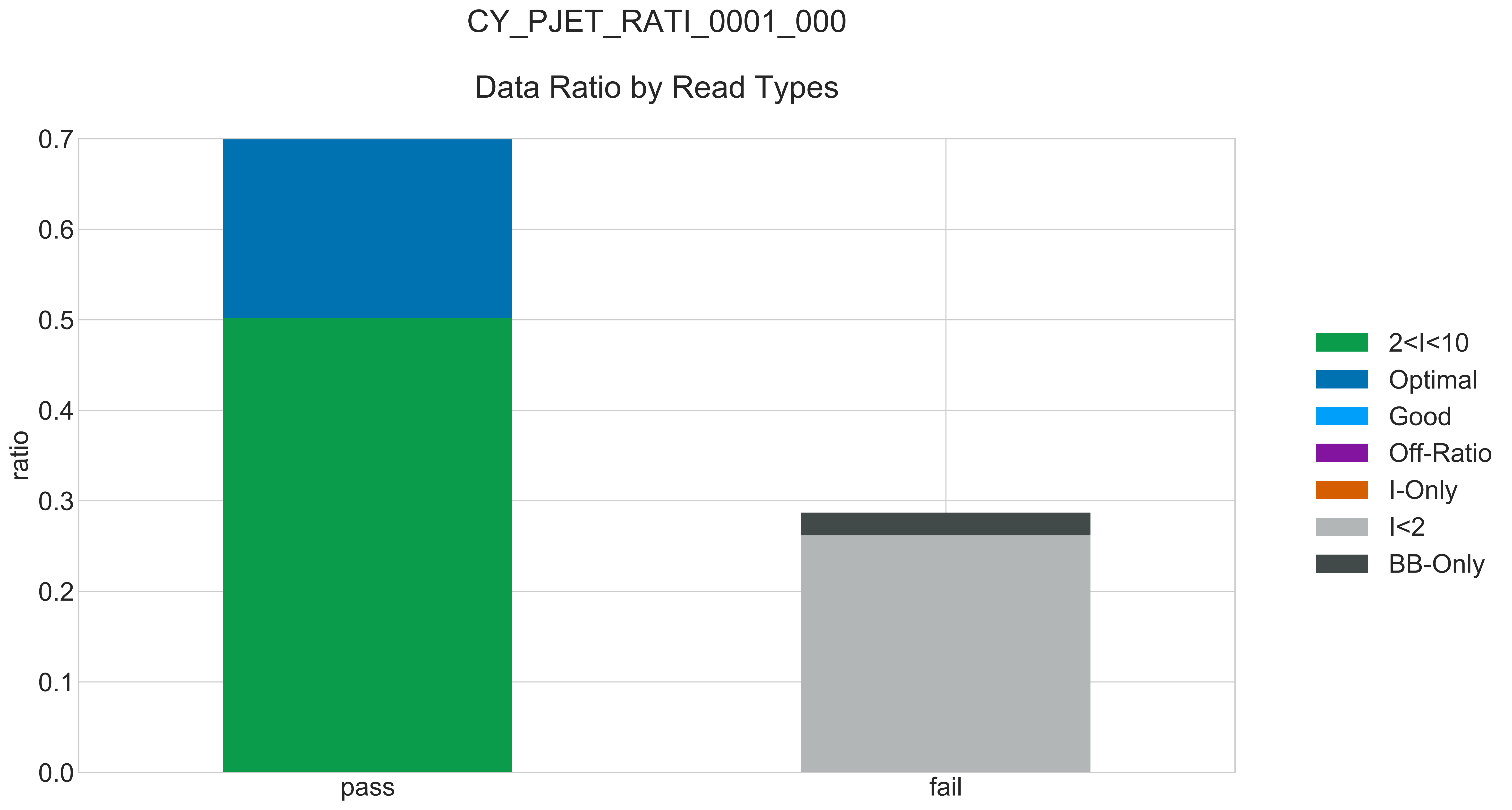

### CY_PJET_RATI_0001_000_READ_LEN_.pdf

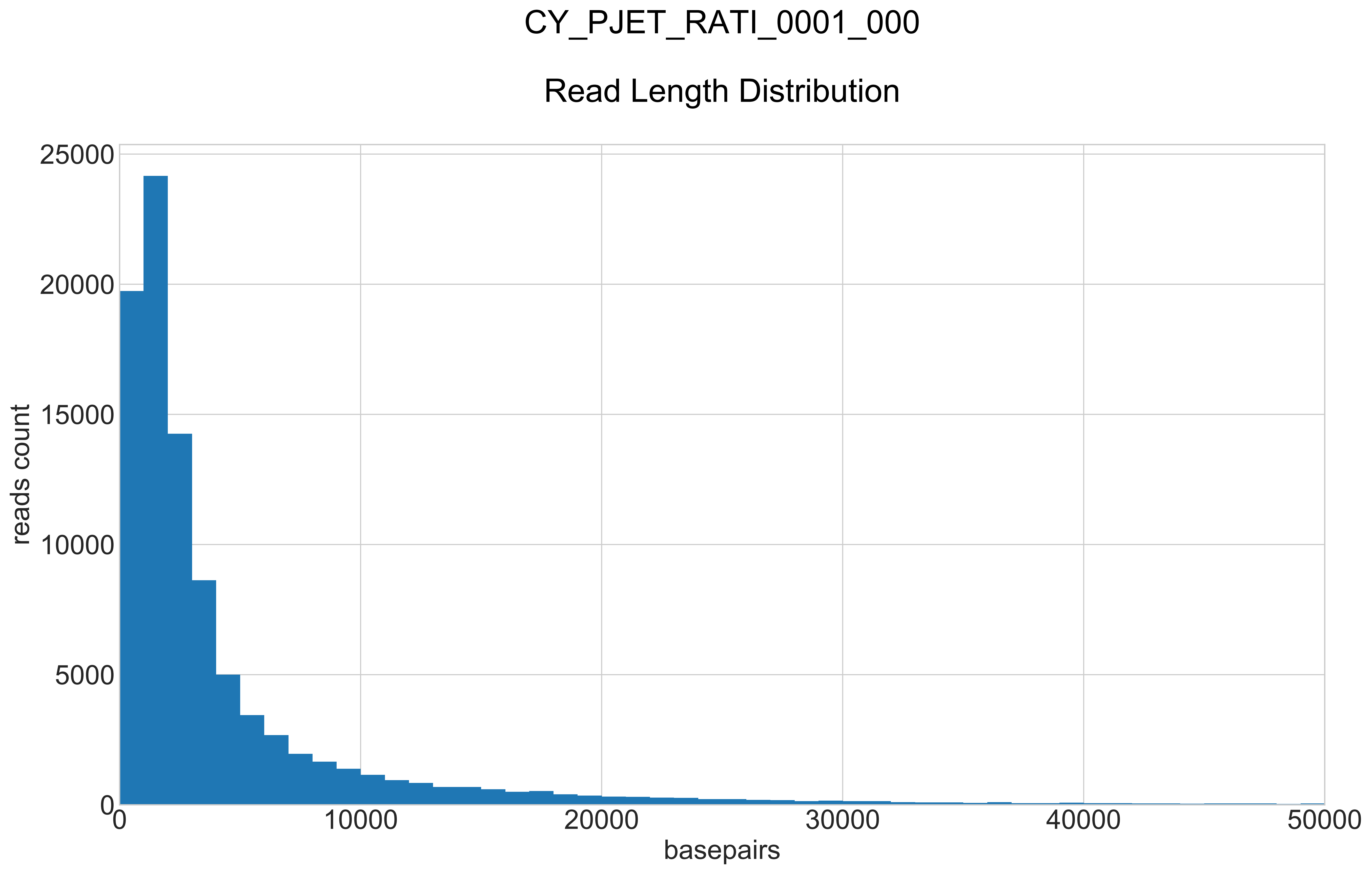

### CY_PJET_RATI_0001_000_REPEATS_COUNT_.pdf

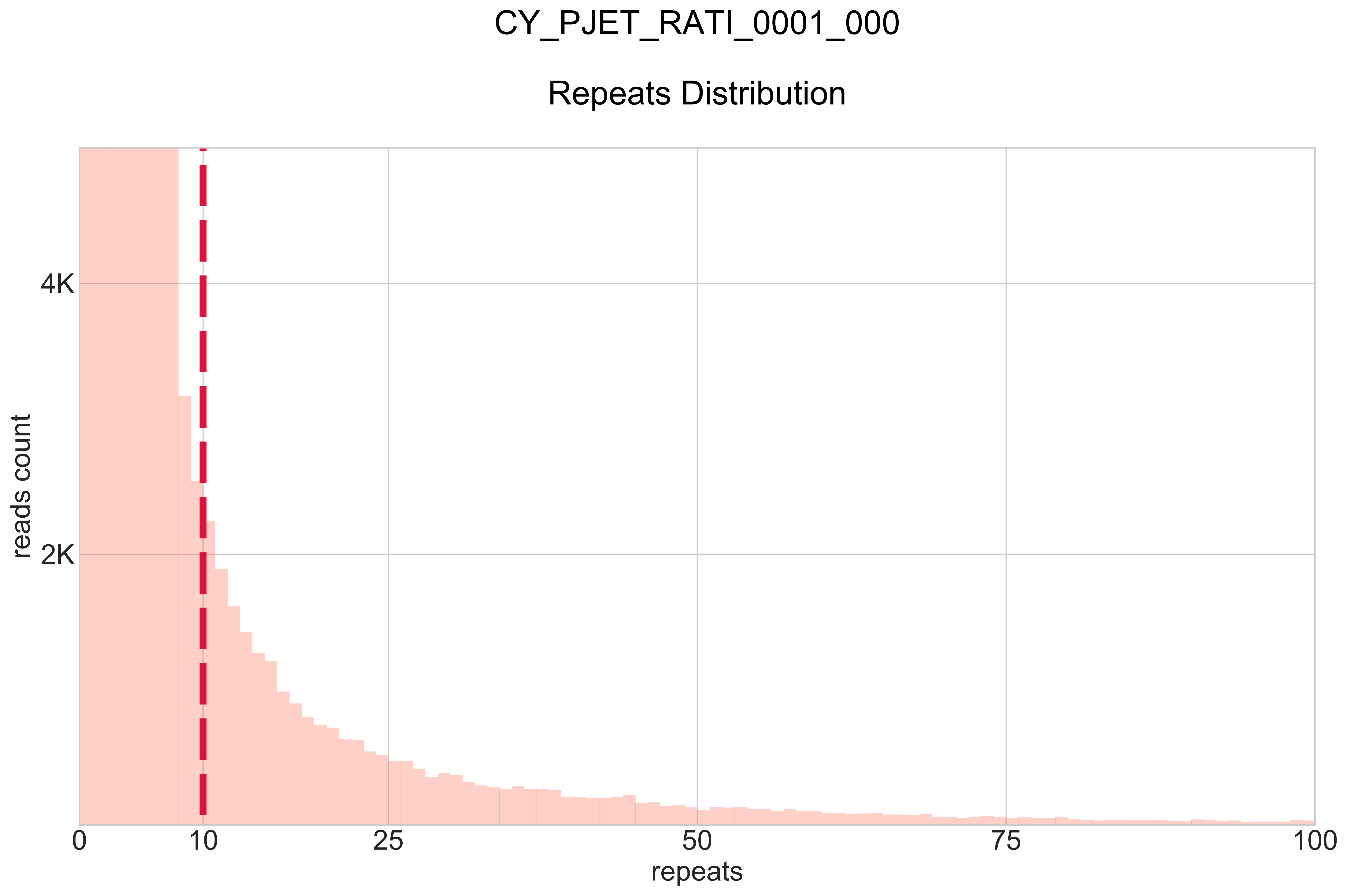

### CY_SM_PC_HC_0002_001_000_BB-I_.pdf

# CY_SM_PC_HC_0002_001_000

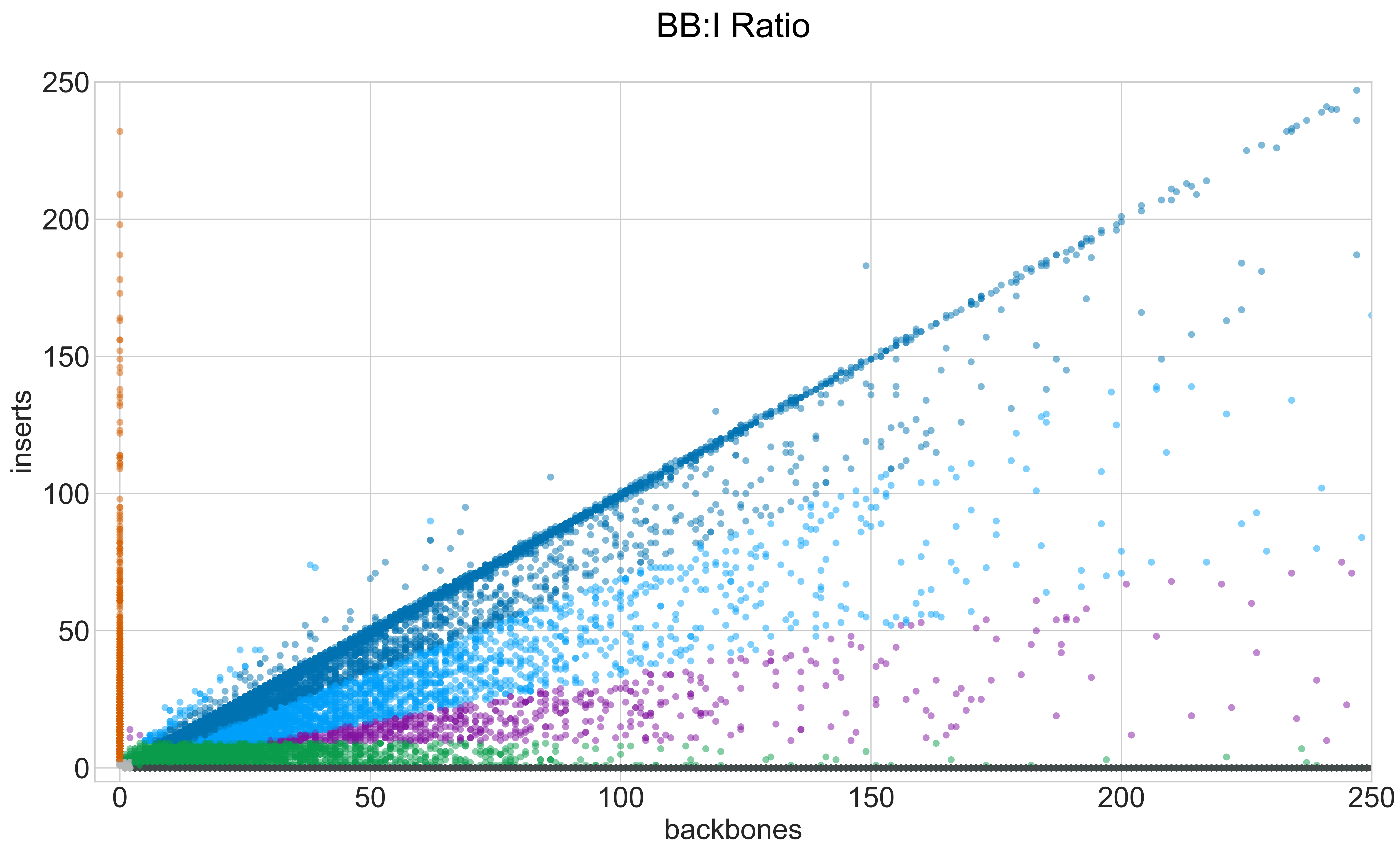

### CY_SM_PC_HC_0002_001_000_GROUPS_STACKED_NORM-LEN_.pdf

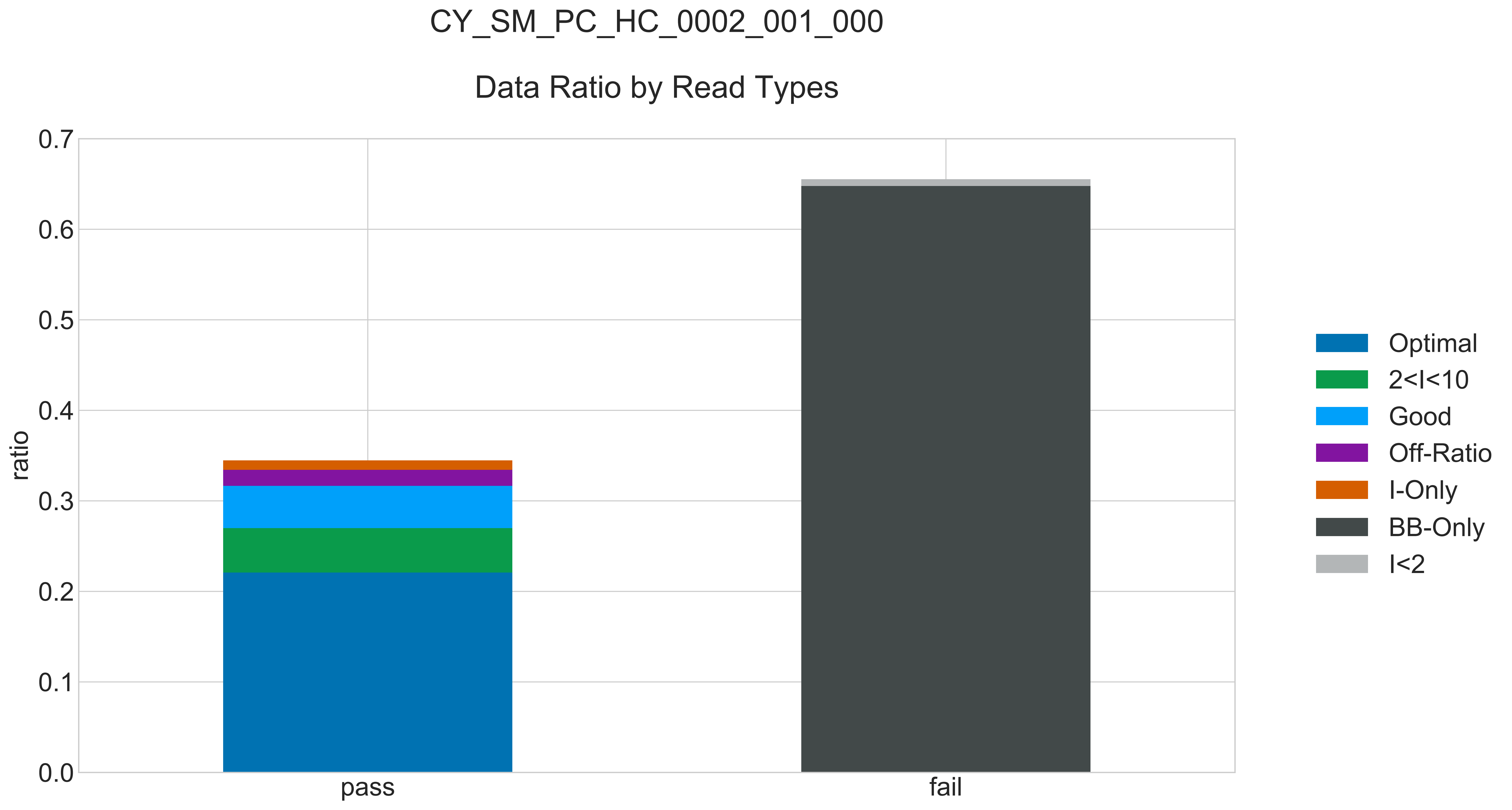

### CY_SM_PC_HC_0002_001_000_READ_LEN_.pdf

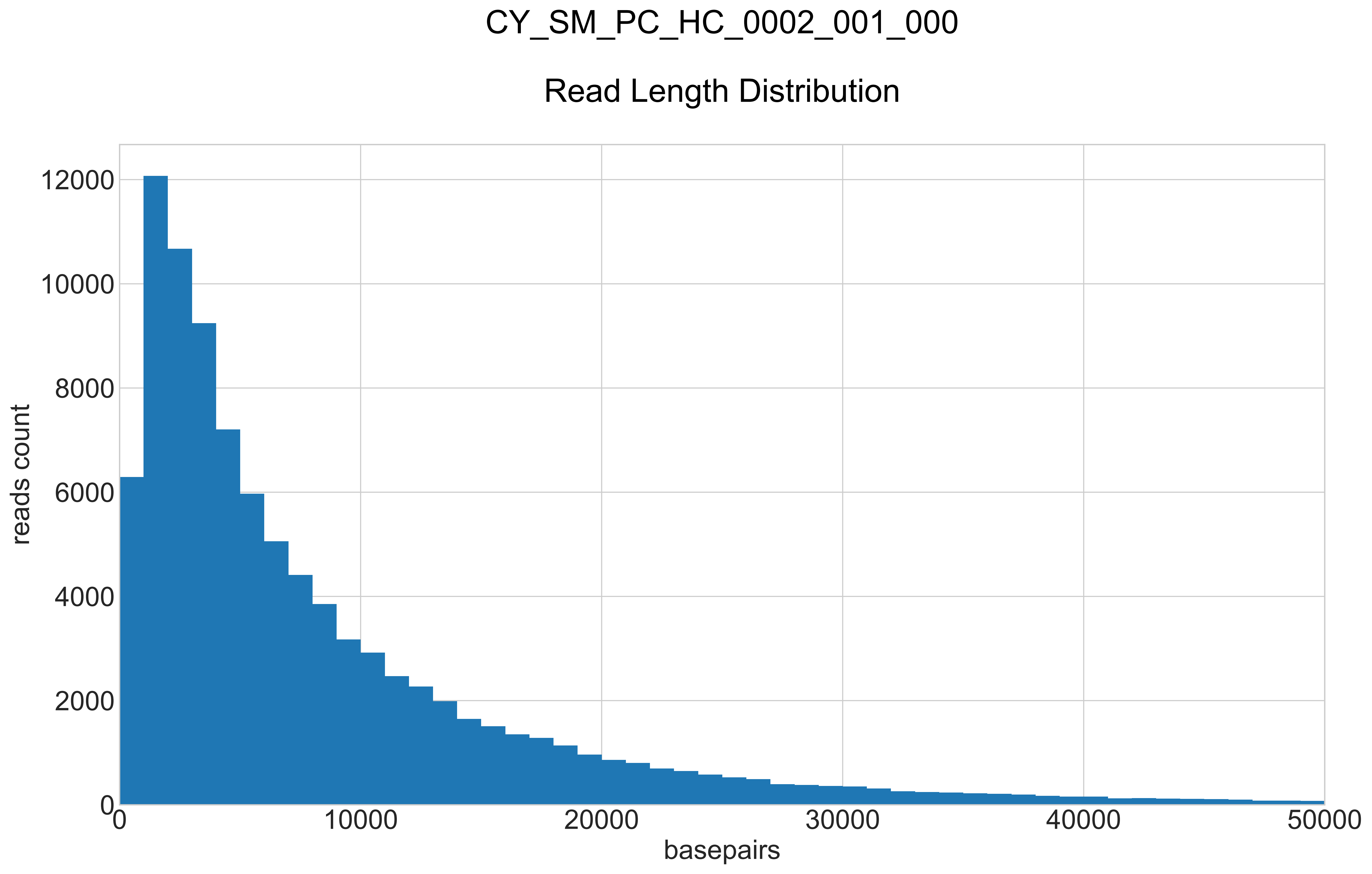

### CY_SM_PC_HC_0002_001_000_REPEATS_COUNT_.pdf

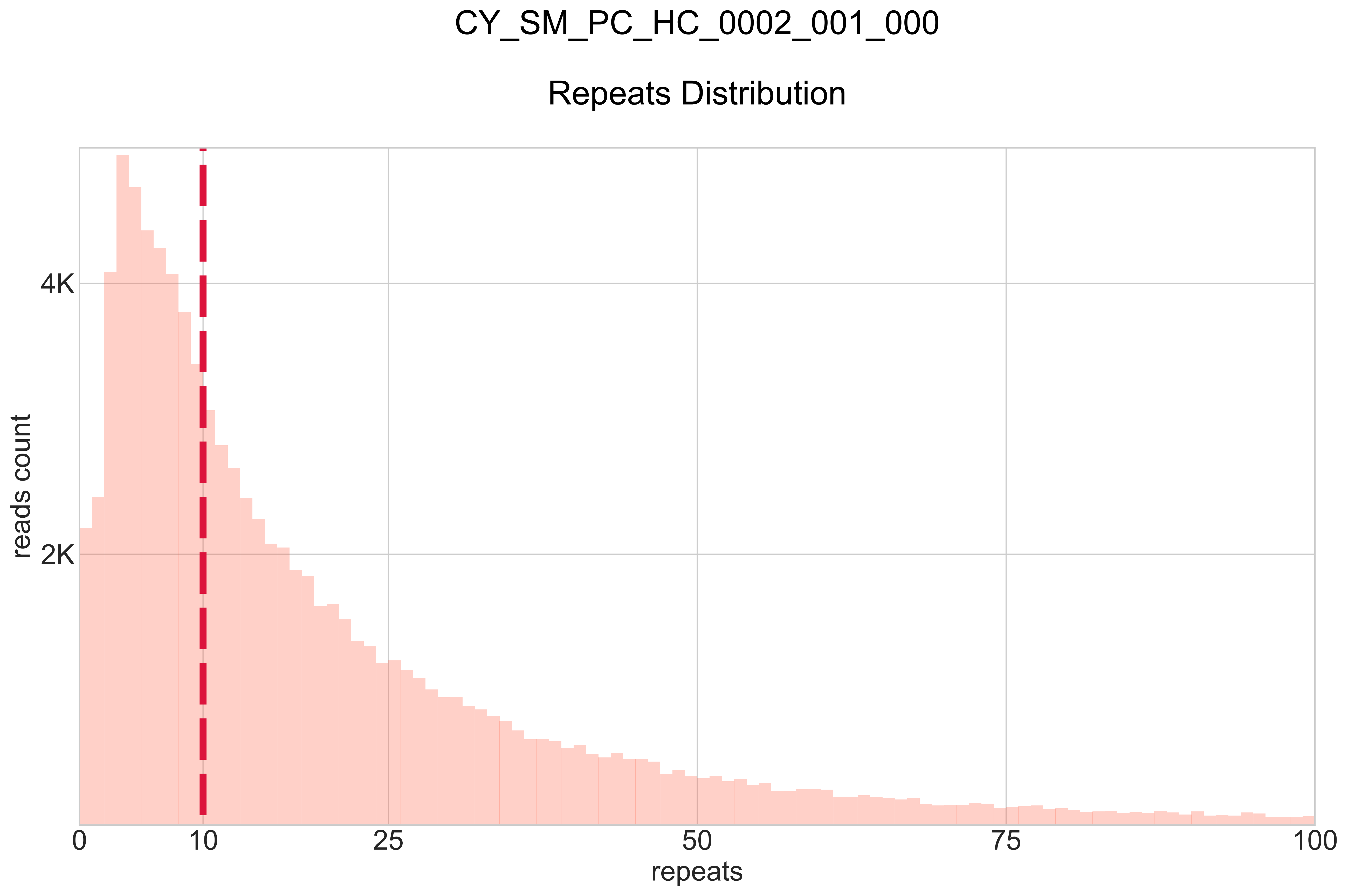

### CY_SM_PC_HC_0004_001_000_BB-I_.pdf

# CY_SM_PC_HC_0004_001_000

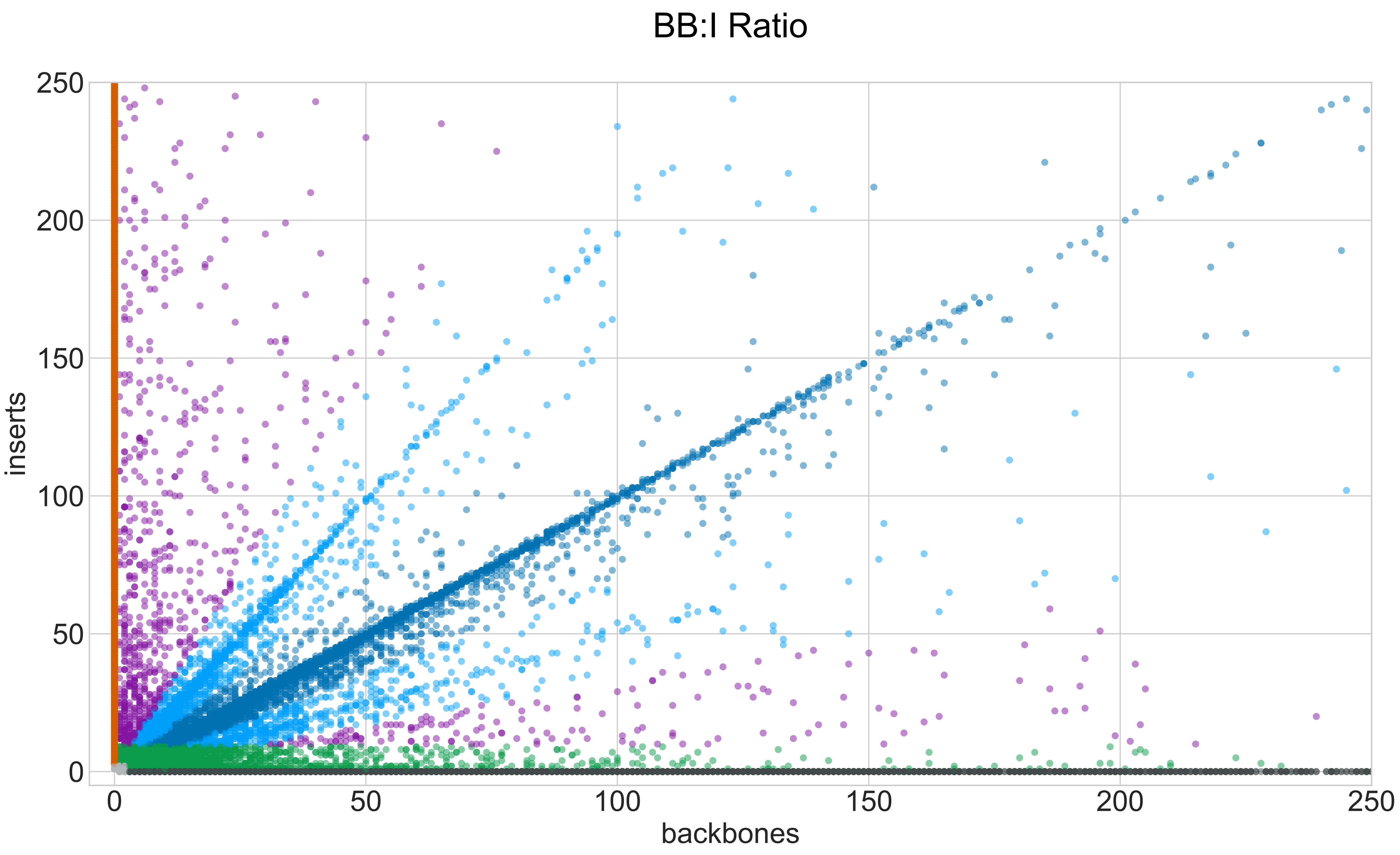

### CY_SM_PC_HC_0004_001_000_GROUPS_STACKED_NORM-LEN_.pdf

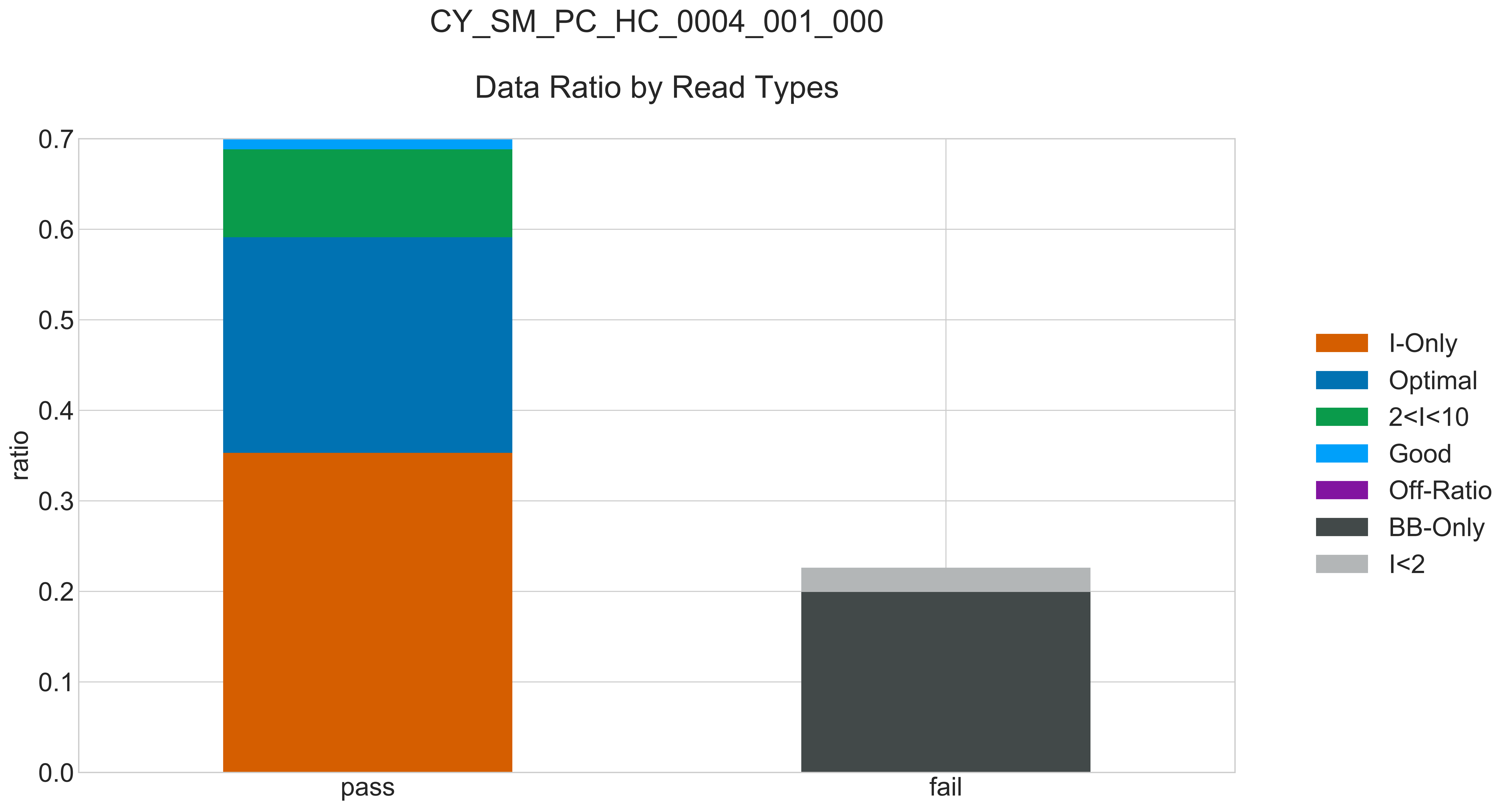

### CY_SM_PC_HC_0004_001_000_READ_LEN_.pdf

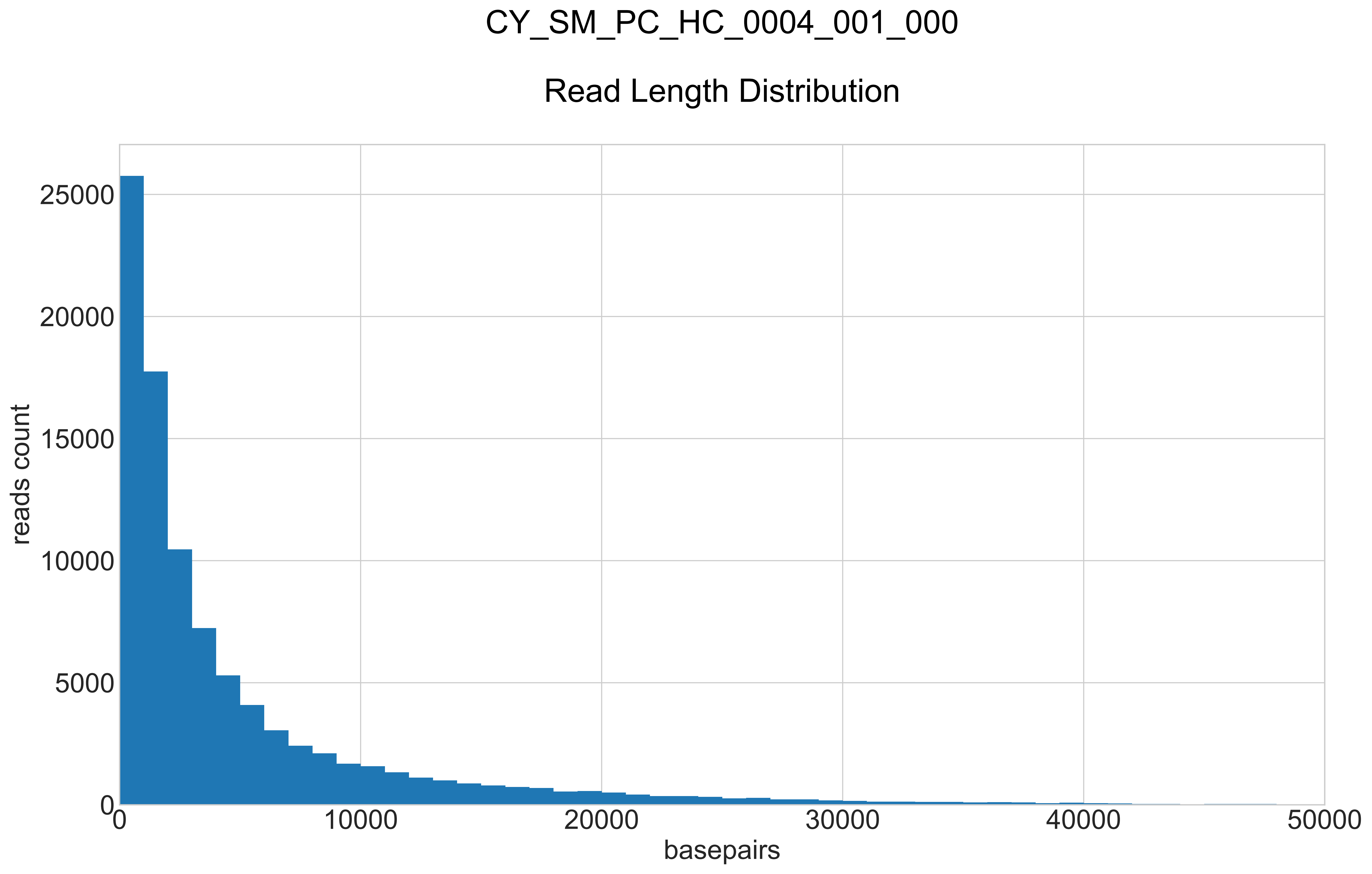

### CY_SM_PC_HC_0004_001_000_REPEATS_COUNT_.pdf

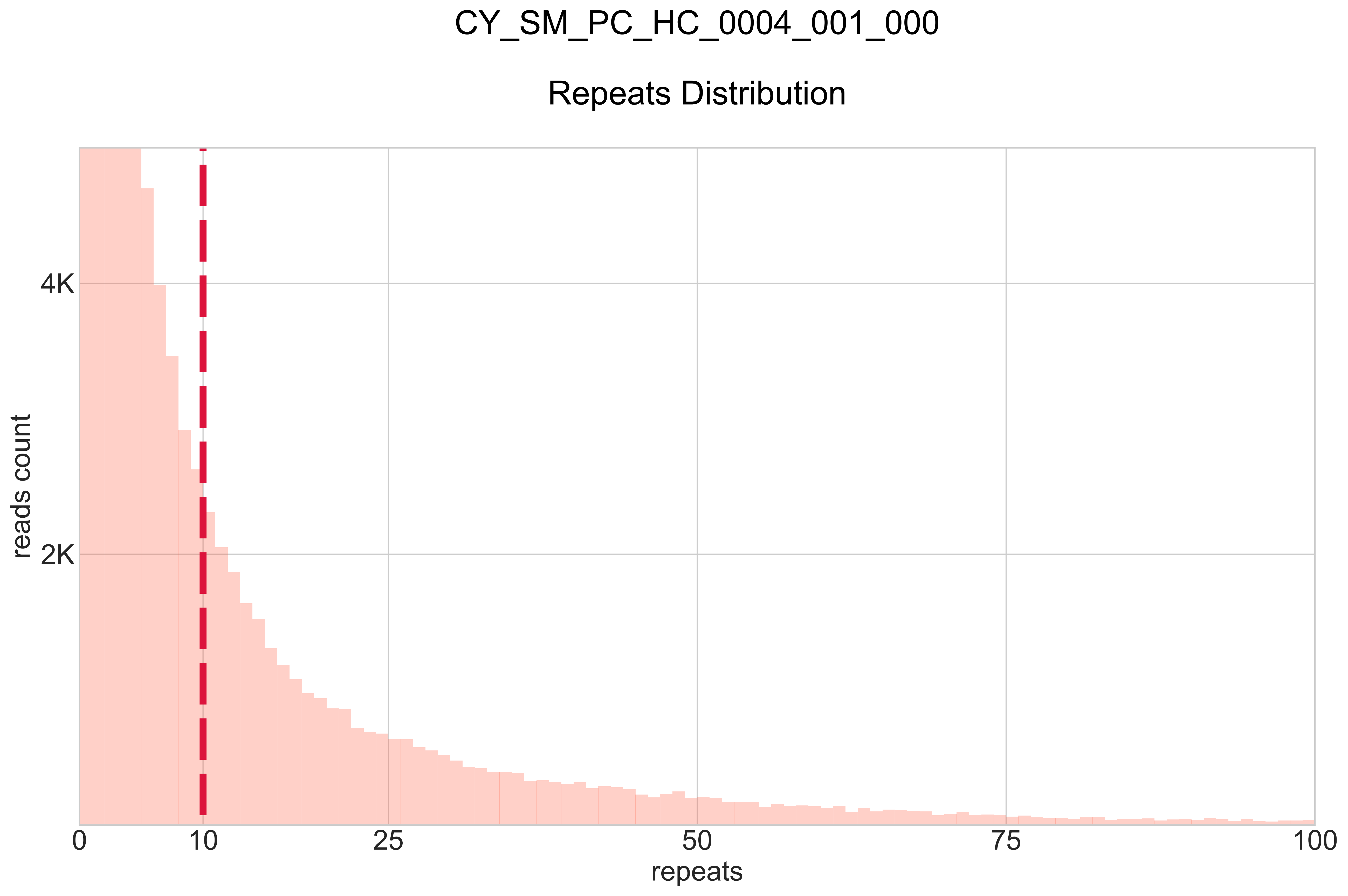

### CY_SM_PC_HC_0004_002_000_BB-I_.pdf

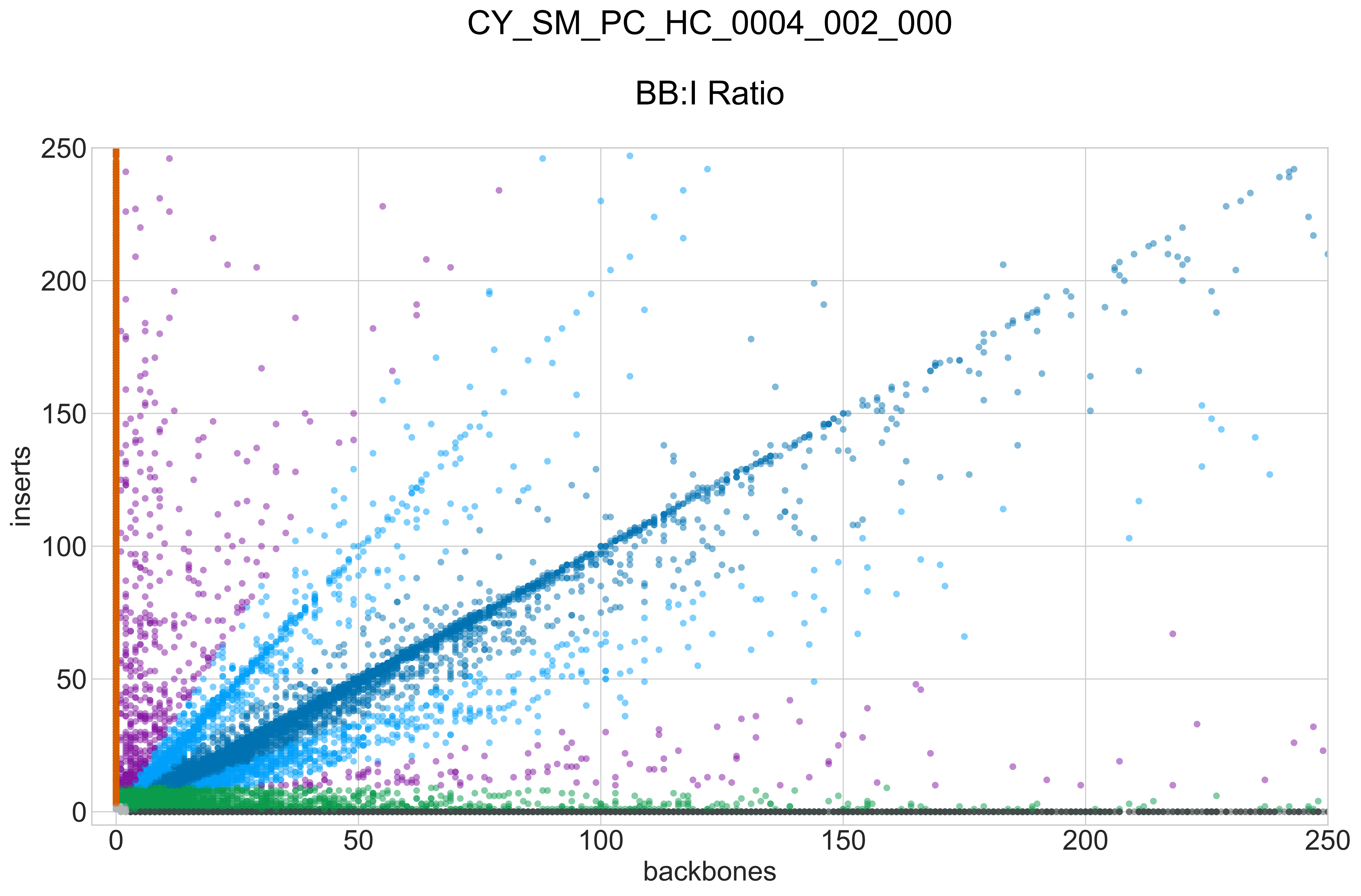

### CY_SM_PC_HC_0004_002_000_GROUPS_STACKED_NORM-LEN_.pdf

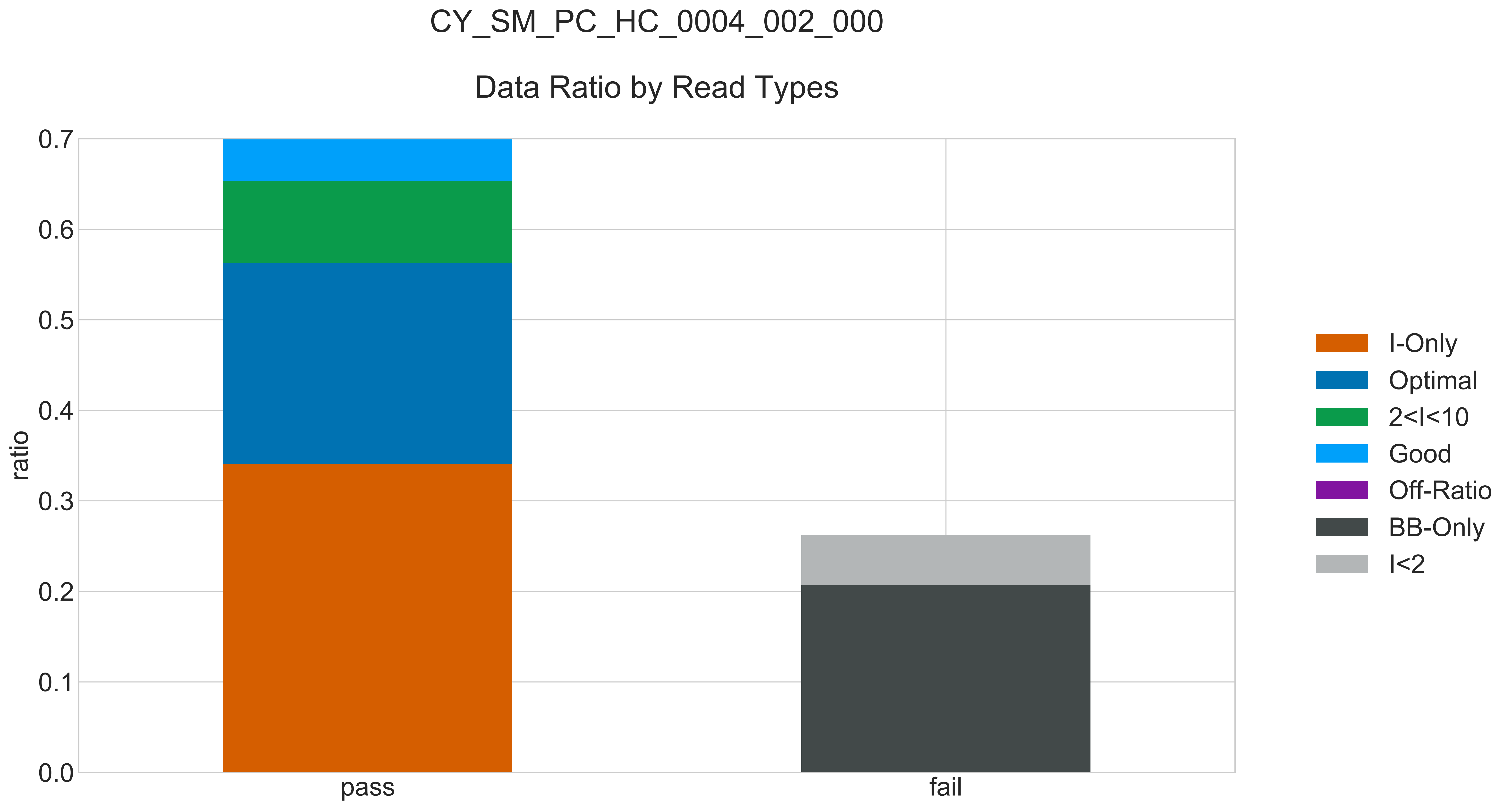

### CY_SM_PC_HC_0004_002_000_READ_LEN_.pdf

# CY_SM_PC_HC_0004_002_000

##### Read Length Distribution

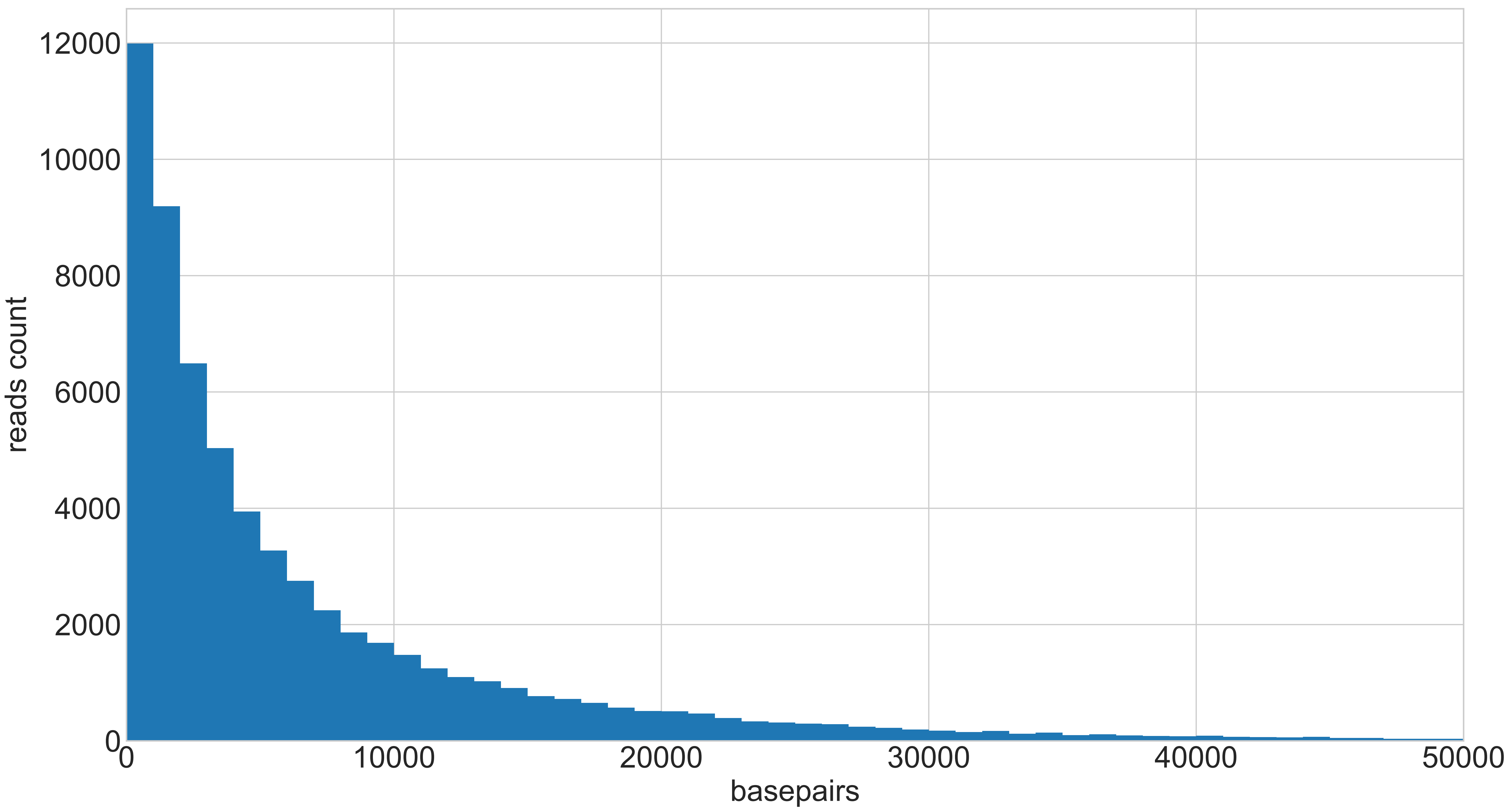

### CY_SM_PC_HC_0004_002_000_REPEATS_COUNT_.pdf

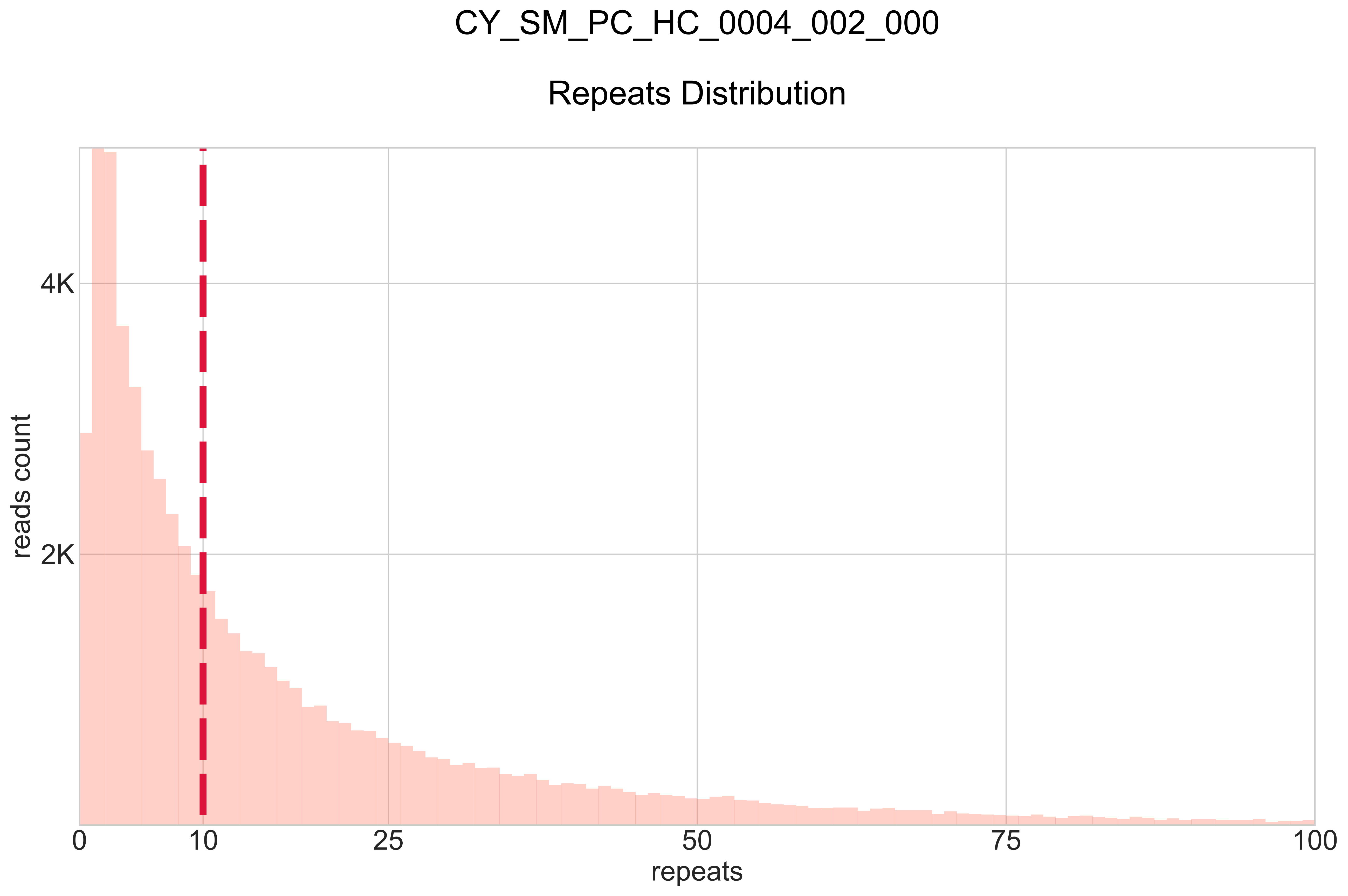

### CY_SM_PC_HC_0004_003_000_BB-I_.pdf

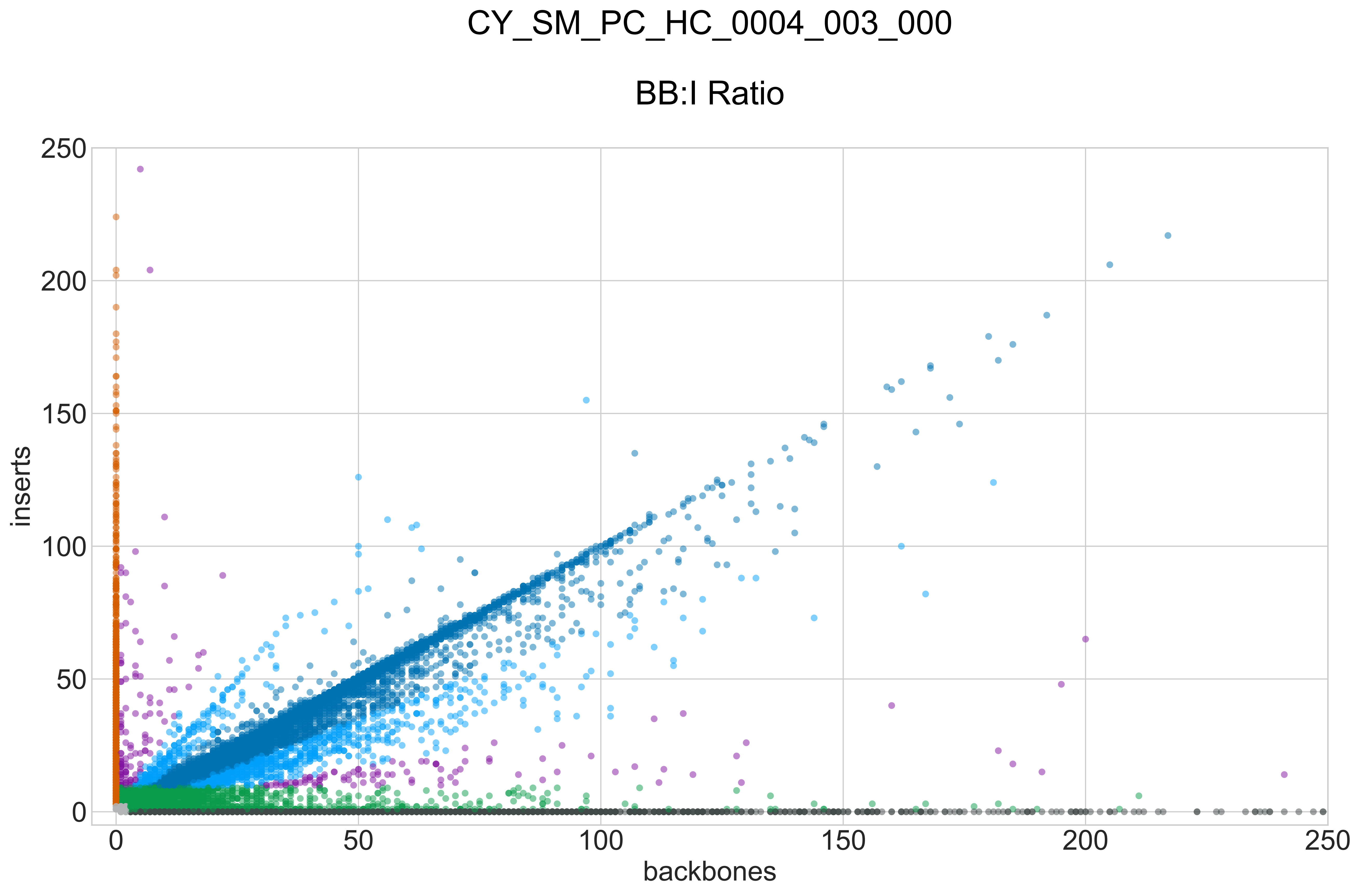

### CY_SM_PC_HC_0004_003_000_GROUPS_STACKED_NORM-LEN_.pdf

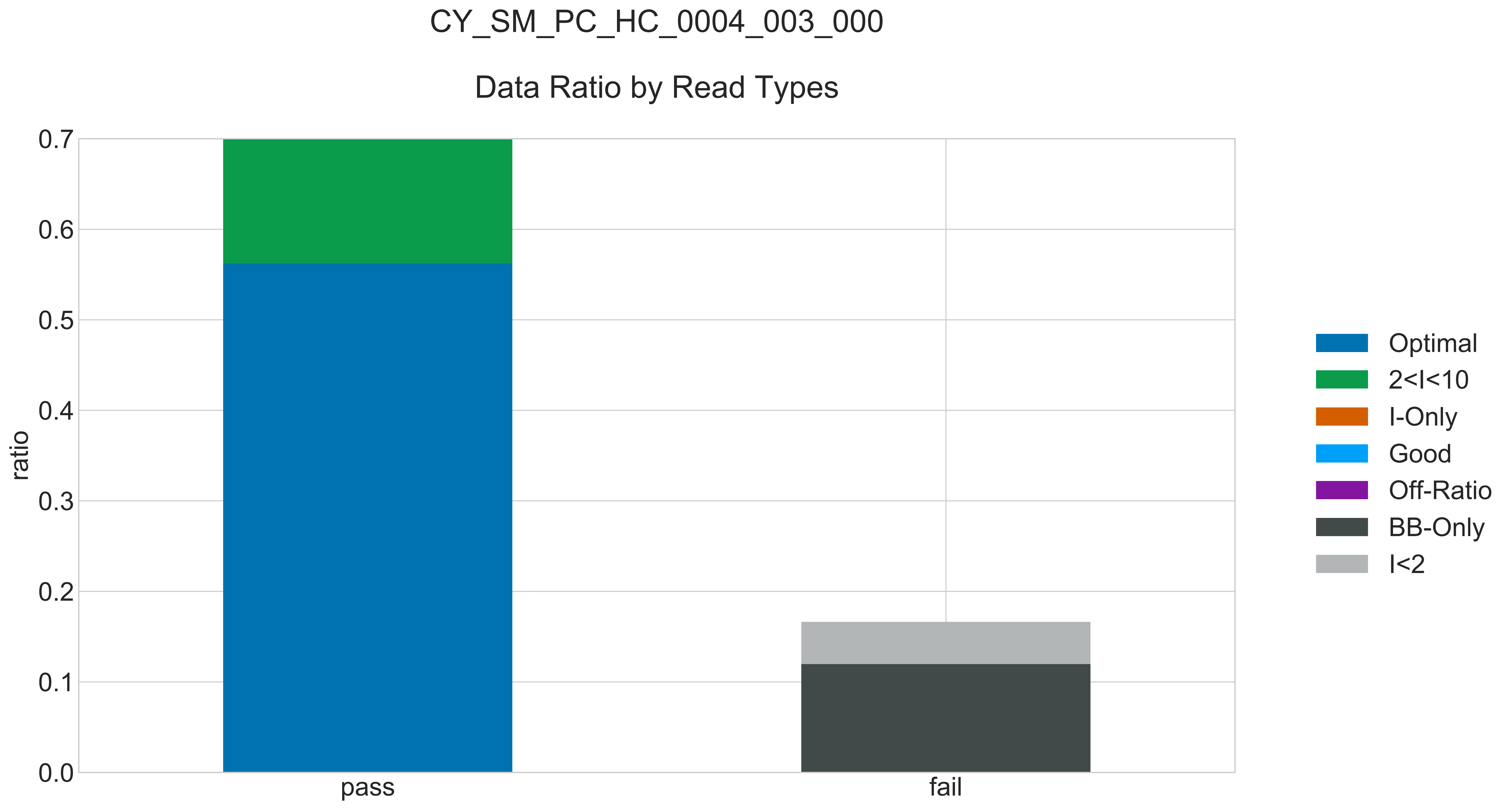

### CY_SM_PC_HC_0004_003_000_READ_LEN_.pdf

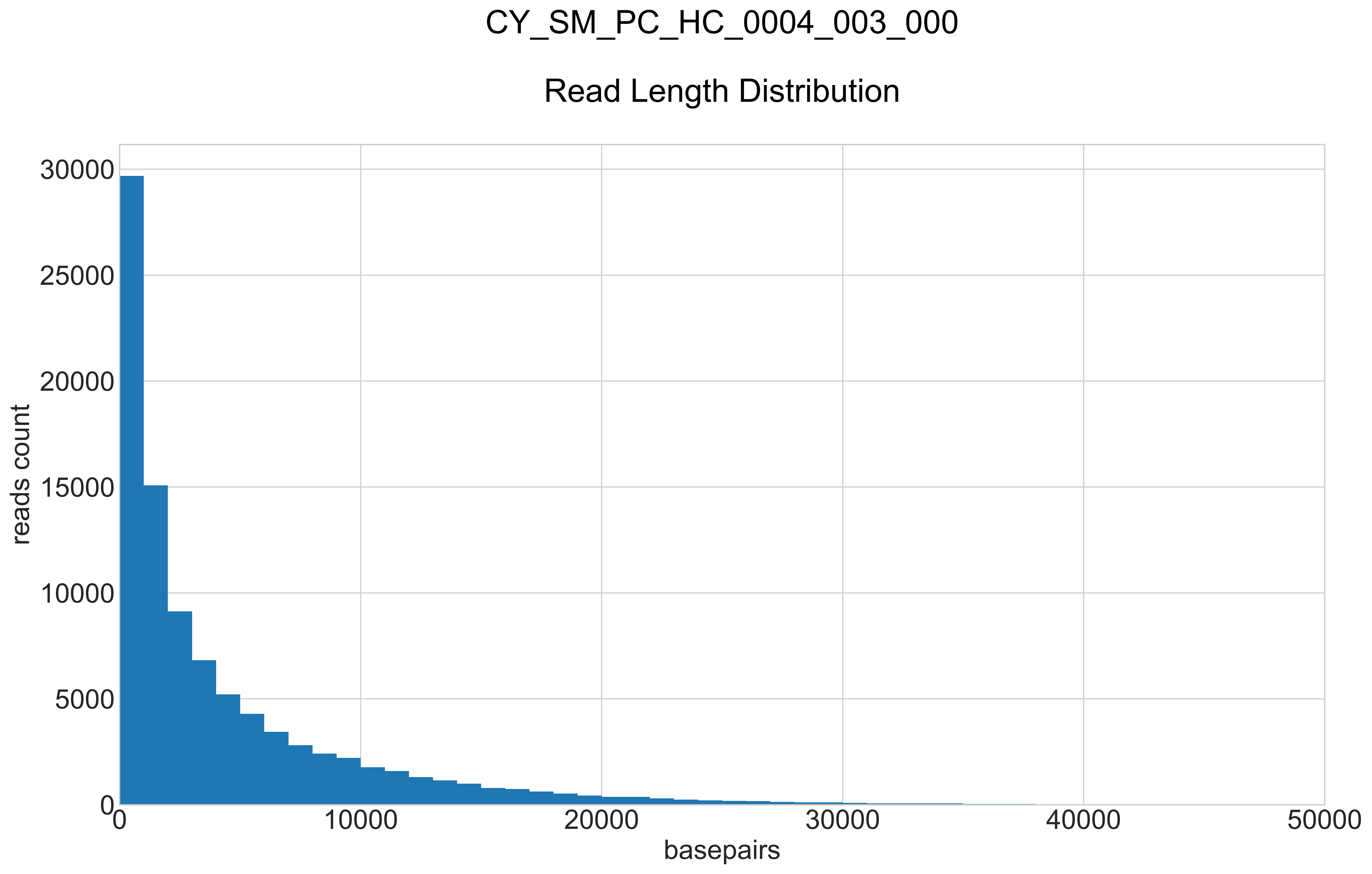

### CY_SM_PC_HC_0004_003_000_REPEATS_COUNT_.pdf

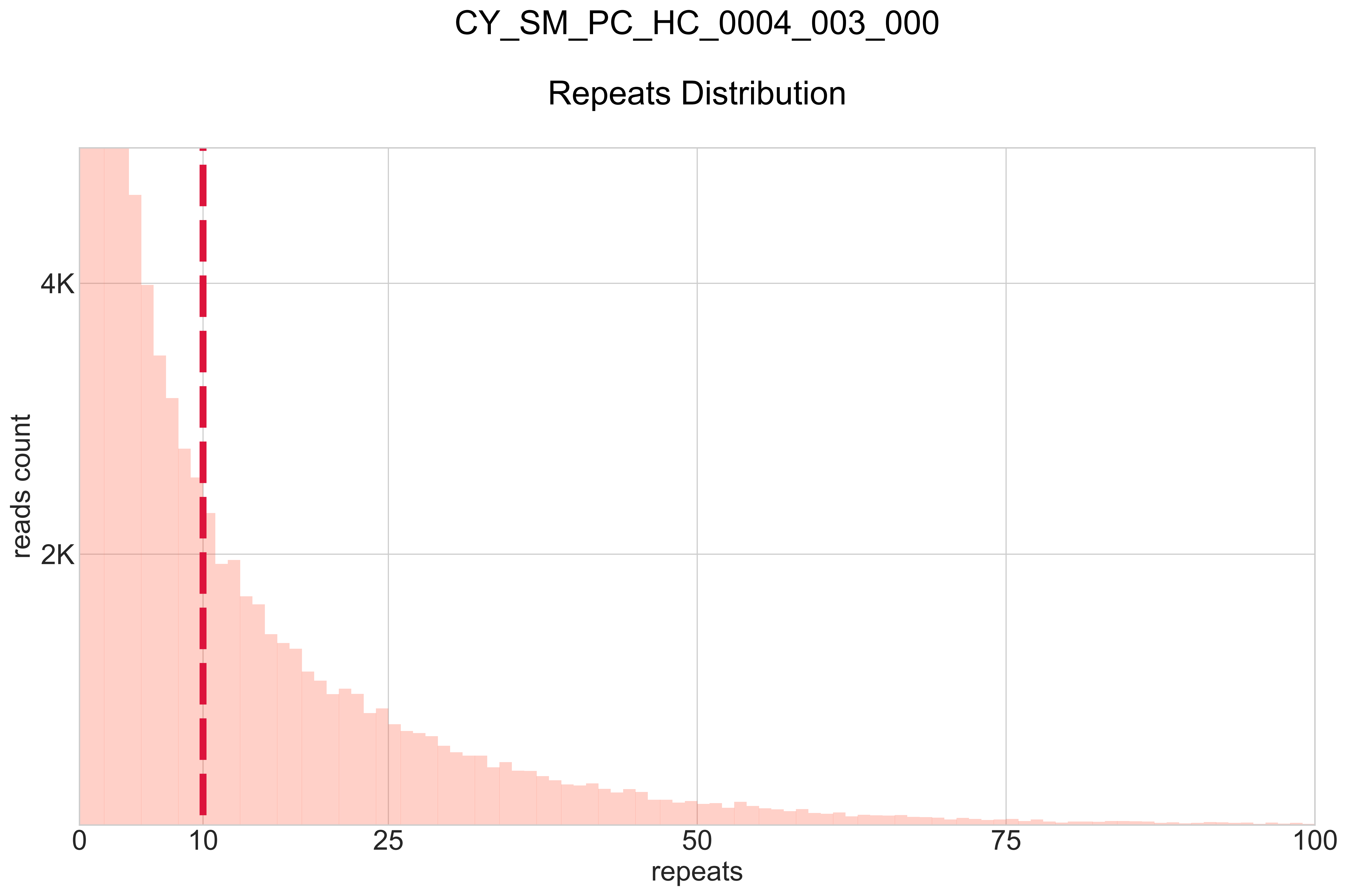

### CY_SM_PC_HC_0004_004_000_BB-I_.pdf

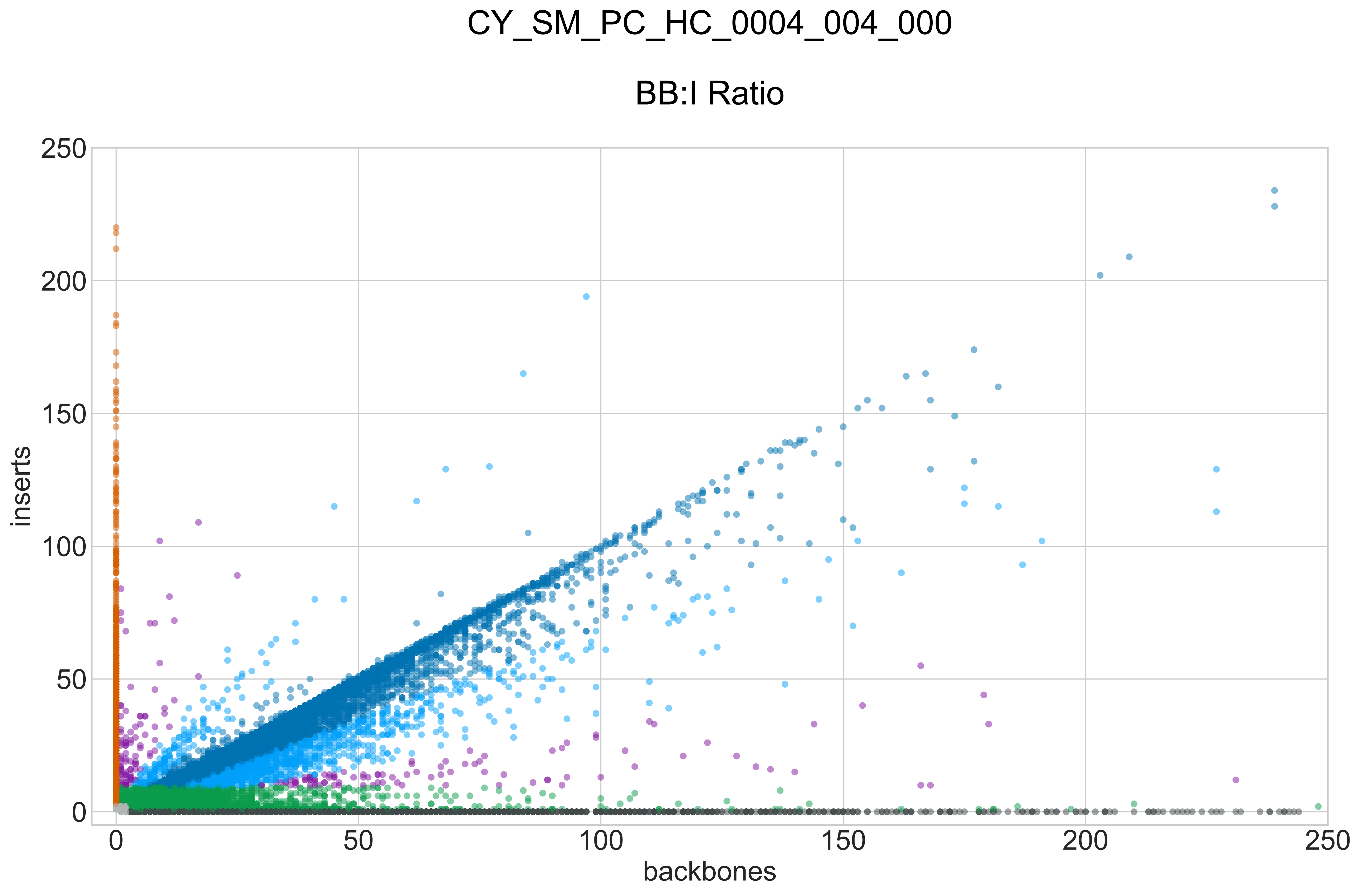

### CY_SM_PC_HC_0004_004_000_GROUPS_STACKED_NORM-LEN_.pdf

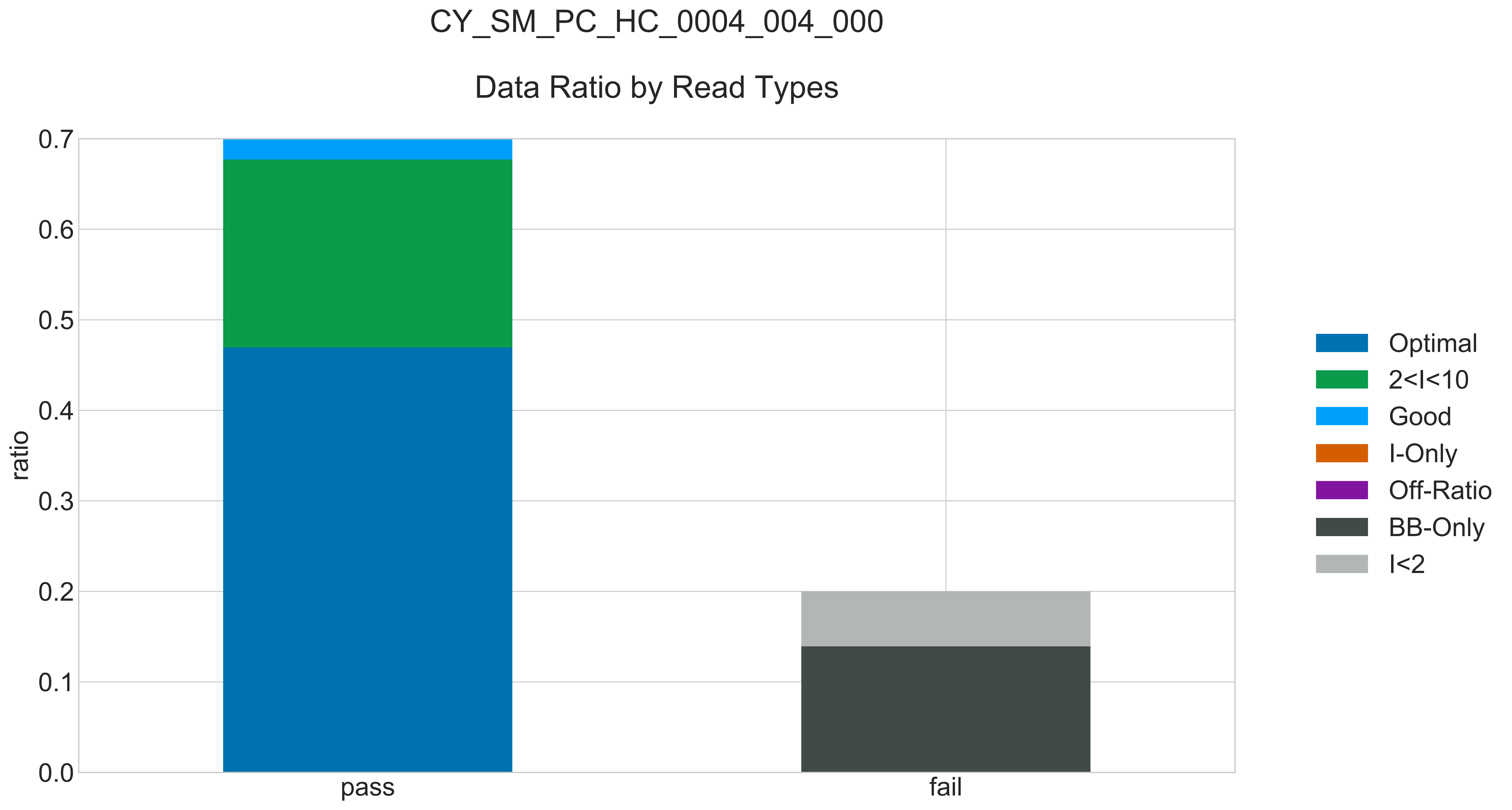

### CY_SM_PC_HC_0004_004_000_READ_LEN_.pdf

# CY_SM_PC_HC_0004_004_000

##### Read Length Distribution

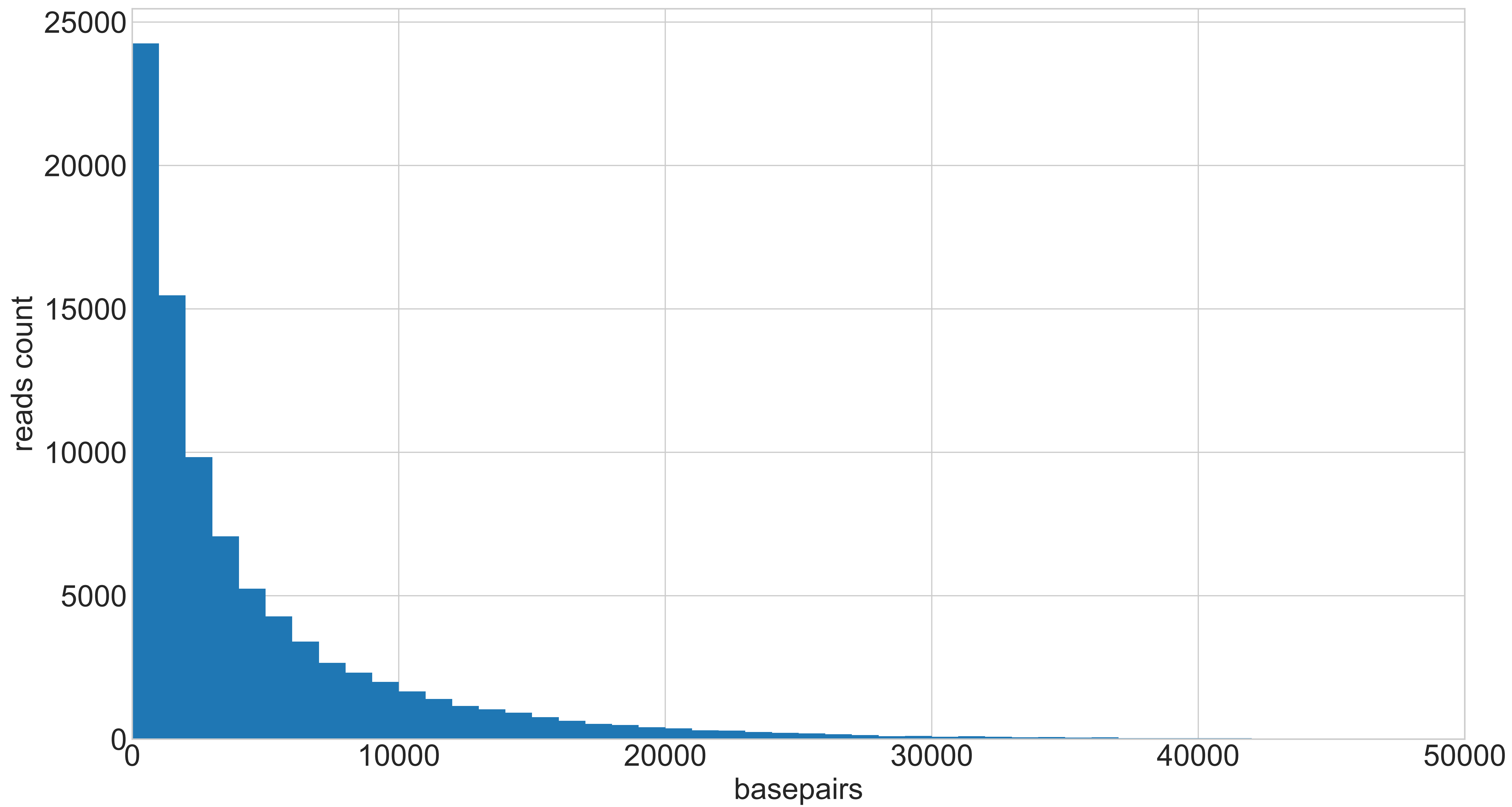

### CY_SM_PC_HC_0004_004_000_REPEATS_COUNT_.pdf

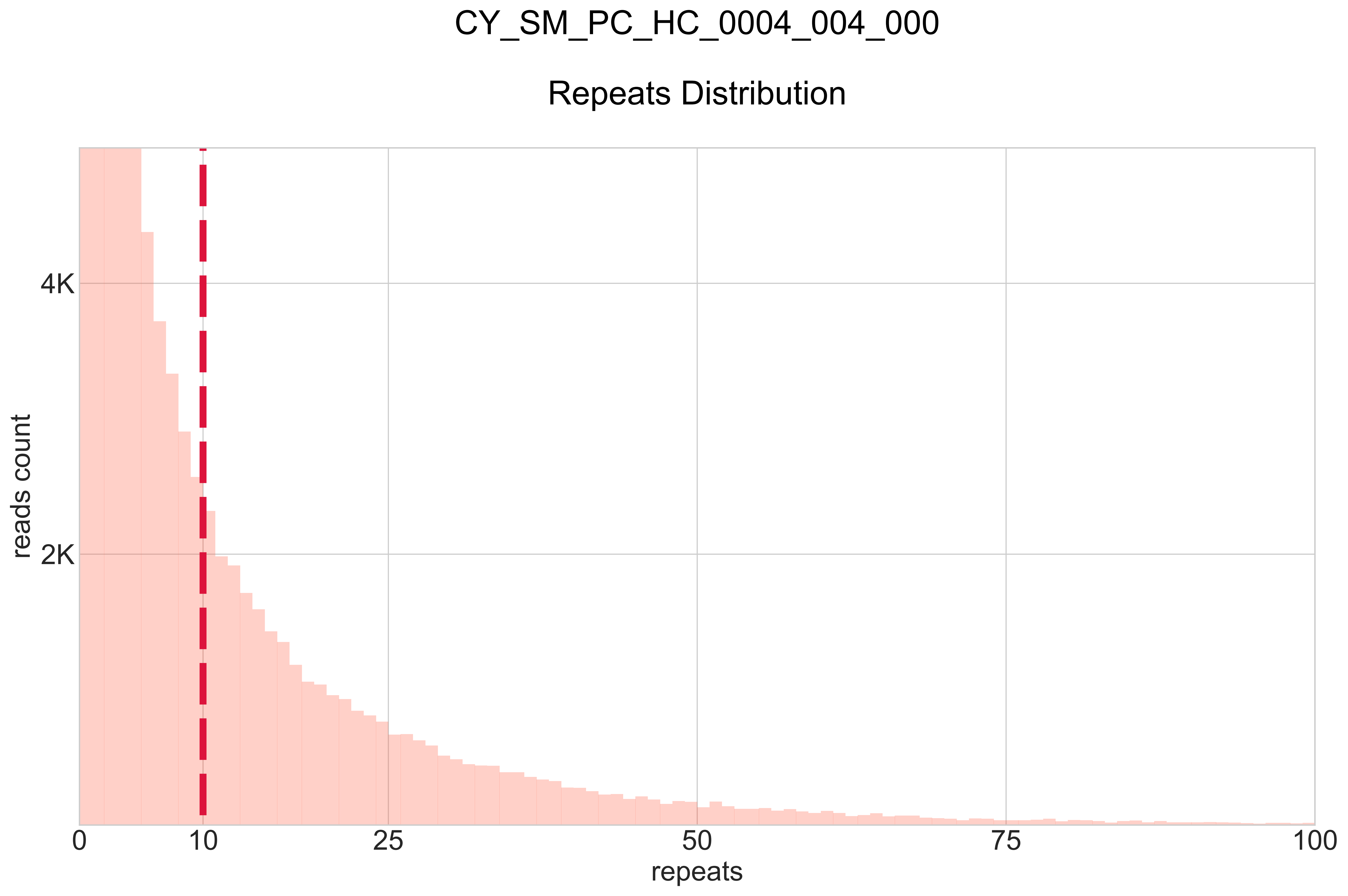

### CY_SM_PC_HN_0002_001_000_BB-I_.pdf

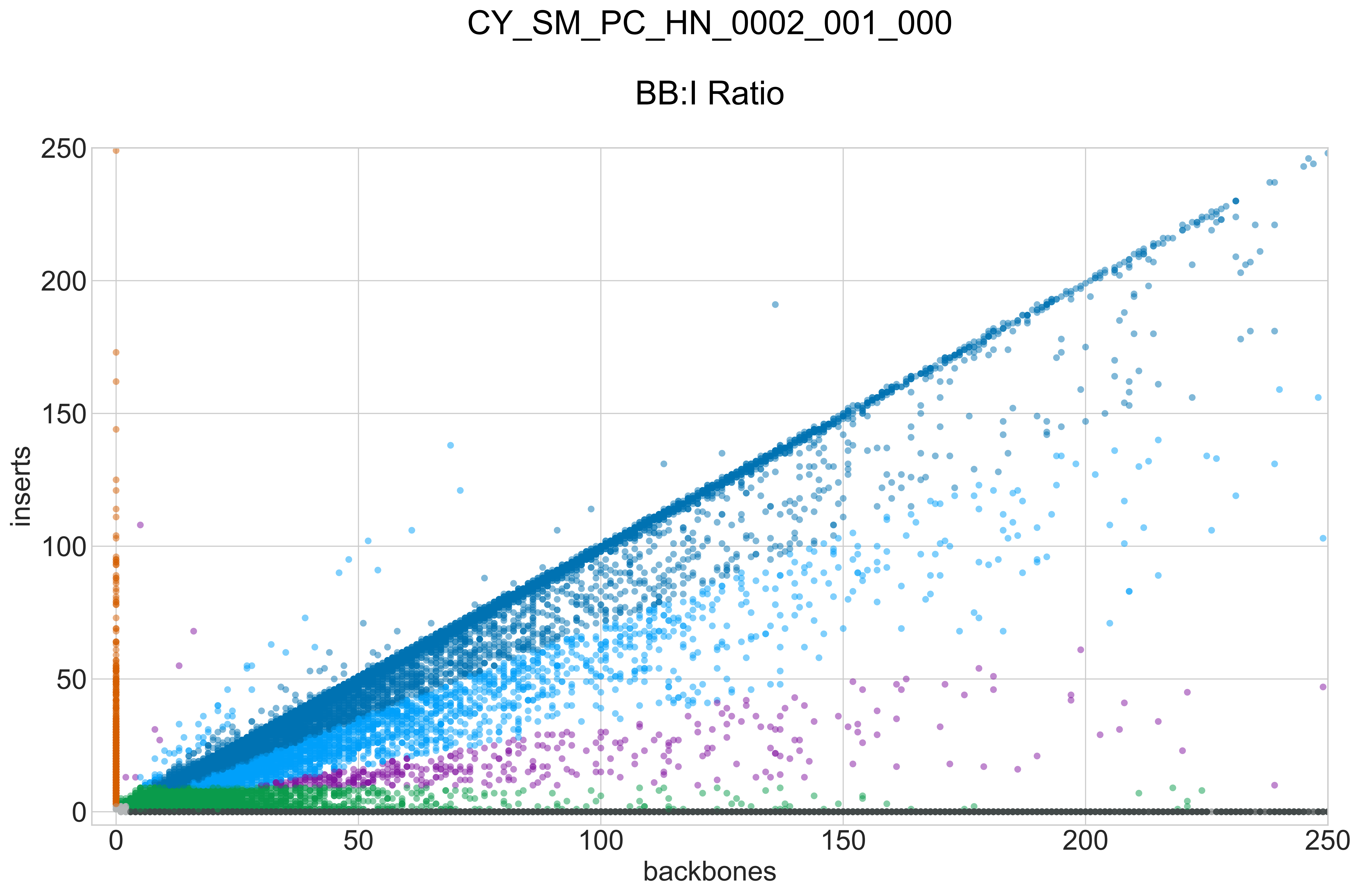

### CY_SM_PC_HN_0002_001_000_GROUPS_STACKED_NORM-LEN_.pdf

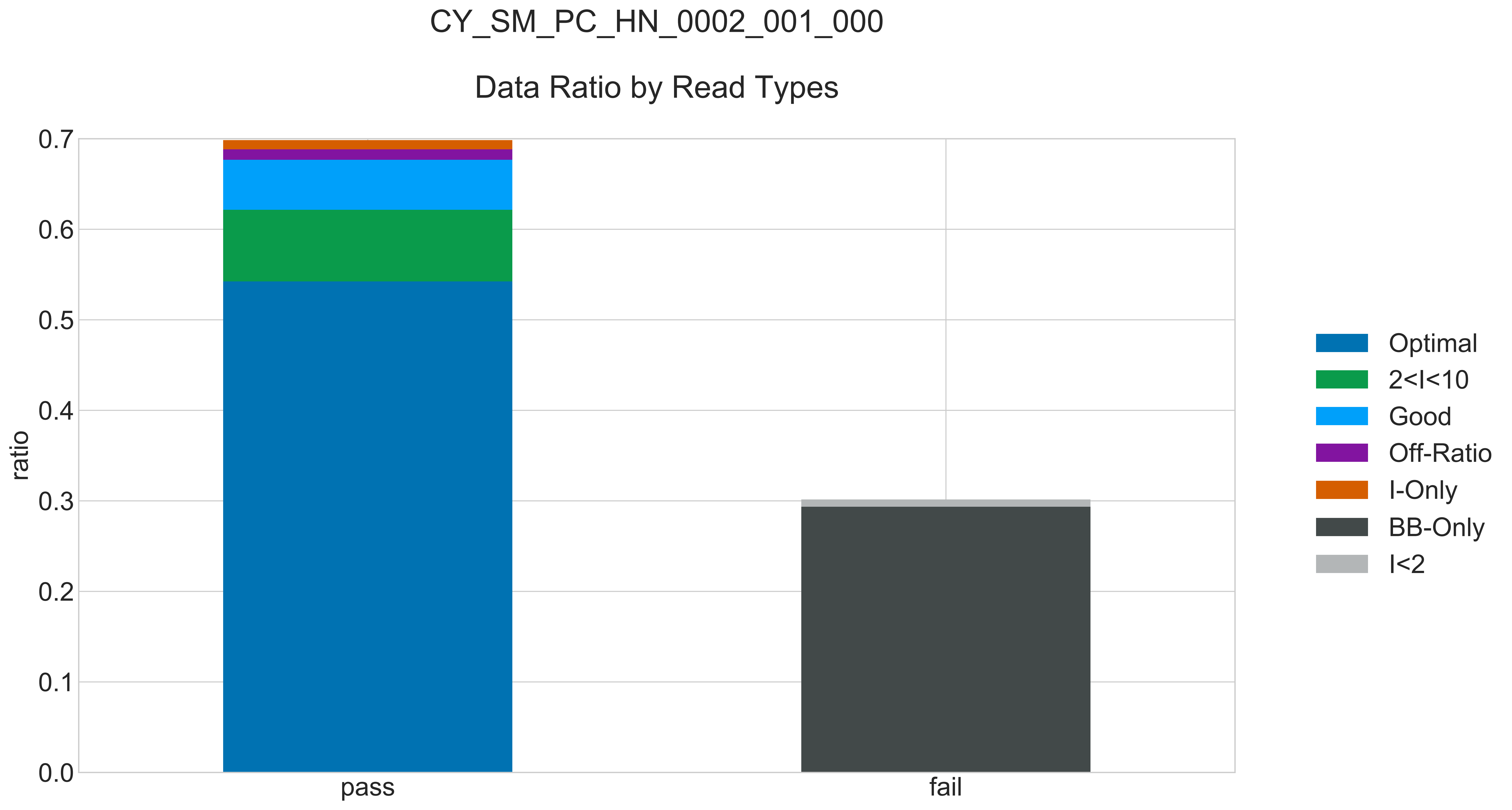

### CY_SM_PC_HN_0002_001_000_READ_LEN_.pdf

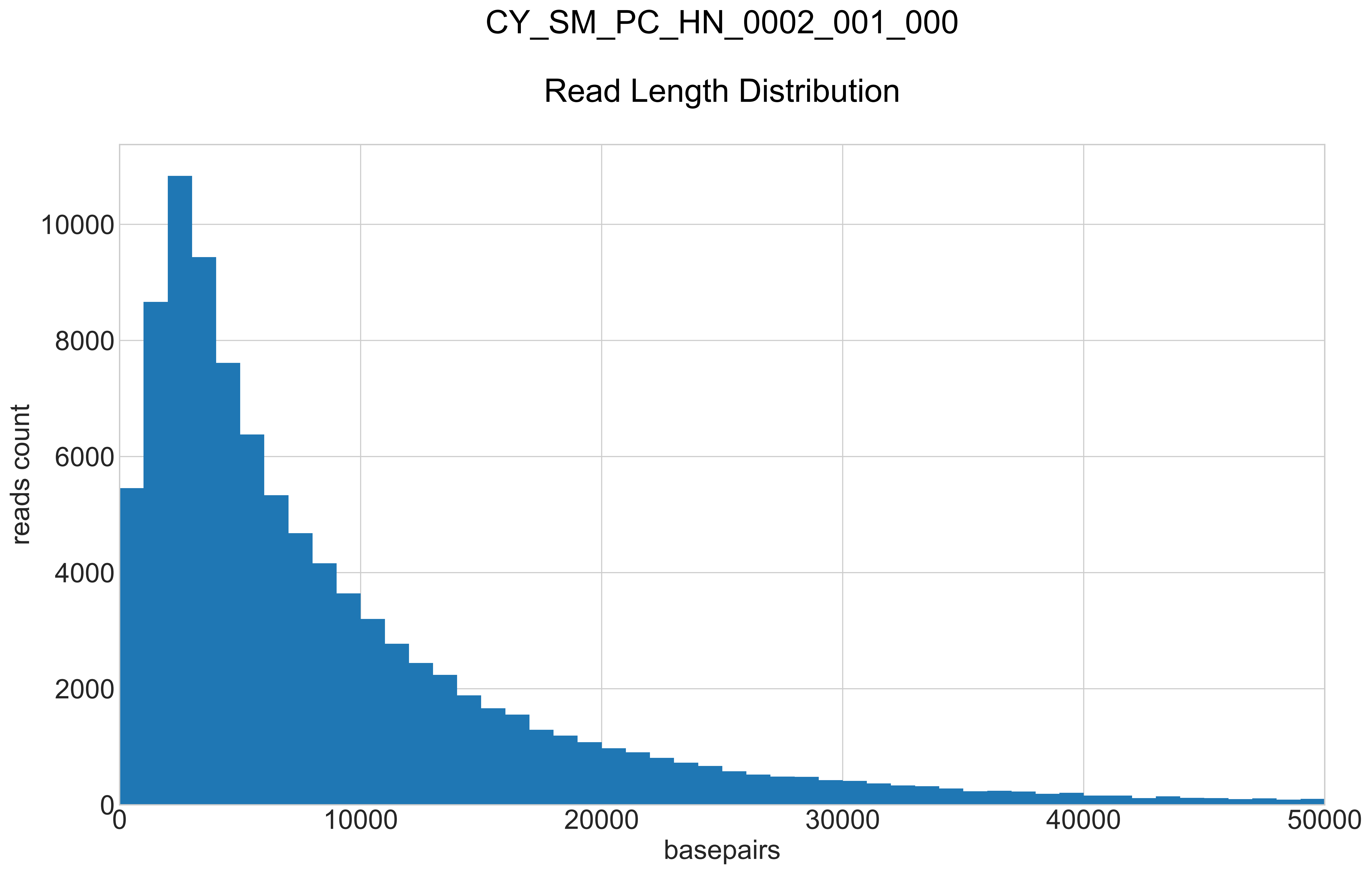

### CY_SM_PC_HN_0002_001_000_REPEATS_COUNT_.pdf

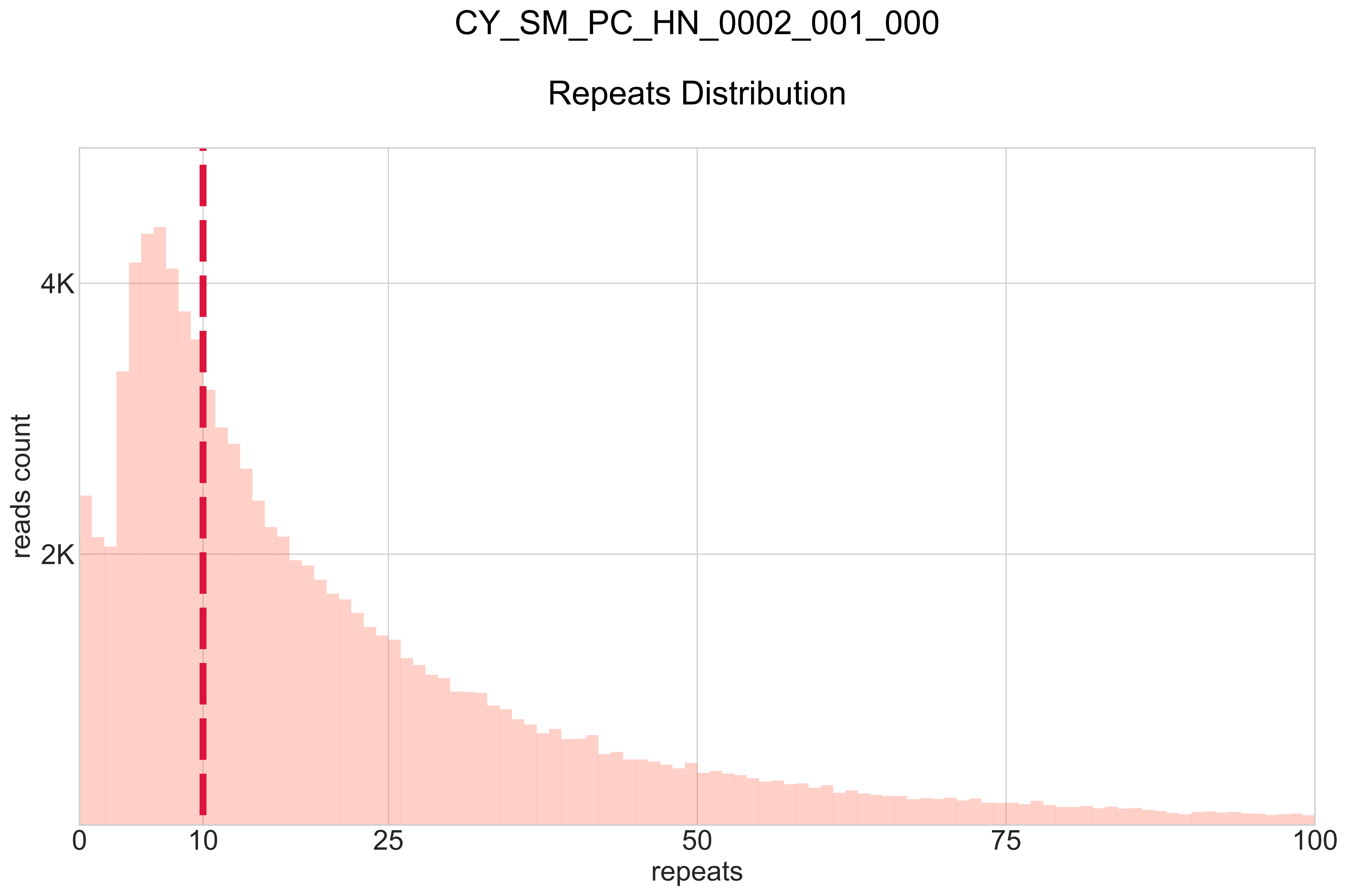

### CY_SM_PC_HN_0002_002_001_BB-I_.pdf

# CY_SM_PC_HN_0002_002_001

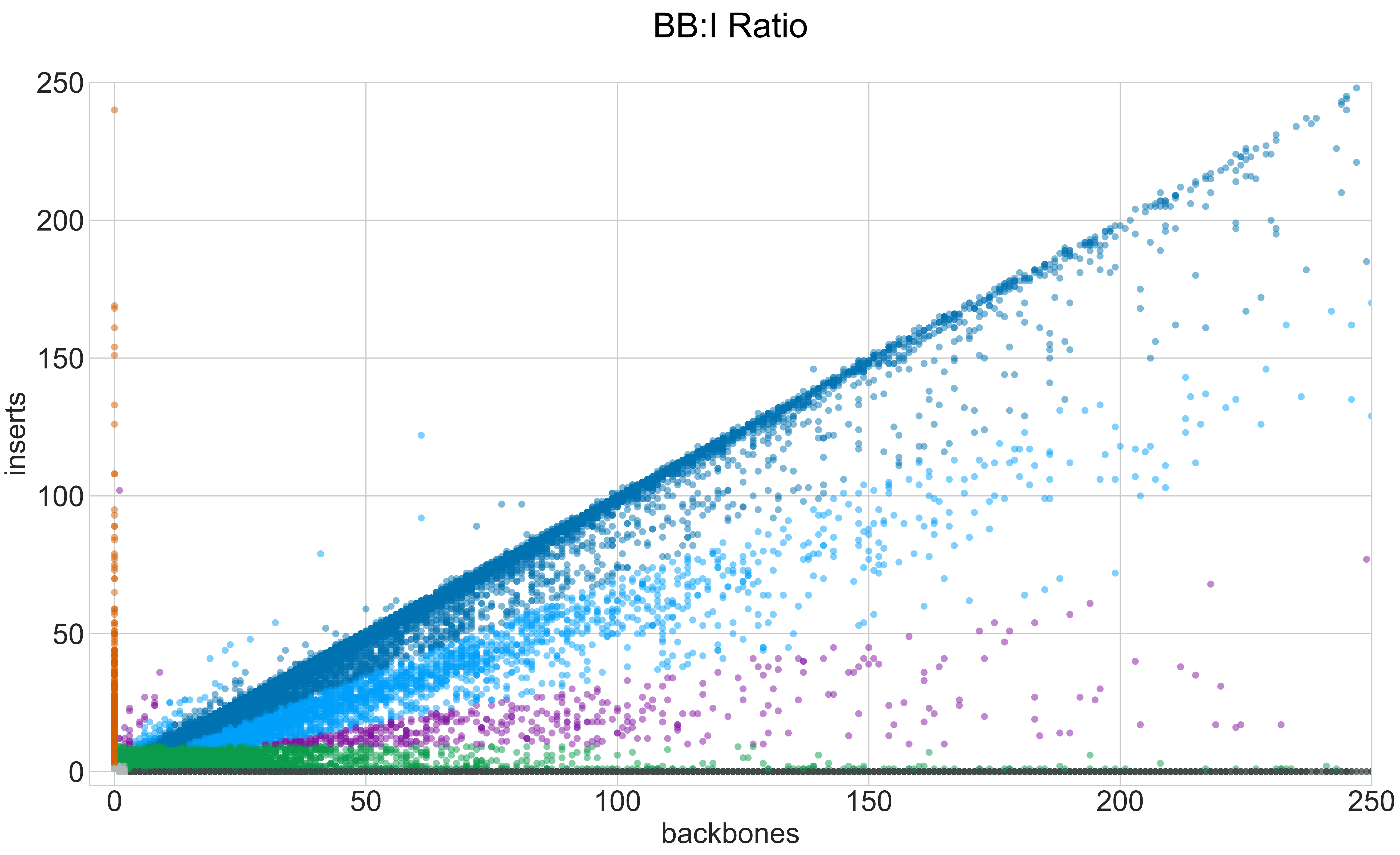

### CY_SM_PC_HN_0002_002_001_GROUPS_STACKED_NORM-LEN_.pdf

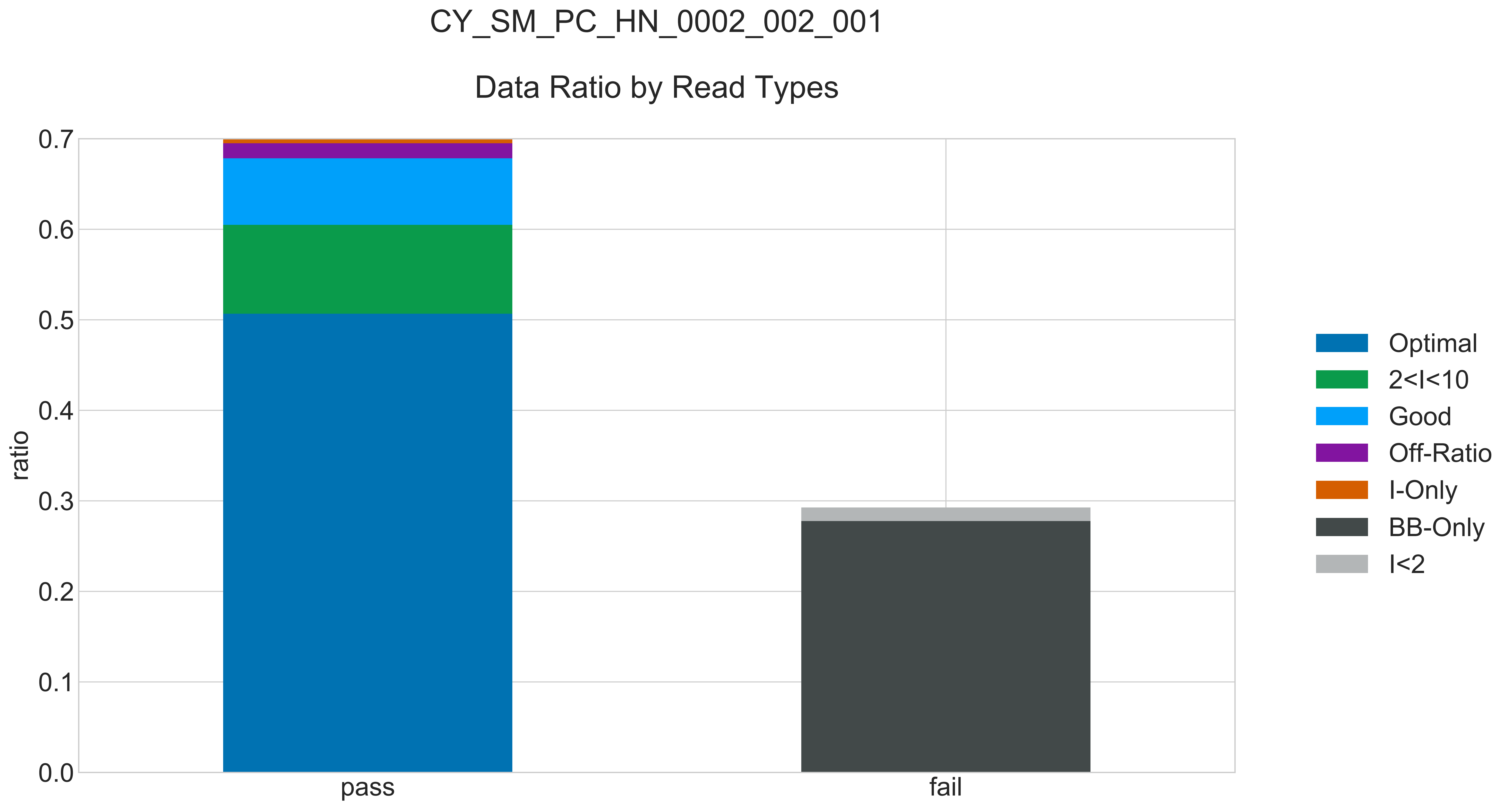

### CY_SM_PC_HN_0002_002_001_READ_LEN_.pdf

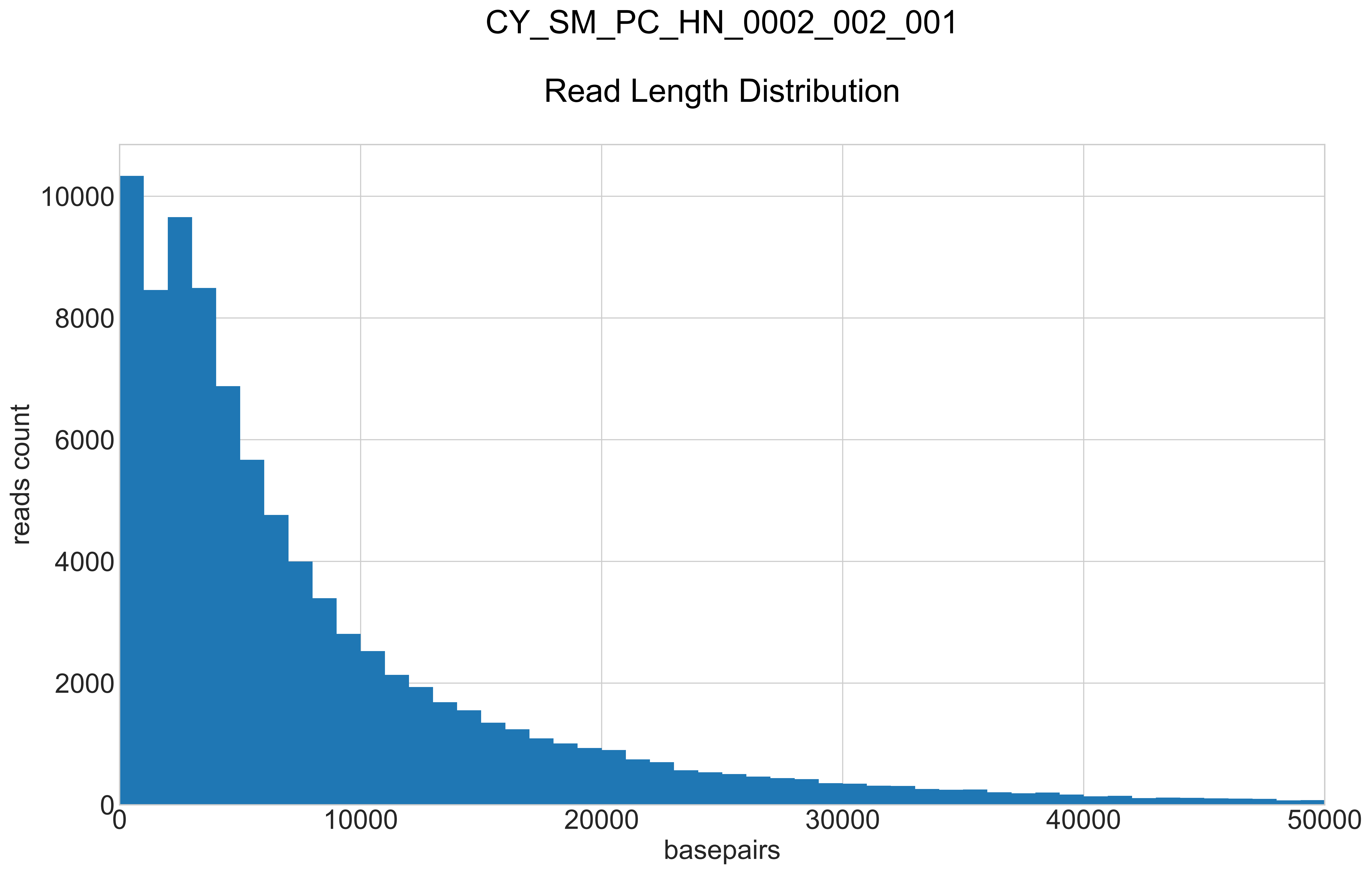

### CY_SM_PC_HN_0002_002_001_REPEATS_COUNT_.pdf

# CY_SM_PC_HN_0002_002_001

##### Repeats Distribution

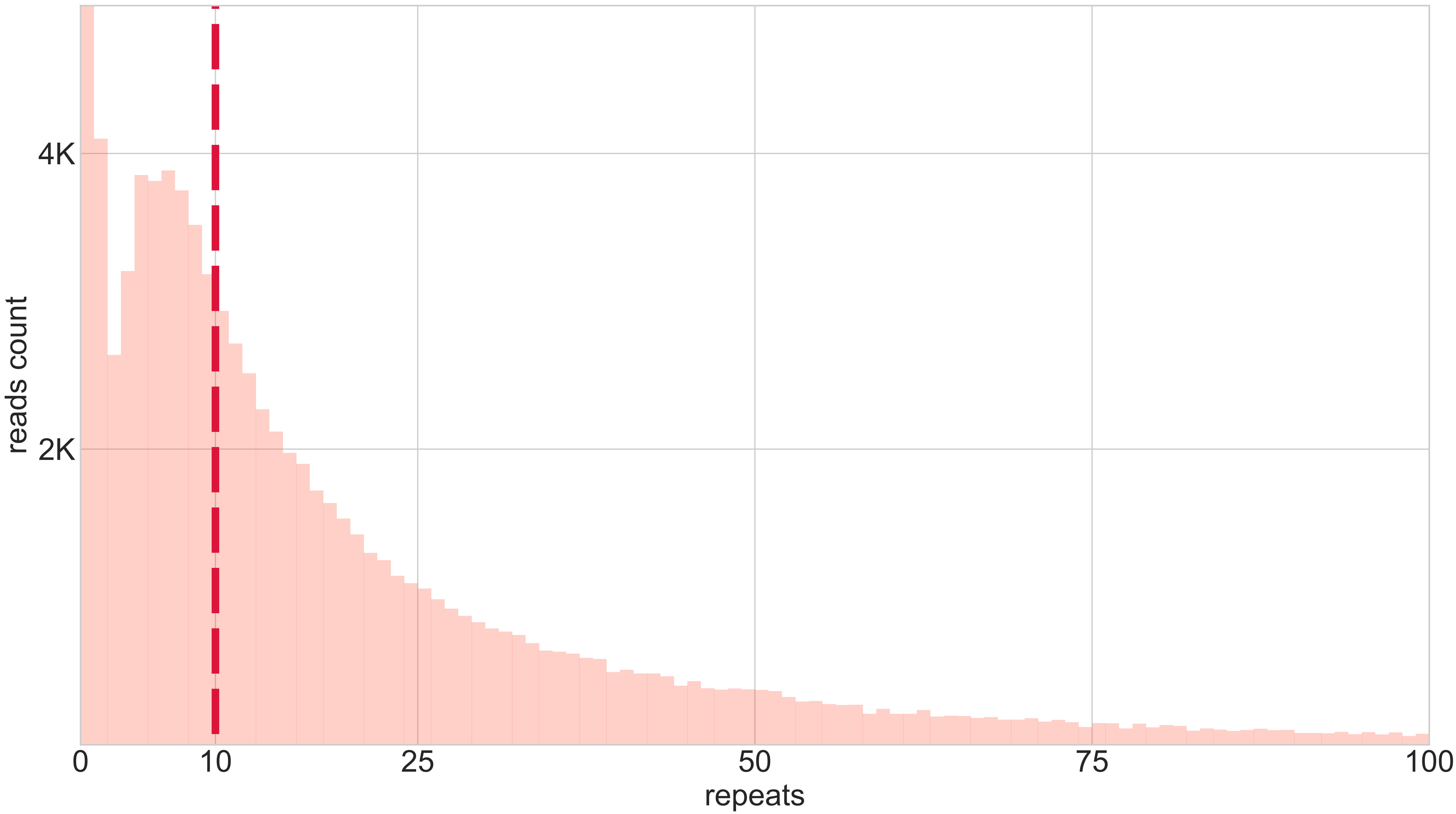

### CY_SM_PC_HN_0002_003_000_BB-I_.pdf

# CY_SM_PC_HN_0002_003_000

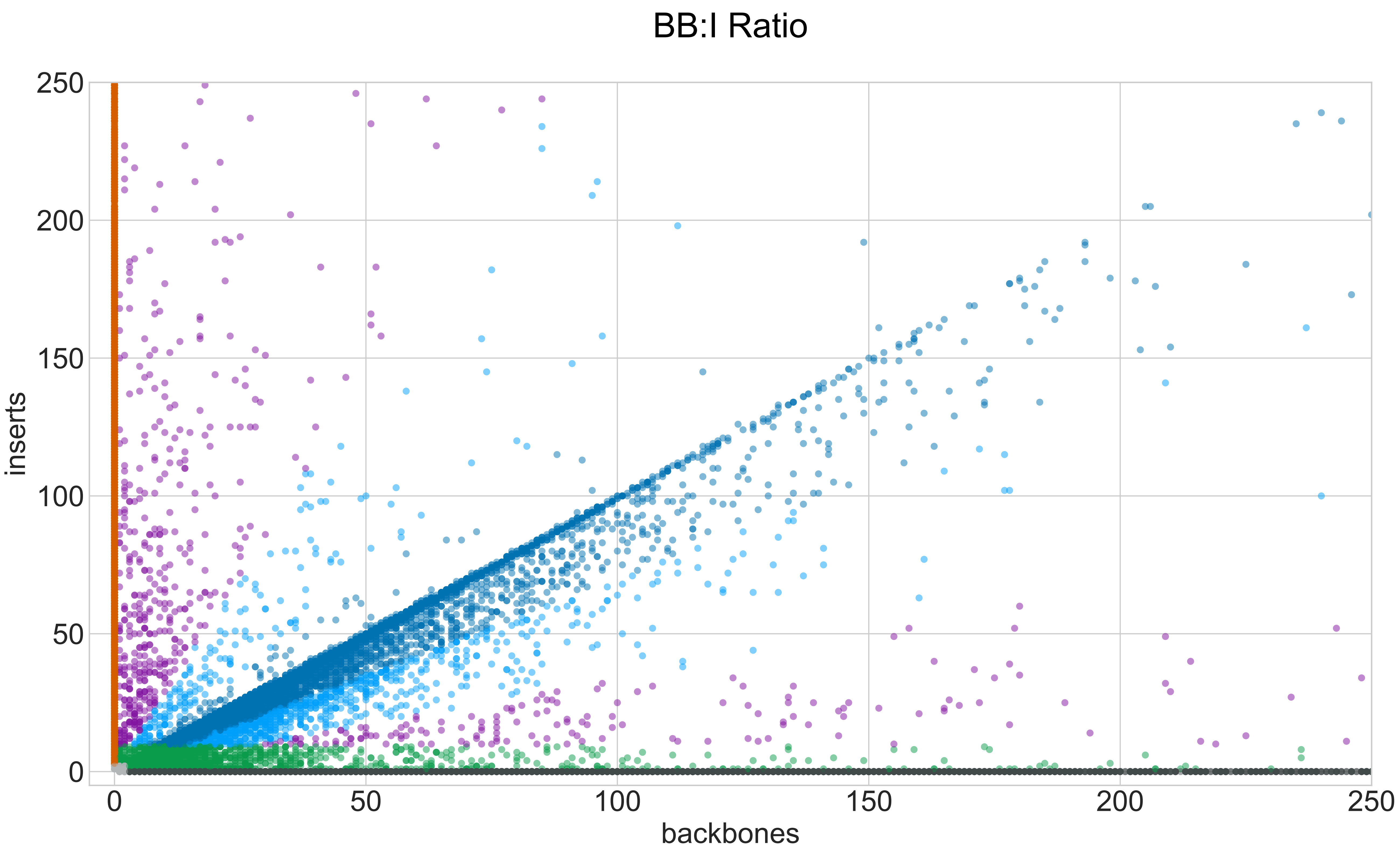

### CY_SM_PC_HN_0002_003_000_GROUPS_STACKED_NORM-LEN_.pdf

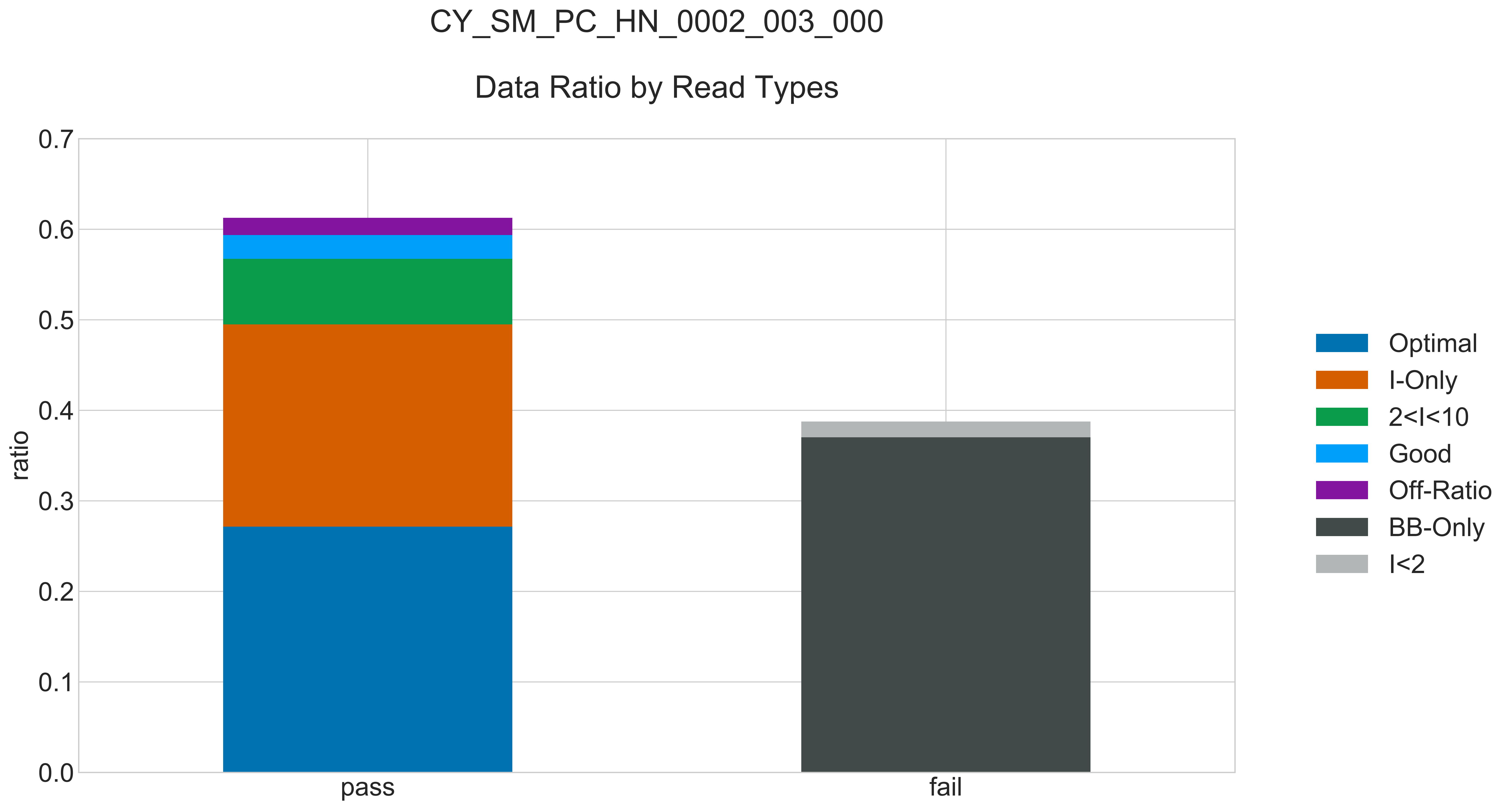

### CY_SM_PC_HN_0002_003_000_READ_LEN_.pdf

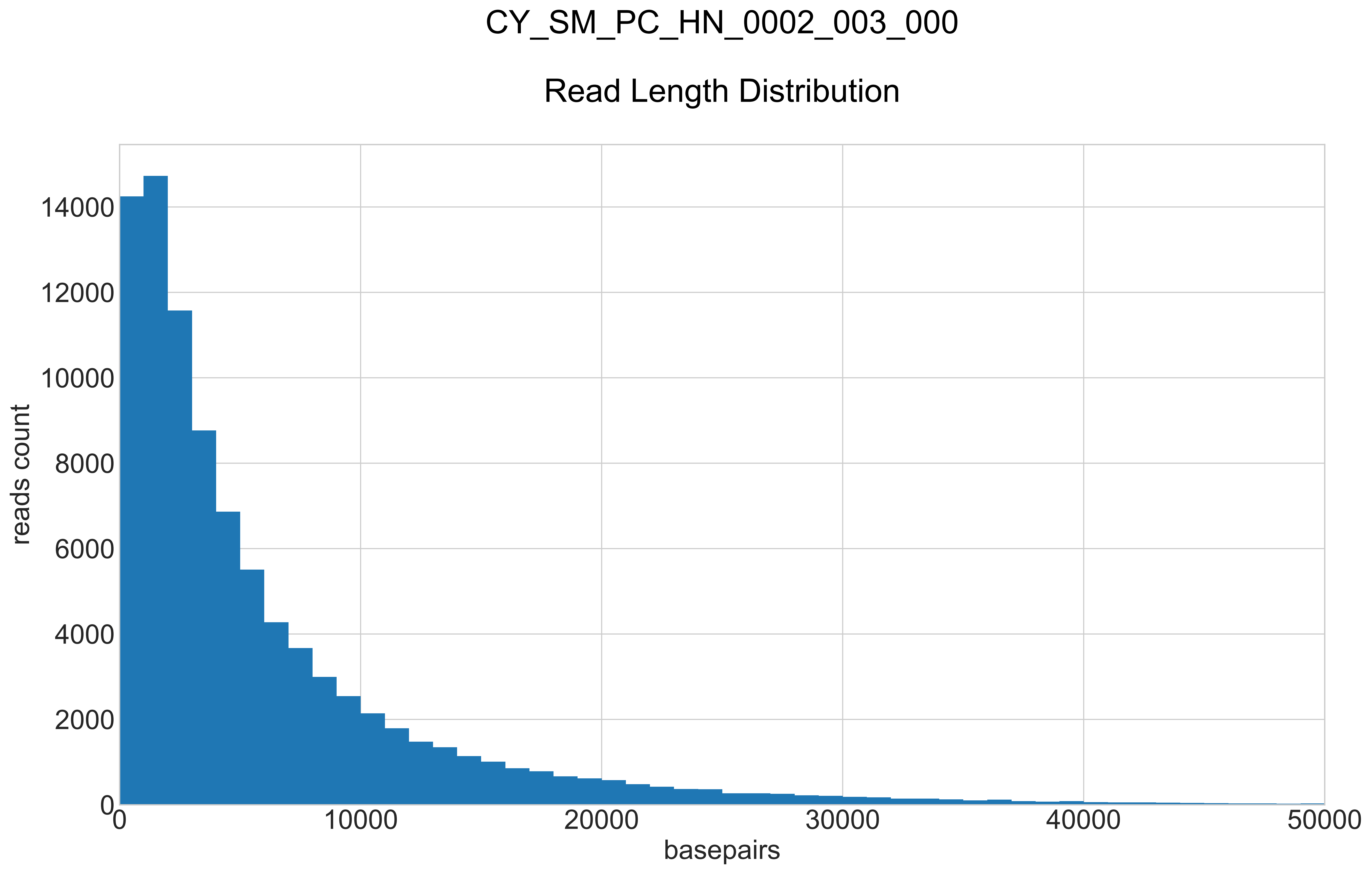

### CY_SM_PC_HN_0002_003_000_REPEATS_COUNT_.pdf

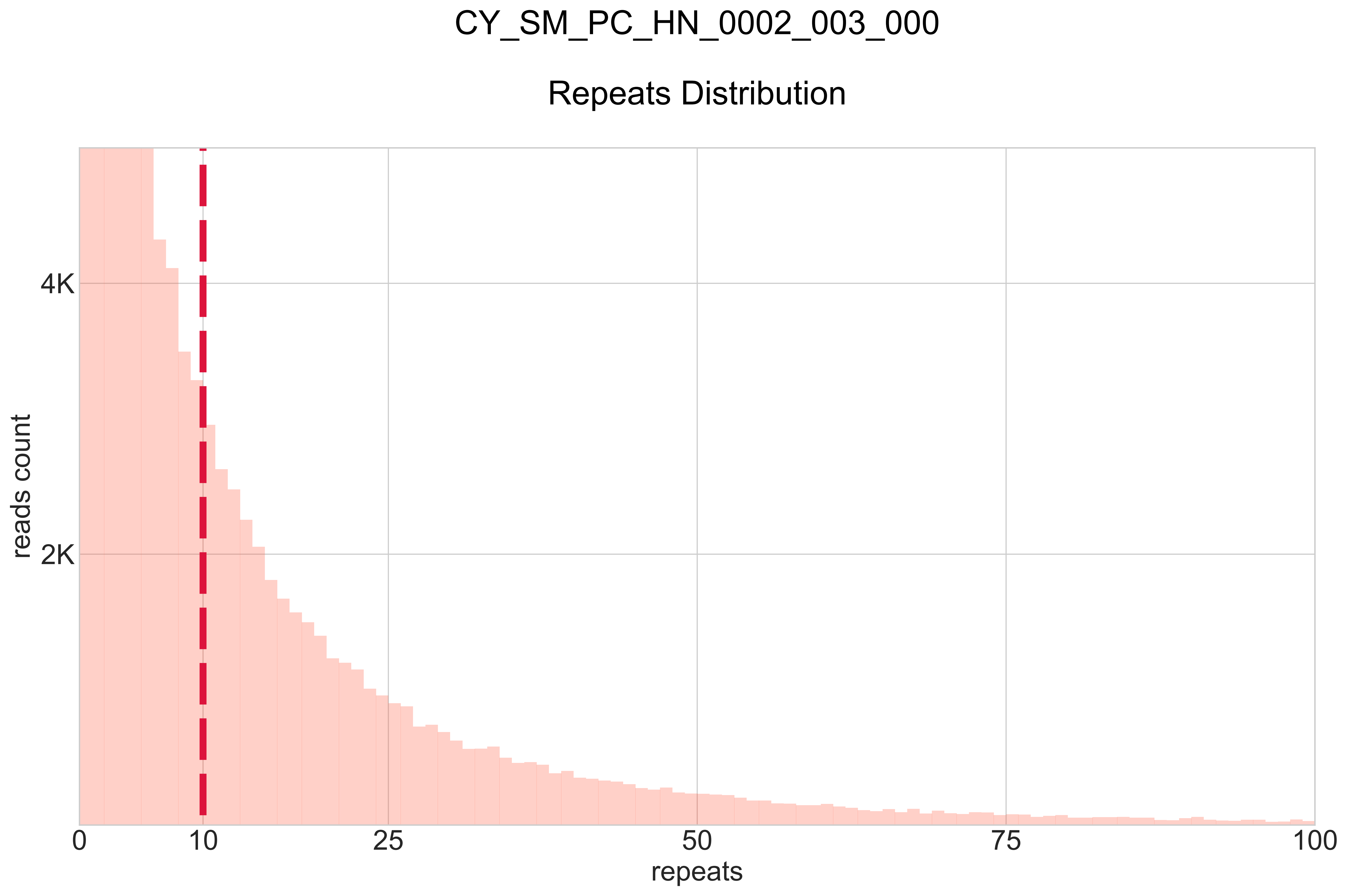

### CY_SS_PC_HC_0001_001_000_BB-I_.pdf

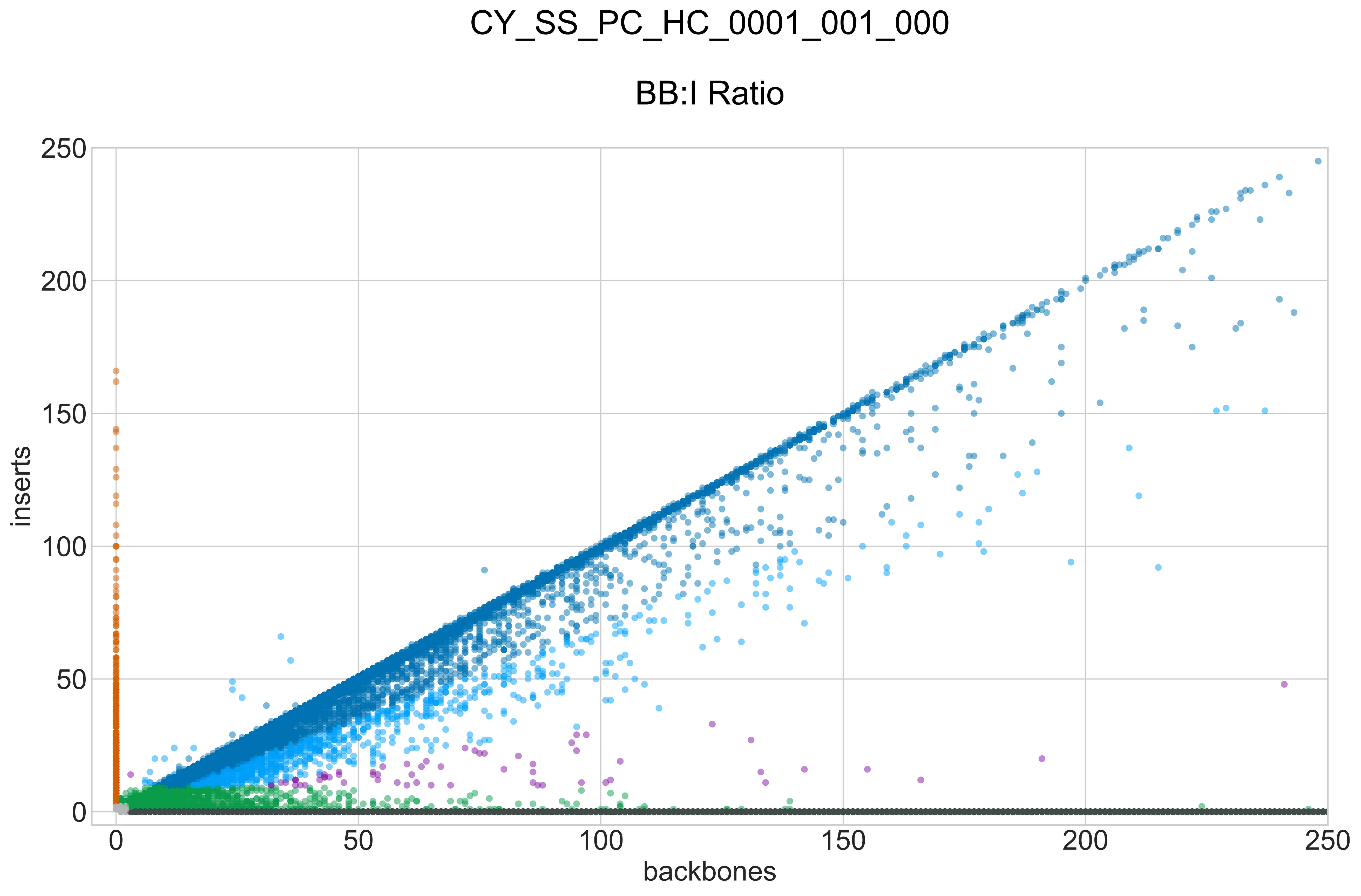

### CY_SS_PC_HC_0001_001_000_GROUPS_STACKED_NORM-LEN_.pdf

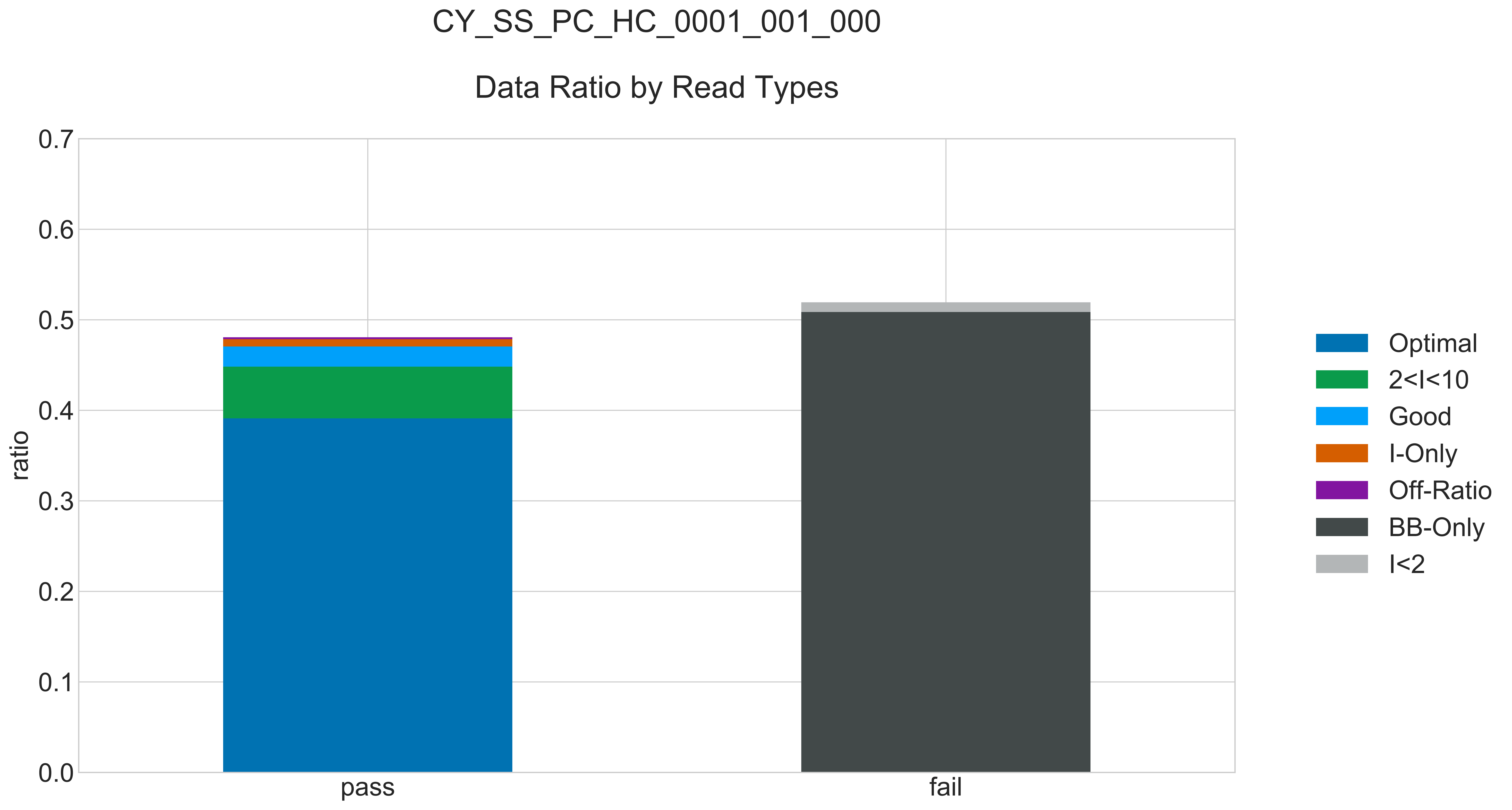

### CY_SS_PC_HC_0001_001_000_READ_LEN_.pdf

# CY_SS_PC_HC_0001_001_000

##### Read Length Distribution

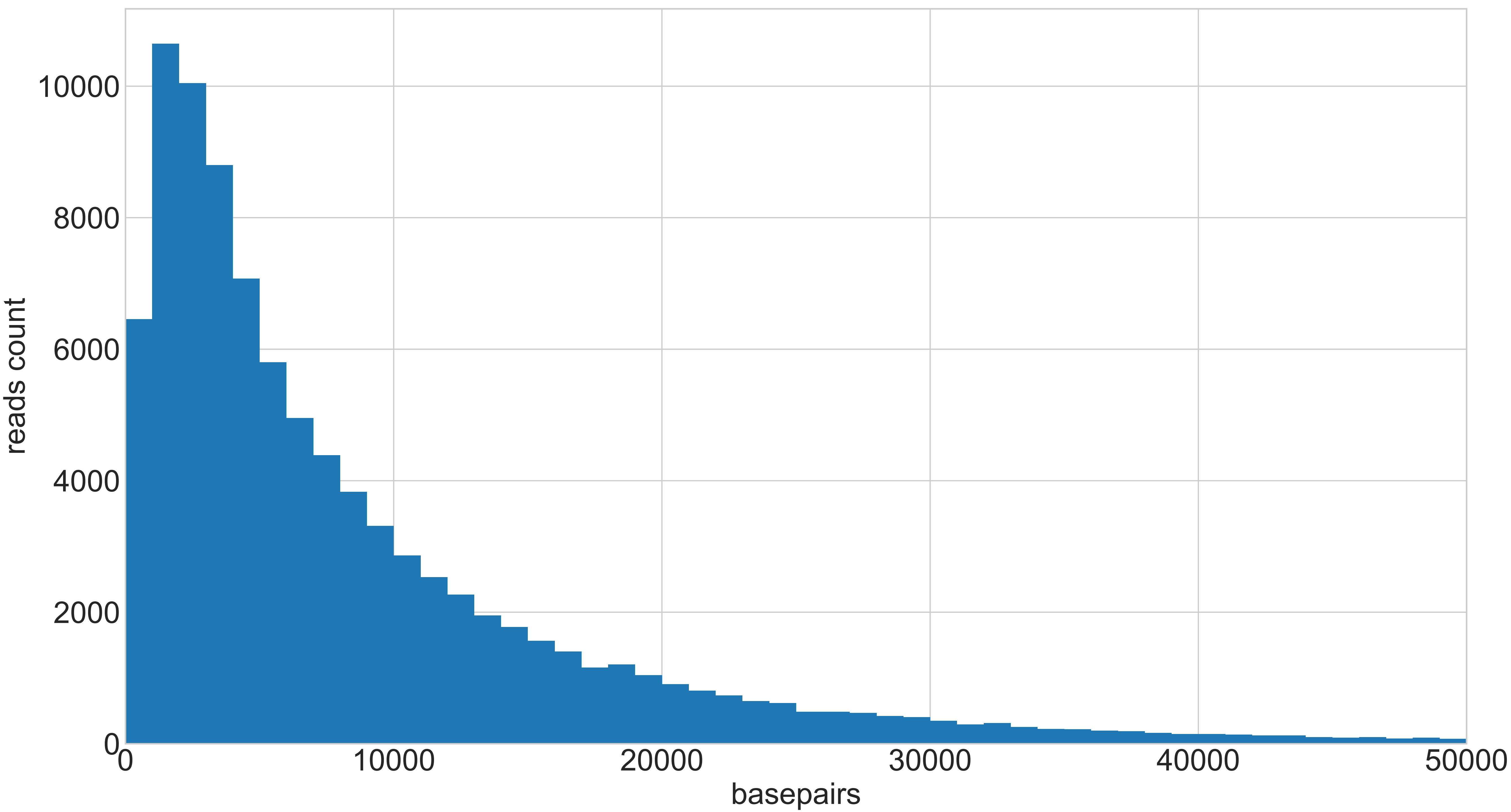

### CY_SS_PC_HC_0001_001_000_REPEATS_COUNT_.pdf

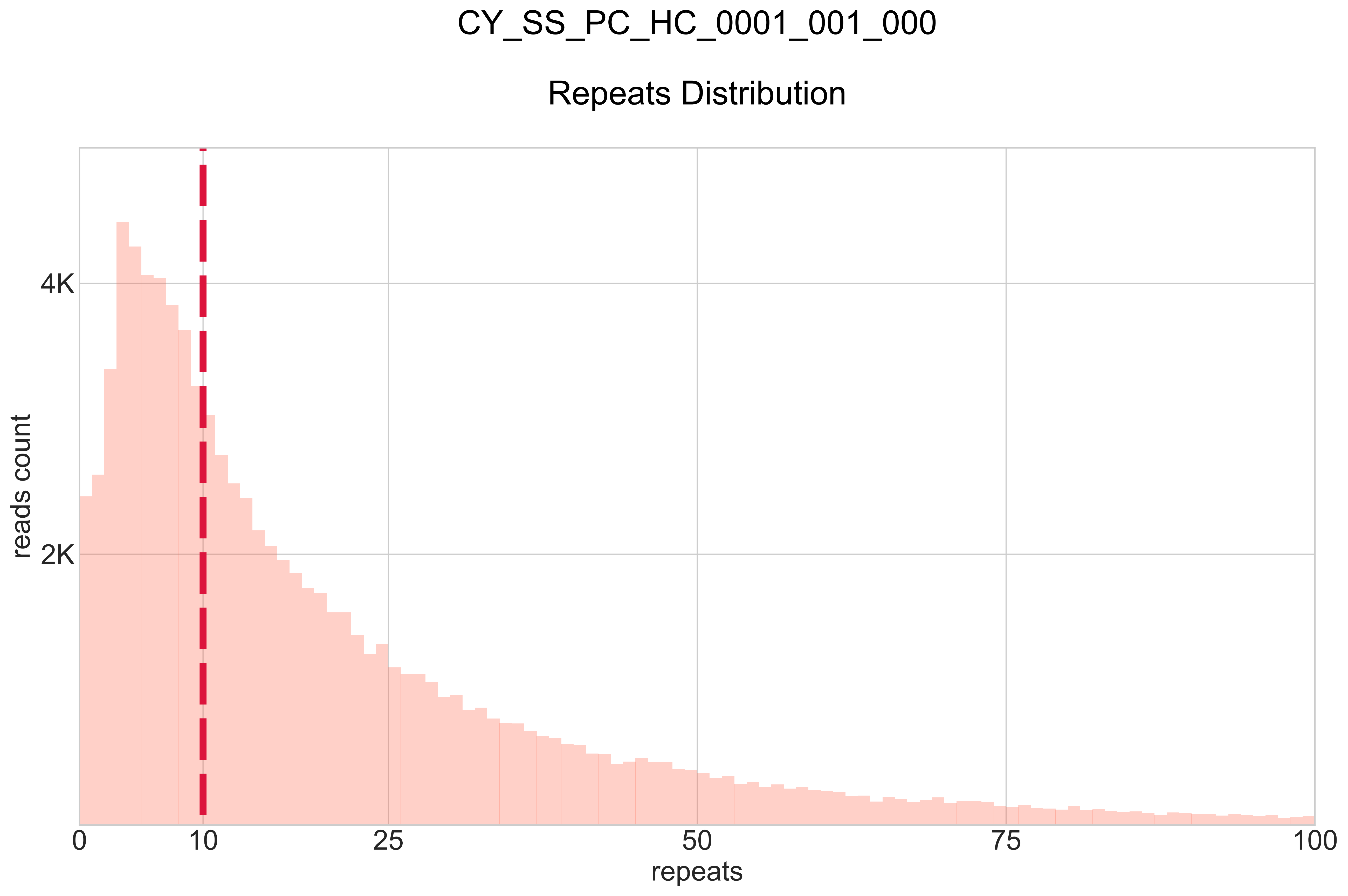

### CY_SS_PC_HC_0005_002_000_BB-I_.pdf

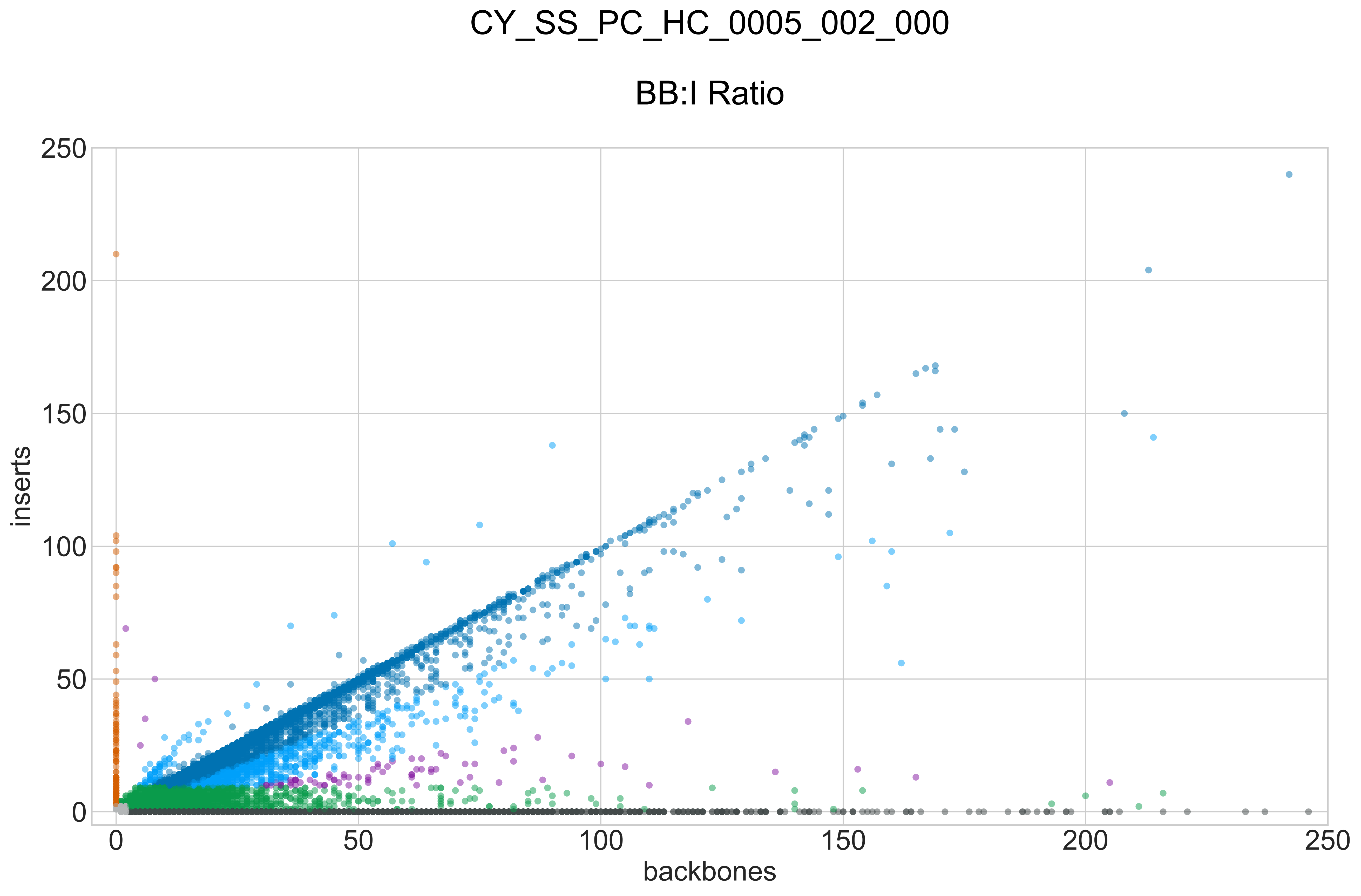

### CY_SS_PC_HC_0005_002_000_GROUPS_STACKED_NORM-LEN_.pdf

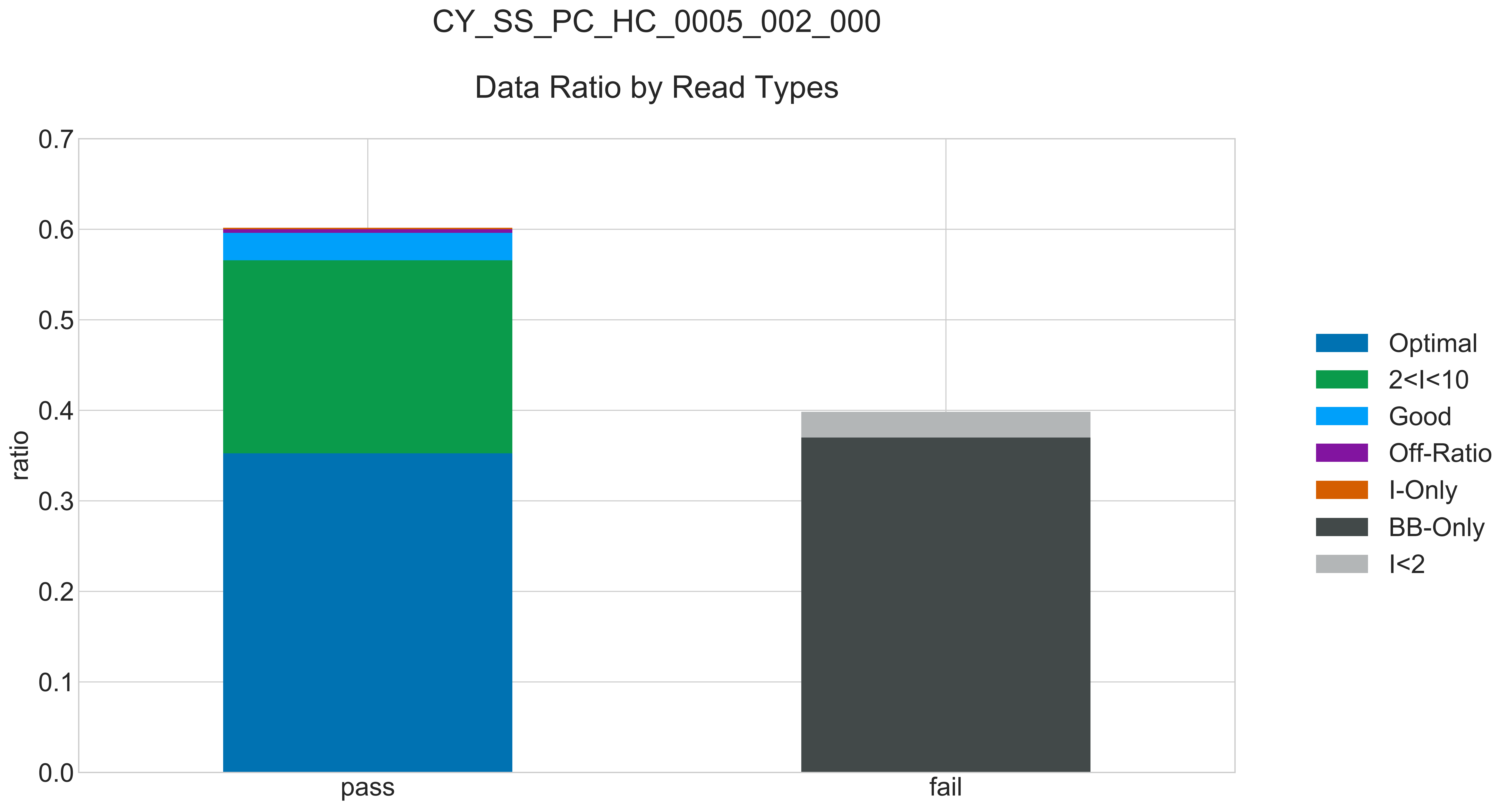

### CY_SS_PC_HC_0005_002_000_READ_LEN_.pdf

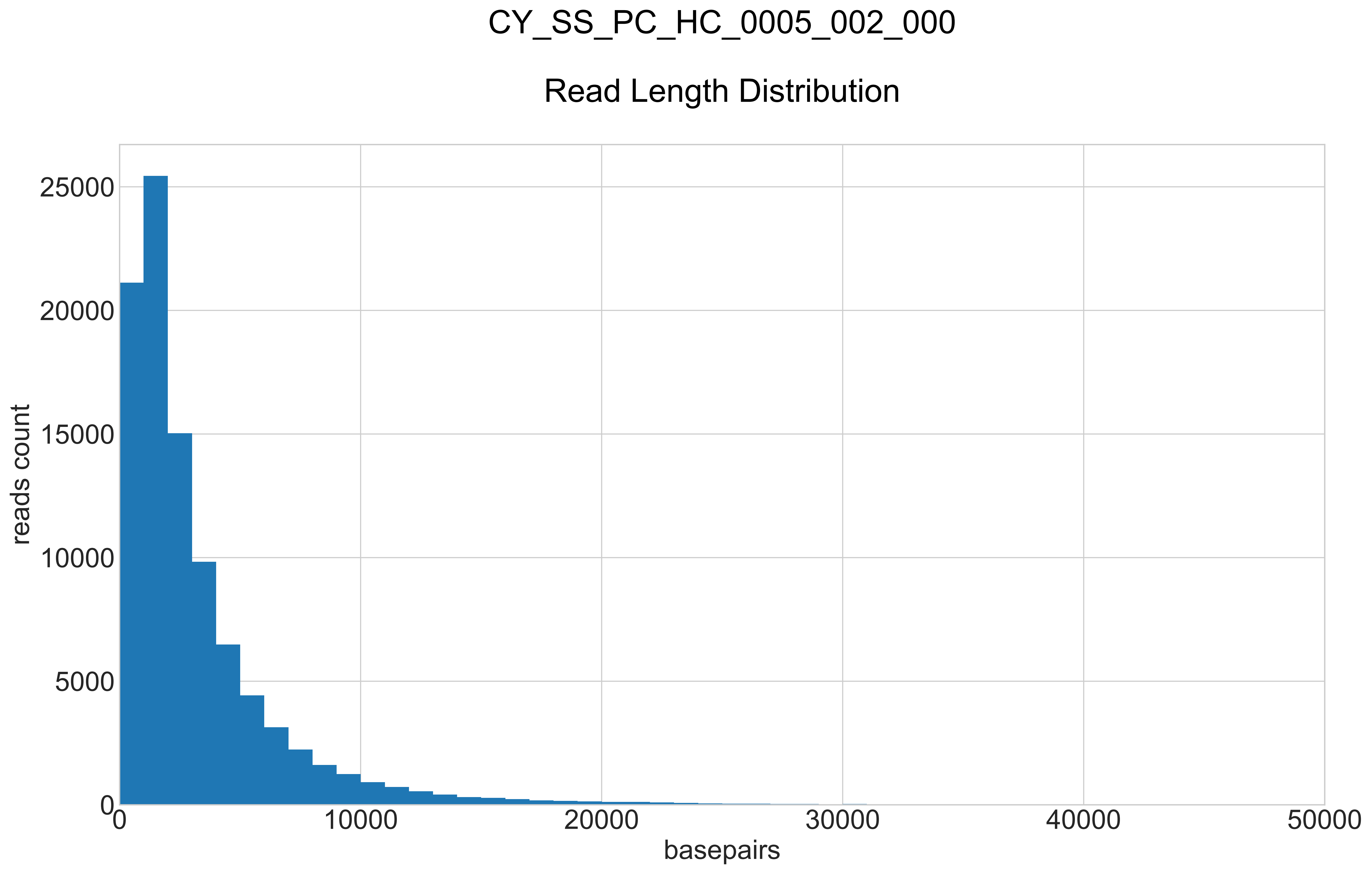

### CY_SS_PC_HC_0005_002_000_REPEATS_COUNT_.pdf

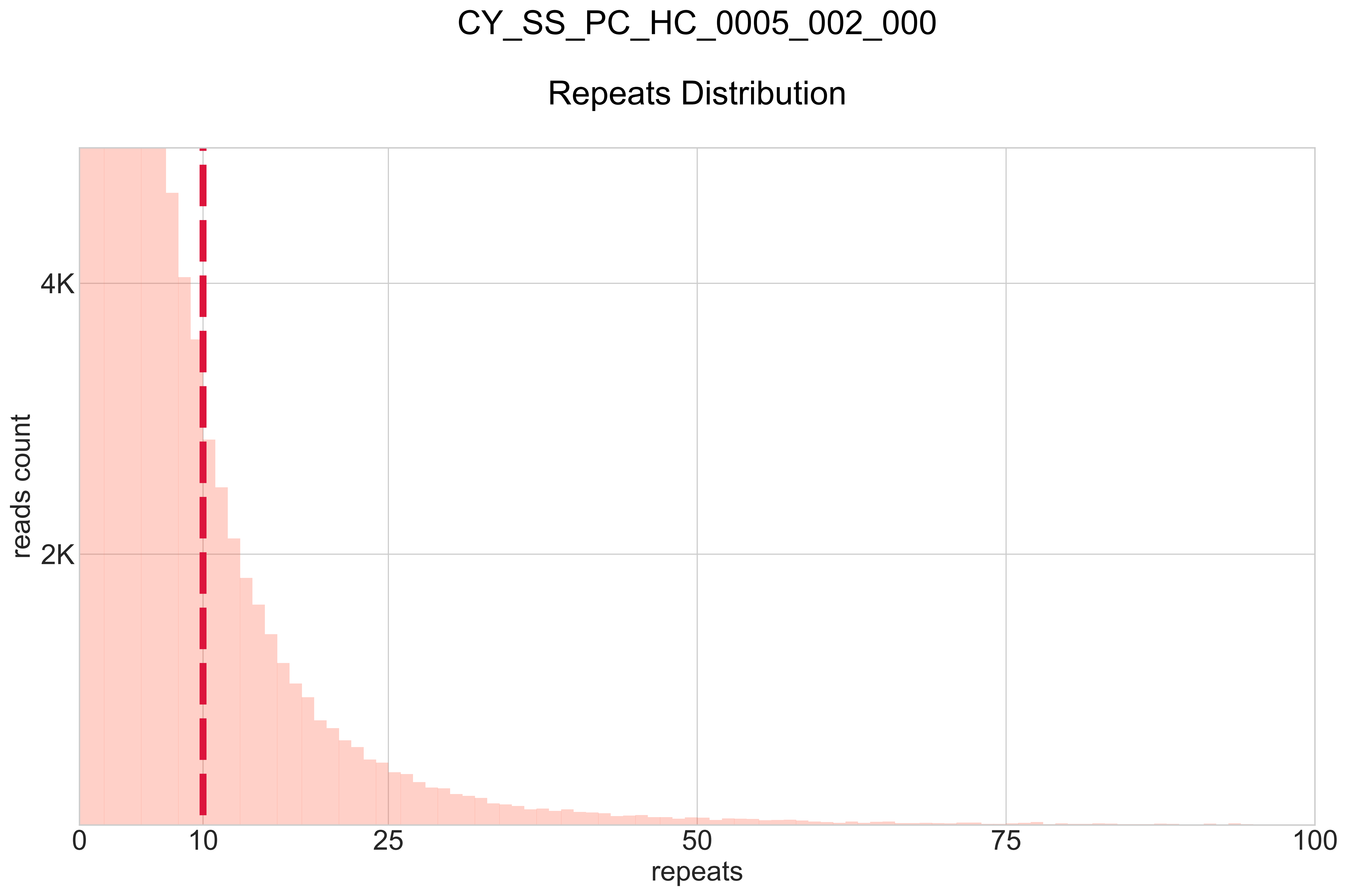

### CY_SS_PC_HN_0001_001_000_BB-I_.pdf

# CY_SS_PC_HN_0001_001_000

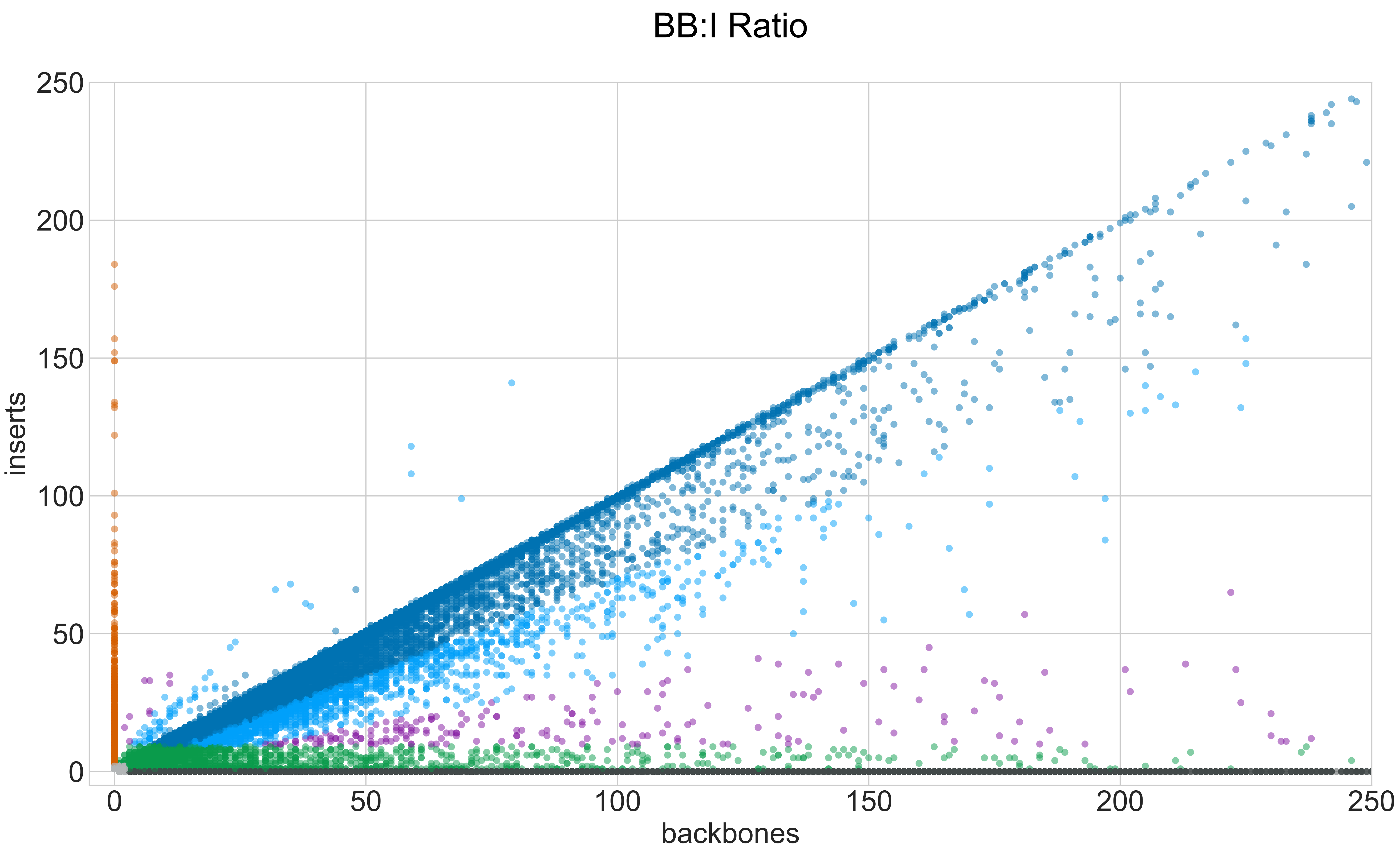

### CY_SS_PC_HN_0001_001_000_GROUPS_STACKED_NORM-LEN_.pdf

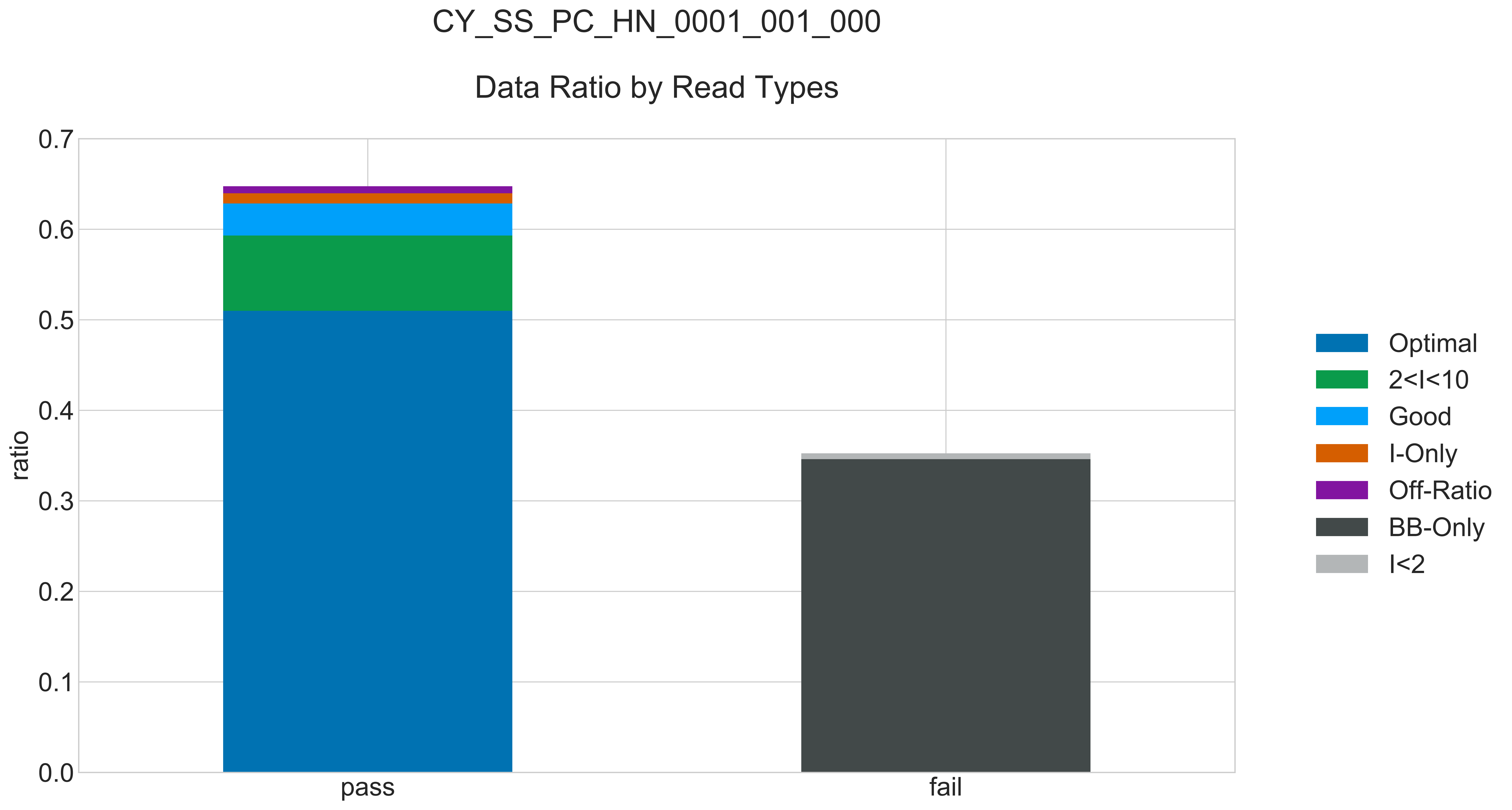

### CY_SS_PC_HN_0001_001_000_READ_LEN_.pdf

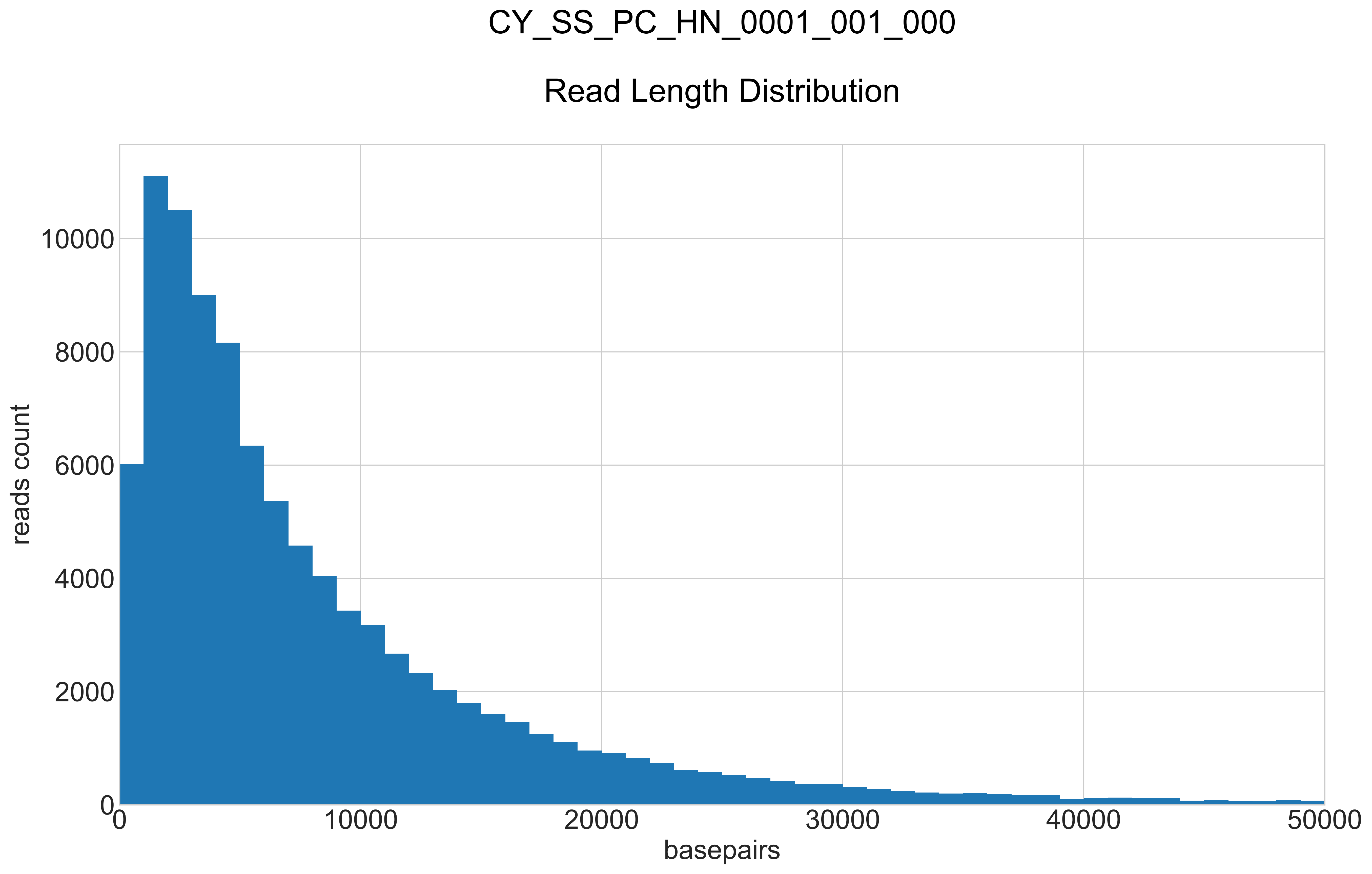

### CY_SS_PC_HN_0001_001_000_REPEATS_COUNT_.pdf

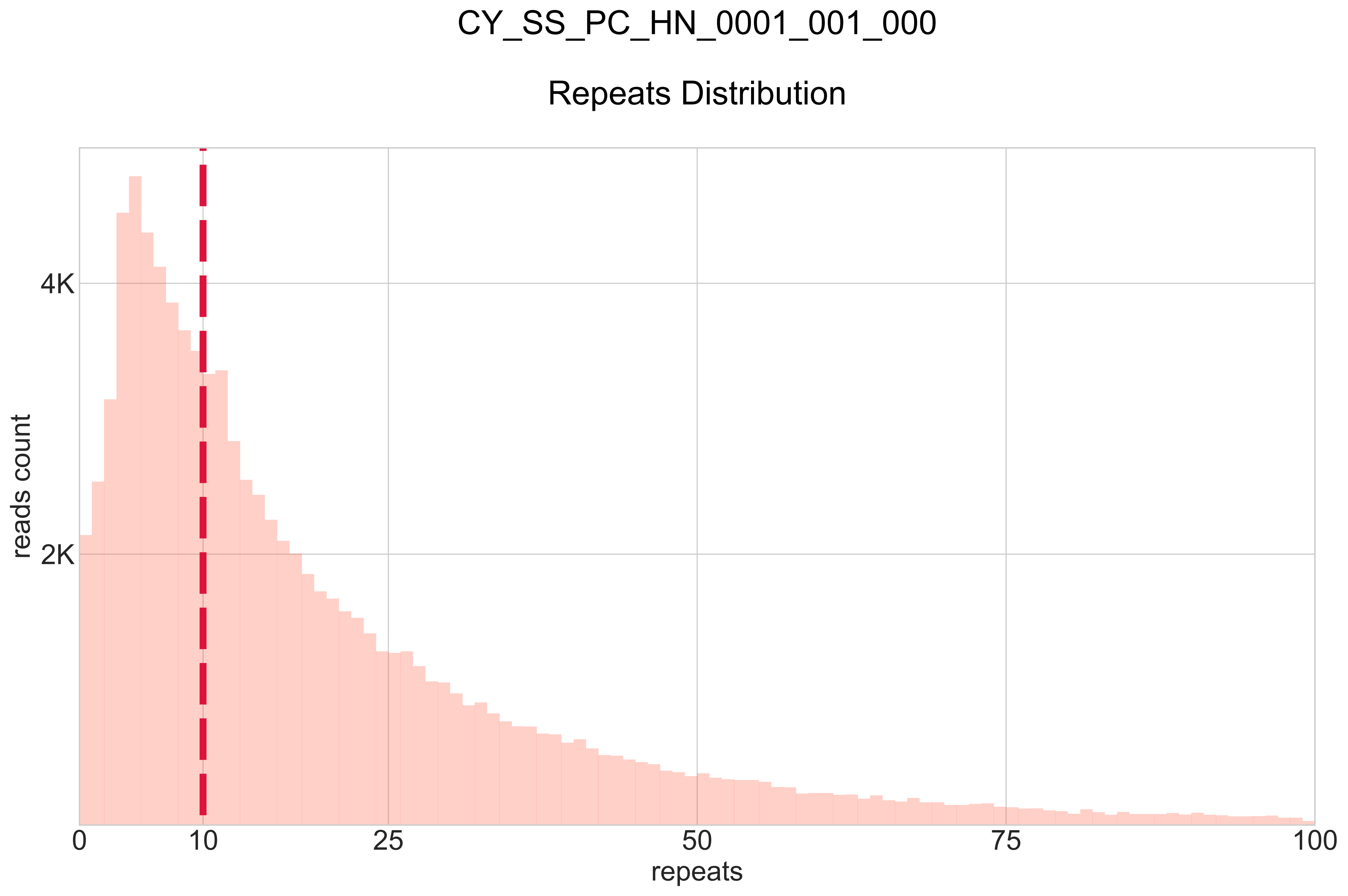

### CY_SS_PC_HN_0001_002_000_BB-I_.pdf

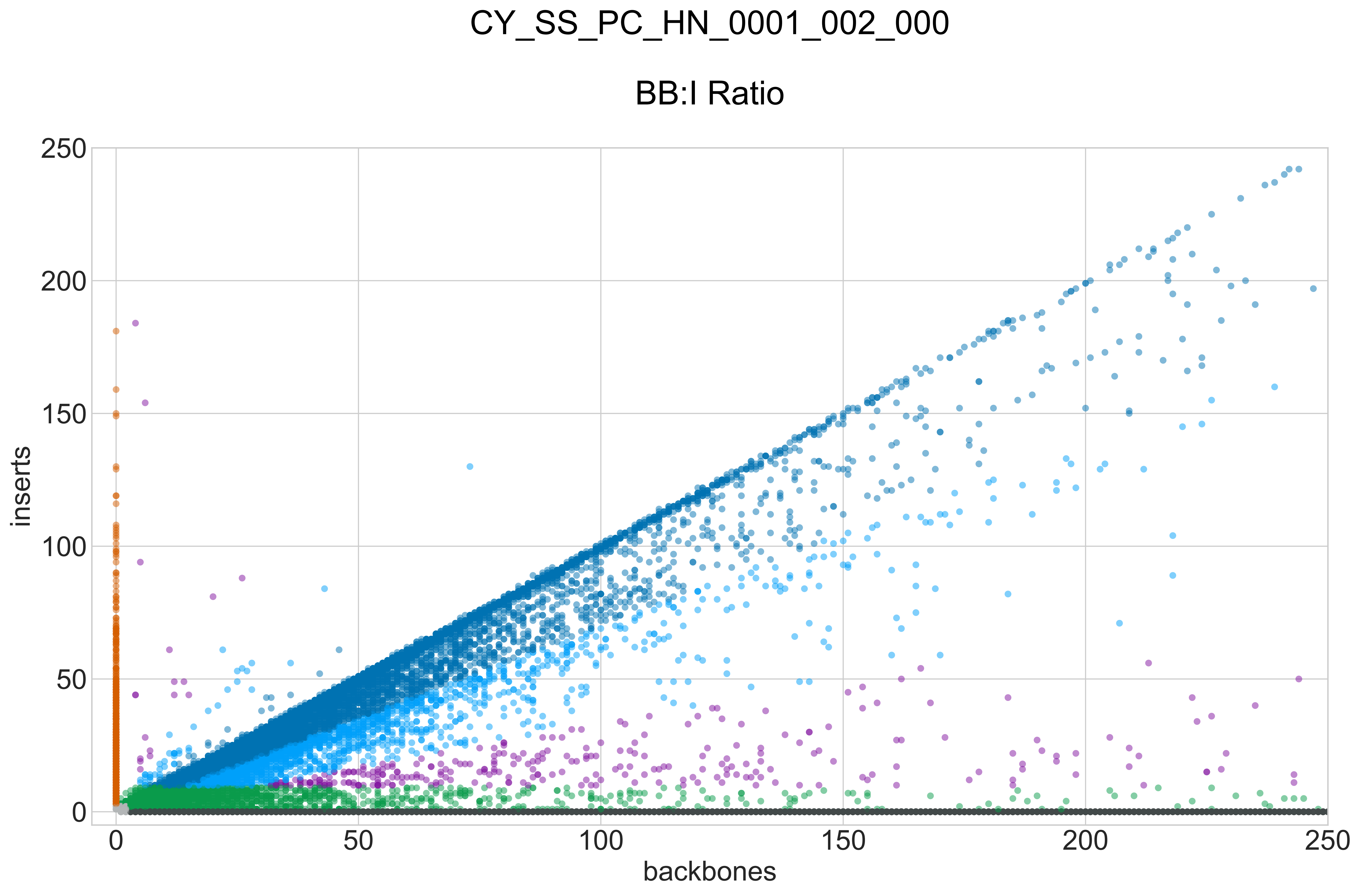

### CY_SS_PC_HN_0001_002_000_GROUPS_STACKED_NORM-LEN_.pdf

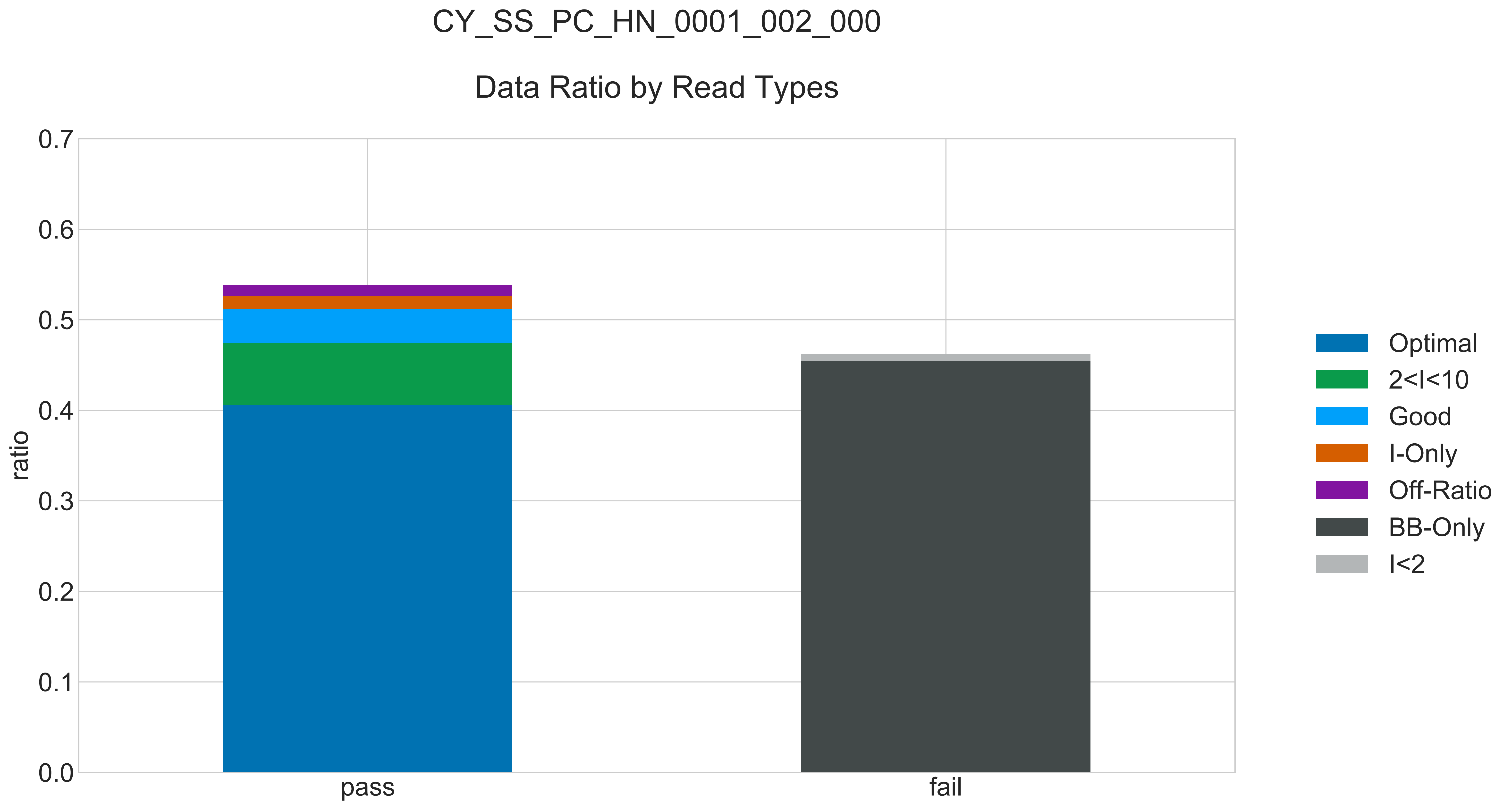

### CY_SS_PC_HN_0001_002_000_READ_LEN_.pdf

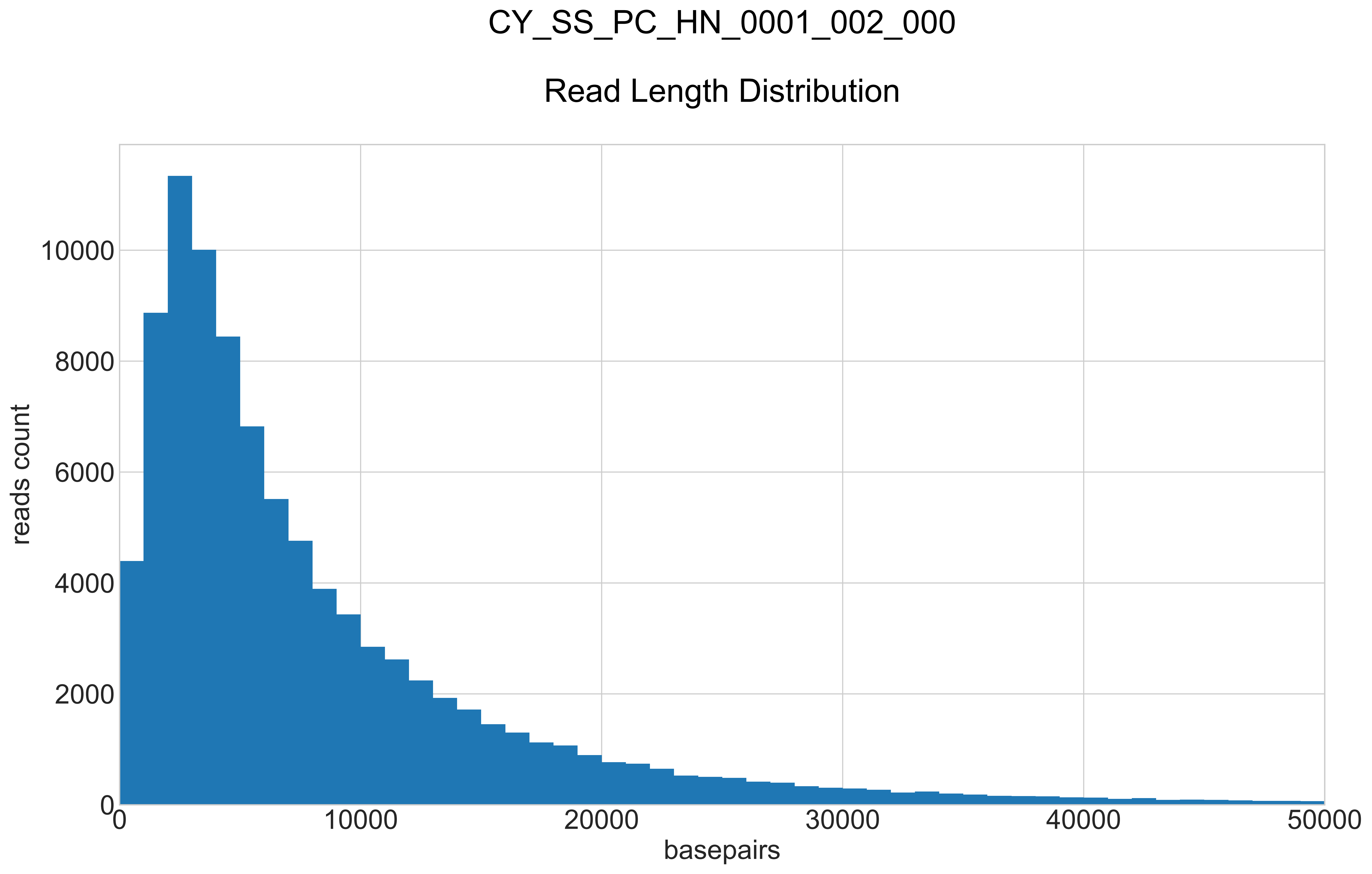

### CY_SS_PC_HN_0001_002_000_REPEATS_COUNT_.pdf

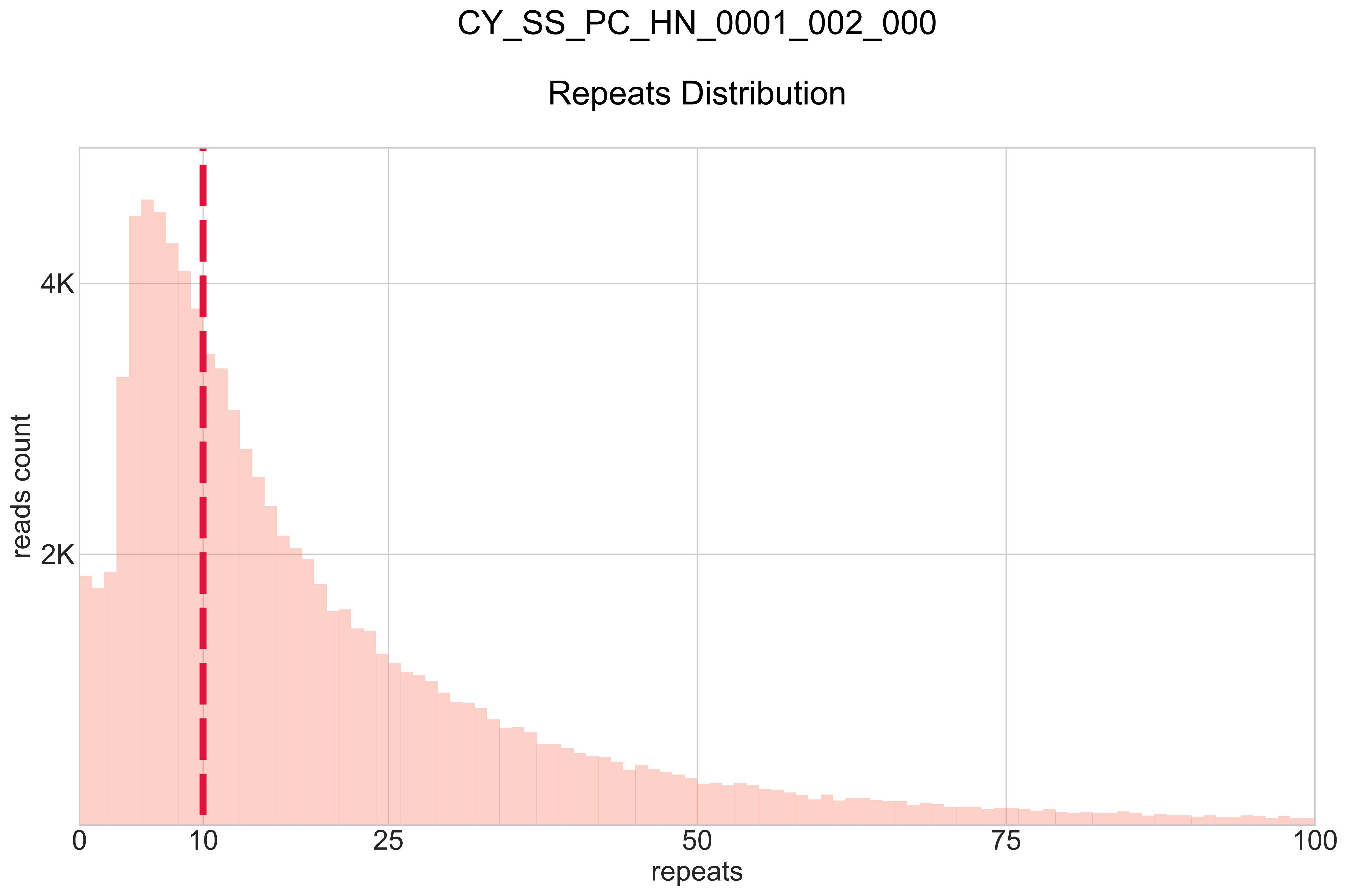

### CY_SS_PC_HN_0001_003_000_BB-I_.pdf

# CY_SS_PC_HN_0001_003_000

##### BB:I Ratio

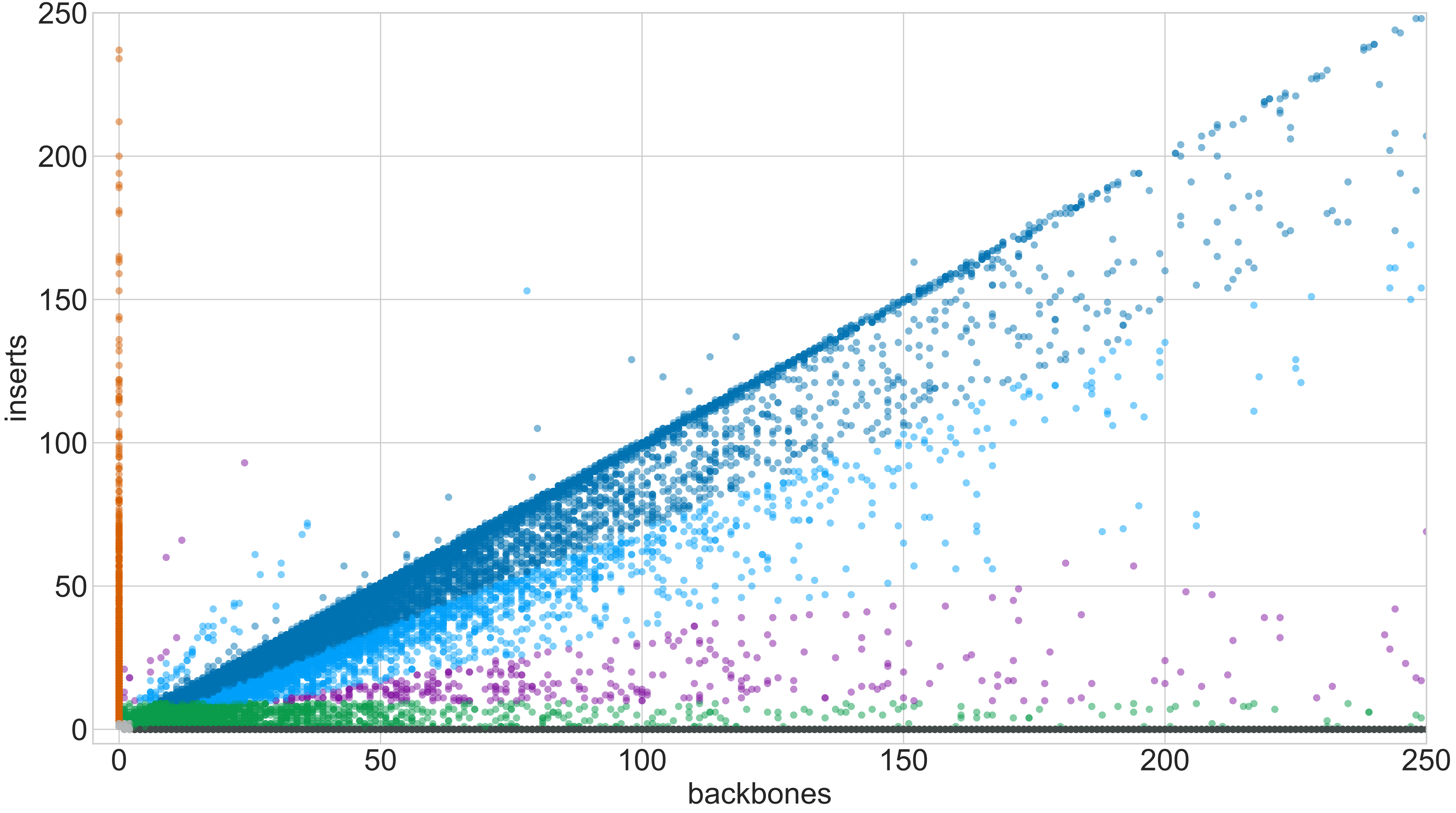

### CY_SS_PC_HN_0001_003_000_GROUPS_STACKED_NORM-LEN_.pdf

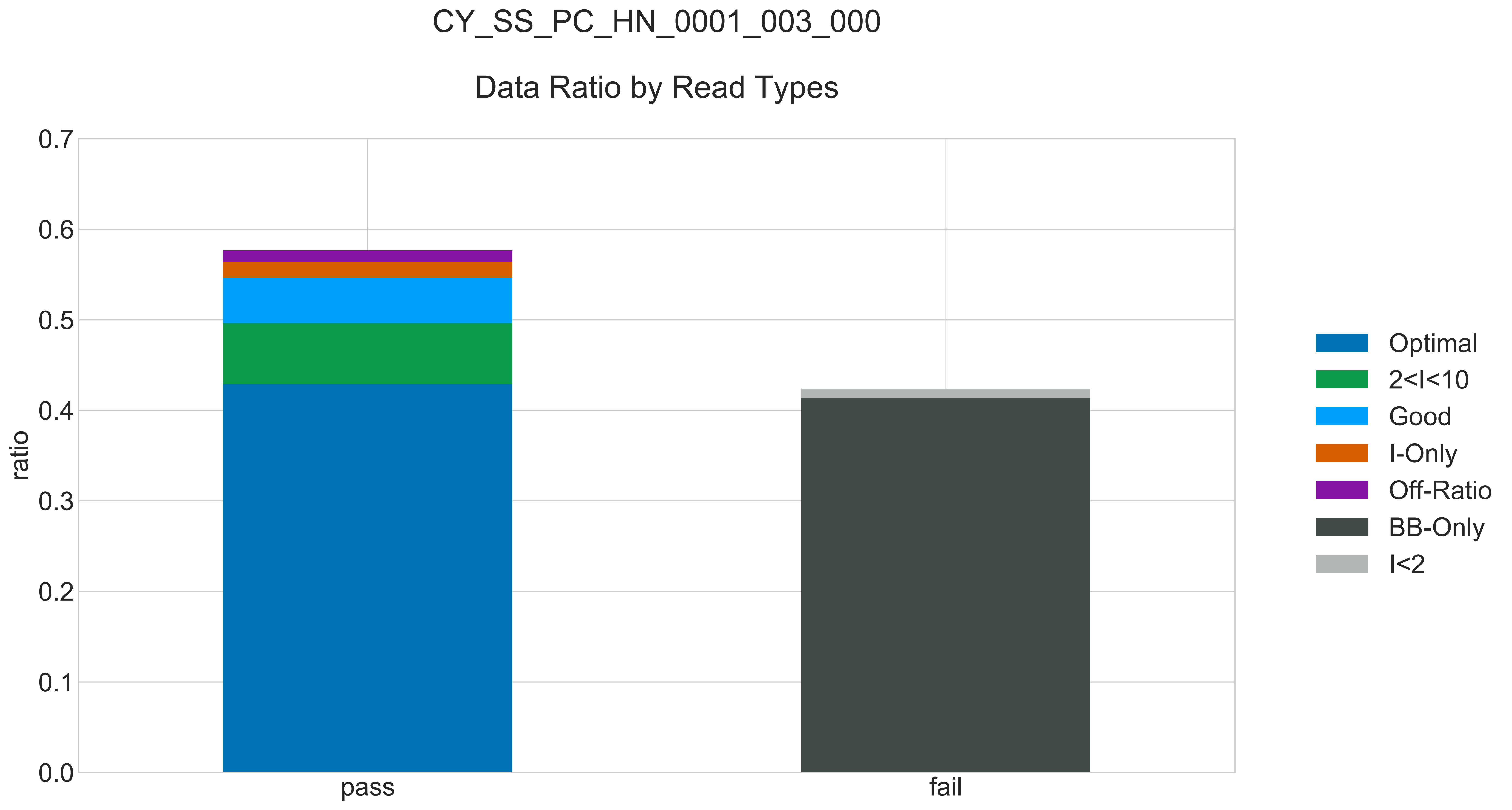

### CY_SS_PC_HN_0001_003_000_READ_LEN_.pdf

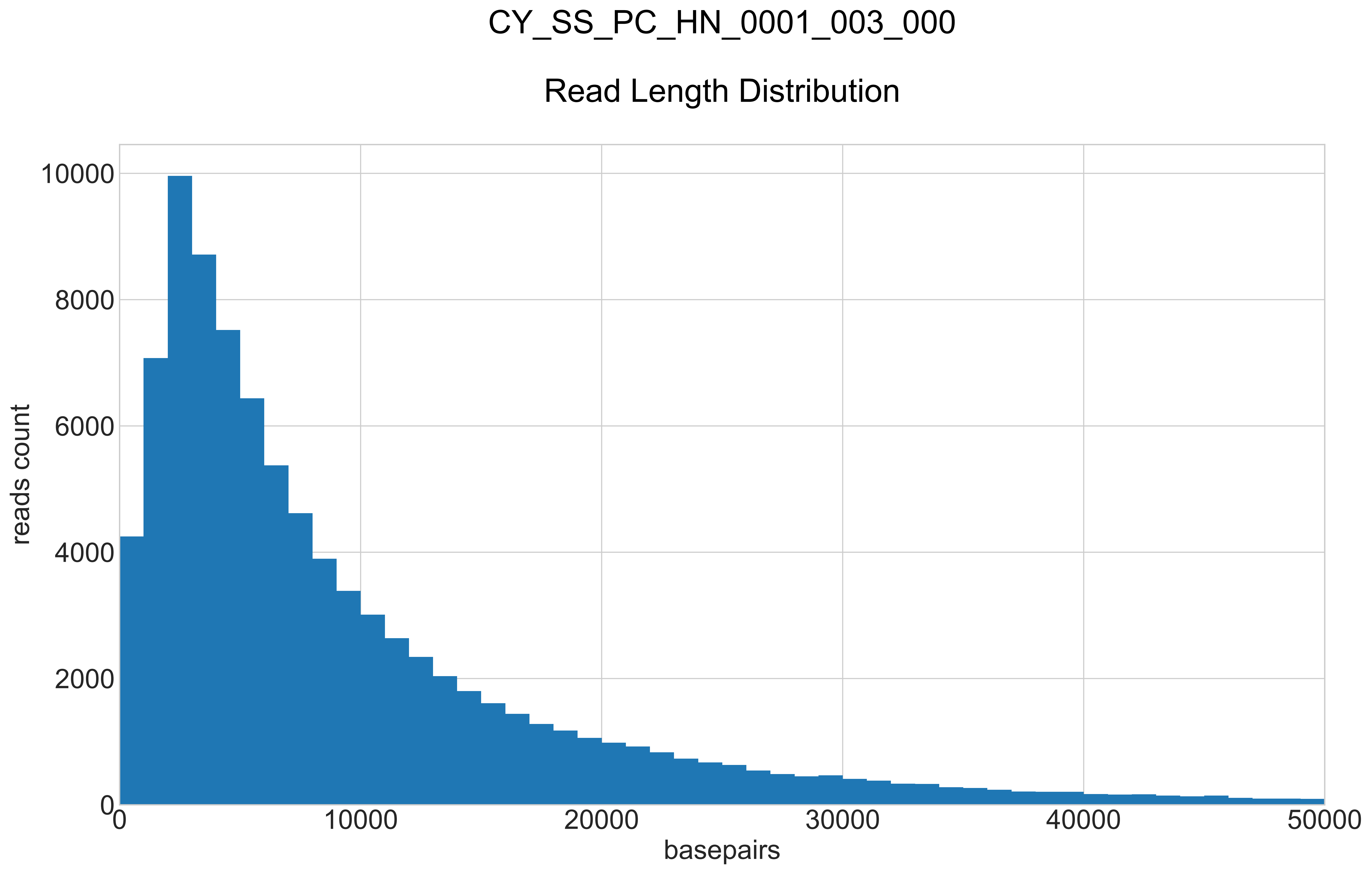

### CY_SS_PC_HN_0001_003_000_REPEATS_COUNT_.pdf

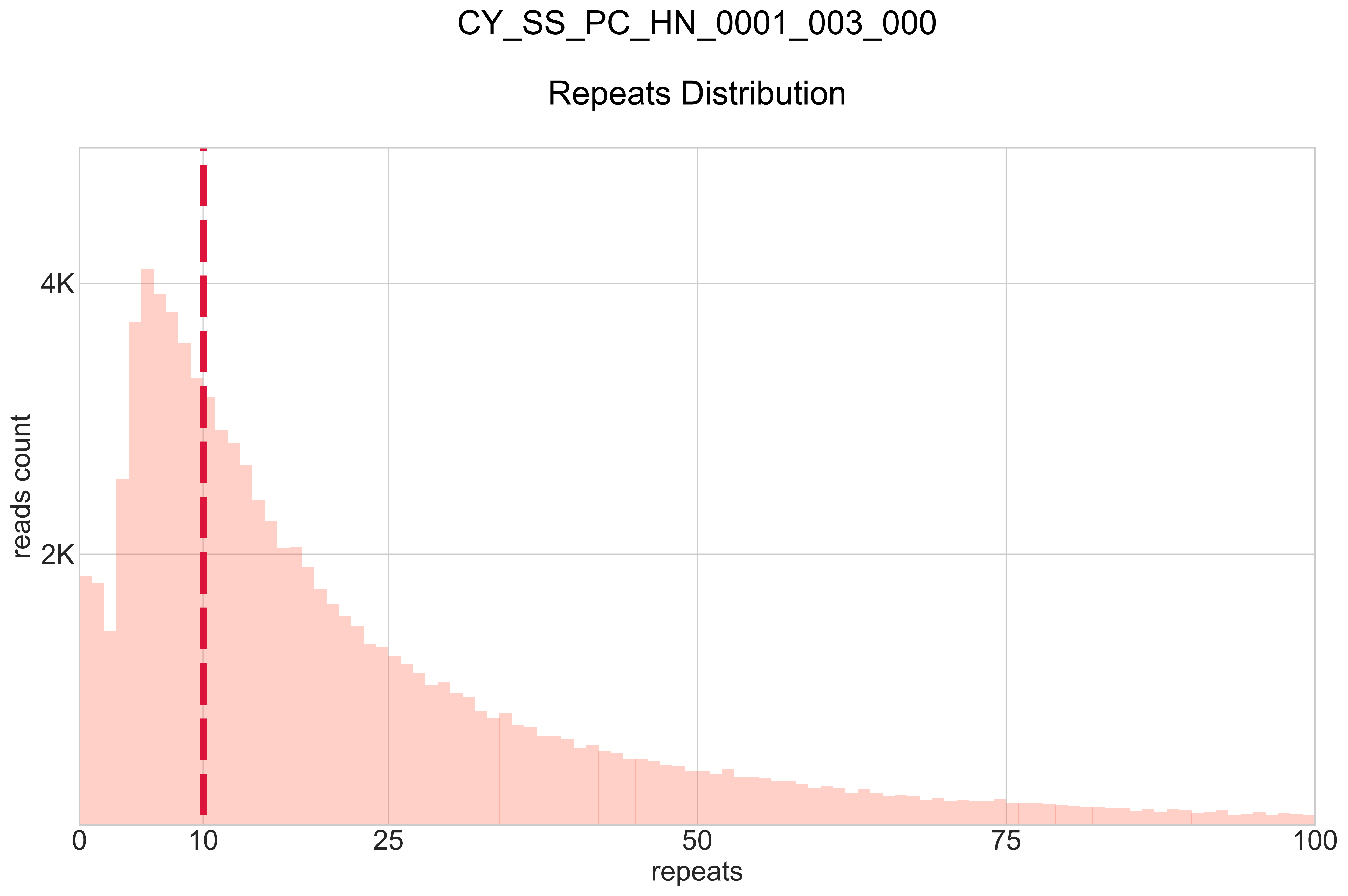

### CY_SS_PC_HN_0001_004_000_BB-I_.pdf

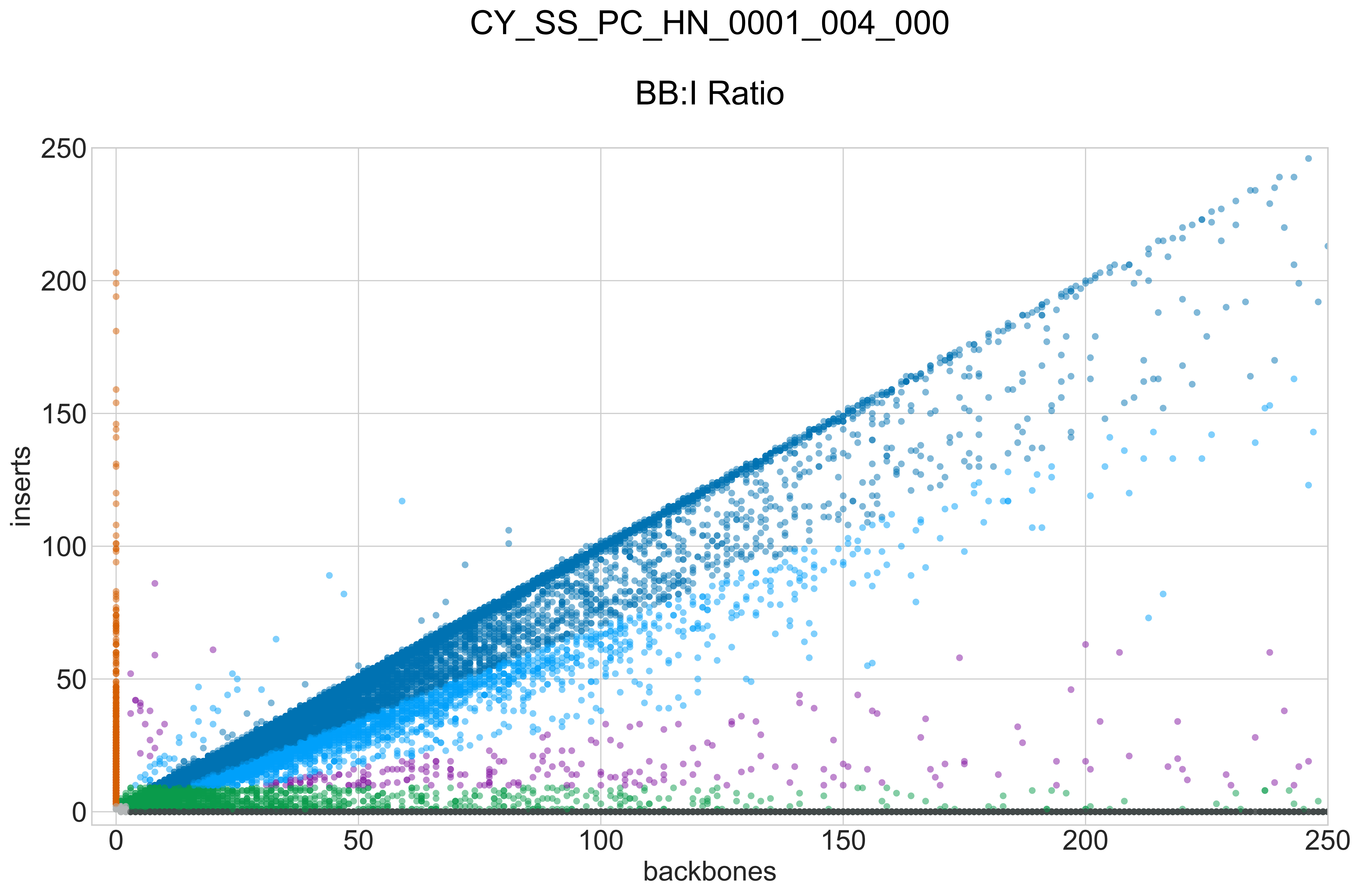

### CY_SS_PC_HN_0001_004_000_GROUPS_STACKED_NORM-LEN_.pdf

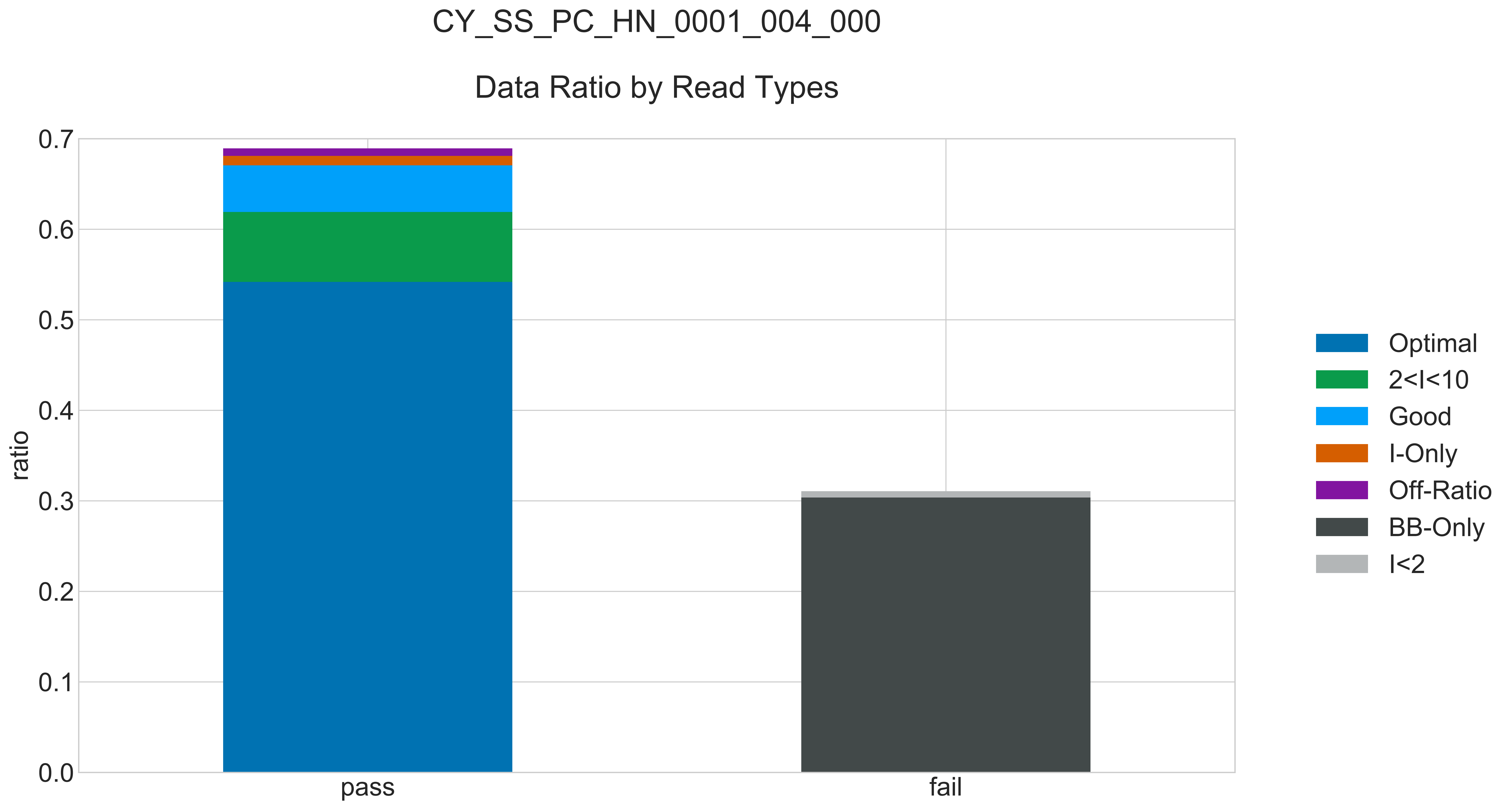

### CY_SS_PC_HN_0001_004_000_READ_LEN_.pdf

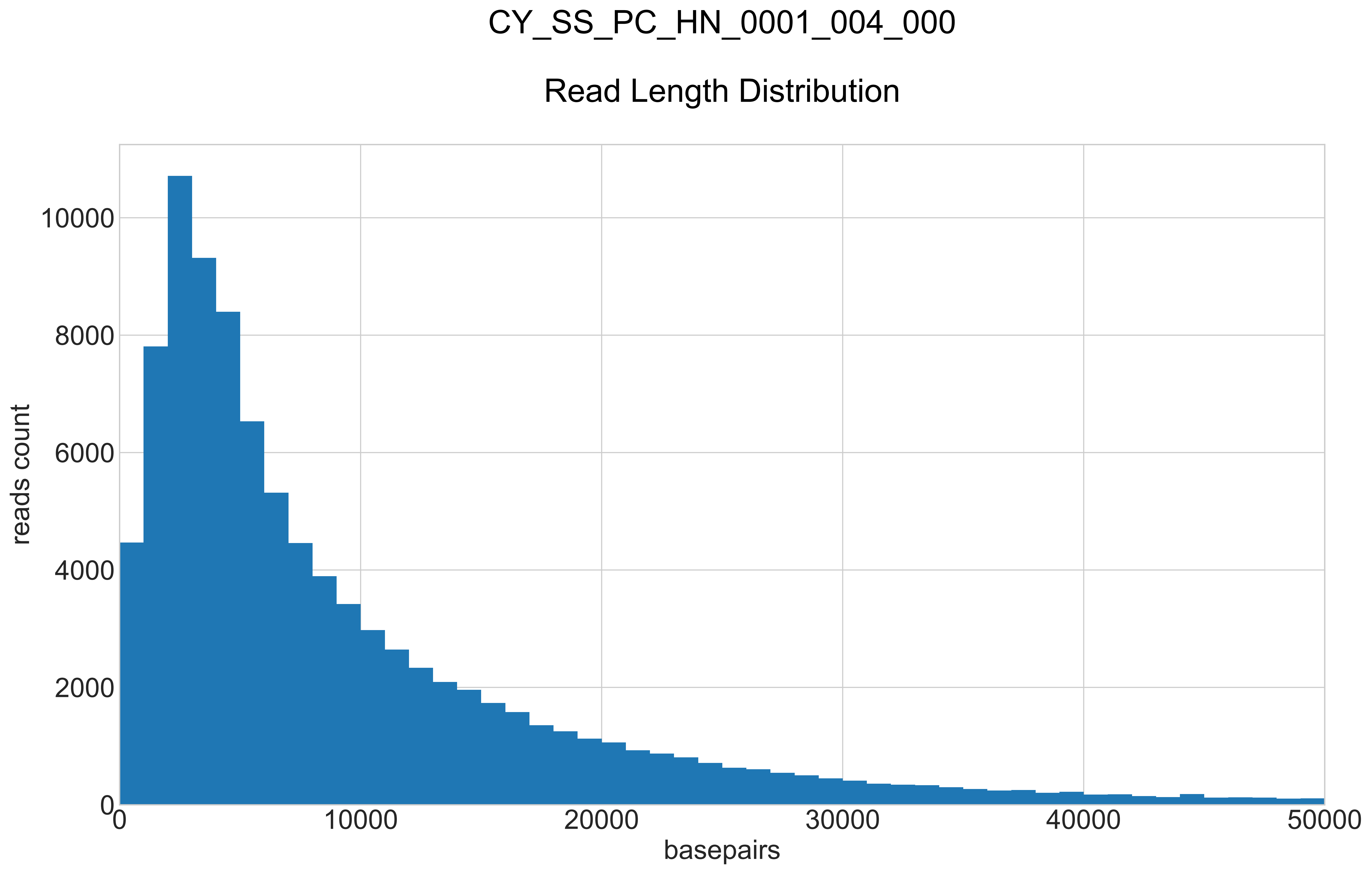

### CY_SS_PC_HN_0001_004_000_REPEATS_COUNT_.pdf

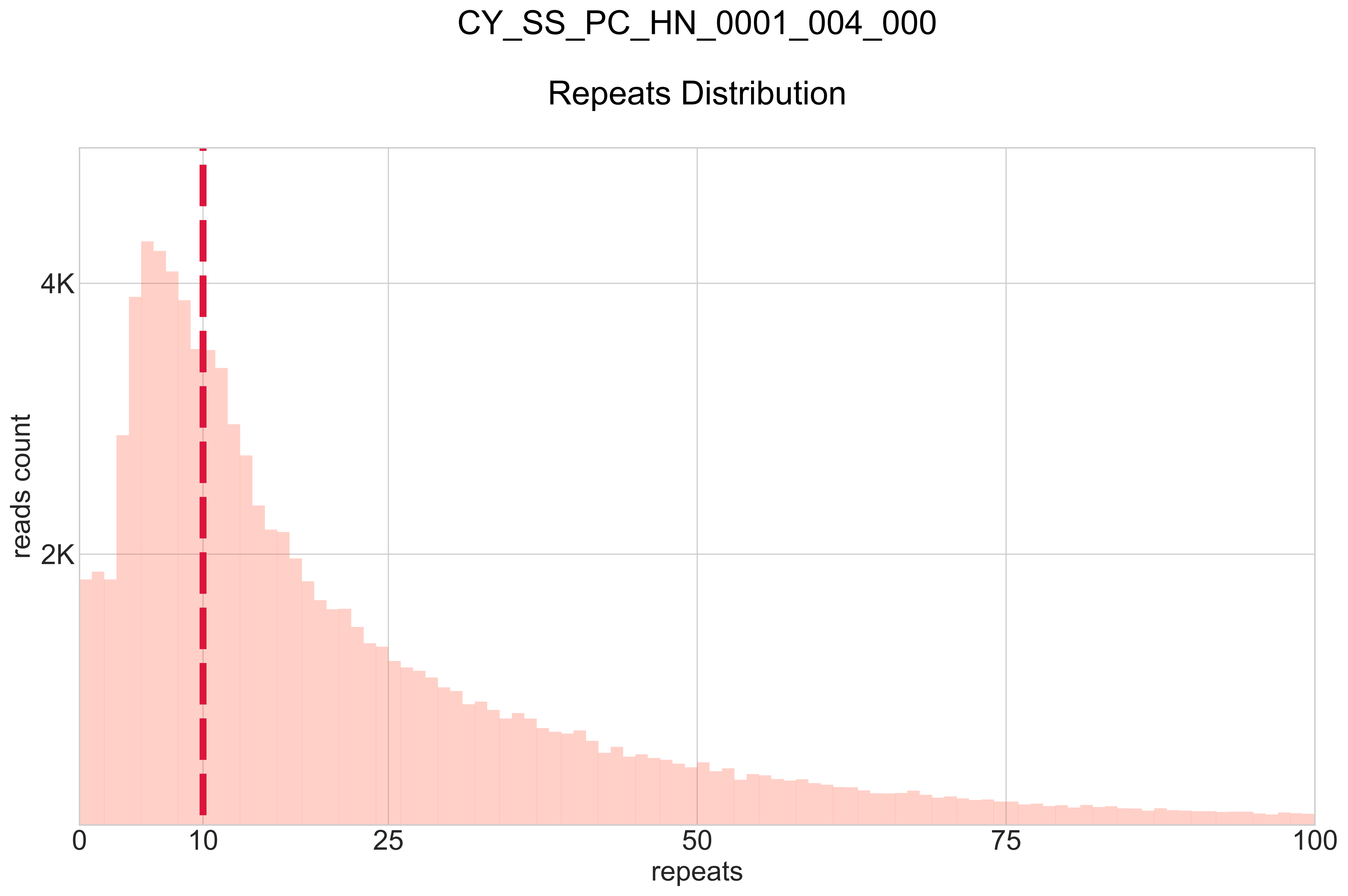

### CY_SS_PC_HN_0001_005_000_BB-I_.pdf

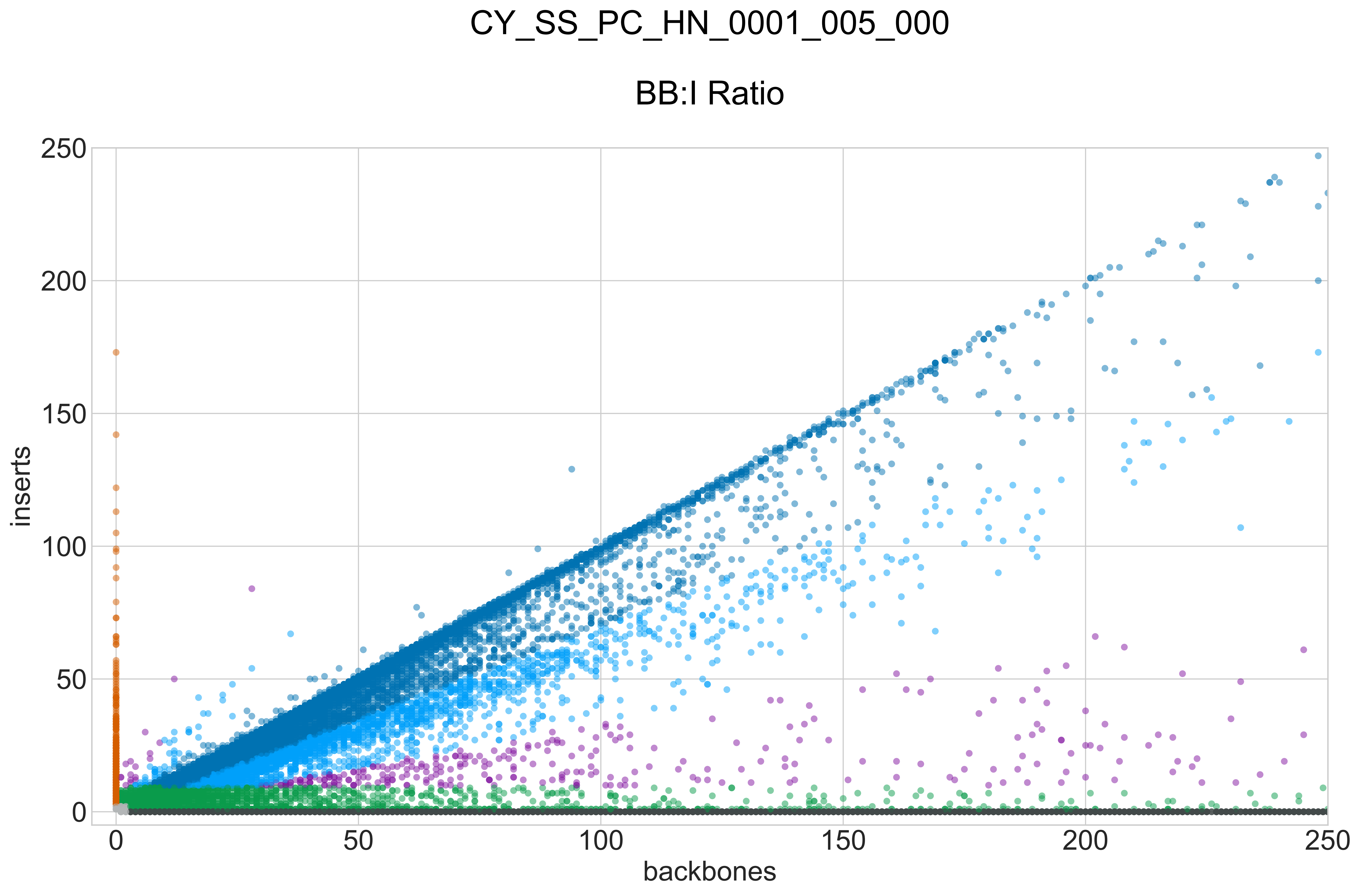

### CY_SS_PC_HN_0001_005_000_GROUPS_STACKED_NORM-LEN_.pdf

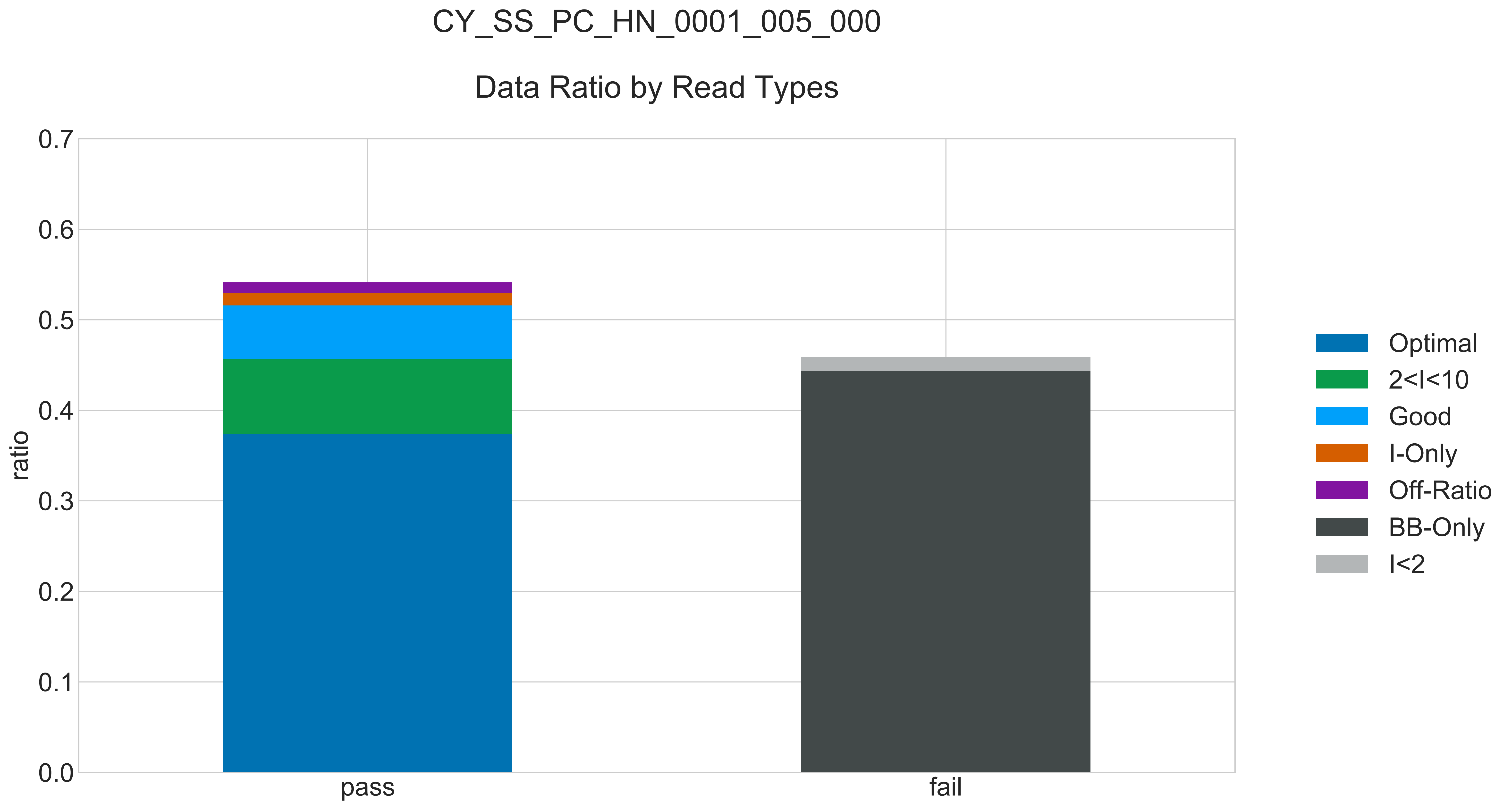

### CY_SS_PC_HN_0001_005_000_READ_LEN_.pdf

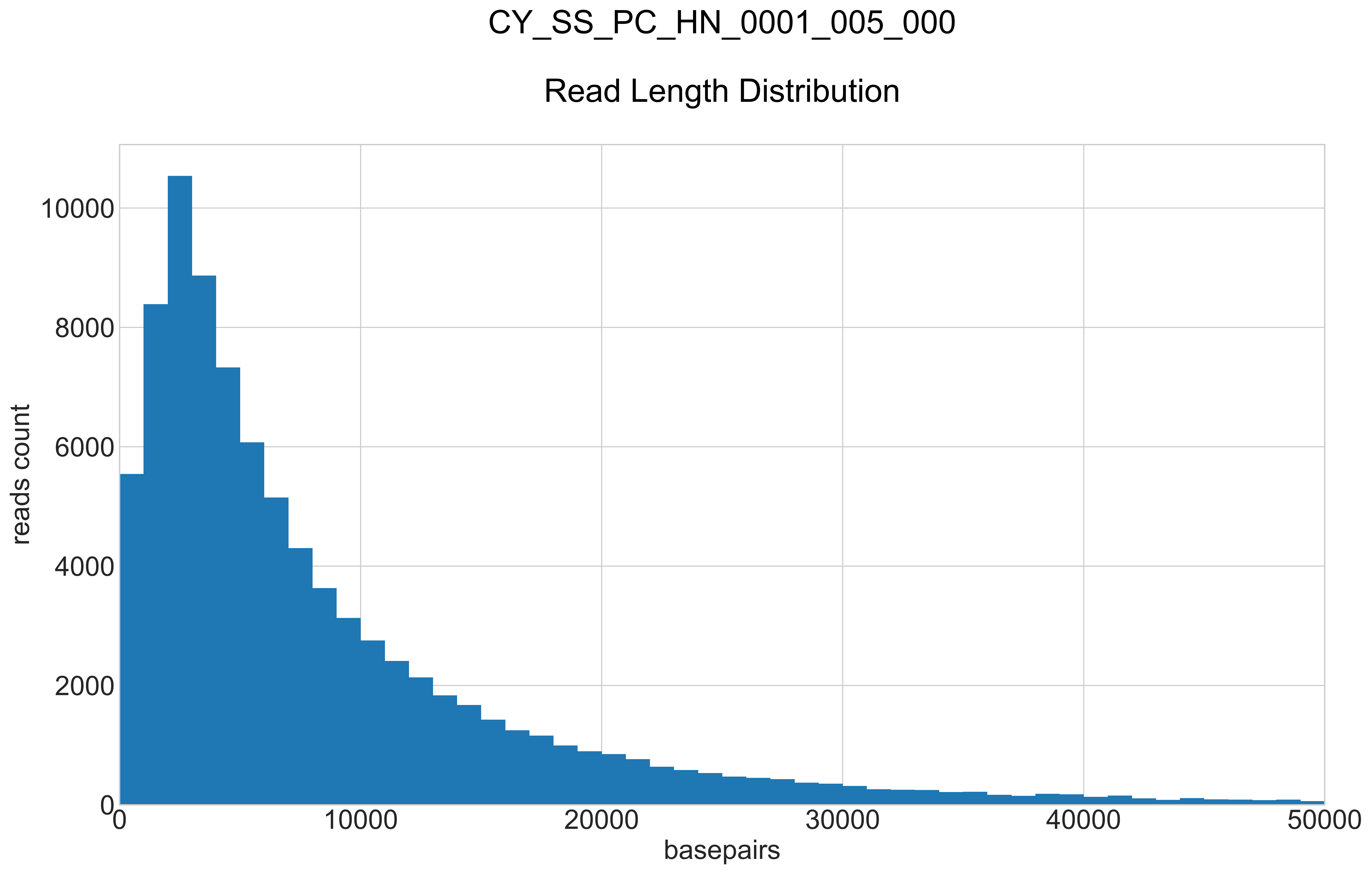

### CY_SS_PC_HN_0001_005_000_REPEATS_COUNT_.pdf

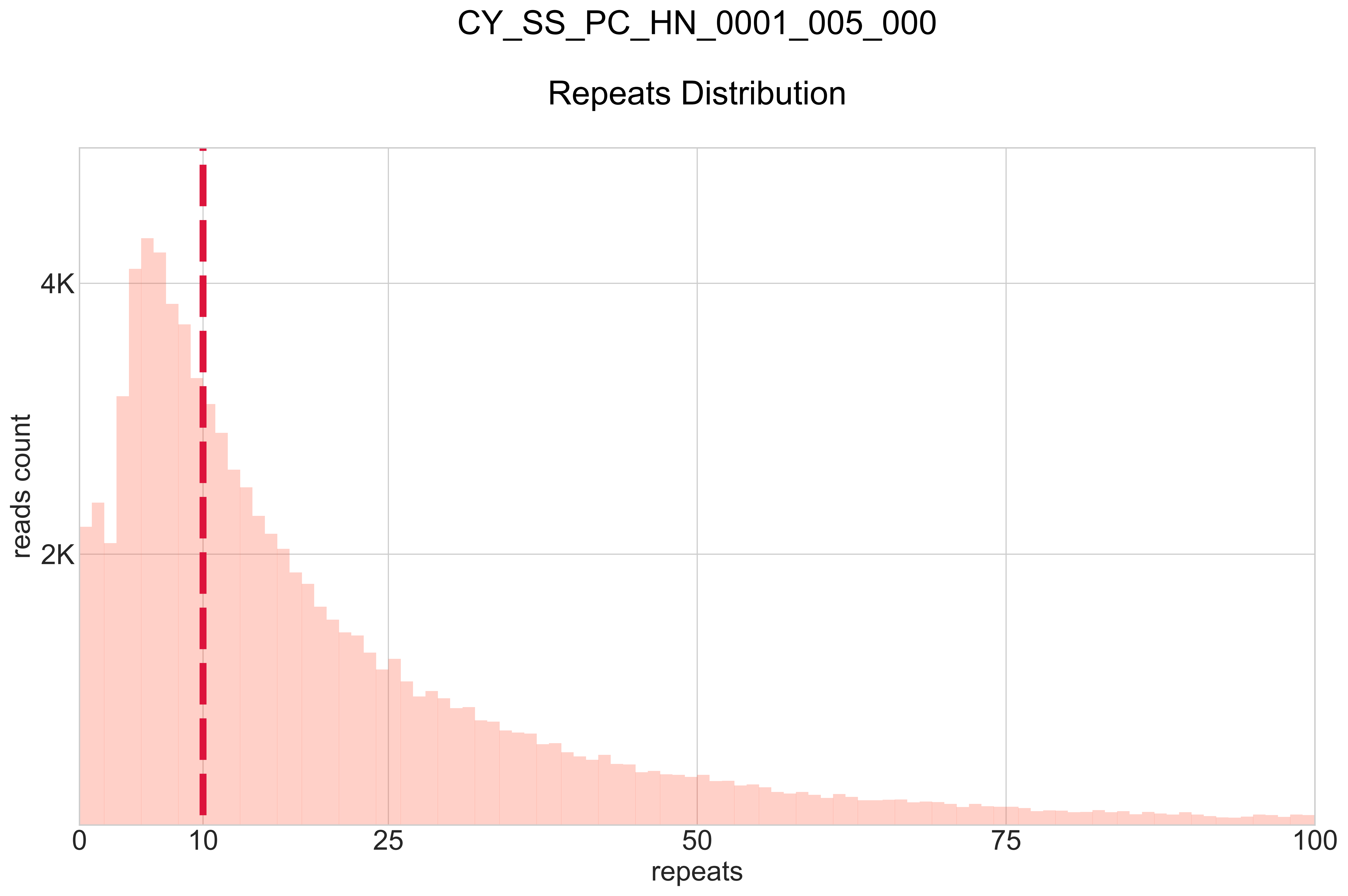
