## Supplementary Material for "Accurate detection of circulating tumor DNA using nanopore consensus sequencing"

**Supplementary Figures**


| | |
|---|---|
| Supplementary Fig. 1 | Detailed CyclomicsSeq protocol. |
| Supplementary Fig. 2 | Approach to design the three backbones used. |
| Supplementary Fig. 3 | Reproducibility of CyclomicsSeq in six technical replicates. |
| Supplementary Fig. 4 | Consensus calling lowers the error rate in the backbone. |
| Supplementary Fig. 5 | The effect of forward/reverse correction is consistent between runs. |
| Supplementary Fig. 6 | Mean single-nucleotide false positive rate across BB22 and BB25. |
| Supplementary Fig. 7 | Consensus calling lowers the error rate in Flongle flow cells. |
| Supplementary Fig. 8 | Consensus calling lowers the error rate in R10 flow cells. |
| Supplementary Fig. 9 | Detecting mutations in a single synthetic TP53 exon in pJET. |


**Supplementary Tables**


| | |
|---|---|
| Supplementary Table S1 | Sequences of backbones and inserts. |
| Supplementary Table S2 | Sample information. |
| Supplementary Table S3 | Sequencing information. |


**Supplementary Data**

Read length distribution and ratios of sequencing data grouped by read type for all CyclomicsSeq runs (provided as a separate file).

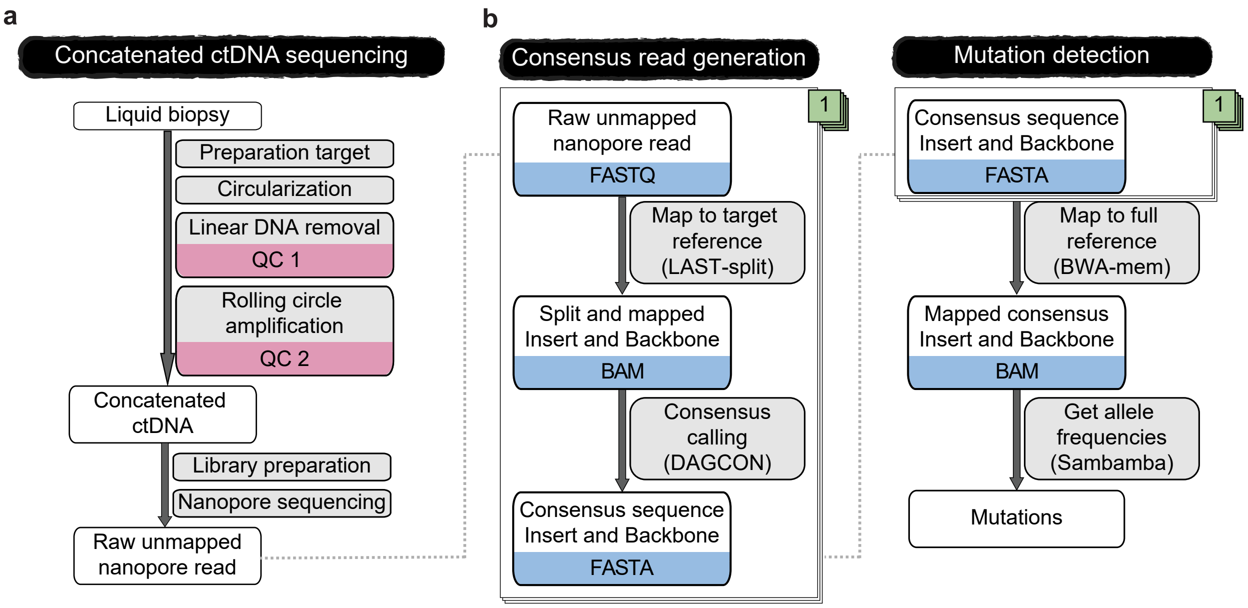


**Supplementary Fig. 1** Detailed CyclomicsSeq protocol. **a** Schematic overview of the wet-lab steps of CyclomicsSeq. QC = quality control. **b** Schematic overview of the dry-lab steps of CyclomicsSeq.

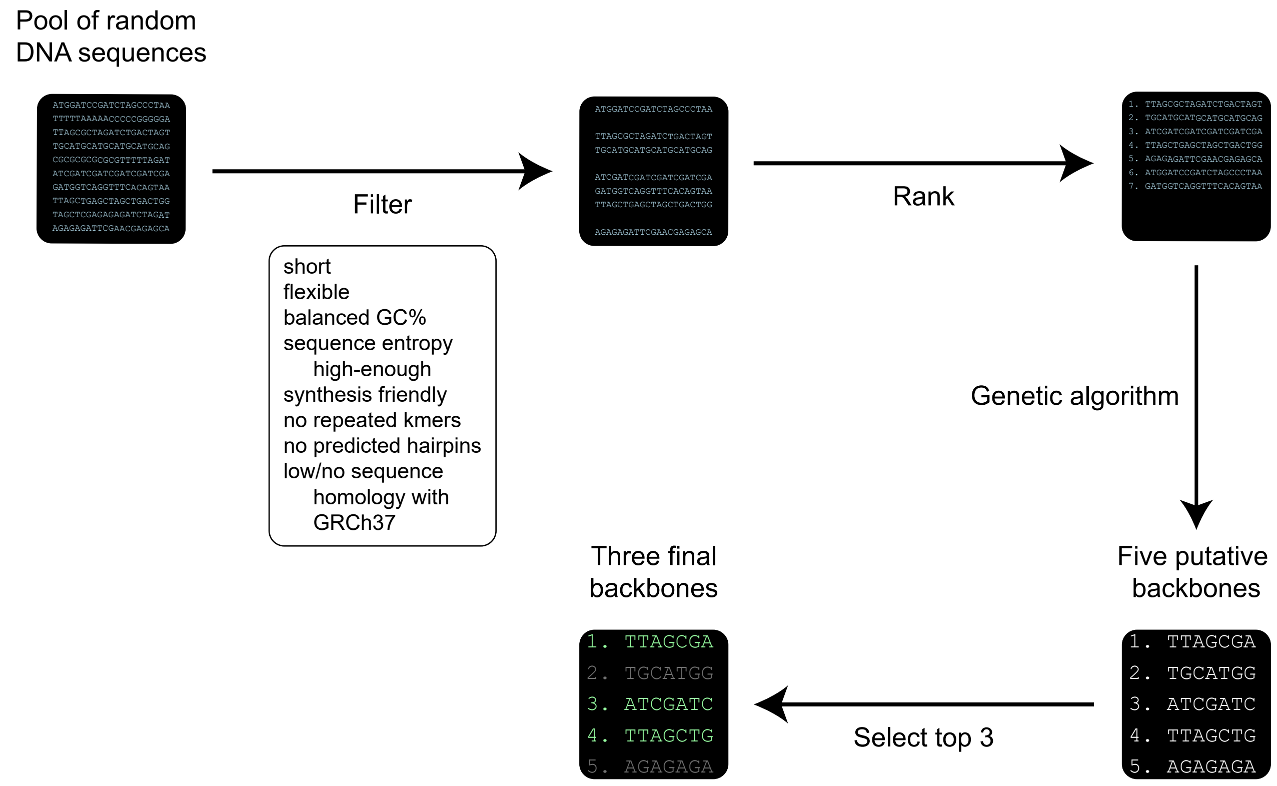


**Supplementary Fig. 2** Approach to design the three backbones used: BB22, BB24 and BB25.

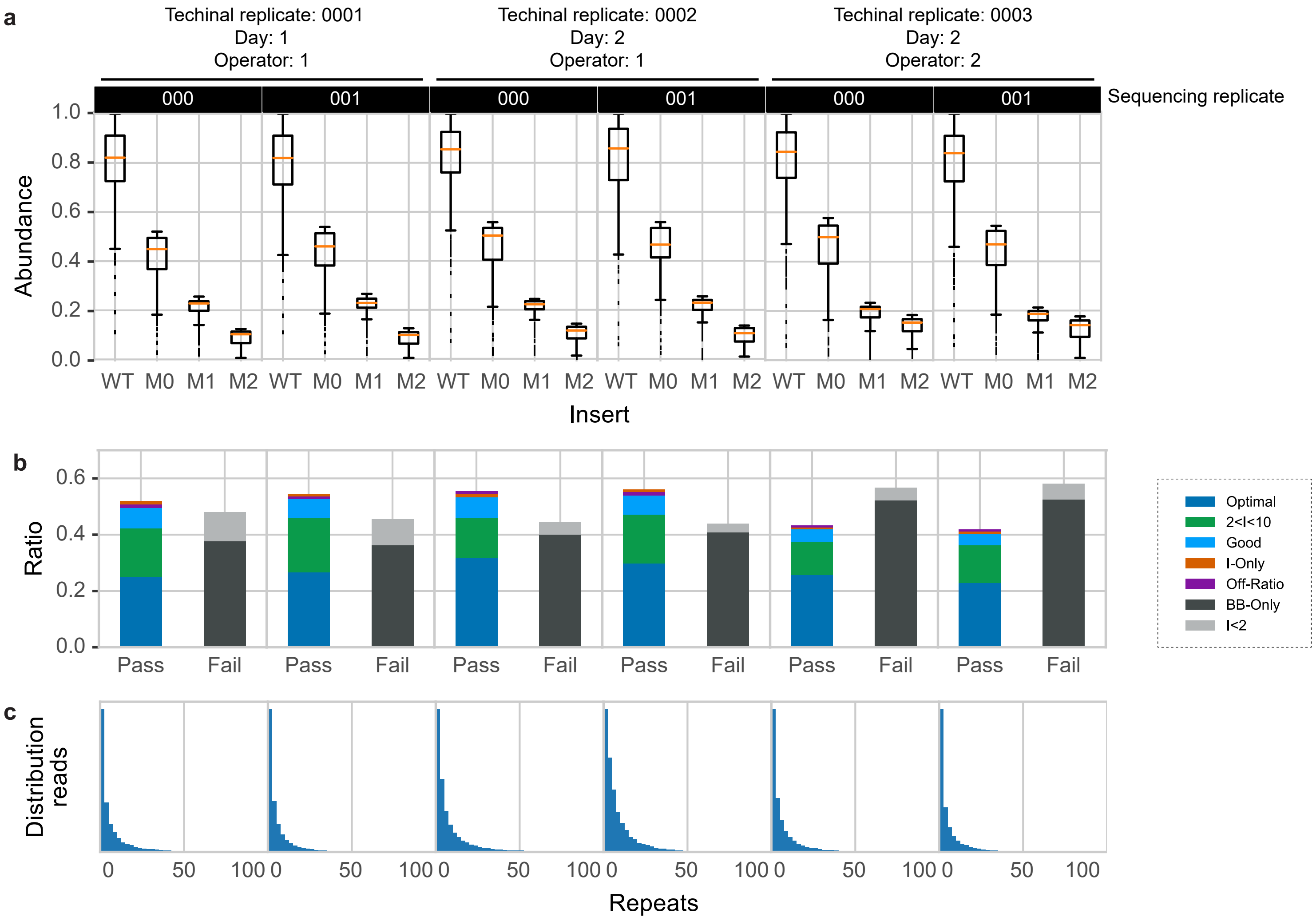


**Supplementary Fig. 3** Reproducibility of CyclomicsSeq in six technical replicates. **a** Box plots (center line = median; box limits = 25th and 75th percentiles; whiskers = 1.5x interquartile range; data points = outliers) depicting abundance of four inserts observed across the six technical replicates. WT = Wild type. M0 = Mutant 0, M1 = Mutant 1, M2 = Mutant 2. **b** Ratio of sequencing data grouped by read type for CyclomicsSeq reads. Colors, noted in the legend, represent the different categories a read can belong to (see Figure 1 for a description). **c** Number of repeats versus the number of reads. 6 Flongle runs were used for these analyses. Day of processing and operator are indicated above the graphs. 000 and 001 are sequencing replicates of the same technical replicate.

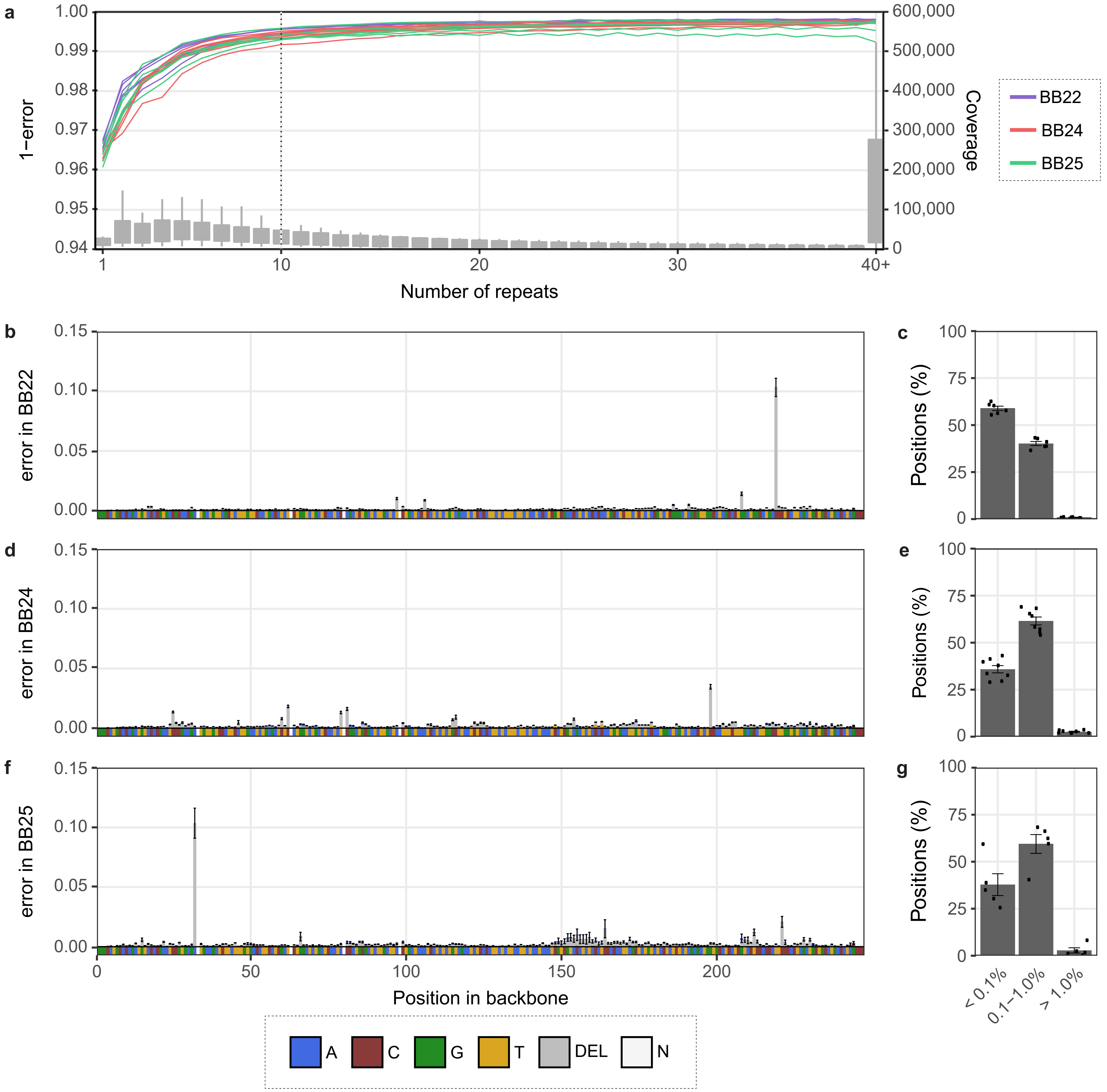


**Supplementary Fig. 4** Consensus calling lowers the error rate in the backbone. **a** 1 - Error rate (FP + deletions) in backbones (BB22, BB24, and BB25) per number of repeats. The dashed line indicates 10 repeats. Colours represent backbone type. Mean error rate across BB22 (**b**), BB24 (**d**) and BB25 (**f**) in reads with at least 10 repeats. Reference sequence is depicted below the x-axis. Colours represent base type. N = any. Percentage of positions in BB22 (**c**), BB24 (**e**) and BB25 (**g**) with indicated error percentage. Data points represent individual sequencing runs. 6 BB22, 8 BB24, and 5 BB25 runs were used for the calculations. Error bars indicate the standard deviation (sd).

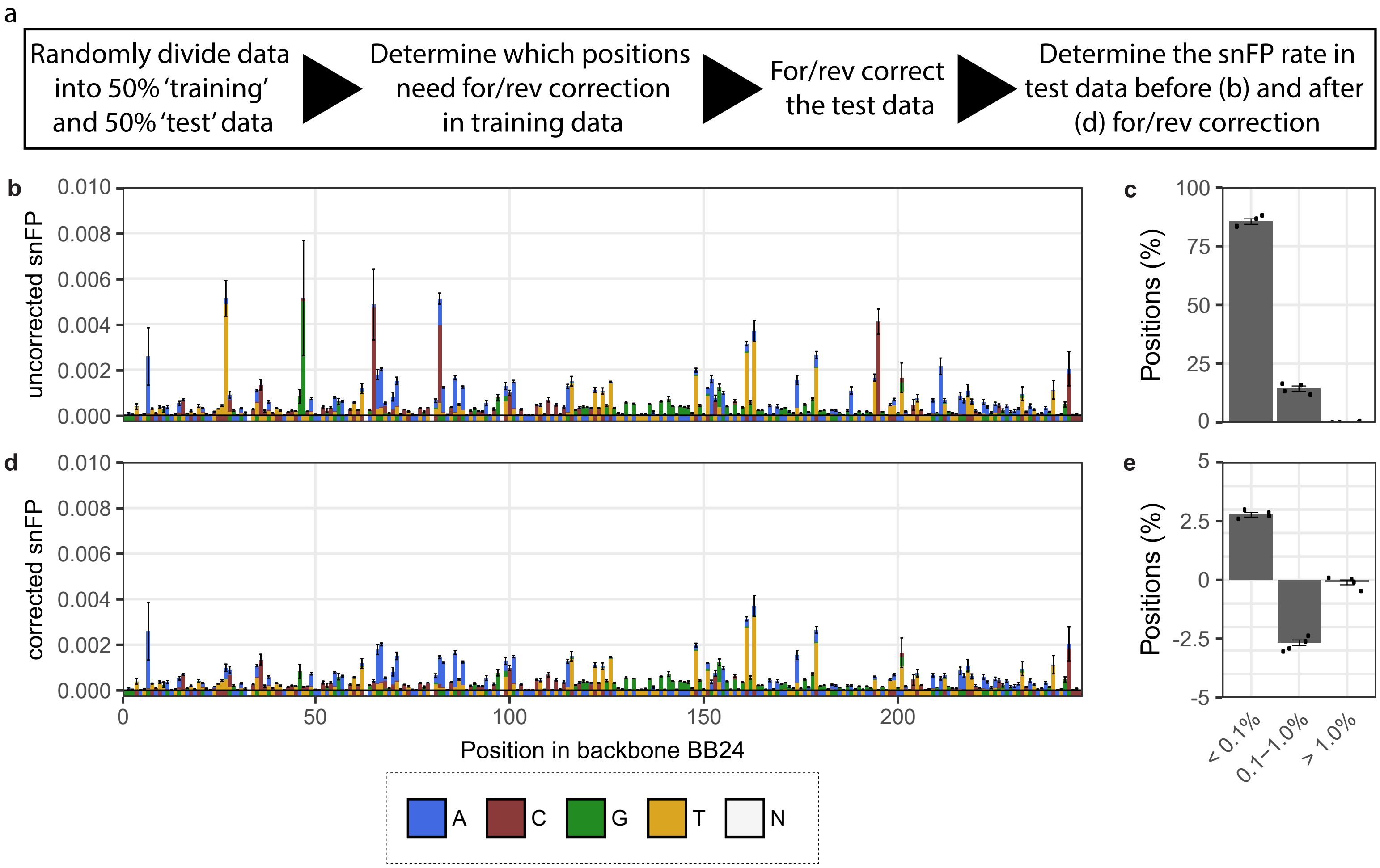


**Supplementary Fig. 5** The effect of forward/reverse correction is consistent between runs. **a** Approach to determine whether the results of the forward/reverse correction are reproducible. The samples were divided into a 50% 'training' and a 50% 'test' set. Next, the bases that require forward/reverse correction were identified in the 'training' data. The forward/reverse correction was subsequently applied to the 'test data'. Finally, the snFP rate before and after forward/reverse correction was compared in the 'test data'. **b** Mean snFP rate across BB24 in the 'test data' in reads with at least 10 repeats prior to and **d** after forward/reverse correction. Reference sequence is depicted below the x-axis. Colours represent base type. N = any. **c** Percentage of positions in BB24 with indicated snFP percentage in the 'test data' prior to forward/reverse correction and **e** the difference after forward/reverse correction. Data points represent individual sequencing runs. 8 BB24 runs were used for the calculations. Error bars indicate the standard deviation (sd). snFP = single nucleotide false positive.

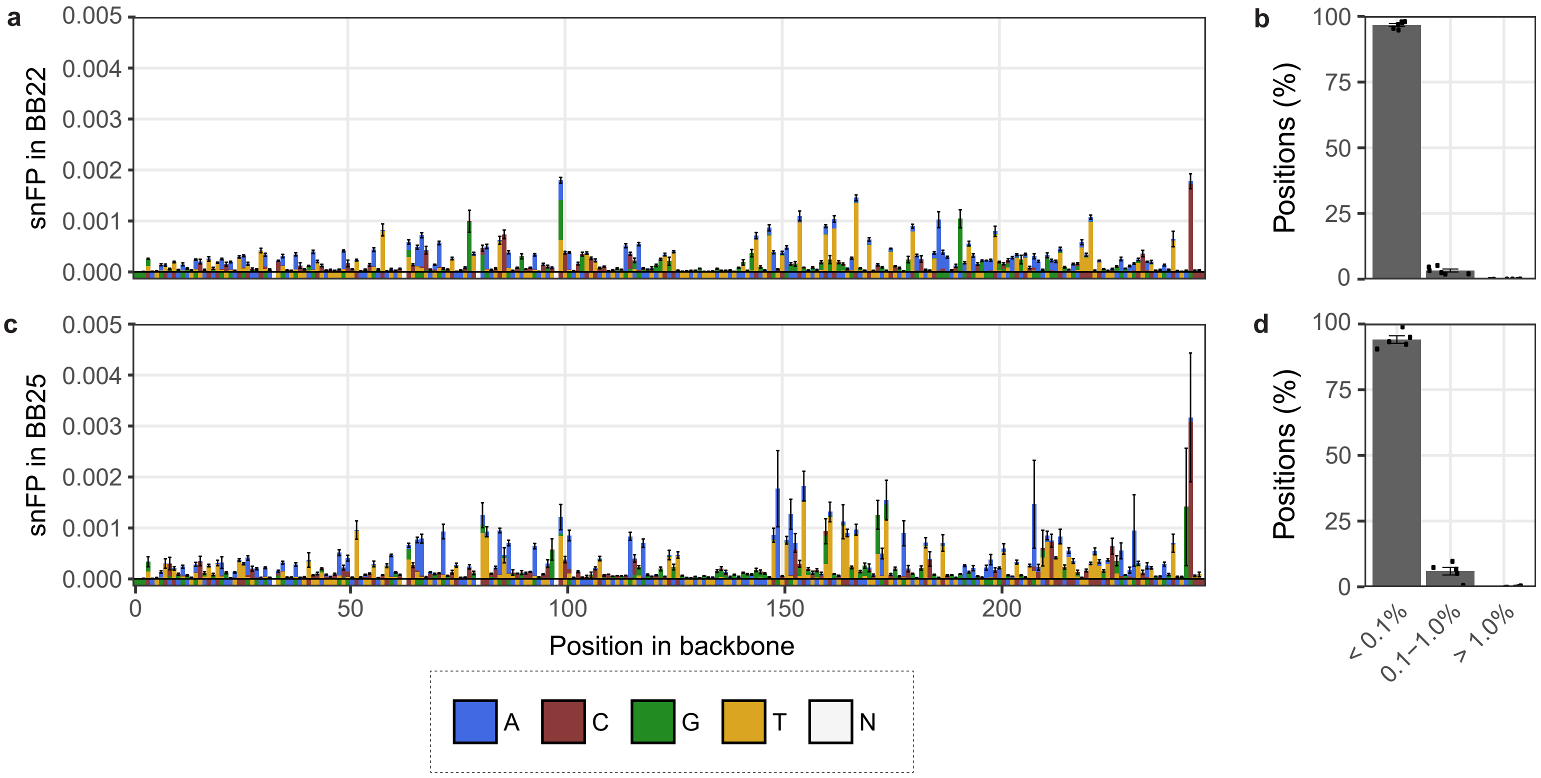


**Supplementary Fig. 6** Mean single-nucleotide false positive rate across BB22 (**a**) and BB25 (**c**) in reads with at least 10 repeats. Reference sequence is depicted below the x-axis. Colours represent base type. N = any. Percentage of positions in BB22 (**b**) and BB25 (**d**) with indicated snFP percentage. Data points represent individual sequencing runs. 6 BB22 and 5 BB25 runs were used for the calculations. Error bars indicate the standard deviation (sd).

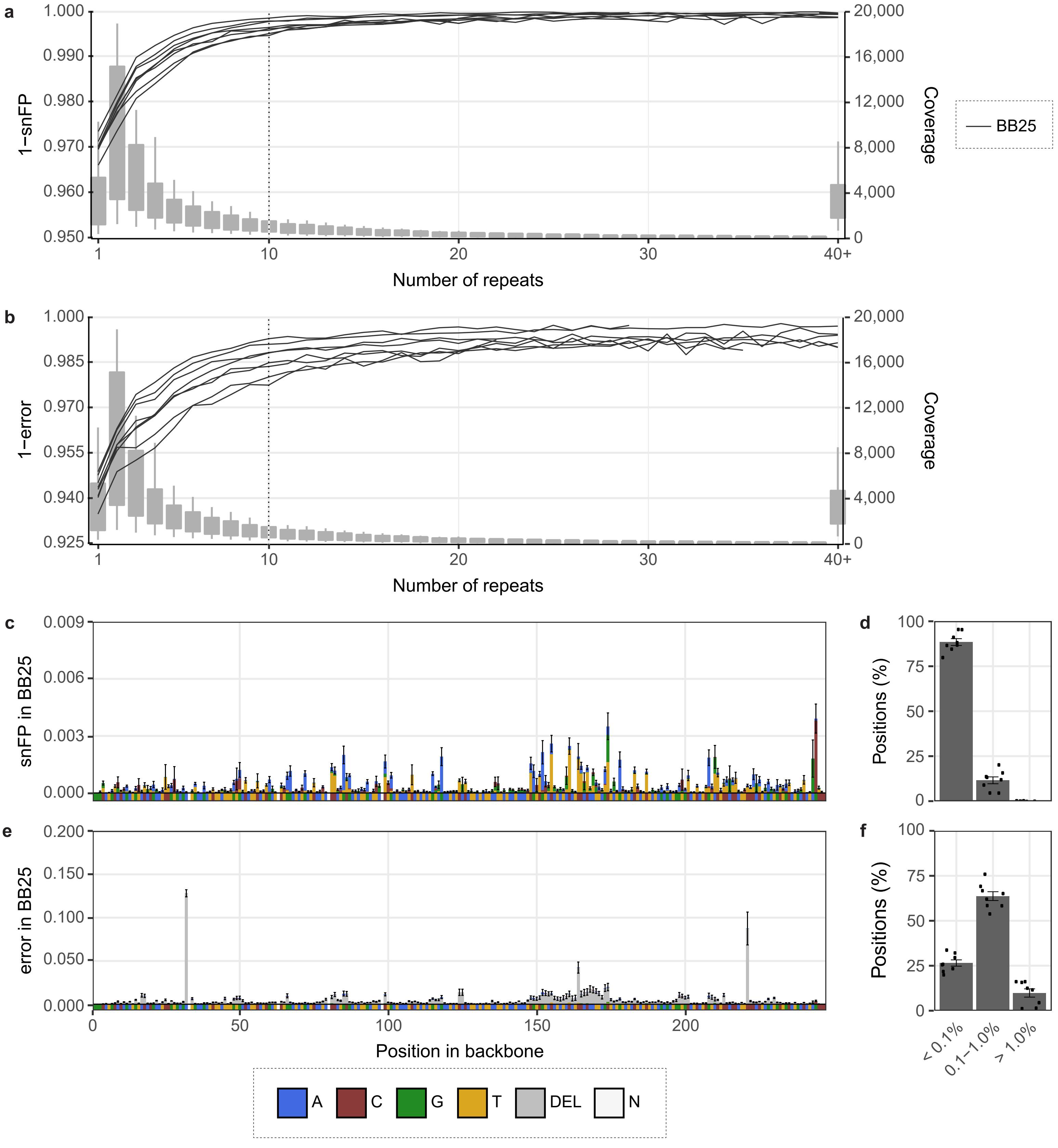


**Supplementary Fig. 7** Consensus calling lowers the number of sequencing errors in the backbone sequenced with Flongle. **a** 1 - single nucleotide false positive (snFP) and **b** 1- error (snFP + deletion) in backbone BB25 per number of repeats. The dashed line indicates 10 repeats. **c** Mean snFP and e error rate across BB25 in reads with at least 10 repeats. Reference sequence is depicted below the x-axis. Colours represent base type. N = any. **d** Percentage of positions in BB25 with indicated snFP and **f** error percentage. Data points represent individual sequencing runs. 8 BB25 runs were used for the calculations. Error bars indicate the standard deviation (sd).

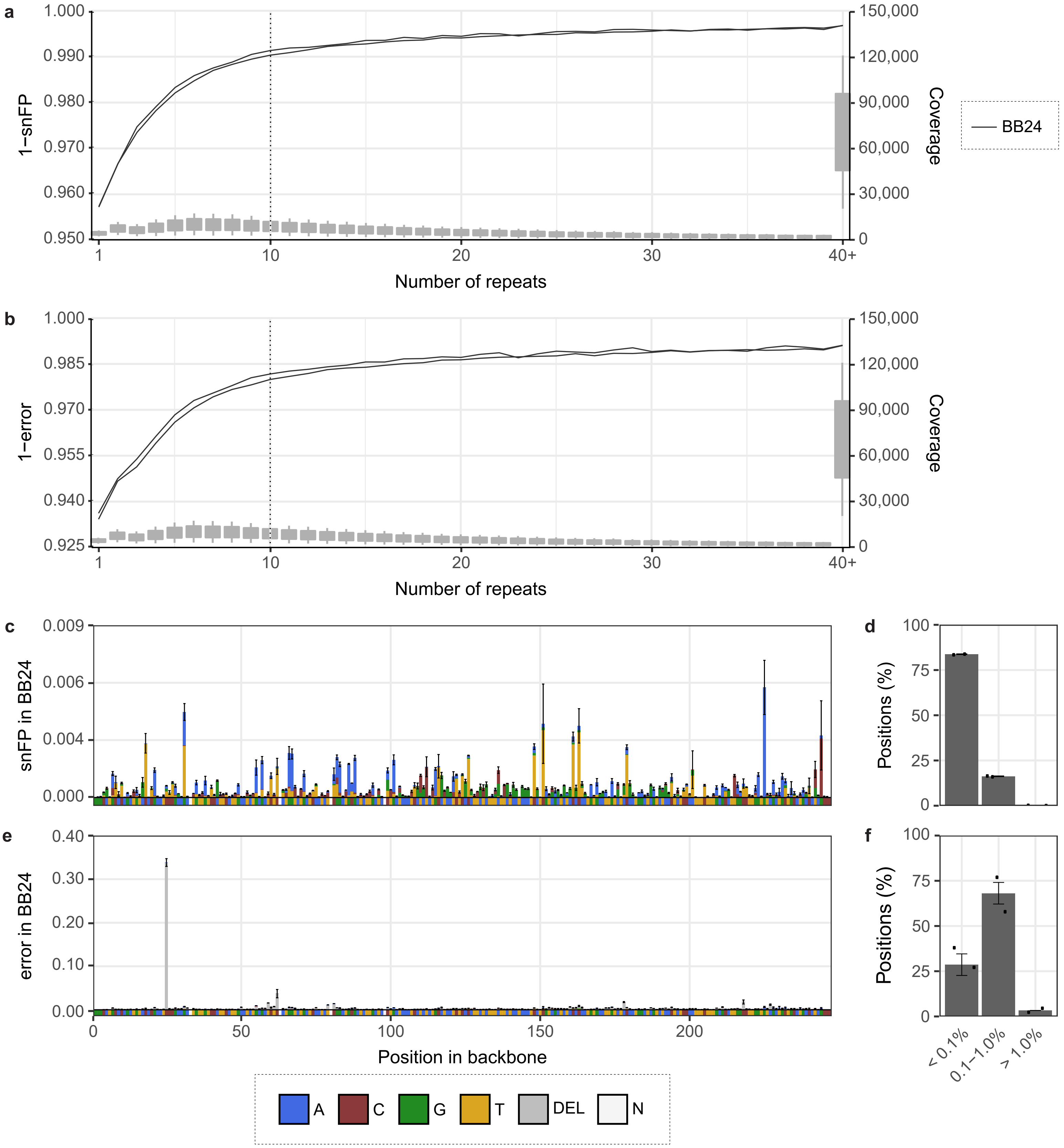


**Supplementary Fig. 8** Consensus calling lowers the number of sequencing errors in the backbone sequenced with R10 flow cells. **a** 1 - single nucleotide false positive (snFP) and **b** 1- error (snFP + deletion) in backbone BB24 per number of repeats. The dashed line indicates 10 repeats. **c** Mean snFP and **e** error rate across BB24 in reads with at least 10 repeats. Reference sequence is depicted below the x-axis. Colours represent base type. N = any. **d** Percentage of positions in BB24 with indicated snFP and **f** error percentage. Data points represent individual sequencing runs. 2 BB24 runs were used for the calculations. Error bars indicate the standard deviation (sd).

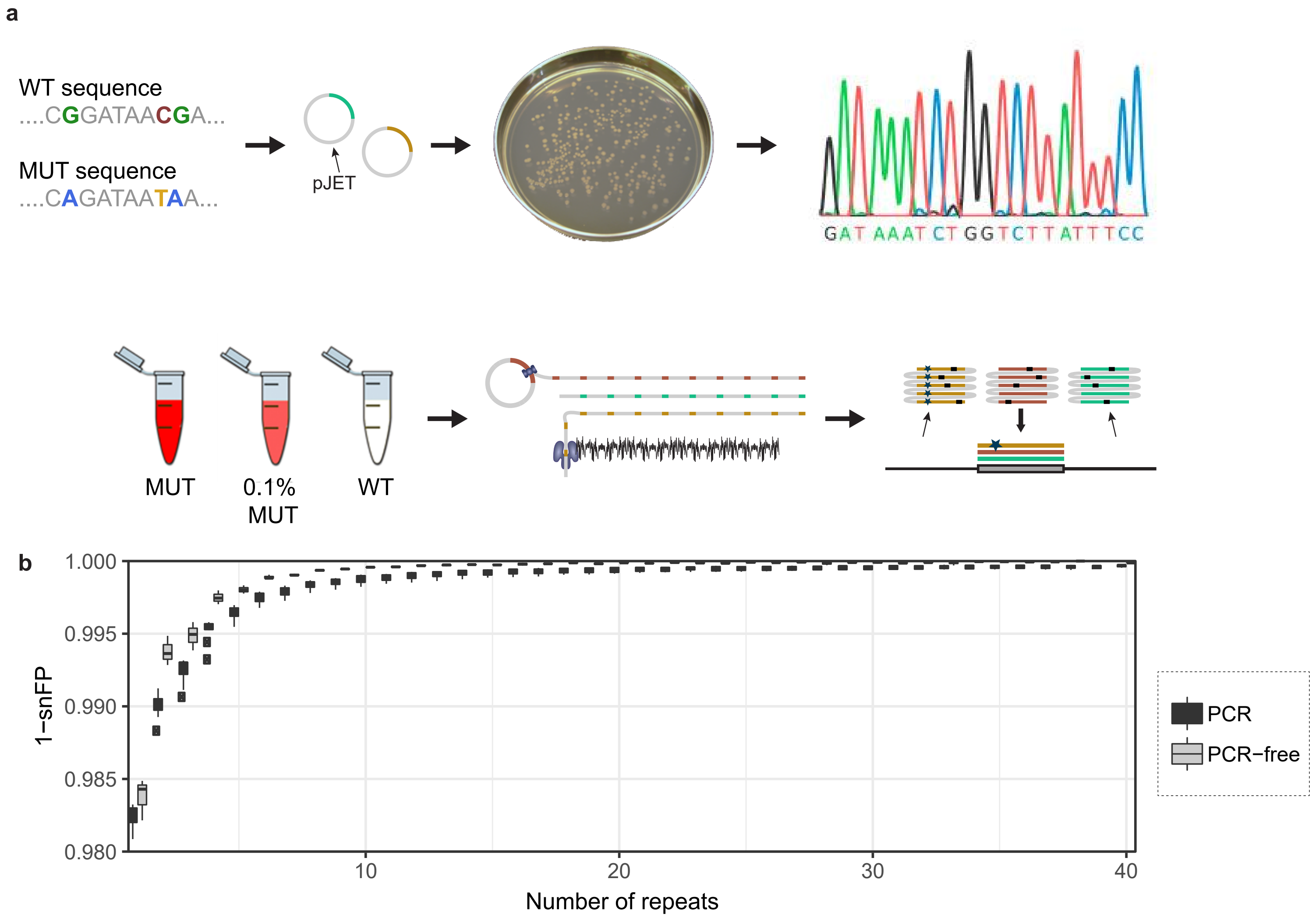


**Supplementary Fig. 9** Detecting mutations in a single synthetic *TP53* exon in pJET using CyclomicsSeq. **a** Experimental setup of the experiment. A synthetic WT sequence and MUT sequence (with three common COSMIC mutations in HNSCC) were cloned into pJET. Both sequences were amplified *in vivo* (PCR-free). Sanger sequencing was used to confirm the correctness of the WT and MUT sequence. CyclomicsSeq was performed on 100% WT, 100% MUT, and WT supplemented with 0.1% MUT. **b** Box plots (center line = median; box limits = 25th and 75th percentiles; whiskers = 1.5x interquartile range; data points = outliers) depicting 1 - single nucleotide false positive (snFP) rate in the insert (17:7577010-7577150 in GRCh37) per number of repeats for 8 PCR and 3 PCR-free inserts.

| Name | Genomic coordinates | Sequence |
|---|---|---|
| BB22 | NA | GGGCATGCACAGATGTACACGTGACGCAACGANTGATGTTAGCTATTTGTTCAATGACATATNCTGGTATGATCAATACNAGATCTGATAT<br>TGATATNCTGATACTCATATATGTAGAATATCACATTATATTTATTATAATACATCGTCGAACATATACACAATGCATCTTATCTATACGTAT<br>CGGGATAGCGTTGGCATAGCACTGGATGGCATGACCCTCATTAGATGCTGCATGACATAGCCC |
| BB24 | NA | GGGCATGCACAGATGTACACGAATCCCGAAGANTGTTGTCCATTCATTGAATATGAGATCTCNATGGTATGATCAATATNCGGATGCGAT<br>ATTGATANCTGATAAATCATATATGCATAATCTCACATTATATTTATTATAATAAATCATCGTAGATATACACAATGTGAATTGTATACAATG<br>GATAGTATAACTATCCAATTTCTTTGAGCATTGGCCTTGGTGTAGATGCTGCATGACATAGCCC |
| BB25 | NA | GGGCATGCACAGATGTACACGAATCCGTGAGANTGAAGATCTTATTTGTGACATTCATCGATNCTGGATATGATCAATANCCATGCGATAT<br>TGATTANCTGATAAATCATATATGTAGAATATCACATTATATTAATTATAATAAATCGTCGTACATATACATCCACAATTAGCTATGTATACT<br>ATCTATAGAGATGGTGCATCATCGTACTCCACCATTCCCACTAGATGCTGCATGACATAGCCC |
| pJET | NA | See https://assets.thermofisher.com/TFS-Assets/LSG/brochures/pJET1.2-plasmid-sequence.txt |
| *TP53* amplicon5 | 17:7573954-7574071 | TCCTTCCCAGCCTGGGCATCCTTGAGTTCCAAGGCCTCATTCAGCTCTCGGAACATCTCGAAGCGCTCACGCCCACGGATCTGCAGCAACA<br>GAGGAGGGGGAGAAGTAAGTATATACA |
| *TP53* amplicon9 | 17:7576810-7576955 | ACTGGAAACTTTCCACTTGATAAGAGGTCCCAAGACTTAGTACCTGAAGGGTGAAATATTCTCCATCCAGTGGTTTCTTCTTTGGCTGGGGA<br>GAGGAGCTGGTGTTGTTGGGCAGTGCTAGGAAAGAGGCAAGGAAAGGTGATAAA |
| *TP53* amplicon12 | 17:7577010-7577150 | CTTGCTTACCTCGCTTAGTGCTCCCTGGGGGCAGCTCGTGGTGAGGCTCCCCTTTCTTGCGGAGATTCTCTTCCTCTGTGCGCCGGTCTCTCC<br>CAGGACAGGCACAAACACGCACCTCAAAGCTGTTCCGTCCCAGTAGAT |
| *TP53* amplicon19 | 17:7578343-7578452 | TGTCGTCTCTCCAGCCCCAGCTGCTCACCATCGCTATCTGAGCAGCGCTCATGGTGGGGGCAGCGCCTCACAACCTCCGTCATGTGCTGTG<br>ACTGCTTGTAGATGGCCAT |
| *TP53* amplicon20 | 17:7578409-7578545 | CTCACAACCTCCGTCATGTGCTGTGACTGCTTGTAGATGGCCATGGCGCGGACGCGGGTGCCGGGCGGGGGTGTGGAATCAACCCACAG<br>CTGCACAGGGCAGGTCTTGGCCAGTTGGCAAAACATCTTGTTGAGGGC |
| *TP53* amplicon21 | 17:7578476-7578592 | GGGGTGTGGAATCAACCCACAGCTGCACAGGGCAGGTCTTGGCCAGTTGGCAAAACATCTTGTTGAGGGCAGGGGAGTACTGTAGGAAG<br>AGGAAGGAGACAGAGTTGAAAGTCAGGG |
| *TP53* ampliconS0 | 17:7579205-7579355 | GAAGCCAAAGGGTGAAGAGGAATCCCAAAGTTCCAAACAAAAGAAATGCAGGGGGATACGGCCAGGCATTGAAGTCTCATGGAAGCCA<br>GCCCCTCAGGGCAACTGACCGTGCAAGTCACAGACTTGGCTGTCCCAGAATGCAAGAAGCCCA |

**Supplementary Table S1.** Sequences of backbones and inserts

| Sample ID | Sample name | Sample type | Sample source | Mutation | Time point |
|---|---|---|---|---|---|
| CY_SS_PC_HC_0001_001_000 | Control A | Healthy control | Plasma cfDNA | NA | NA |
| CY_SM_PC_HC_0002_001_000 | Control B | Healthy control | Plasma cfDNA | NA | NA |
| CY_SM_PC_HC_0004_001_000 | Control C | Healthy control | Plasma cfDNA | NA | NA |
| CY_SM_PC_HC_0004_002_000 | Control C | Healthy control | Plasma cfDNA | NA | NA |
| CY_SM_PC_HC_0004_003_000 | Control C | Healthy control | Plasma cfDNA | NA | NA |
| CY_SM_PC_HC_0004_004_000 | Control C | Healthy control | Plasma cfDNA | NA | NA |
| CY_BB25_19WT_0001_000 | Control D | Healthy control | Plasma cfDNA | NA | NA |
| CY_SS_PC_HC_0005_002_000 | Control E | Healthy control | Plasma cfDNA | NA | NA |
| CY_SS_PC_HN_0001_001_000 | Patient A 0B | HNSCC | Plasma cfDNA | 17:7577121 G>A | 0 |
| CY_SS_PC_HN_0001_002_000 | Patient A 1B | HNSCC | Plasma cfDNA | 17:7577121 G>A | 1 |
| CY_SS_PC_HN_0001_005_000 | Patient A 1B | HNSCC | Plasma cfDNA | 17:7577121 G>A | 1 |
| CY_SS_PC_HN_0001_003_000 | Patient A 2B | HNSCC | Plasma cfDNA | 17:7577121 G>A | 2 |
| CY_SS_SC_HN_0001_006_000 | Patient A 2S | HNSCC | Saliva cfDNA | 17:7577121 G>A | 2 |
| CY_SS_PC_HN_0001_004_000 | Patient A 3B | HNSCC | Plasma cfDNA | 17:7577121 G>A | 4 |
| CY_SM_PC_HN_0002_001_000 | Patient B 0B | HNSCC | Plasma cfDNA | 17:7577095-7577123 Deletion | 0 |
| CY_SM_PC_HN_0002_002_001 | Patient B 0B | HNSCC | Plasma cfDNA | 17:7577095-7577123 Deletion | 0 |
| CY_SM_PC_HN_0002_003_000 | Patient B 1B | HNSCC | Plasma cfDNA | 17:7577095-7577123 Deletion | 1 |
| CY_SS_PC_HN_0003_001_000 | Patient C 0B | HNSCC | Plasma cfDNA | 17:7578403 C>T | 0 |
| CY_SS_PC_HN_0003_006_000 | Patient C 19B | HNSCC | Plasma cfDNA | 17:7578403 C>T | 19 |
| CY_SS_PC_HN_0003_002_000 | Patient C 1B | HNSCC | Plasma cfDNA | 17:7578403 C>T | 1 |
| CY_SS_PC_HN_0003_003_000 | Patient C 2B | HNSCC | Plasma cfDNA | 17:7578403 C>T | 2 |
| CY_SS_PC_HN_0003_004_000 | Patient C 3B | HNSCC | Plasma cfDNA | 17:7578403 C>T | 3 |
| CY_SS_PC_HN_0003_005_000 | Patient C 4B | HNSCC | Plasma cfDNA | 17:7578403 C>T | 4 |
| CY_PJET_12MU_0001_000 | MUT | Synthetic control | Synthetic | 17:7577094 G>A ; 17:7577120 C>T ; 17:7577121 G>A | NA |
| CY_LOT1_QC_0001_000 | REP mix 1 | Synthetic control | Synthetic | Multiple mixed | NA |
| CY_LOT1_QC_0001_001 | REP mix 1 | Synthetic control | Synthetic | Multiple mixed | NA |
| CY_LOT1_QC_0002_000 | REP mix 2 | Synthetic control | Synthetic | Multiple mixed | NA |
| CY_LOT1_QC_0002_001 | REP mix 2 | Synthetic control | Synthetic | Multiple mixed | NA |
| CY_LOT1_QC_0003_000 | REP mix 3 | Synthetic control | Synthetic | Multiple mixed | NA |
| CY_LOT1_QC_0003_001 | REP mix 3 | Synthetic control | Synthetic | Multiple mixed | NA |
| CY_PJET_12WT_0001_000 | WT | Synthetic control | Synthetic | NA | NA |
| CY_PJET_RATI_0001_000 | WT/MUT mix | Synthetic control | Synthetic | 17:7577094 G>A ; 17:7577120 C>T ; 17:7577121 G>A | NA |

NA = Not applicable; WT = Wild type; MUT = Mutant; REP = reproducibility; HNSCC = head and neck squamous cell carcinoma; cfDNA = cell free DNA

**Supplementary Table S2.** Sample information

| Sample ID | Backbone | Insert | Flow cell | Reads |
|---|---|---|---|---|
| CY_SS_PC_HC_0001_001_000 | BB24 | *TP53* amplicon 12 | R9 | 1459482 |
| CY_SM_PC_HC_0002_001_000 | BB24 | *TP53* amplicon 5,9 | R9 | 1783724 |
| CY_SM_PC_HC_0004_001_000 | BB25 | *TP53* amplicon 9,19,20,21 | R9 | 3681526 |
| CY_SM_PC_HC_0004_002_000 | BB25 | *TP53* amplicon 9,19,20,21 | Flongle | 73591 |
| CY_SM_PC_HC_0004_003_000 | BB25 | CleanPlex *TP53* Panel* | R9 | 2450755 |
| CY_SM_PC_HC_0004_004_000 | BB25 | CleanPlex *TP53* Panel* | Flongle | 173694 |
| CY_BB25_19WT_0001_000 | BB25 | *TP53* amplicon 19 | R9 | 11854864 |
| CY_SS_PC_HC_0005_002_000 | BB25 | *TP53* amplicon 12 | R9 | 5345433 |
| CY_SS_PC_HN_0001_001_000 | BB24 | *TP53* amplicon 12 | R9 | 1340417 |
| CY_SS_PC_HN_0001_002_000 | BB24 | *TP53* amplicon 12 | R9 | 1343902 |
| CY_SS_PC_HN_0001_005_000 | BB24 | *TP53* amplicon 12 | R10 | 585206 |
| CY_SS_PC_HN_0001_003_000 | BB24 | *TP53* amplicon 12 | R9 | 1089610 |
| CY_SS_SC_HN_0001_006_000 | BB24 | *TP53* amplicon 12 | R9 | 1583579 |
| CY_SS_PC_HN_0001_004_000 | BB24 | *TP53* amplicon 12 | R9 | 860412 |
| CY_SM_PC_HN_0002_001_000 | BB24 | *TP53* amplicon 5,9,12 | R9 | 1019983 |
| CY_SM_PC_HN_0002_002_001 | BB24 | *TP53* amplicon 5,9,12 | R10 | 108000 |
| CY_SM_PC_HN_0002_003_000 | BB25 | *TP53* amplicon 9 | R9 | 2453166 |
| CY_SS_PC_HN_0003_001_000 | BB22 | *TP53* amplicon 19 | R9 | 3776045 |
| CY_SS_PC_HN_0003_006_000 | BB22 | *TP53* amplicon 19 | R9 | 4680602 |
| CY_SS_PC_HN_0003_002_000 | BB22 | *TP53* amplicon 19 | R9 | 4208840 |
| CY_SS_PC_HN_0003_003_000 | BB22 | *TP53* amplicon 19 | R9 | 3611187 |
| CY_SS_PC_HN_0003_004_000 | BB22 | *TP53* amplicon 19 | R9 | 2409222 |
| CY_SS_PC_HN_0003_005_000 | BB22 | *TP53* amplicon 19 | R9 | 3987095 |
| CY_PJET_12MU_0001_000 | pJET | *TP53* amplicon 12 | R9 | 3848342 |
| CY_LOT1_QC_0001_000 | BB25 | *TP53* amplicon S0 | Flongle | 45573 |
| CY_LOT1_QC_0001_001 | BB25 | *TP53* amplicon S0 | Flongle | 125446 |
| CY_LOT1_QC_0002_000 | BB25 | *TP53* amplicon S0 | Flongle | 21844 |
| CY_LOT1_QC_0002_001 | BB25 | *TP53* amplicon S0 | Flongle | 12242 |
| CY_LOT1_QC_0003_000 | BB25 | *TP53* amplicon S0 | Flongle | 70747 |
| CY_LOT1_QC_0003_001 | BB25 | *TP53* amplicon S0 | Flongle | 72964 |
| CY_PJET_12WT_0001_000 | pJET | *TP53* amplicon 12 | R9 | 2702138 |
| CY_PJET_RATI_0001_000 | pJET | *TP53* amplicon 12 | R9 | 2521207 |

* Paragon Genomics kit: https://www.paragongenomics.com/product/cleanplex-tp53-panel/

**Supplementary Table S3.** Sequencing information
