## Supplementary Figure 1 for "Accurate detection of circulating tumor DNA using nanopore consensus sequencing"

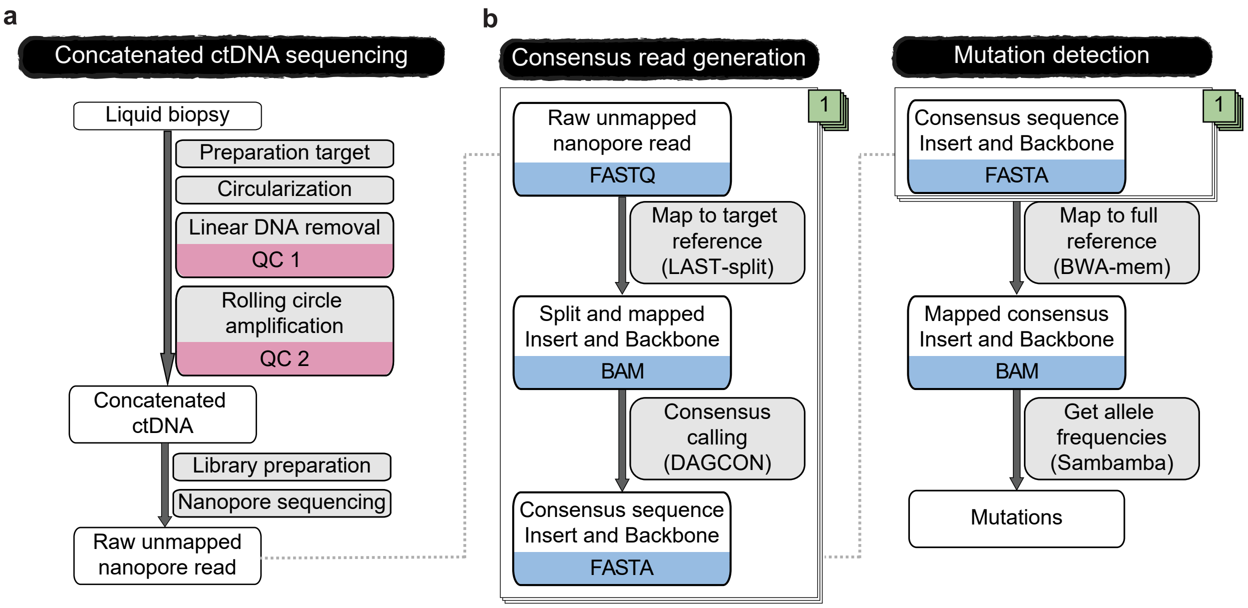


**Supplementary Fig. 1** Detailed CyclomicsSeq protocol. **a** Schematic overview of the wet-lab steps of CyclomicsSeq. QC = quality control. **b** Schematic overview of the dry-lab steps of CyclomicsSeq.
