## Supplementary Figure 3 for "Accurate detection of circulating tumor DNA using nanopore consensus sequencing"

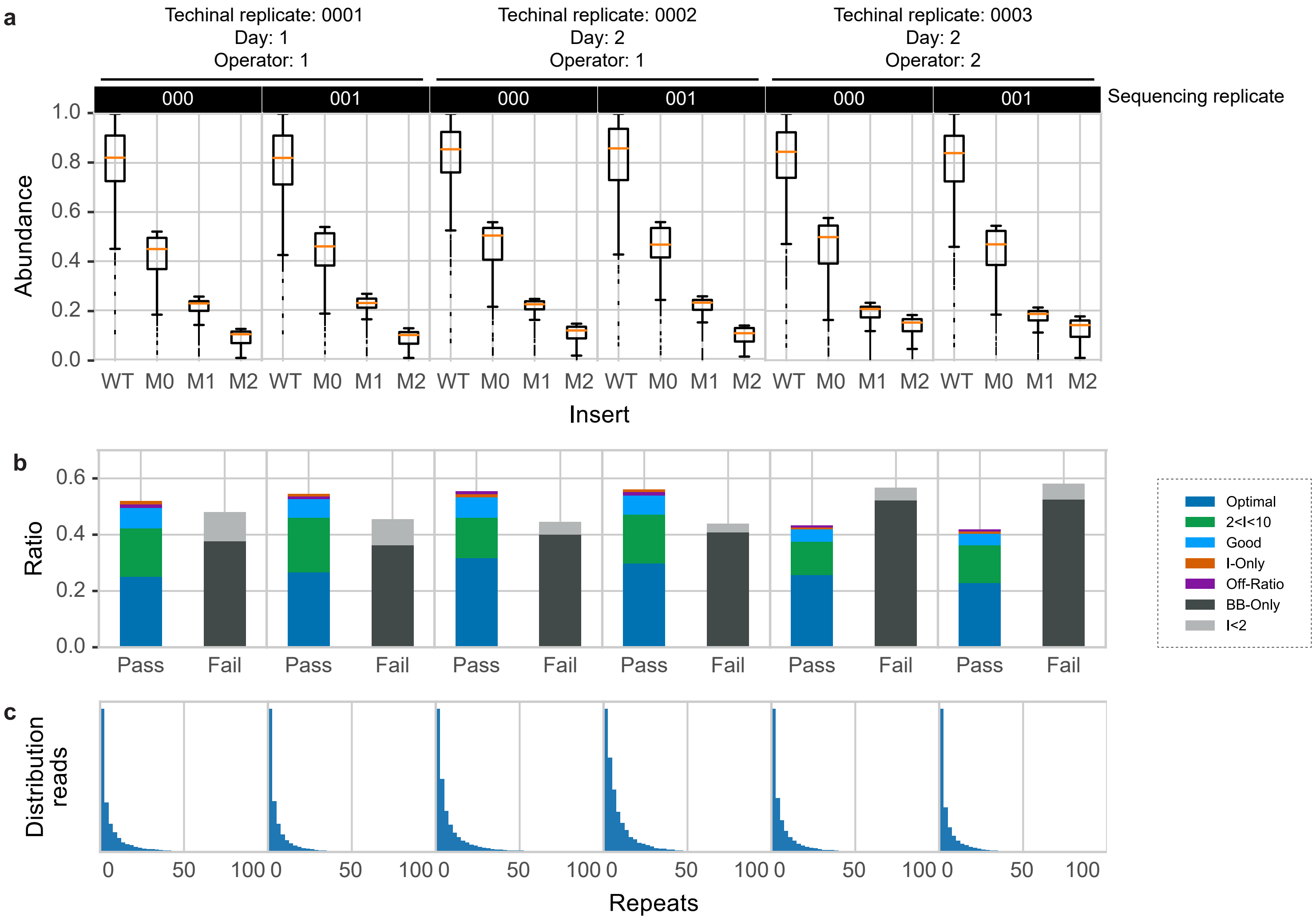


**Supplementary Fig. 3** Reproducibility of CyclomicsSeq in six technical replicates. **a** Box plots (center line = median; box limits = 25th and 75th percentiles; whiskers = 1.5x interquartile range; data points = outliers) depicting abundance of four inserts observed across the six technical replicates. WT = Wild type. M0 = Mutant 0, M1 = Mutant 1, M2 = Mutant 2. **b** Ratio of sequencing data grouped by read type for CyclomicsSeq reads. Colors, noted in the legend, represent the different categories a read can belong to (see Figure 1 for a description). **c** Number of repeats versus the number of reads. 6 Flongle runs were used for these analyses. Day of processing and operator are indicated above the graphs. 000 and 001 are sequencing replicates of the same technical replicate.
