## Supplementary Figure 4 for "Accurate detection of circulating tumor DNA using nanopore consensus sequencing"

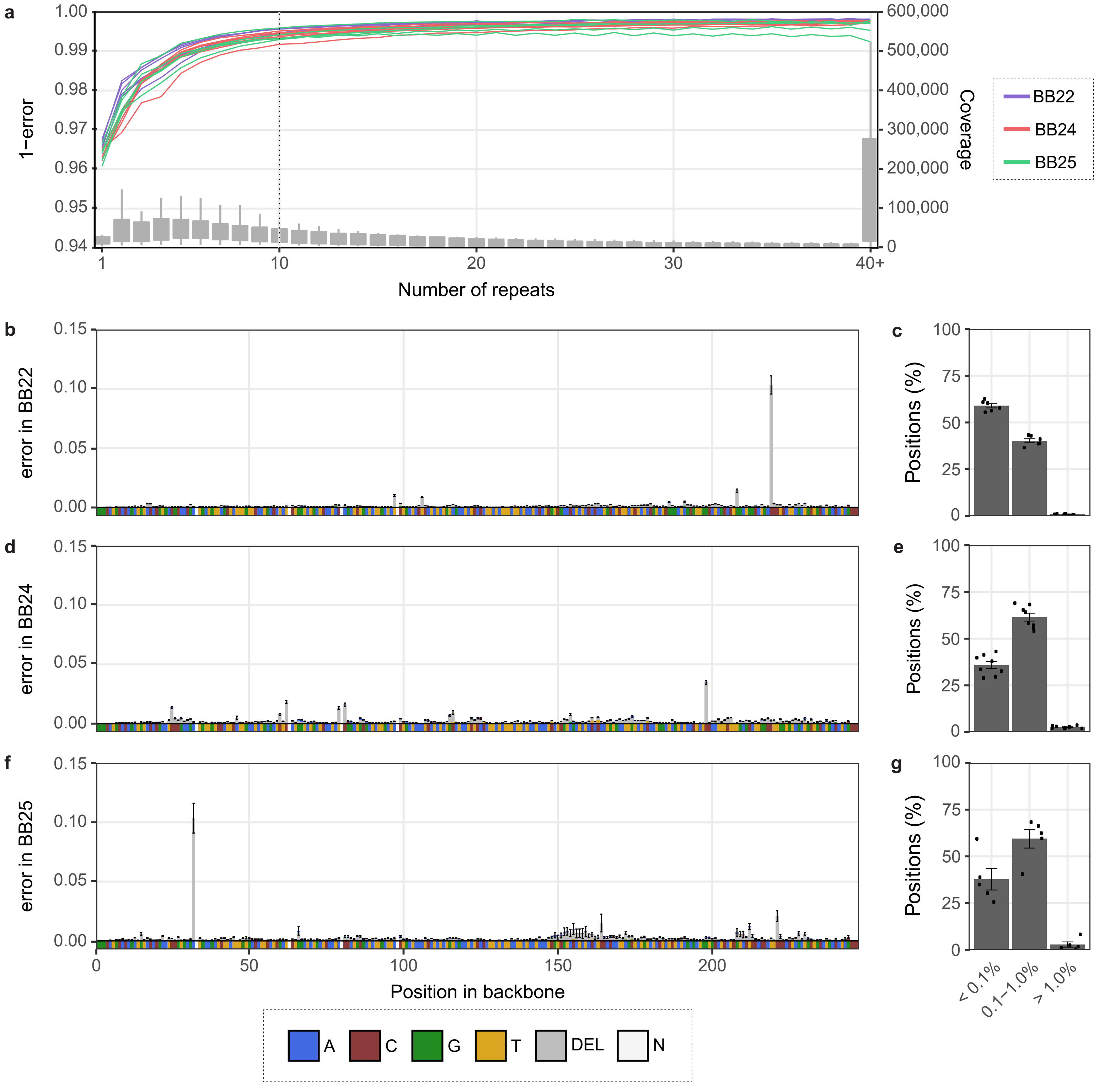


**Supplementary Fig. 4** Consensus calling lowers the error rate in the backbone. **a** 1 - Error rate (FP + deletions) in backbones (BB22, BB24, and BB25) per number of repeats. The dashed line indicates 10 repeats. Colours represent backbone type. Mean error rate across BB22 (**b**), BB24 (**d**) and BB25 (**f**) in reads with at least 10 repeats. Reference sequence is depicted below the x-axis. Colours represent base type. N = any. Percentage of positions in BB22 (**c**), BB24 (**e**) and BB25 (**g**) with indicated error percentage. Data points represent individual sequencing runs. 6 BB22, 8 BB24, and 5 BB25 runs were used for the calculations. Error bars indicate the standard deviation (sd).
