## Supplementary Figure 5 for "Accurate detection of circulating tumor DNA using nanopore consensus sequencing"

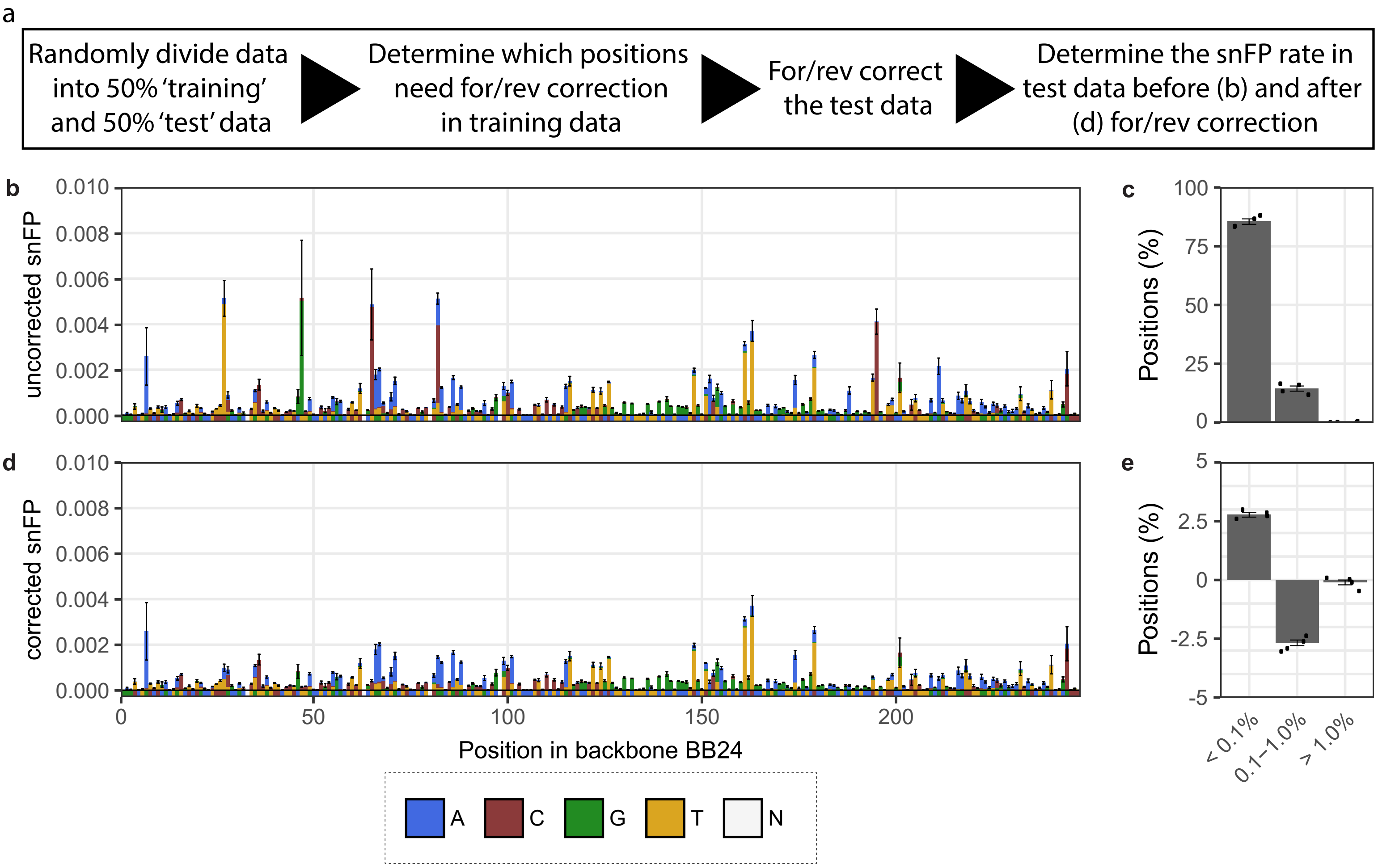


**Supplementary Fig. 5** The effect of forward/reverse correction is consistent between runs. **a** Approach to determine whether the results of the forward/reverse correction are reproducible. The samples were divided into a 50% 'training' and a 50% 'test' set. Next, the bases that require forward/reverse correction were identified in the 'training' data. The forward/reverse correction was subsequently applied to the 'test data'. Finally, the snFP rate before and after forward/reverse correction was compared in the 'test data'. **b** Mean snFP rate across BB24 in the 'test data' in reads with at least 10 repeats prior to and **d** after forward/reverse correction. Reference sequence is depicted below the x-axis. Colours represent base type. N = any. **c** Percentage of positions in BB24 with indicated snFP percentage in the 'test data' prior to forward/reverse correction and **e** the difference after forward/reverse correction. Data points represent individual sequencing runs. 8 BB24 runs were used for the calculations. Error bars indicate the standard deviation (sd). snFP = single nucleotide false positive.
