## Supplementary Figure 6 for "Accurate detection of circulating tumor DNA using nanopore consensus sequencing"

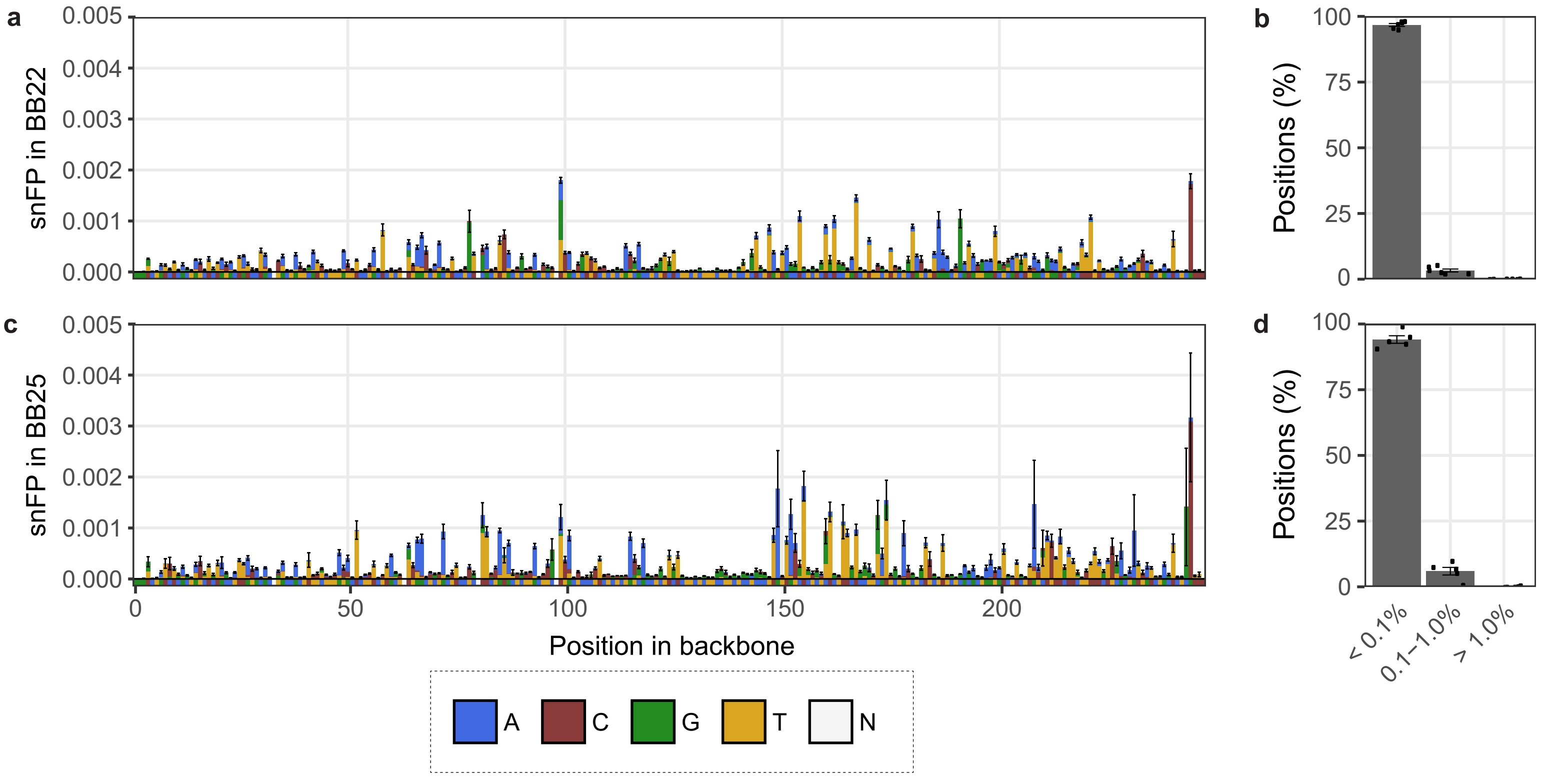


**Supplementary Fig. 6** Mean single-nucleotide false positive rate across BB22 (**a**) and BB25 (**c**) in reads with at least 10 repeats. Reference sequence is depicted below the x-axis. Colours represent base type. N = any. Percentage of positions in BB22 (**b**) and BB25 (**d**) with indicated snFP percentage. Data points represent individual sequencing runs. 6 BB22 and 5 BB25 runs were used for the calculations. Error bars indicate the standard deviation (sd).
