## Supplementary Figure 7 for "Accurate detection of circulating tumor DNA using nanopore consensus sequencing"

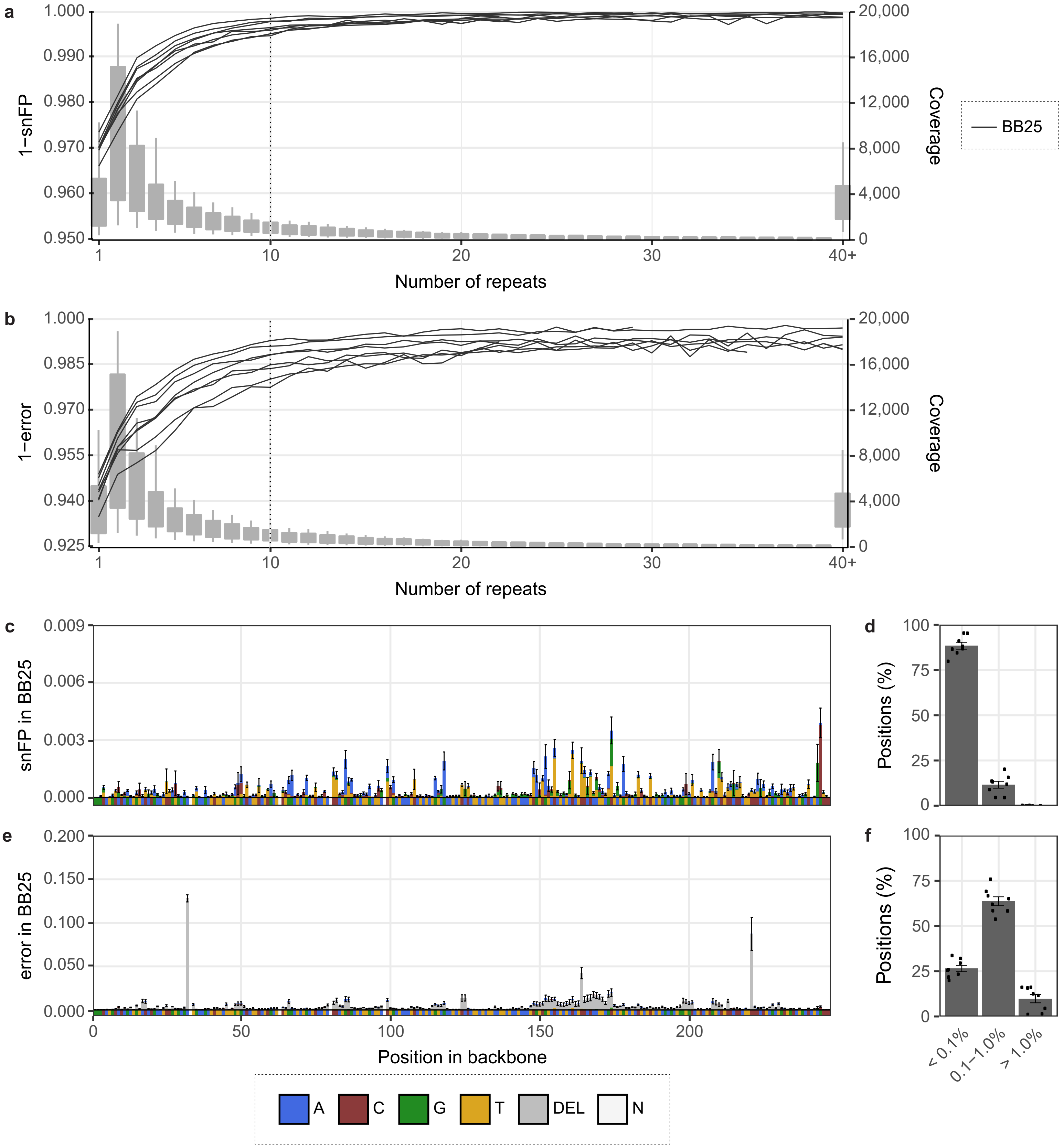


**Supplementary Fig. 7** Consensus calling lowers the number of sequencing errors in the backbone sequenced with Flongle. **a** 1 - single nucleotide false positive (snFP) and **b** 1- error (snFP + deletion) in backbone BB25 per number of repeats. The dashed line indicates 10 repeats. **c** Mean snFP and e error rate across BB25 in reads with at least 10 repeats. Reference sequence is depicted below the x-axis. Colours represent base type. N = any. **d** Percentage of positions in BB25 with indicated snFP and **f** error percentage. Data points represent individual sequencing runs. 8 BB25 runs were used for the calculations. Error bars indicate the standard deviation (sd).
