## Supplementary Figure 9 for "Accurate detection of circulating tumor DNA using nanopore consensus sequencing"

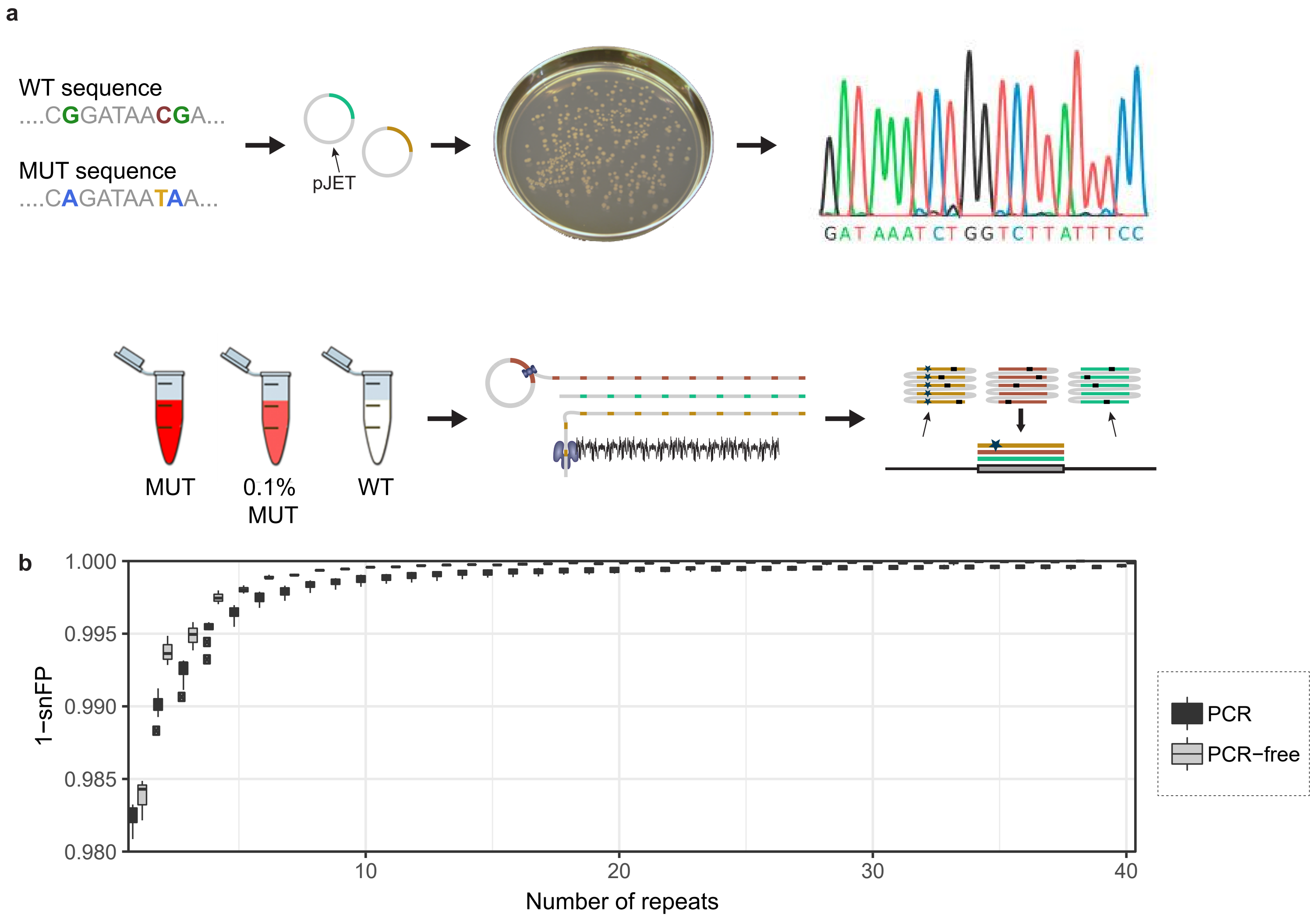


**Supplementary Fig. 9** Detecting mutations in a single synthetic *TP53* exon in pJET using CyclomicsSeq. **a** Experimental setup of the experiment. A synthetic WT sequence and MUT sequence (with three common COSMIC mutations in HNSCC) were cloned into pJET. Both sequences were amplified *in vivo* (PCR-free). Sanger sequencing was used to confirm the correctness of the WT and MUT sequence. CyclomicsSeq was performed on 100% WT, 100% MUT, and WT supplemented with 0.1% MUT. **b** Box plots (center line = median; box limits = 25th and 75th percentiles; whiskers = 1.5x interquartile range; data points = outliers) depicting 1 - single nucleotide false positive (snFP) rate in the insert (17:7577010-7577150 in GRCh37) per number of repeats for 8 PCR and 3 PCR-free inserts.
