## Supplementary Figure 10 for "Accurate detection of circulating tumor DNA using nanopore consensus sequencing"

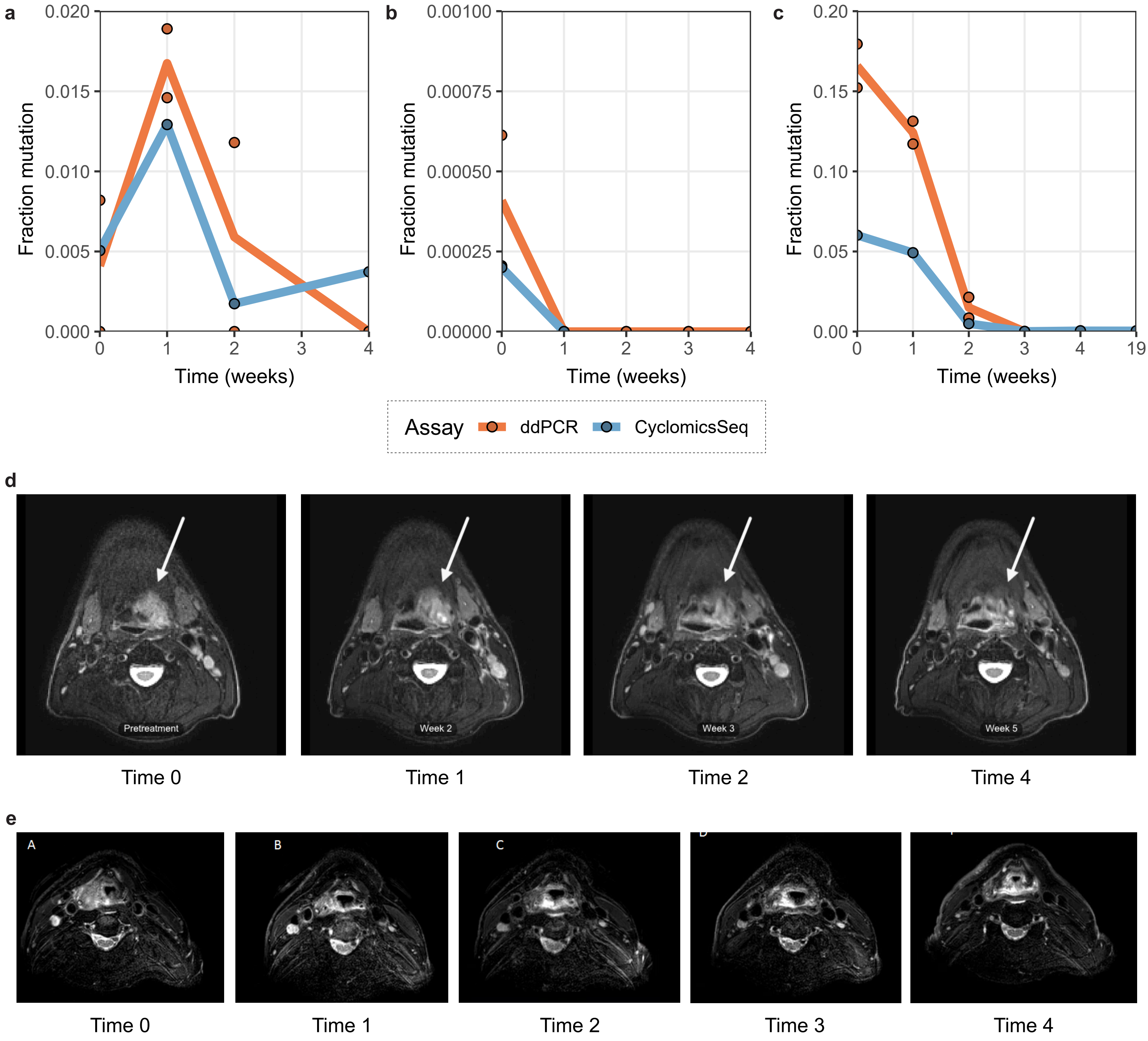


**Supplementary Fig. 10** ddPCR results and MRIs of patients. **a** 17:7577121 G>A in Patient A in ddPCR and CyclomicsSeq. **b** 17:7576870 C>A and 17:7577095-7577123 deletion in Patient B in ddPCR and CyclomicsSeq, respectively. **c** 17:7578403 C>T in Patient C in ddPCR and CyclomicsSeq. Each data point is a single measurement and lines show the mean measurement per time in weeks. **d** MRIs of patient A. White arrow indicates the primary tumor. **e** MRIs patient B. Time indicates the time in weeks after treatment initiation.
